# A machine-learning approach for accurate detection of copy-number variants from exome sequencing

**DOI:** 10.1101/460931

**Authors:** Vijay Kumar Pounraja, Gopal Jayakar, Matthew Jensen, Neil Kelkar, Santhosh Girirajan

## Abstract

Copy-number variants (CNVs) are a major cause of several genetic disorders, making their detection an essential component of genetic analysis pipelines. Current methods for detecting CNVs from exome sequencing data are limited by high false positive rates and low concordance due to the inherent biases of individual algorithms. To overcome these issues, calls generated by two or more algorithms are often intersected using Venn-diagram approaches to identify “high-confidence” CNVs. However, this approach is inadequate, as it misses potentially true calls that do not have consensus from multiple callers. Here, we present CN-Learn, a machine-learning framework (https://github.com/girirajanlab/CN_Learn) that integrates calls from multiple CNV detection algorithms and learns to accurately identify true CNVs using caller-specific and genomic features from a small subset of validated CNVs. Using CNVs predicted by four exome-based CNV callers (CANOES, CODEX, XHMM and CLAMMS) from 503 samples, we demonstrate that CN-Learn identifies true CNVs at higher precision (~90%) and recall (~85%) rates while maintaining robust performance even when trained with minimal data (~30 samples). CN-Learn recovers twice as many CNVs compared to individual callers or Venn diagram-based approaches, with features such as exome capture probe count, caller concordance and GC content providing the most discriminatory power. In fact, about 58% of all true CNVs recovered by CN-Learn were either singletons or calls that lacked support from at least one caller. Our study underscores the limitations of current approaches for CNV identification and provides an effective method that yields high-quality CNVs for application in clinical diagnostics.

## INTRODUCTION

Copy-number variants (CNVs) are a major source for genomic variation, evolution, and disease (Sebat et al. 2004; Redon et al. 2006; Perry et al. 2008; Girirajan et al. 2011). About 15% of affected individuals referred for clinical genetic testing carry a disease-associated CNV (Miller et al. 2010), making CNV detection an essential aspect of genetic analysis pipelines (Sathirapongsasuti et al. 2011; Krumm et al. 2012). Although the clinical utility of microarrays has not diminished (Coughlin et al. 2012), exome sequencing is becoming a prevalent technology for genetic testing (Yang et al. 2013; Stark et al. 2016; Tan et al. 2017; Retterer et al. 2016; de Ligt et al. 2013). Several algorithms are available to call CNVs from exome sequencing data in both clinical (Retterer et al. 2016) and disease-specific cohorts, including autism (Krumm et al. 2012), schizophrenia (Fromer et al. 2012), epilepsy (Epilepsy Phenome/Genome Project & Epi4K Consortium 2015), and cancer (Koboldt et al. 2012). A common strategy employed by these CNV callers is to apply various statistical distributions to model the aggregate read depth of the exons and use read depth fluctuations between adjacent exons to identify duplication or deletion events (Backenroth et al. 2014; Jiang et al. 2015; Packer et al. 2015; Krumm et al. 2012; Fromer et al. 2012).

Several themes have emerged due to variations in the approaches employed by different CNV callers to model read-depth distributions. *First*, the distributions and algorithms chosen to model read depth depend on the expertise of the researchers and their subjective assumptions about the underlying data. For example, callers such as XHMM (Fromer et al. 2012) assume the read-depth distribution to be Gaussian, while CANOES (Backenroth et al. 2014) assumes a negative binomial distribution, CODEX (Jiang et al. 2015) assumes a Poisson distribution, and CoNIFER (Krumm et al. 2012) makes no assumptions about the read-depth distribution. *Second*, while every method normalizes data to eliminate noise and outliers resulting from GC and repeat content biases, the number of samples required for normalization and the definition of outliers are inconsistent among the callers. For example, a principal component analysis (PCA) based method such as XHMM requires at least 50 unrelated samples for effective normalization, while CANOES only requires as low as 15 samples (Backenroth et al. 2014). Similarly, the annotations for “extreme” GC content differ among XHMM (<0.1 or >0.9), CODEX (<0.2 or >0.8), and CLAMMS (<0.3 or >0.7) (Packer et al. 2015). Further, XHMM only considers exome-capture targets between the size range of 10 bp and 10 kbp and with average coverage >10X across all samples, while CODEX uses targets that are >20 bp long and with median coverage >20X. CODEX eliminates targets with mappability scores <0.9, while CLAMMS eliminates regions with scores <0.75 along with a custom list of “blacklisted” regions (Packer et al. 2015). *Fourth*, the validation methods and the subsets of CNV calls used to estimate sensitivity and specificity measures vary widely among the callers. For example, large CNVs (>100 kbp) validated with microarrays and analyzed with PennCNV (Wang et al. 2007) were used to estimate the performance of CANOES (Backenroth et al. 2014), while only a subset of CLAMMS calls with minor allele frequency <0.1 were validated using PCR. Also, not all callers report confidence scores for the CNVs they identify. Even when they do, their confidence scales are not directly comparable. These issues influence the number and type of CNVs detected by each caller in a given sample, resulting in pronounced differences in accuracy, false positive rates, and concordance among the callers (Hong et al. 2016; Yao et al. 2017). *Finally*, studies using exome-sequencing data to detect CNVs either utilize predictions made by a single caller (Krumm et al. 2013; Poultney et al. 2013) or use a Venn-diagram approach to identify calls with concordance among multiple callers as “high-confidence” CNVs (Kataoka et al. 2016; Priest et al. 2016; Bademci et al. 2016; Krumm et al. 2015). While using data from multiple callers minimizes false positive rates, this approach discards a large subset of non-concordant CNVs, thereby reducing the overall CNV yield (Hong et al. 2016). In addition to a low CNV yield, the reported breakpoints of the concordant calls do not necessarily agree between callers. These limitations associated with individual CNV callers as well as the methods used to integrate predictions from different CNV callers necessitate a better approach to identify and prioritize clinically relevant CNVs.

Here, we propose a machine-learning method called CN-Learn that overcomes the limitations of high false positive and low concordance rates among calls generated by different CNV algorithms to identify high-confidence CNVs. Our method leverages several attributes intrinsic to each CNV call, such as GC content, mappability of the genomic region, and CNV size, in addition to concordance among the callers. CN-Learn learns the associations between these attributes and the presence or absence of CNVs using a small subset of validated CNVs in the cohort (known truth), and then segregates true CNVs from false positives in the test samples with high precision. Using exome-sequencing data and validations from 503 samples, we demonstrate CN-Learn’s ability to recover more than twice as many potentially true variants compared to a Venn-diagram approach. Our study reiterates the limitations of existing CNV detection and integration methods, and offers a better alternative that yields a set of high-quality CNVs for application in clinical diagnostics.

## RESULTS

We developed CN-Learn as a binary Random Forest classifier that can be trained to differentiate true CNV predictions from false positives using a small subset of validated CNVs (**Figure 1**). We identified twelve features that represented the extent of support from individual algorithms and the genomic context for each CNV (see Methods). We detected statistically significant correlations for several pairs of the quantitative features (**Figure 2; Supplemental Table 1**). Based on a collection of decision trees built using the twelve features extracted from CNVs in the training samples, CN-Learn estimates the probability of each CNV in the test sample to be true (**Figure 2**). In addition to a Random Forest classifier, CN-Learn can also be built as a Logistic Regression (LR) or Support Vector Machine (SVM) based classifier (**Supplemental file**).

**Figure 1:**
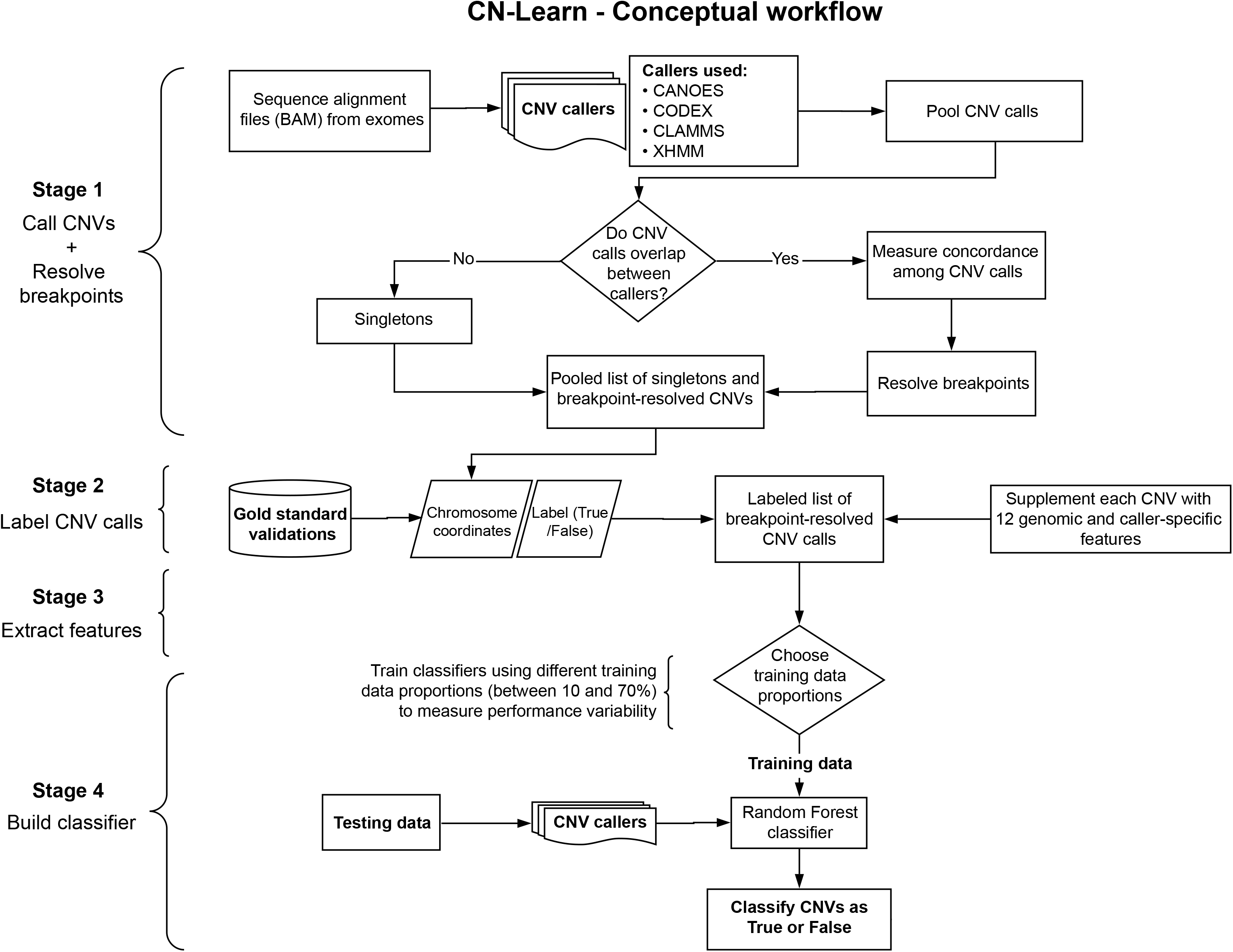
Overview of the CN-Learn pipeline. The CN-Learn pipeline consists of pre-processing steps (Step 1 and 2), followed by building the classifier using training data and discriminatory features, and finally running the classifier on the test data. The complete pipeline is outlined as follows. ***Step 1***: CNV predictions were made using four exome-based CNV callers. While CANOES, CODEX, CLAMMS and XHMM were used in this study, a generic pipeline can be constructed with a different set or number of callers. Breakpoints of overlapping calls from multiple callers were then resolved. ***Step 2:*** Breakpoint-resolved CNVs were labeled as “True” or “False” based on the overlap with “gold standard” calls and subsequently used to train CN-Learn. ***Step 3:*** Caller-specific and genomic features were extracted for the labeled CNVs in the training and testing set. ***Step 4:*** CN-Learn was trained as a Random Forest classifier using the extracted features of the CNVs in the training set to make predictions on the CNVs from the testing set.

**Figure 2:**
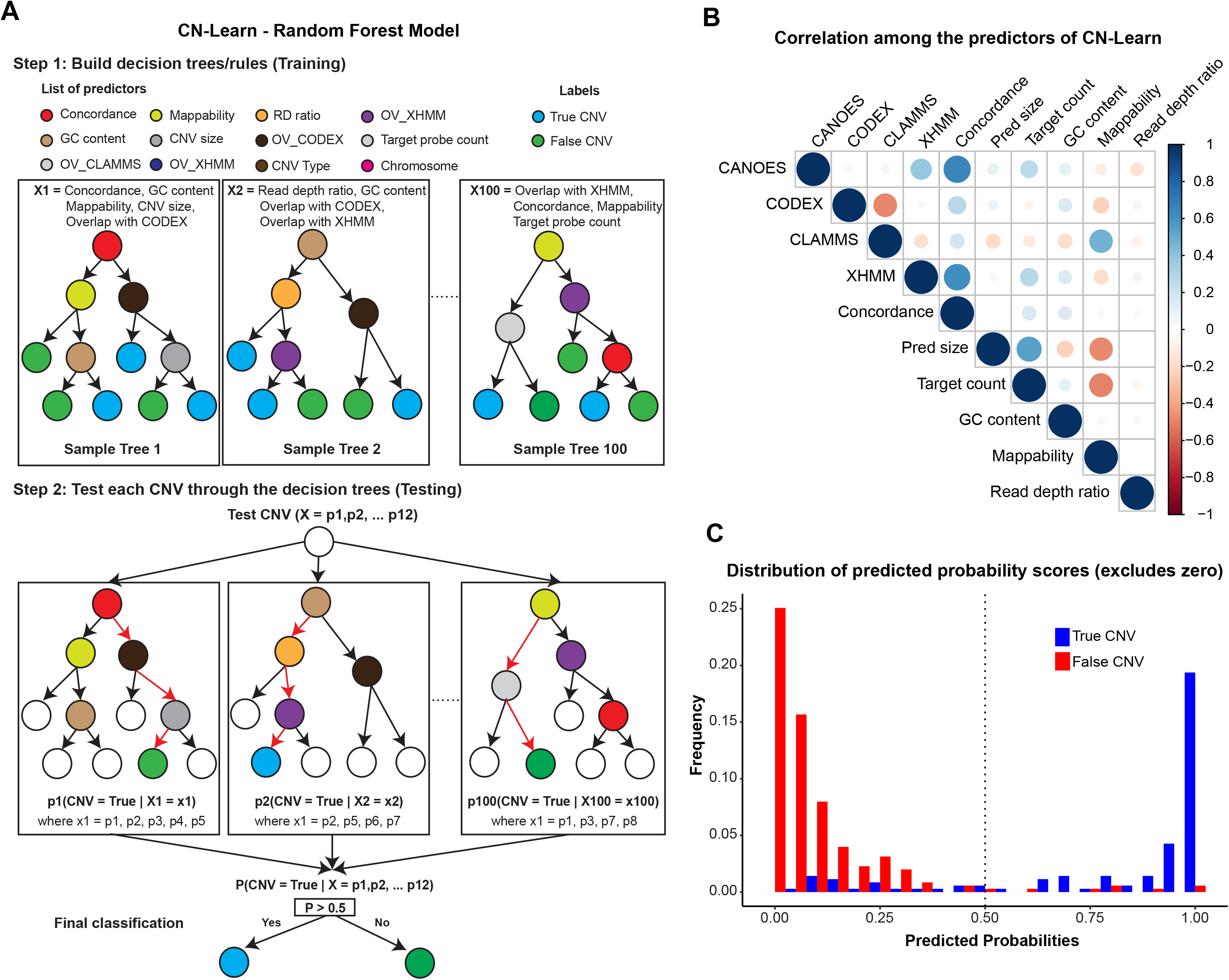
Illustration of the Random Forest model used to build CN-Learn. **(A)** The inner workings of the random forest model used for training CN-Learn is shown. Twelve features were used to grow 100 trees with different subsamples of predictors and training data to classify each CNV in the test set as either true or false. If the predicted probability score was greater than 0.5, the CNV call was classified as true. Calls with predicted probability score <0.5 were labeled as false. **(B)** A Spearman rank correlation between pairs of quantitative predictors used by the CN-Learn classifier is shown. The color of the circles indicates the direction of the correlation, while the size of the circles indicates the strength of the correlation. The correlation scores are provided in the supplemental file (**Supplemental Table 1**). **(C)** The frequency of microarray validated and invalidated CNVs, distributed across 20 bins of increasing predicted probability scores, is shown. For the probability bins less than 0.5, the proportion of CNVs that were validated was higher than the proportion of CNVs that was not validated. This indicated that classification score of 0.5 is an appropriate threshold for distinguishing true and false CNVs.

### CN-Learn detects high-confidence CNVs with high precision and recall rates

To build the CN-Learn classifier, we first identified 41,791 CNV predictions from 503 samples using four exome-based CNV callers (CANOES, Codex, CLAMMS, and XHMM). Using a read-depth based method (**Supplemental Fig. S1**) to resolve breakpoint conflicts of overlapping CNV predictions obtained from different callers, we identified 29,101 unique CNV events among the 503 samples (**Supplemental file**). An alternate approach to resolve breakpoint conflicts (**Supplemental Fig. S2**) also provided the same number of unique CNV events. We selected 2,506 of these CNVs from 291 samples with microarray validations that were between 50 kbp and 5 Mbp and spanned regions covered by microarray probes (**Supplemental Fig. S3**). After determining the proportion of CNVs that overlapped with microarray validations at different thresholds (**Supplemental Table 2**), we labeled each of the selected CNVs as either “True” or “False” based on a 10% reciprocal overlap threshold. We next used CNVs from 70% of the 291 samples to train CN-Learn as a Random Forest classifier and the remaining 30% of samples to test its performance. Given the uneven distribution of the labels between the two CNV classes (11% true vs. 89% false), we chose precision and recall as the measures of classifier performance (**Supplemental Table 2**). We captured the aggregate performance of CN-Learn using ten random draws of training data (10-fold cross validation), stratified by sample. CN-Learn used the twelve predictors supplied with each CNV in the test set (see Methods) and classified the calls as either true or false at 91% precision and 86% recall rates. The overall diagnostic ability of the binary classifier, measured as the area under the Receiver Operating Characteristic (ROC) curve, was 95% (**Figure 3A**). Despite sampling different sets of training data for each iteration during cross-validation, the performance of CN-Learn was consistent across all ten iterations for both precision (91 ±5%) and recall rates (86±5%) (**Figure 3B**).

**Figure 3:**
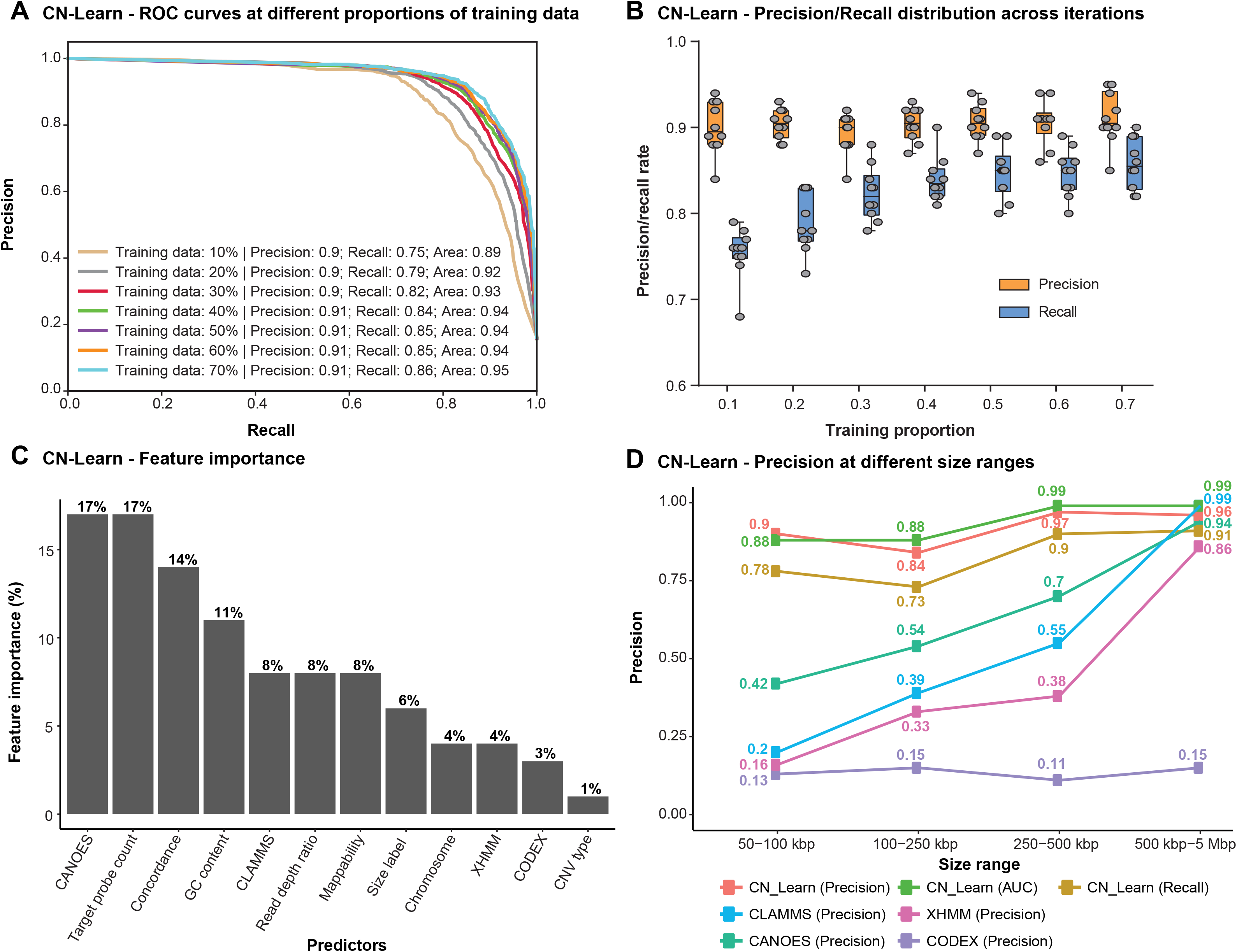
Characteristics of the CN-Learn binary Random Forest classifier. **(A)** Receiver operating characteristic (ROC) curves indicating the trade-off between the precision and recall rates when CN-Learn was trained as a Random Forest classifier are shown. Each curve represents the performance achieved when using different proportions of samples to train CN-Learn, starting from 10% up to 70% in increments of 10%. The results shown were from experiments aggregated across 10-fold cross validation. **(B)** Variability observed in the precision and recall measures during the 10-fold cross validation at various proportions of training data is shown. Both measures varied within ±5% of their corresponding averages. **(C)** The relative importance of each genomic and caller-specific feature supplemented to CN-Learn is shown. Data shown here are the averages of the values obtained across 10-fold cross-validation after using 70% of the samples for training. **(D)** Precision rates for CNVs when CN-Learn was trained at four different size ranges compared to the precision rates of CNVs from individual callers are shown. Precision rates for CN-Learn were estimated as its classification accuracy (True positives/ [True positives + False positives]), while the precision rates for the individual callers were calculated as the proportion of CNVs at each size range that were validated by the microarray calls.

We also assessed the relative importance of each feature towards making accurate CNV predictions by calculating the Gini index (defined as the total decrease in improvements to the node purity for all splits on each feature averaged over all trees in the forest) (Breiman 2001). We found that features such as the number of exome capture probes spanning a given call, the extent to which CANOES agreed with a CNV prediction, concordance among the callers, and GC content provided the most discriminatory power to the classifier (**Figure 3C**). Post-classification analysis of the concordance profile indicated that only 34% of all CNVs classified as true had support from all the four callers, while the remaining 66% lacked support from at least one caller (**Supplemental Fig. S4**). We further assessed the performance of CN-Learn on an independent set of 90 samples from the 1000 Genomes project (Jiang et al. 2015), and observed a precision rate of 93% and recall rate of 86% when using 70% training data (**Supplemental Fig. S5-S6**). When we generated precision/recall rates that are directly comparable with the concordance-based methods, we observed a consistently superior performance for CN-Learn (**Supplemental File, Supplemental Tables 3-4, Supplemental Fig. S7**). Overall, these results highlight the ability of CN-Learn to look beyond the single measure of concordance typically used in a Venn-diagram based approach, and to utilize the discriminatory power of additional variables to identify high-confidence CNVs in a systematic manner.

### Performance of CN-Learn is robust across varying CNV sizes, frequencies and training sets

We independently trained CN-Learn using varying proportions of training data (between 10% and 70% in increments of 10%) and observed steady performance gains with increase in the number of training samples (**Figure 3A**). Interestingly, even when the classifier was built using just 10% of the total samples (n=29 samples), we obtained 90% precision and a recall rate of 75%, indicating the robustness of the classifier when learning from minimal training data. We further trained CN-Learn independently at four size ranges of CNVs and observed a modest increase in precision with increase in CNV size (90% for 50-100 kbp CNV to 97% for 0.5-5 Mbp CNV) (**Figure 3D**). In fact, the precision achieved by CN-Learn at each size interval was substantially higher than the precision achieved by the individual CNV callers. We also observed comparable precision and recall rates when CN-Learn was run on breakpoint-resolved CNVs obtained by merging overlapping predictions (**Supplemental Fig. S8**). We then tested the performance of CN-Learn for different classes of CNVs based on their frequency in the cohort (very rare, rare, common, very common), and found precision (> 90%) and recall (> 80%) rates that were consistent across the CNV frequency spectrum (**Supplemental File; Supplemental Fig. S9**). In fact, when added as an additional predictor to CN-Learn, CNV frequency showed the highest discriminatory power (16%) of any predictor, but did not contribute to significant improvements in the performance of the classifier (**Supplemental Fig. S10, S11**). While the performance of the Random Forest classifier was robust, Logistic Regression and SVM classifiers failed to match its performance (**Supplemental File; Supplemental Figs. S12, S13**).

We next assessed the performance of CN-Learn by considering calls made by CLAMMS as the truth set and classified CNVs obtained from predictions made by the other three callers (CANOES, CODEX and XHMM). We categorized 25,019 breakpoint-resolved CNVs (**Supplemental Fig. S3**) from all 503 samples as either “True” or “False” based on their intersection (10% reciprocal overlap) with CNVs predicted by CLAMMS. Given the higher resolution of CNVs detected from exome data relative to SNP microarrays, we used a total of 16,497 CNVs between 5 kbp and 5 Mbp in size to build CN-Learn (**Supplemental Fig. S3**). CN-Learn achieved an aggregate precision rate of 94% with an overall recall rate of 85% during the 10-fold cross validation, and achieved comparable performance when independently trained with CNVs at different training sample proportions (**Supplemental Fig. S14A**). Performance variability observed both during crossvalidation and across the size ranges were comparable to the variability observed when microarray was used as the truth set (**Supplemental Figs. S14B-C**). Among the features used by CN-Learn to classify the CNVs, the relative importance of the mappability score was the highest, with GC content being the next important feature (**Supplemental Fig. S14C**). While caller-specific features contributed to the discriminatory power of CNV classification when microarray was used for validation, genomic features played a more prominent role when a sequence-based method was used for validation (**Supplemental Fig. S15**). These results show that the performance of CN-Learn is robust with minimal training data, at different size ranges and even when orthogonal validations are not available.

### CN-Learn recovers true CNVs that lack complete concordance among callers

To assess the ability of CN-Learn to correctly identify true CNVs that lack support from multiple callers, we analyzed the concordance profile of all CNVs classified by CN-Learn as true, based on microarray validations, before and after CN-Learn classification. Using a single random draw of 29 samples (10%) to train CN-Learn and 262 samples as the test set, we obtained predictions for 2,245 CNVs with microarray validations (**Figure 4A**). Among these predictions, only 122/2,245 CNVs (5.4%) were supported by all four CNV callers prior to classification by CN-Learn. The strong concordance of the four methods for these CNV predictions was corroborated by a high microarray validation rate (116/122, 95%) (**Figure 4A**). In contrast, CNVs that lacked support from one or more callers were less likely to intersect with microarray data. For example, only 41% (11/27) of the CNVs with support from XHMM, CODEX and CLAMMS intersected with microarray data. After classification by CN-Learn, 84% (266/315) of all CNVs labeled as “True” intersected with microarray calls, indicating the high classification accuracy achieved by the classifier (**Figure 4A**). All CNVs classified as true and false from this analysis are provided in **Supplemental Tables 5 and 6**. Of the 266 CNVs validated by microarrays and correctly identified as true by the classifier, only 42% (112/266) were supported by all four CNV callers. The remaining 58% (154/266) were either singletons or calls that lacked support from at least one caller, and would have been excluded if complete concordance was used as the only determinant for selecting high confidence CNVs. For example, 13% of all true CNVs (35/266) recovered by CN-Learn were missed by CLAMMS but were identified by one or more of the other callers, reiterating the limitations of using a single exome-based CNV caller for variant predictions. Further, CN-Learn managed to recover 97% (112/116) of the true CNVs validated by microarrays that were supported by all of the four callers. Although these 112 CNVs could have been identified by a simple caller intersection approach, CN-Learn was uniquely able to recover CNVs that lacked support from at least one other caller. For example, CN-Learn classified 66 CNVs supported by CANOES, CODEX and XHMM as true, of which 51% (34/66) were also validated by microarrays. This result is notable because, without using CN-Learn for classification, only 31% (46/148) of CNVs supported by CANOES, CODEX and XHMM would intersect with the microarray calls, indicating the inherently high false positive rate associated with simply intersecting calls from individual callers using a Venn diagram. Our results indicate that in addition to correctly identifying almost every true CNV reported by the four callers, CN-Learn overcame the limitations of the Venn-diagram based approach and recovered 154 additional high-confidence CNVs with sub-optimal concordance, thereby improving the CNV yield by 2.37-fold (266/112). Further, the true positive calls recovered by CN-Learn spanned the entire spectrum of CNV size range (**Figure 4B**). We also obtained comparable improvements in CNV yield when CNVs predicted by CLAMMS were used as the truth set (**Supplemental file; Supplemental Figs. S16-S17, Supplemental Tables 5 and 6**).

**Figure 4:**
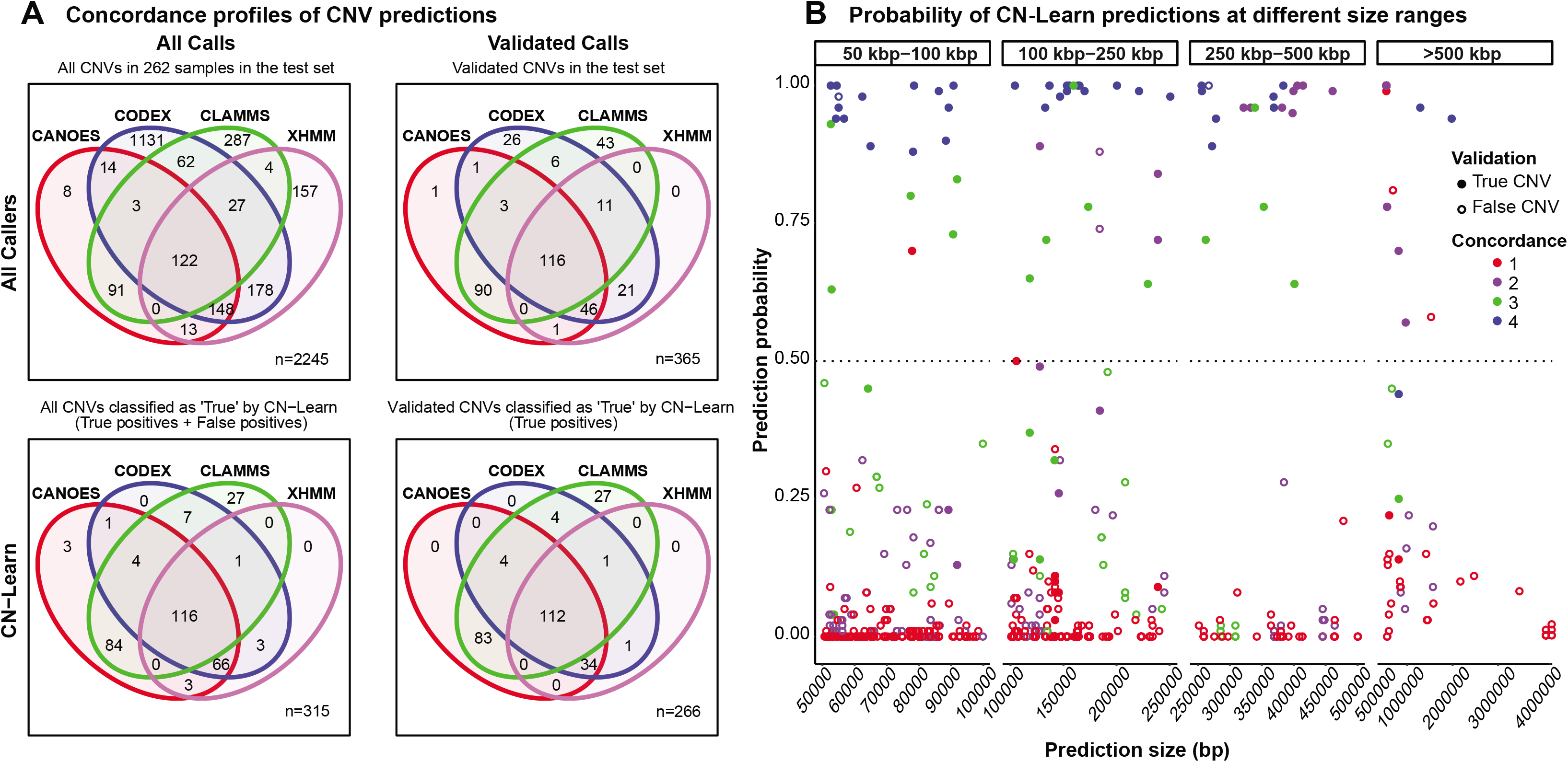
Concordance profile of CNVs before and after classification by CN-Learn using microarray calls as the “gold standard” validation set. **(A)** Venn diagrams are shown for CNVs (≥50 kbp) identified from a random draw of 262 samples out of the total 291 samples before (**Top panels**) and after (**Bottom panels**) classification by CN-Learn. The Venn diagrams show the overlap of calls among the four callers (**Top left**) versus those that were validated by microarrays (**Top right**). Venn diagrams depicting the overlap of all true calls among the four callers after classification by CN-Learn (**Bottom left**) and true calls that were also validated by microarrays are shown (**Bottom right**). (**B**) The distribution of all calls within the 262 samples based on the probability scores (Y-axis) predicted by CN-Learn across four size ranges (X-axis) is shown. The number of overlaps of each CNV with exome-based CNV callers is represented by different colors. CNVs that validated with microarrays are indicated by filled circles, while CNVs that did not validate with microarrays are represented by hollow circles.

## DISCUSSION

Exome sequencing is a cost-effective assay to accurately identify single base pair mutations and small insertions or deletions within protein-coding genes, which are more likely to cause disease than mutations in other regions of the genome (Bamshad et al. 2011). Even with the emergence of whole genome sequencing (WGS) techniques, exome sequencing has continued to flourish, resulting in an increase in the availability of datasets for analysis. This widespread data availability has allowed for repurposing data generated for single nucleotide variant detection to identify CNVs as well (Koboldt et al. 2013). Given the high false positive rates among calls reported by individual exome-based callers and platform-specific biases (Tan et al. 2014), existing CNV identification pipelines that leverage more than one calling algorithm have depended on naïve Venn diagram-based integration approaches to identify high-confidence CNVs. One reason for the overwhelming trust in such approaches could be the fact that the self-reported performance measures of each individual algorithm are typically high. Therefore, there is little reason to doubt that performance variations among the callers could affect the precision of Venn diagram-based integration approaches. While we found that the likelihood of a CNV prediction to be true increased with an increase in the number of callers supporting it (Supplemental Fig. S18), there are two key limitations of this approach. *First*, even among completely concordant predictions, the observed false positive rates were not zero. Inflated false positives pose a large hurdle for researchers interested in identifying clinically relevant CNVs, as it is time consuming to validate a large number of false positive CNVs using orthogonal methods. *Second*, the Venn diagram approach failed to identify a large subset of non-concordant or singleton CNVs supported by microarray validations. In fact, approximately one true CNV (>50 kbp) per individual in our cohort would have been discarded under a Venn-diagram based approach. This evidence reiterates that Venn diagram-based approaches do not have the required precision for usage in both clinical and research settings.

The utility of alternate methods for CNV detection hence rests on the ability to both eliminate false positives among completely concordant predictions and recover true CNVs that lack adequate support from multiple callers. Therefore, instead of addressing the shortcomings of existing methods by developing yet another CNV detection tool, our study focused on offering a reliable integrative approach. In this study, we demonstrate a machine-learning approach that leverages caller-specific and genomic contexts from a subset of validated calls to identify high-confidence CNVs more thoroughly than individual callers on their own or Venn diagram-based approaches. CN-Learn achieved precision as high as 94% while doubling the CNV yield, showing its ability to capture singletons that would have been missed by other approaches. Importantly, the precision of CN-Learn is robust to variation in CNV size, ratio of training to testing samples, and validation method/type, indicating the utility of CN-Learn in a variety of clinical or research contexts. The use of multiple variables that capture the genomic context of each CNV in addition to caller concordance is a major reason for the high recovery and precision rates achieved by CN-Learn. In fact, GC content and mappability scores were some of the most useful features in predicting true CNVs.

One of the limitations of CN-Learn is its dependency on a small set of biologically validated CNVs. Our study leverages microarray validations to evaluate the predictions of each caller and to label the breakpoint-resolved CNV predictions as either true or false. While microarrays may not be considered as a “gold standard” for CNV detection, we were able to use microarray calls as an orthogonal validation to demonstrate the utility of CN-Learn. Our study also serves as a proof-of-principle for future studies that could utilize CN-Learn with “gold-standard” CNVs curated from multiple genomic technologies, such as PacBio SMRT sequencing (John et al. 2009), Illumina long-read sequencing (Voskoboynik et al. 2013), 10X linked-read sequencing (Zheng et al. 2016), BioNano Genomics genome mapping (Lam et al. 2012; Mak et al. 2016) or PCR. Another limitation of our study is the time and computational capacity required to run four different CNV callers. Each CNV calling algorithm extracts read-depth information for each sample using tools such as Samtools (Li et al. 2009), Bedtools (Quinlan and Hall 2010) or Genome Analysis Toolkit (GATK) (McKenna et al. 2010), which are often the rate-limiting steps for each pipeline. Future studies could simplify this data extraction layer by using a single read-depth tool without adversely impacting the results of the individual callers. Finally, we also acknowledge the complexity associated with resolving breakpoint conflicts from multiple callers that arise during data integration. While we presented two strategies to resolve breakpoints of concordant CNVs (see Methods), future studies could explore more effective strategies before using CN-Learn. As population-scale projects continue to generate large exome-sequencing datasets, the need and importance of robust CNV integration approaches such as CN-Learn is apparent.

Overall, CN-Learn integrates predictions from multiple CNV callers and overcomes the limitations of existing integration approaches, even when the availability of samples with biological validations is limited. Although we chose a set of four CNV calling algorithms with microarray validations, CN-Learn framework can be extended to use different sets of CNV callers or validation types to identify high-confidence CNVs, making our framework easy to adopt and customize. This can allow clinicians and researchers to use their preferred callers and validation methods to detect CNVs from exomes. Our results suggest that a small set of high-quality validated CNVs and an objective machine learning method can help alleviate several shortcomings of existing integration approaches to generate an informed set of clinically relevant CNVs.

## METHODS

### Samples

We obtained exome sequencing data from 503 individuals in the Simons Variation in Individual Project (SVIP) which were generated using the Agilent Sure-Select Human All Exon v2.0 capture kit containing 182,430 autosomal probes (Spiro and Chung 2012). Overall, 187 individuals within this cohort carry the 16p11.2 deletion, and 143 individuals carry the reciprocal duplication. Single nucleotide polymorphism (SNP) based microarray data were available for 291 samples, and CNVs >100 kbp were confirmed experimentally using array CGH (Duyzend et al. 2016). We also used 90 samples from the 1000 Genomes project (Durbin et al. 2010), of which 46 samples were sequenced at the Washington University Genome Sequencing Center using the Agilent SureSelect All Exon v2.0 capture kit and 44 samples were sequenced at Baylor College of Medicine using the Roche HGSC VCRome capture kit. We used 186,065 autosomal probes from the consensus probe set (ftp://ftp.1000genomes.ebi.ac.uk/vol1/ftp/technical/reference/exome_pull_down_targets/20130108.exome.targets.bed) to make CNV predictions.

### Exome CNV callers

We chose four exome CNV callers to obtain the initial sets of CNV calls: CANOES, CLAMMS (v1.1), CODEX (v0.2.2), and XHMM (Backenroth et al. 2014; Packer et al. 2015; Jiang et al. 2015; Fromer et al. 2012). As the SVIP samples were sequenced in two sets (312 and 191 samples), each set of samples was treated as individual batches for running the CNV-calling pipelines. CANOES and CODEX were run using the default parameters. CLAMMS models were built with the assumption that the samples were independent without accounting for batch effects. All of the parameters used to make CNV calls using the four callers for this study can be found in the pipelines provided as part of the CN-Learn distribution (**Supplemental_Code.zip**, https://github.com/girirajanlab/CN_Learn). Running the four CNV callers yielded 41,791 calls of varying sizes (**Supplemental Fig. S19**). These original CNV calls (deletions and duplications) were then characterized based on their level of concordance among the four callers (**Supplemental file**).

### Resolving breakpoint conflicts for overlapping predictions

Multiple CNV callers can make predictions that overlap with each other in a given genomic region. Treating such overlapping predictions with different breakpoints as separate CNVs would result in double-counting the calls for the same CNV event. Therefore, it is important to merge concordant predictions and represent them as a single event for downstream analyses. We developed a five-step procedure that uses fluctuations in local read depth to resolve breakpoint conflicts among overlapping predictions by identifying the most likely start and end coordinates of the underlying event. A detailed explanation of this strategy is presented in the **Supplemental file** (**Supplemental Figs. S1 & S20**). As an alternate strategy, we resolved breakpoint conflicts by simply selecting the smallest and largest coordinates among the overlapping predictions as the start and end coordinates of the underlying CNV event. This strategy is also described in the Supplemental file (**Supplemental Fig. S2**). Applying both methods to the 41,791 CNVs detected in the SVIP samples yielded 8,382 unique breakpoint-resolved CNV events, in addition to the 20,719 singleton calls (**Supplemental Fig. S3**).

### CNV validations

SNP microarray data were generated by the SVIP consortium (Duyzend et al. 2016) using Illumina OmniExpress-12 microarrays for 272 samples and OmniExpress-14 for 19 samples. PennCNV (v.1.0.3) was used to identify CNVs from microarray data for all 291 individuals using standard parameters (Wang et al. 2007). Individual and family-based (trios and quads) CNV calls were combined for autosomal chromosomes, while CNVs on chromosome X were called only at the individual level. CNV calls with ≥1 bp overlap or gaps <20% of the total CNV length and <50 kbp were merged. CNVs ≥50 kbp in length and containing ≥5 SNP target probes were subsequently considered for further analyses.

For the 1000 Genomes samples, we pooled the validated calls from three sources (Altshuler et al. 2010; Conrad et al. 2012; McCarroll et al. 2008) that were used to measure the performance of CODEX (Jiang et al. 2015). Merging the overlaps among the 10,235 CNVs resulted in 7,302 CNVs (**Supplemental Table 4**) that were used to label the CNVs predicted for the 90 samples from the 1000 Genomes project.

### Feature selection for CN-Learn classifier

We identified twelve features that represent the extent of support provided by the four individual callers and genomic context as predictors of true CNVs to the CN-Learn Random Forest classifier. Since we used four callers in our study, the extent of overlap calculated during the breakpoint-resolution process (**Supplemental file & Supplemental Fig. S1**) served as individual predictors. Concordance count and read-depth ratio (RD_ratio_) for both breakpoint-resolved concordant calls and singletons were also supplied as features. As individual algorithms use different GC and repeat content (mappability) thresholds to classify CNV predictions as outliers, CNVs with extreme GC content or low mappability could be predicted by one caller but discarded as an outlier by the other callers. To take this into account, we extracted GC content data using the “nuc” option in bedtools (Quinlan and Hall 2010) and mappability scores (Derrien et al. 2012) using the “bigWigAverageOverBed” option in kentUtils (Kent et al. 2010) for use as predictors. Similarly, the efficacy of CNV detection can vary across chromosomes, size ranges, and CNV type (duplication/deletion). To take these variations into account, we used chromosome number, CNV size, CNV type and the number of exome capture probes as the final set of CN-Learn features.

### Probability estimation and classification using CN-Learn

CN-Learn leverages the features extracted for the CNVs in the training samples to build a Random Forest (Breiman 2001) classifier with hundreds of decision trees and estimates the probability of each CNV in the test samples to be true (**Figure 2A**). Decision trees have been shown to perform well when the distribution of observations is unbalanced between the classes of interest (Cieslak and Chawla 2008). Given the high false positive rates of CNV callers, an uneven split between the number of true and false predictions is likely to occur in clinical samples. In the samples we analyzed, 10% of the CNVs overlapped with the truth sets derived from microarray calls at a 10% reciprocal overlap threshold (Supplemental Table 2). To address the imbalance between the number of true and false CNVs, we stratified the training data by sample to accurately reflect the clinical setting as follows: CNVs in p% of the samples were used for training, and the remaining (1-p)% were used as the testing set. For a random forest built with “N” trees, “M” predictors and “C” classes, the probability of an observation “o” belonging to the class “c” (true or false) can be expressed as p_*o,c*_ = Pr(Y = c | X = x_*i*_), where x_*i*_ is a vector that captures the values for each of the 12 predictors and Y is the outcome variable. The probability p_o,true_ of the CNV “o” being true in the test set was then measured as the proportion of trees in the forest that assigned it to the true class. Specifically, the probability of a CNV prediction can be represented as:

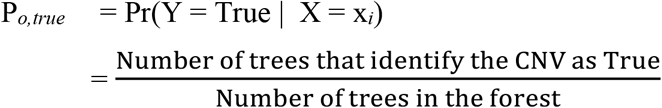

where X represents the values of the following features for the observation “o”:

x_1_ = Overlap proportion with CANOES
x_2_ = Overlap proportion with CODEX
x_3_ = Overlap proportion with CLAMMS
x_4_ = Overlap proportion with XHMM
x_5_ = Concordance among callers
x_6_ = Read depth ratio
x_7_ = Chromosome number
x_8_ = CNV type (Duplication/Deletion)
x_9_ = CNV size
x_10_ = Target probe count
x_11_ = GC content
x_12_ = Mappability

CNV calls with predicted probability score >0.5 were then classified as true. We selected this cutoff based on the distribution of validated (true) and invalidated (false) CNVs across the predicted probability scores (ranging between 0 and 1). For both duplications and deletions, the proportion of true calls compared to false calls was higher for CNVs with probability scores >0.5 (**Figure 2C**). This indicated that at a 0.5 threshold, the classifier recovers as many true CNVs (recall) as possible without compromising on the false positive rate (precision).

### Statistical analysis

All statistical analyses, including the calculation of precision-recall rates, feature importance, and ROC areas, were performed using the Python library scikit-learn (v 0.18.1) (Pedregosa et al. 2012). Plots were generated using the R package ggplot2 and the Python library matplotlib.

### Code availability

CN-Learn is available as an open-source software at *https://github.com/girirajanlab/CN_Learn* and provided as a supplemental file (**Supplemental_Code.zip**). In addition to the scripts necessary to run CN-Learn, we also provide simplified and easily parallelizable pipelines for each of the original CNV calling algorithms (CANOES, CLAMMS, CODEX and XHMM) used in this study.

Furthermore, in order to simplify installation and avoid software version incompatibilities, we provide a Docker image with all the required tools and software packages preinstalled to run CN-Learn. Users can download the image using the command ‘ *docker pull girimjanlab/cnlearri*’ on any Linux platform to run CN-Learn. Detailed instructions to install and use Docker and CN-Learn are provided in the README.md file in the supplemental file (**Supplemental_Code.zip**) and at the landing page for the software on GitHub.

## DECLARATIONS

### Ethics approval and consent to participate

As these data were de-identified, all of our samples were exempt from IRB review and conformed to the Helsinki Declaration. No other approvals were needed for the study.

### Consent for publication

All authors agree and consent for publication of the manuscript.

### Availability of data and materials

BAM files for all samples, aligned to the hg19 reference genome, were obtained from the Simons Foundation Autism Research Initiative (SFARI) via SFARI Base following appropriate approvals (*https://www.sfari.org/resource/sfari-base/*) and from the ftp site of the 1000 Genomes data collection (ftp://ftp.1000genomes.ebi.ac.uk/vol1/ftp/phase3/data). Realigning the reads to hg38 (GRCh38) would not significantly affect the conclusions of the manuscript, as the CNV predictions are restricted to sequences captured by the exome capture probes. The script used to generate the plots are provided in the supplemental file (Supplemental_Code.zip) and in the “research” directory of CN-Learn repository. Any intermediate datasets generated during this study are available from the corresponding author upon request.

### Authors’ contributions

VK and SG conceived the project. VK performed the analyses, generated the plots/images, and wrote and revised the manuscript; MJ organized the images for publication and assisted with revision to the manuscript; GJ and NK collected data and performed parts of the analyses; SG supervised the research and wrote and revised the manuscript. All authors read and approved the final draft of the manuscript.

## Acknowledgements

We thank Naomi Altman (Penn State), Dajiang Liu (Penn State), Arjun Krishnan (Michigan State), Cooduvalli Shashikant (Penn State), Lucilla Pizzo (Penn State), and Shaun Mahony (Penn State) for constructive comments on the manuscript.

