## Supplemental File for "A machine-learning approach for accurate detection of copy-number variants from exome sequencing"

**Supplemental Figure S1: Illustration of the read-depth based breakpoint resolution method.** ... 14

### DEFINITIONS

**Caller intersection:** A strategy used to assess overlap between CNVs predicted by multiple CNV calling methods. Caller intersection strategy is often used to prioritize high-confidence calls identified by multiple callers.

**CNV:** Copy-number variants, or large deletions or duplications in the genome.

**CNV Calls:** Predictions for the likely existence of CNVs in a given genomic region.

**Concordance:** A term for agreement among callers for a given CNV event. In our study, calls made by two callers were considered concordant when the overlap between their coordinates was >1%. We use the term "complete concordance" when all tested callers predict the same CNV event and "suboptimal concordance" when a subset of callers predict the CNV event while other callers do not.

**Known truth/"gold standard"**: A set of CNV calls that has been validated extensively or identified using a reliable genomic technology (such as high-resolution microarrays) or a high confidence algorithm independent of the input exome-based callers.

**Label:** A discrete identity ("True" or "False") assigned to each CNV prediction based on their intersection with the known truth set (validated calls).

**Singleton:** A CNV prediction made by a single caller but not by other callers. Singleton calls have no concordance with other callers.

### SUPPLEMENTAL METHODS AND RESULTS

We used four exome CNV callers (CANOES, CLAMMS, CODEX, and XHMM) to predict CNVs within 503 samples. Since microarray validations were available for 291 samples, our analysis was restricted to these samples when microarray data were used as the validation set or "gold standard". First, we pooled the CNV predictions from 503 samples and resolved the breakpoints of the predictions that overlapped among each other. Next, to account for the differences in resolution and coverage between the exome and microarray probes, we chose only those CNVs that intersected with at least one SNP microarray probe for subsequent analysis. We then measured the extent of overlap among callers for each CNV event and labeled the calls as either "True" or "False" based on their overlap with microarray data. Finally, using the breakpoint-resolved CNVs >50 kbp (or >5 kbp when CLAMMS calls were treated as the "truth set"), we trained CN-Learn to make predictions on the test data and measured the performance of the classifier by comparing the classifier-assigned label (true or false) for each call to the "truth set" label.

#### Characterization of CNVs predicted by four exome CNV callers

The four CNV callers predicted an aggregate of 41,791 CNVs from the 503 SVIP exome samples. The number of CNV predictions varied widely among the callers, but the number of duplications and deletions were consistent for each individual caller (**Supplemental Fig. S19**. The frequency of predictions varied between 10 and 50 calls per sample. Further, 20,966 of the 41,791 CNV predictions were singletons, while the remaining 20,795 predictions overlapped with at least one prediction from another caller (**Supplemental Fig. S3)**. A total of 24,456 calls were made on the 291 samples for which 'gold-standard' microarray validations were available. In order to assess the characteristics of CNVs predicted by the individual callers, we calculated the overlap of raw CNV calls with microarray validations. We found that the likelihood of a CNV call to overlap with microarray validations increased as the level of support from multiple callers increased (**Supplemental Fig. S18**). While this result provides support to integrative approaches using caller intersections, it also highlights a few key limitations of the approach. *First*, even for CNV predictions with complete concordance, we observed false positive rates between 5-24% for the four callers. In fact, we found false positive rates as high as 94% for CNVs supported by two or three callers. *Second*, between 3-92% of the predictions from the four callers that lacked support from at least one of the other three callers were in fact true CNVs as detected by microarrays (**Supplemental Fig. S18**). Therefore, the strategy to identify high-confidence CNVs based on intersections among multiple callers is not adequate compared to our CN-Learn method.

**Breakpoint resolution methods**

**Method 1: Read Depth approach (see Supplemental Fig. S1)**

***Step 1: Group overlapping predictions***

We identified and grouped overlapping CNV predictions made by multiple callers across the genome and assigned them to unique groups. Each group represented an underlying CNV event, and the genomic coordinates of the predictions within each group were potential candidates for the likely start and end positions of the underlying event.

***Step 2: Measure the extent of overlap among predictions***

Within each group, we first calculated the amount of overlap between each CNV prediction with all other predictions in the group. When the overlap between two calls was greater than 1%, we considered the CNV calls between those two callers to be concordant. For example, for callers C1, C2, and C3 with predictions P1, P2 and P3 of length L1, L2 and L3, three sets of overlap proportions (OV1, OV2, OV3) can be measured as follows:

**OV1** = [(L1 ∩ L1)/L1, (L1 ∩ L2)/L1, (L1 ∩ L3)/L1] for C1;

**OV2** = [(L1 ∩ L2)/L2, (L2 ∩ L2)/L2, (L2 ∩ L3)/L2] for C2;

**OV3** = [(L1 ∩ L3)/L3, (L2 ∩ L3)/L3, (L3 ∩ L3)/L3] for C3.

Each set of overlaps for a prediction represents the overlap architecture of a particular genomic region from the perspective of a particular caller. In turn, the elements within each set represent the extent of support conferred to the prediction by its overlapping counterparts, including overlap with itself. For example, a non-overlapping (singleton) prediction made by caller C3 would be represented as [0,0,1], while a prediction made by caller C1 that has a 50% overlap with a prediction from C2 would be represented as [1,0.5,0].

***Step 3: Generate consensus coordinate pairs for overlapping CNVs***

For each group, we generated all possible pairs of start and end coordinates using the coordinates predicted by each caller. For the predictions P1, P2 and P3 from *Step 2*, nine possible coordinate pairs can be generated if and only if $P_{end} – P_{start} > 0$: [($P1_{start}$ & $P1_{end}$), ($P1_{start}$ & $P2_{end}$), ($P1_{start}$ &

$P3_{end}$), ($P2_{start}$ & $P1_{end}$), ($P2_{start}$ & $P2_{end}$), ($P2_{start}$ & $P3_{end}$), ($P3_{start}$ & $P1_{end}$), ($P3_{start}$ & $P2_{end}$), ($P3_{start}$ & $P3_{end}$)]. As these newly generated coordinate pairs do not correspond to their original caller identities, we assigned the overlap information of the caller that contributed the start coordinate to each coordinate pair for subsequent analysis. For example, OV1 can be assigned to ($P1_{start}$ & $P1_{end}$), ($P1_{start}$ & $P2_{end}$), and ($P1_{start}$ & $P3_{end}$); OV2 to ($P2_{start}$ & $P1_{end}$), ($P2_{start}$ & $P2_{end}$), and ($P2_{start}$ & $P3_{end}$); and OV3 to ($P3_{start}$ & $P1_{end}$), ($P3_{start}$ & $P2_{end}$), and ($P3_{start}$ & $P3_{end}$).

We also tested the performance of CN-Learn both when assigning the overlap information of the caller contributing the end coordinate and when assigning the averaged overlap information of the callers contributing the start and end coordinates to the breakpoint-resolved CNV (**Supplemental Fig. S20**). For example, running CN-Learn when OV1 was assigned to breakpoint pairs containing $P1_{end}$, OV2 to breakpoint pairs containing $P2_{end}$ and OV3 to breakpoint pairs containing $P3_{end}$ yielded precision of 91% and recall of 86% with a 70% training set.

***Step 4: Measure average read depth ratio***

For each consensus coordinate pair generated in Step 3, we identified the number of exome capture probes located within the putative CNV coordinates, and selected half of this number of probes from the immediate flanking regions. We calculated the average read depth from all capture probes within the coordinate pairs ($RD_{pair}$) and the average depth from the probes within the left and right flanking regions ($RD_{left}$, $RD_{right}$). We then used these three measures to calculate the ratio of read depths for each coordinate pair as follows:

$RD_{ratio} = RD_{pair} / [RD_{left} + RD_{right}]$

***Step 5: Select the coordinate pair that best represents the underlying CNV***

Within each group, we selected the coordinate pair with either the largest (duplication) or the smallest (deletion) $RD_{ratio}$ to represent the underlying CNV event.

**Method 2: Merge overlapping predictions (see Supplemental Fig. S2)**

***Step 1: Group overlapping predictions***

We identified and grouped overlapping CNV predictions made by multiple callers across the genome and assigned them to unique groups, with each group representing an underlying CNV event.

***Step 2: Select start and end coordinates of the underlying CNV***

We selected the smallest and largest genomic coordinates among the predictions within each group as the start and end positions of the underlying event. For example, for callers C1, C2, and C3 with predictions P1, P2 and P3 of length L1, L2 and L3, the underlying CNV event P4 of length L4 can be derived by assigning the smallest start coordinate among P1, P2 and P3 as the P4 start coordinate. Similarly, the largest end coordinate among the three predictions was assigned as the P4 end coordinate.

***Step 3: Measure the extent of overlap of each prediction with the breakpoint resolved CNV***

Within each group, we then calculated the amount of overlap between each overlapping prediction P1, P2 and P3 and the breakpoint-resolved CNV (P4). The overlap proportion (OV) for the breakpoint resolved CNV was measured as follows:

**OV** = $[(L1 \cap L4)/L4, (L2 \cap L4)/L4, (L3 \cap L4)/L4]$

A step-by-step outline of the two approaches to resolve breakpoint conflicts are illustrated in **Supplemental Figures S1 and S2**. Using the two methods independently, we resolved the breakpoints for 20,795 concordant predictions in 503 samples to obtain 8,382 CNVs. The total number of CNVs obtained from both the breakpoint-resolution methods was the same as the same number of CNV events was interrogated. This resulted in a final set of 29,101 unique CNVs (8,382 concordant CNVs + 20,719 Singletons). The number of singletons before and after breakpoint resolution was slightly different, as singletons with less than 1% overlap with other CNVs were not counted in the concordance calculations but were assimilated into their overlapping CNVs during breakpoint resolution. We then labelled the CNVs as either "True" or "False" based on their overlap with microarray-validated breakpoints. Among the 291 samples with microarray validations, 8,085 of the 29,101 breakpoint-resolved CNVs overlapped with regions covered by at least one microarray probe, and 2,506 of these calls were larger than 50 kbp. This final set of CNVs was subsequently used to train CN-Learn (**Figure S3**).

**CN-Learn recovers CNVs that lack complete concordance (with CLAMMS as "gold standard")**

We analyzed the concordance profile of CNVs labelled using the predictions made by CLAMMS as the "gold standard" before and after CN-Learn classification. Post-classification analysis indicated that 45% of all CNVs classified as true had support from the remaining three callers, while the other 55% lacked support from at least one of the three callers (**Supplemental Fig. S16**). Using a single random draw of 30 samples to train CN-Learn and 453 samples as a test set, we obtained predictions for 14,923 CNV events from the test samples. Among these 14,923 CNV calls, only 1,612 (11%) were supported by all four methods (**Supplemental Fig. S16**), and about 44% (715/1,612) of these fully concordant calls intersected with the "gold standard" predictions made by CLAMMS. This proportion is much lower than the 97% observed for CNVs supported by all four callers when microarray calls were used as the "gold standard", reiterating the low concordance rates among the callers. After classification, 92% (1,376/1,495) of all CNVs classified by CN-Learn as true intersected with the "gold standard" CLAMMS calls (**Supplemental Fig. S16**). This indicated that CN-Learn also classified an additional 119 CNVs with support from each of the three callers as true, even when they were not called by CLAMMS. Although such calls are considered false positives for measuring the accuracy of the classifier, given the extremely high precision demonstrated by CN-Learn, this small subset of calls could still be considered for further investigation. CN-Learn therefore offers a robust strategy to prioritize CNVs with complete concordance when using an exome-based CNV caller as a gold standard.

**CN-Learn using Support Vector Machine and Logistic Regression**

In addition to building CN-Learn as a Random Forest classifier, we also evaluated the ability to build the classifier as either a Support Vector Machine (SVM) classifier with a linear kernel or a Logistic Regression (LR) classifier. If the performances of these linear classifiers were robust, it would reiterate the discriminatory power provided by the features supplied to the model and alleviate concerns of possible overfitting by the non-linear Random Forest classifier. Using microarray calls as the truth set, the precision rates achieved by SVM and LR were very similar to the rates achieved by the Random Forest classifier (**Supplemental Fig. S12**). However, the recall rates achieved by SVM and LR were on average 10% lower than the recall rates achieved by the Random Forest classifier. These conclusions did not hold when CLAMMS calls were used as the truth set (**Supplemental Fig. S12)**. In this case, the precision rates achieved by the linear classifiers were 20% lower than the rate achieved by the Random Forest classifier, while the recall rates were only half as much as the 85% achieved by Random Forest (**Supplemental**

**Fig. S12**). Although the CN-Learn software package provides the ability to use any of these three models, we recommend Random Forest as the selected classifier given the higher recall rates it achieves without compromising the precision.

**Independent assessment of CN-Learn using the 1000 Genomes data**

We analyzed 90 samples from the 1000 Genomes project that were also used to measure the performance of CODEX (Jiang et al. 2015). An exhaustive list of all validated CNVs for these 90 samples were available from three different sources (Altshuler et al. 2010; Conrad et al. 2012; McCarroll and Altshuler 2007). We pooled the validated CNVs from the three sources to obtain 10,053 CNVs, and merged the coordinates of overlapping validated CNVs to obtain a final list of 7,303 CNVs (**Supplemental Table 4**). Next, we made 19,453 CNV predictions using the four callers on the 90 samples, and labeled them based on their overlap with the 7,303 validated CNVs. We trained and tested CN-Learn using 9,697 CNVs between the size range 5 kbp and 5 Mbp (**Supplemental Fig. S5**) to obtain performance metrics for CN-Learn that are directly comparable with naïve concordance-based approaches.

**(1) Performance evaluation of CN-Learn**

When trained with 70% of the samples (63/90), CN-Learn achieved a high precision rate of 93% (**Supplemental Figs. S6A** & **S10**). However, the recall rate (73%) was slightly lower than the recall rate obtained when using the SVIP samples (86%). This decrease in performance could be due to the smaller number of 1000 Genomes samples (n=63 from 1000 Genomes versus n=204 from the SVIP) used to train CN-Learn. While a caller-specific predictor (CANOES) and a genomic predictor (target probe count) showed the highest discriminatory power with the SVIP samples, mappability and read-depth ratio (both genomic predictors) displayed the most discriminatory power when analyzing the 1000 Genomes samples (**Supplemental Fig. S6B**). We also generated Venn diagrams for concordance profiles of calls before and after classification by CN-Learn (**Supplemental Fig. S6C**). Overall, the results obtained are comparable to those obtained for the SVIP cohort.

**(2) Measure precision/recall rates directly comparable with concordance approaches**

Using only the labeled calls for calculating precision and recall rates is a standard practice for measuring the performance of machine learning classifiers (Saito and Rehmsmeier 2015; Liang et al. 2019). Since the original CNV calling algorithms do not use such labels to make CNV predictions, the recall rates for the naïve concordance-based methods could only be calculated as the number of calls predicted from individual callers or concordant calls from two or more callers divided by the total number of validated CNVs (**Supplemental Fig. S7**). The availability of validated CNVs for the 1000 Genomes samples from multiple sources enabled us to make reliable and comparable precision/recall estimates. We first trained CN-Learn using CNVs between 5 kbp and 5 Mbp in size from 70% of the data (63/90 samples), and then made predictions on CNVs within the same size range in the remaining 27 samples that contained 2,270 distinct validated CNVs. We calculated precision and recall independently for each individual caller (CANOES, CLAMMS, CODEX and XHMM), for combinations of individual callers (concordant calls within pairs of callers, combination of three callers, and all four callers), and for CN-Learn as follows (**Supplemental Table 3**):

1. **Precision:** For individual callers and their combinations, the fraction of predicted CNVs that overlapped with the 2,270 validated CNVs. For CN-Learn, the fraction of CNVs classified as true that overlapped with the 2,270 validated CNVs.
2. **Recall:** For individual callers and their combinations, the fraction of 2,270 validated CNVs that were correctly identified. For CN-Learn, the fraction of 2,270 CNVs classified as true by the classifier.

This analysis allowed us to measure the recall rates of CN-Learn based on both the labels of CNV predictions and the set of all validated CNVs in the 27 samples. While the recall estimates are low for most callers and combinations of callers, these estimates are similar to those reported by Jiang and colleagues (Jiang et al. 2015). These results serve as a heuristic for comparing the naïve concordance-based methods with CN-Learn.

**CNV frequency and performance of CN-Learn**

We tested two aspects of CNV frequency and its influence on the performance of CN-Learn. *First*, using the 90 samples from the 1000 Genomes project, we tested whether using the frequency of CNVs within the cohort as a predictor would improve the performance of CN-Learn. *Second*, using the 291 samples from the SVIP cohort, we tested whether the classification performance varies for CNVs with different frequencies in the cohort.

**(1) CNV frequency as an additional predictor to CN-Learn**

When we included CNV frequency as an additional predictor for CN-Learn, we observed a minor improvement in classifier performance. For example, after training with 70% of the 90 samples, CN-Learn achieved 94% precision and 71% recall rates when CNV frequency was used as a predictor, compared with the 93% precision and 73% recall rates obtained without utilizing CNV frequency information (**Supplemental Fig. S10**). Interestingly, CNV frequency showed the highest predictive power (16%) among all of the predictors (**Supplemental Fig. S11**), but it did not significantly improve the performance of the classifier.

**(2) Classification performance for CNVs at different frequencies**

To test the performance of CN-Learn at different frequencies, we classified each CNV in the 291 SVIP samples as (i) very rare (frequency ≤2 occurrences), (ii) rare (frequency between 2 and 10 occurrences), (iii) common (frequency between 10 and 30 occurrences), and (iv) very common (frequency >30 occurrences). We then used two different approaches to evaluate the predictive power of CN-Learn:

(a) First, we trained CN-Learn using CNVs of all frequencies in the training set, and independently classified CNVs in each of the four frequency classes in the testing set.
(b) Second, we trained CN-Learn using only the very rare CNVs in the training set, and independently classified CNVs in each of the four frequency classes in the testing set.

These two approaches resulted in eight independent experiments from which the performance of CN-Learn could be assessed (**Supplemental Fig. S9**). CN-Learn achieved > 90% precision and > 80% recall rates in every experiment, except when classifying very rare CNVs after training with very rare CNVs (precision=78%, recall=75%). In particular, the first four experiments indicate that the classification performance of CN-Learn for very rare CNVs approaches the precision and recall rates achieved for very common CNVs. These results highlight the robust performance and the ability of the twelve predictors used by CN-Learn to capture the latent genomic and caller-specific signatures of true CNVs, regardless of their frequency in the cohort.

### SUPPLEMENTAL FIGURES

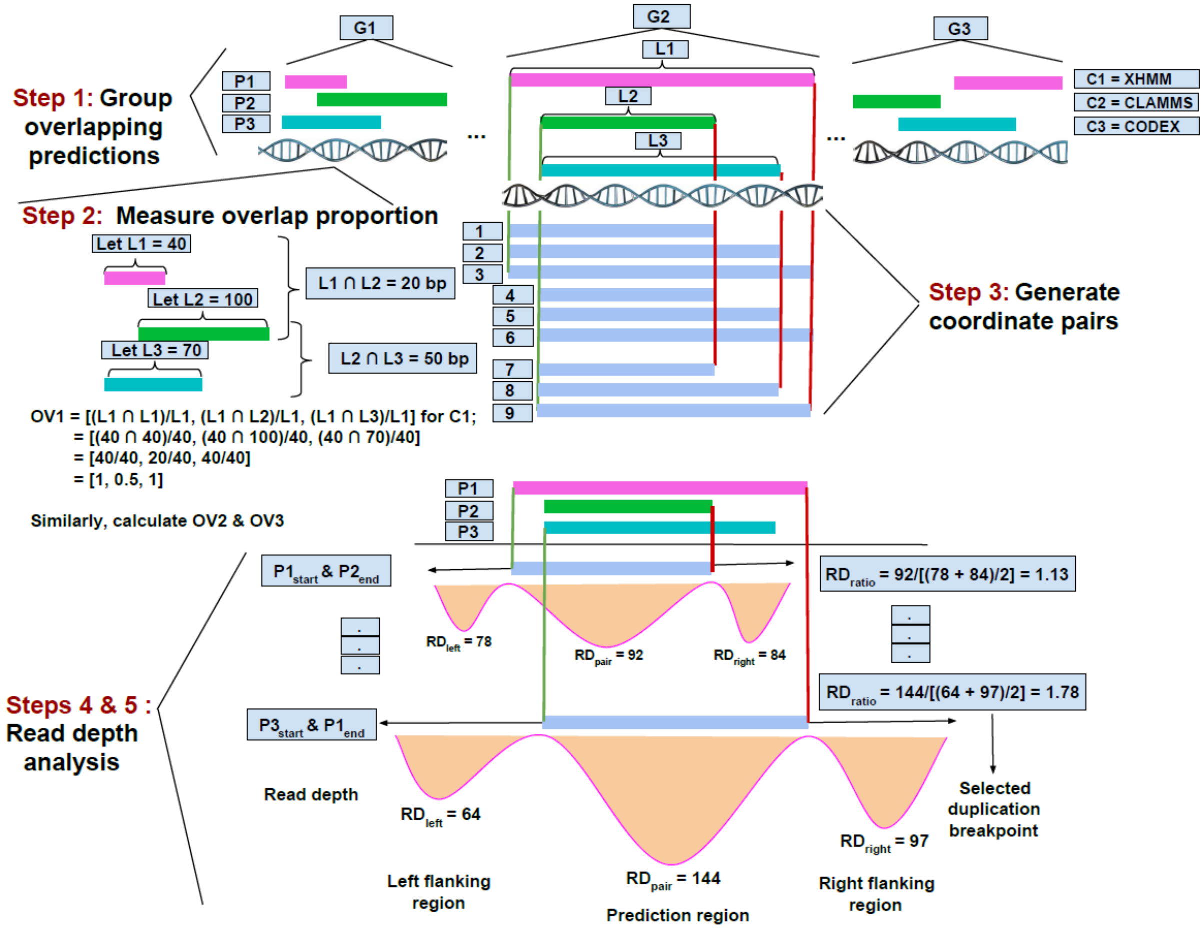

**Supplemental Figure S1: Illustration of the read-depth based breakpoint resolution method.** ***Step 1*****:** Overlapping CNV predictions from the four exome CNV callers were assigned into groups. ***Step 2*****:** Within each group, the amount of overlap between overlapping predictions made by different callers was measured. ***Step 3*****:** The start and end coordinates of the overlapping predictions were then used to generate all possible start and end coordinate pairs to represent the underlying CNV event. ***Steps 4 & 5*****:** For each newly generated coordinate pair, the average read depth of the exome capture probes within the CNV region and the average read depth of probes in the corresponding left and right flanking regions were measured. The ratio of average read depth between the putative CNV and the flanking regions was then measured. The coordinate pairs with the smallest read depth ratio were chosen for deletions, while the coordinates with the highest read depth ratio were chosen for duplications.

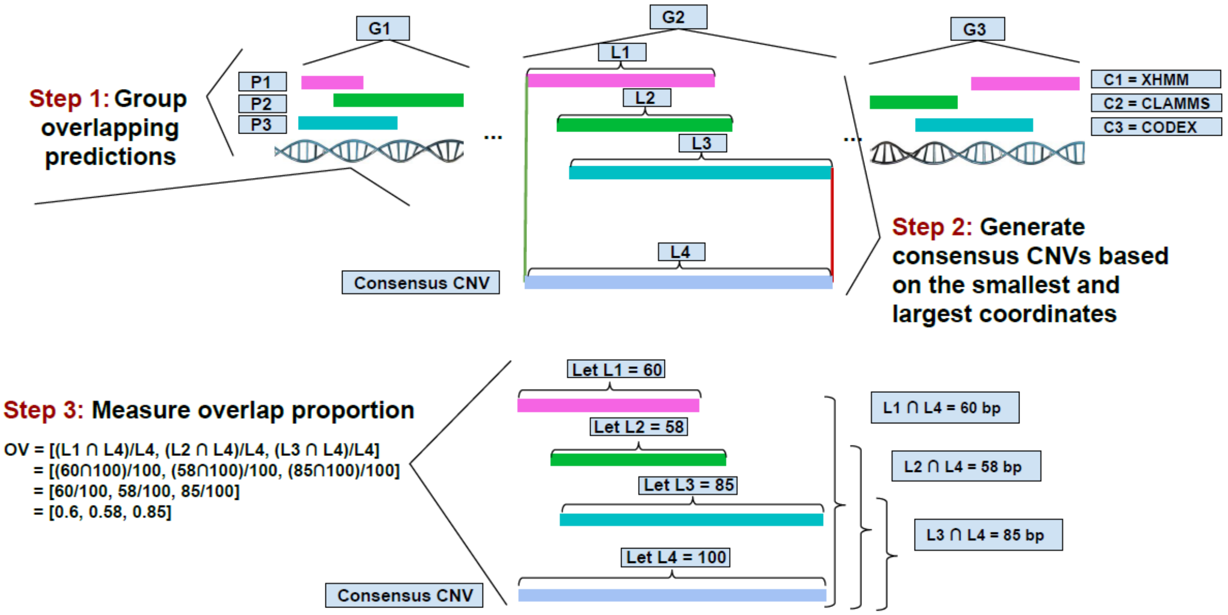

**Supplemental Figure S2: Illustration of breakpoint resolution by merging overlapping CNVs method.** ***Step 1***: Overlapping CNV predictions from the four exome CNV callers were assigned into groups. ***Step 2***: Within each group, the coordinates for the underlying CNV event were derived using the smallest and largest coordinates among the predicted CNVs within each group. ***Step 3***: The amount of overlap of each call from individual callers with the consensus CNV was then measured.

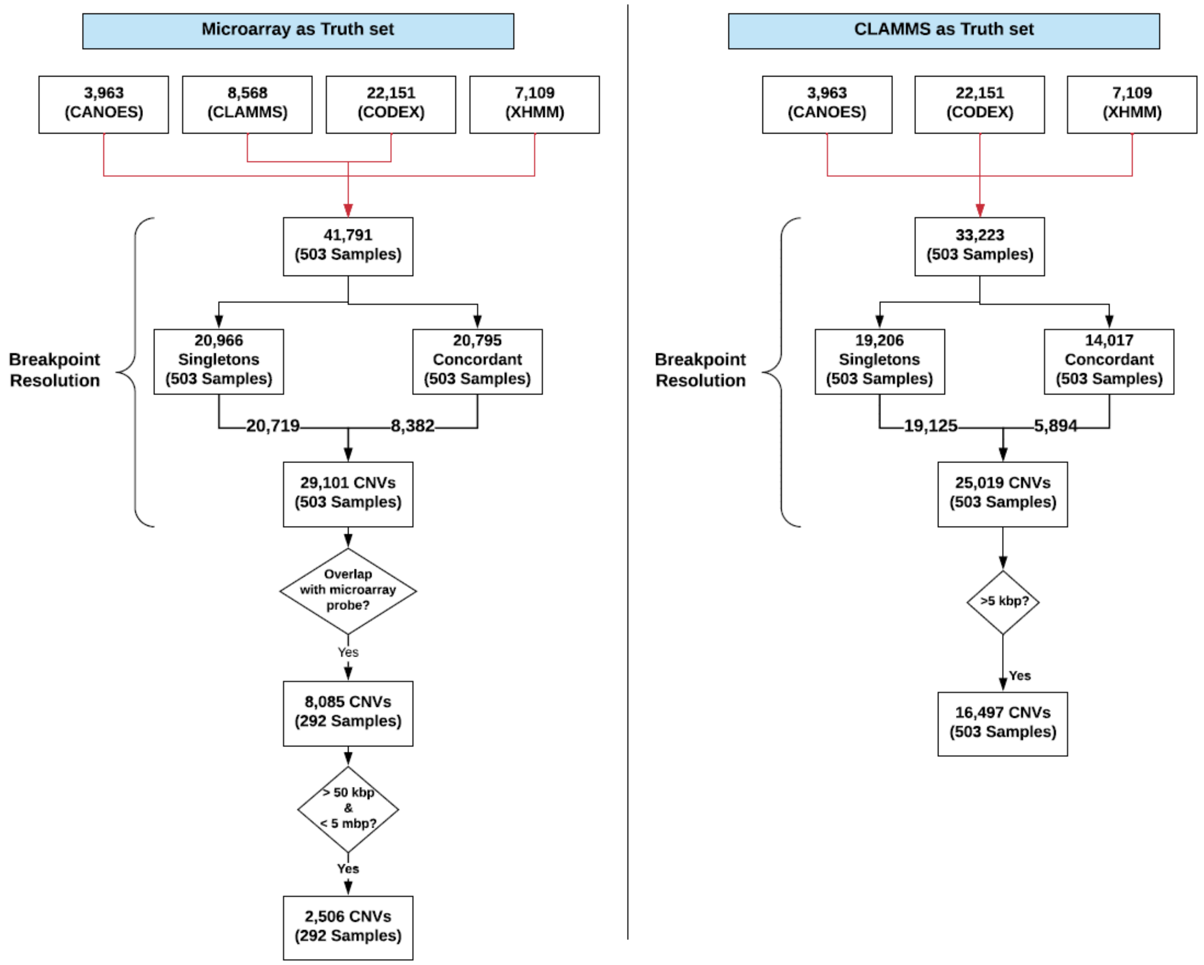

**Supplemental Figure S3: A flowchart for the number of CNVs processed through different stages of the CN-Learn pipeline.** The flowchart shows the number of CNVs from the 503 samples obtained from the SVIP cohort processed through different stages of the CN-Learn pipeline, when microarrays (left) and CLAMMS calls (right) were used as gold standards.

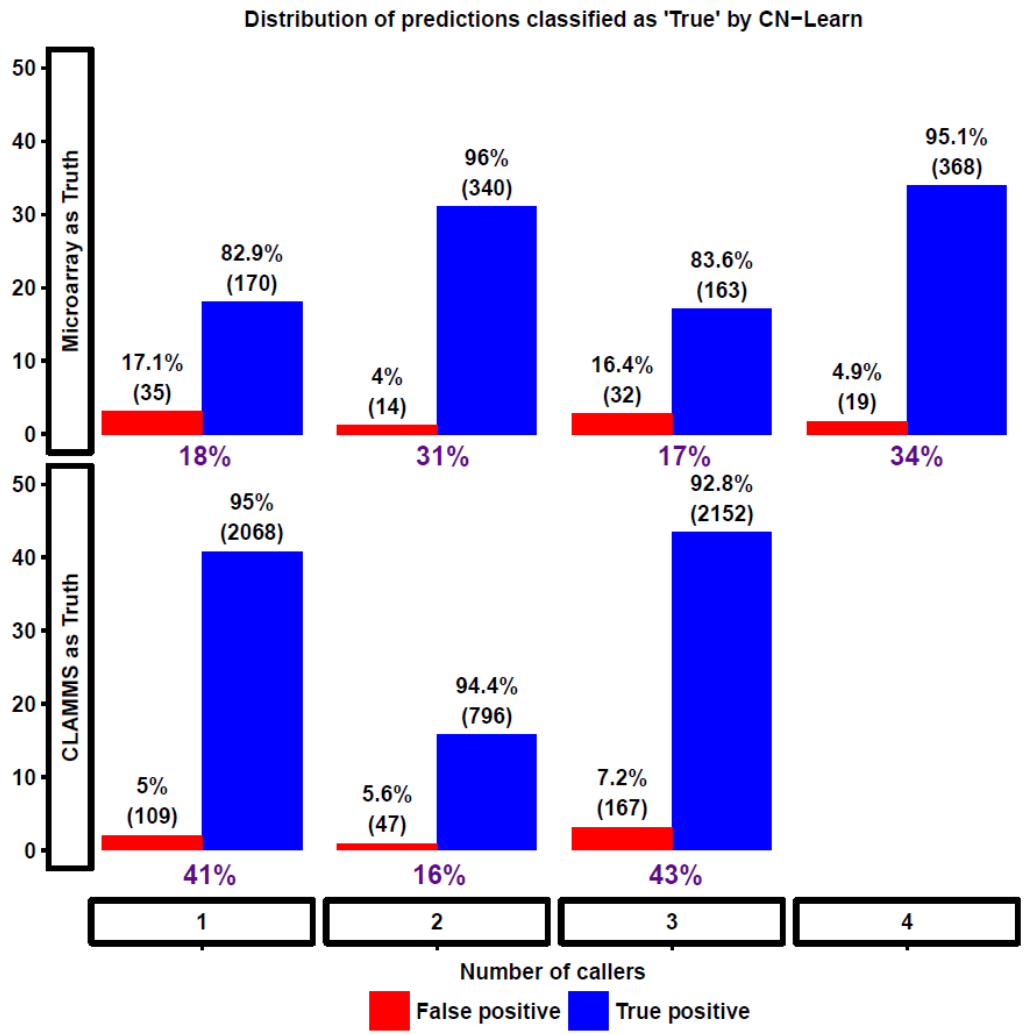

**Supplemental Figure S4: Bar plot illustrating the distribution of CNVs classified by CN-Learn as true based on concordance with one or more callers.** This plot shows the fraction of all CNVs classified as true within each concordance bin when microarray (**bottom panel**) or CLAMMS (**top panel**) validations were used as gold standard. The blue bars indicate the proportion of CNVs within each concordance bin that overlap with the validated set of CNVs, while the red bars indicate the proportion of CNVs within each concordance bin that fail to overlap with the validated CNVs.

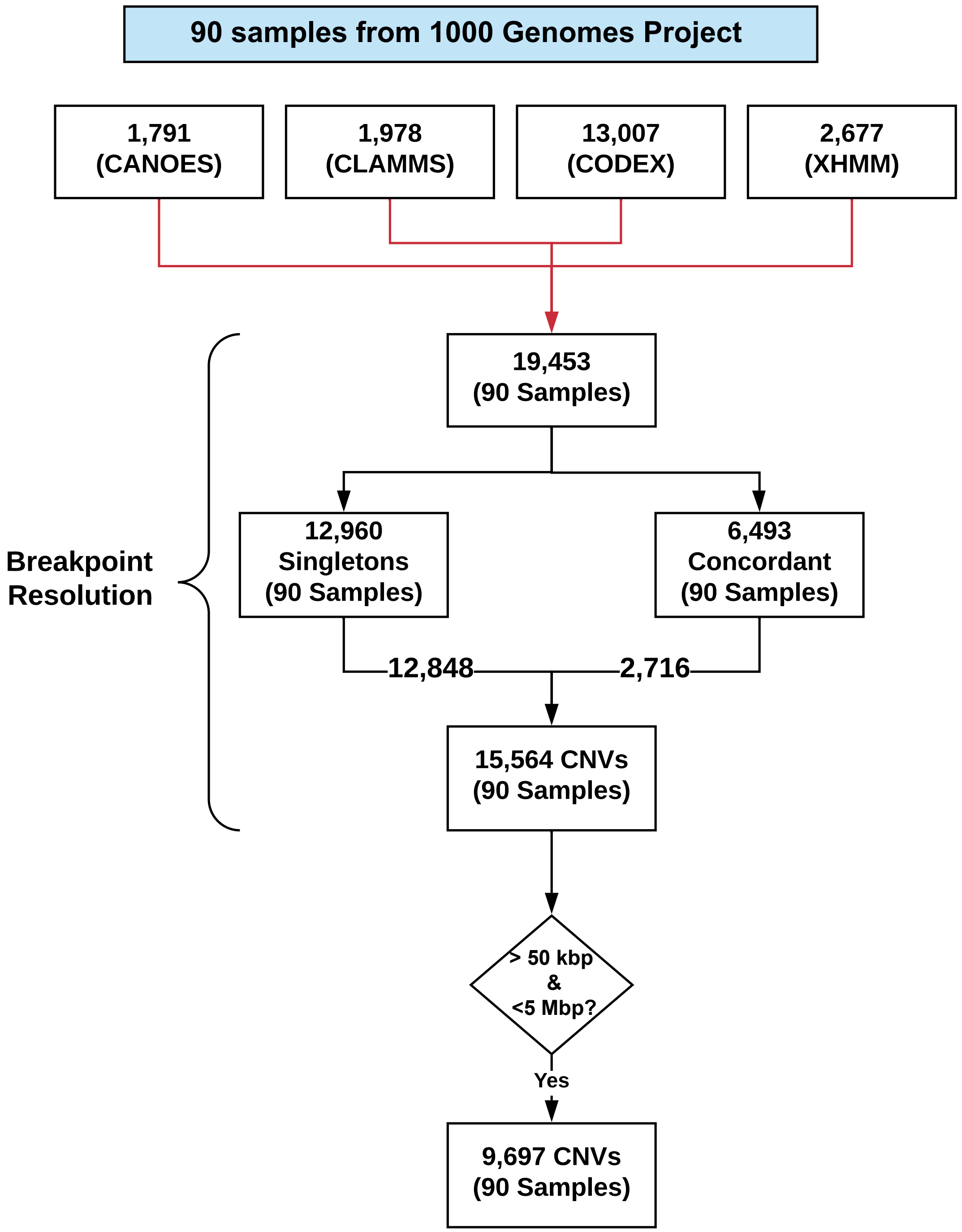

**Supplemental Figure S5: Flowchart for analysis of 1000 Genomes data using CN-Learn.** The flowchart shows the number of CNVs from the 90 samples obtained from the 1000 Genomes project processed through different stages of the CN-Learn pipeline.

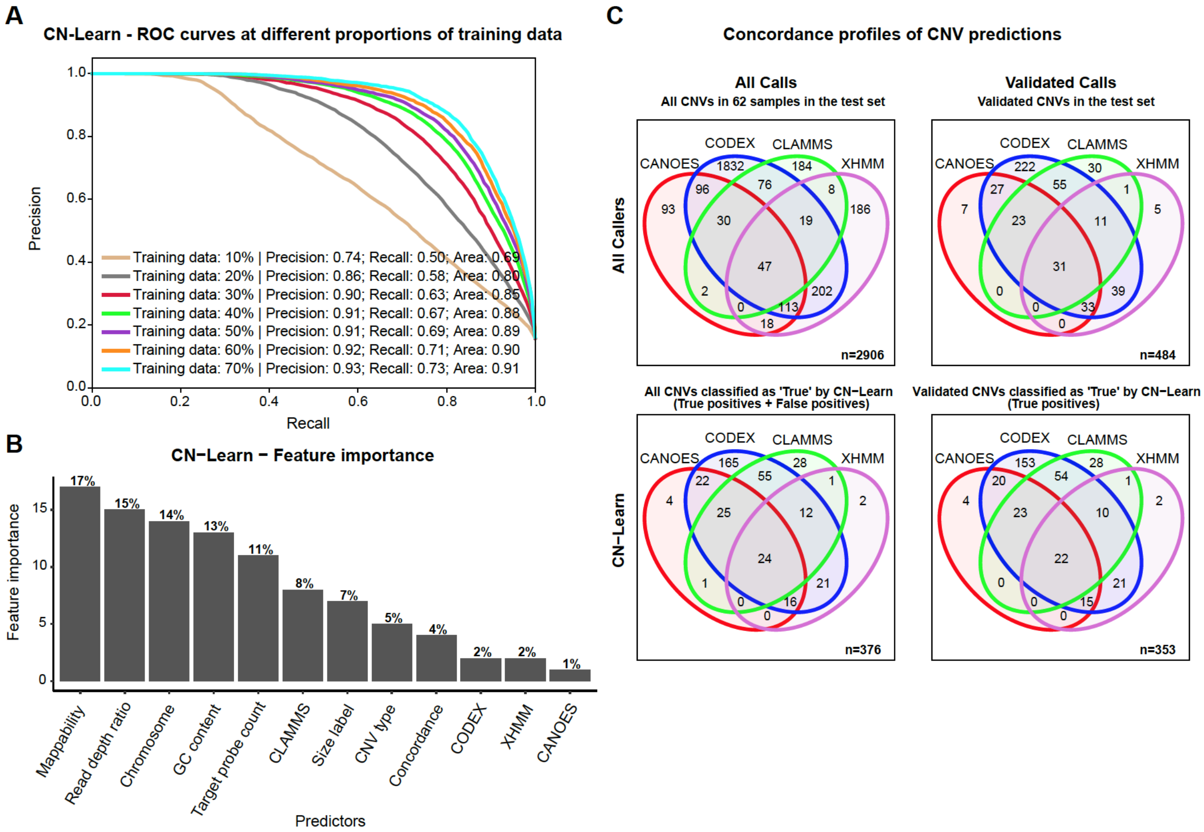

**Supplemental Figure S6: Performance evaluation of CN-Learn using 1000 Genome datasets. (A)** Receiver operating characteristic (ROC) curves indicate the trade-off between the precision and recall rates when CN-Learn was trained as a Random Forest classifier using CNV calls of samples from the 1000 Genomes project. Each curve represents the performance achieved when using different proportions of samples to train CN-Learn, ranging from 10% to 70% in increments of 10%. The results shown were from experiments aggregated across 10-fold cross validation. **(B)** The relative importance of genomic and caller-specific features supplemented to CN-Learn for 1000 Genomes CNV calls is shown. Data shown here are the averages of values obtained across 10-fold cross-validation after using 70% of the samples for training. **(C)** The concordance profile of 1000 Genomes CNVs before and after classification by CN-Learn is shown. Venn diagrams are shown for CNVs (≥5 kbp) identified from a random draw of 27 samples out of the 90 total 1000 Genomes samples before (Top panels) and after (Bottom panels) classification by CN-Learn. The top Venn diagrams show the overlap of calls among the four callers (left) versus the subset with validations (right). The bottom Venn diagrams depict the overlap of all true calls among the four callers after classification by CN-Learn (left) and the validated subset among the calls classified as true (right).

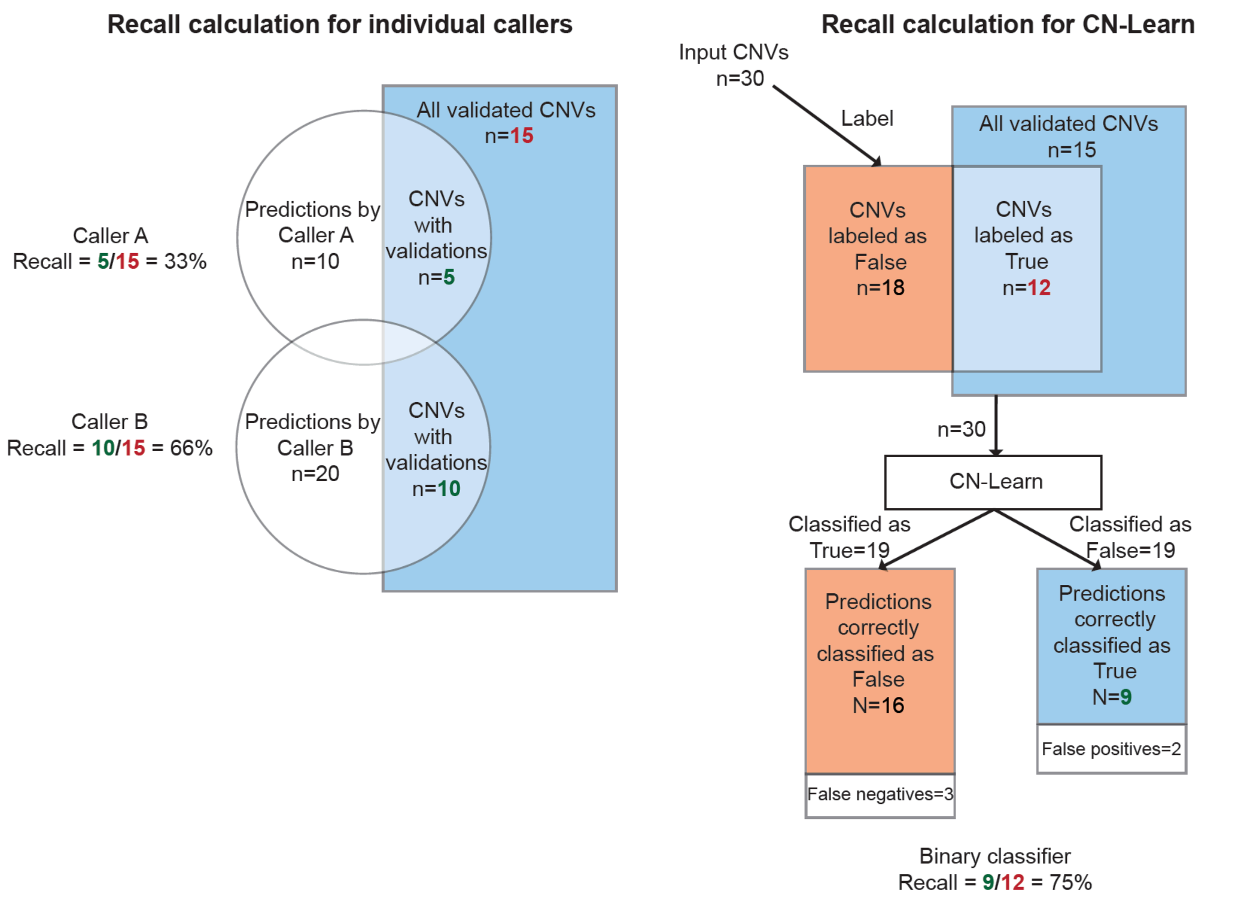

**Supplemental Figure S7: Illustration of sample recall calculations for the individual callers and CN-Learn.** For individual callers as well as the naïve concordance methods, the recall rate is the fraction of all validated CNVs that are correctly identified by the caller. For CN-Learn, the recall rate is the fraction of all CNVs labeled as "True" based on overlap with gold-standard validations that were correctly classified as true by the algorithm.

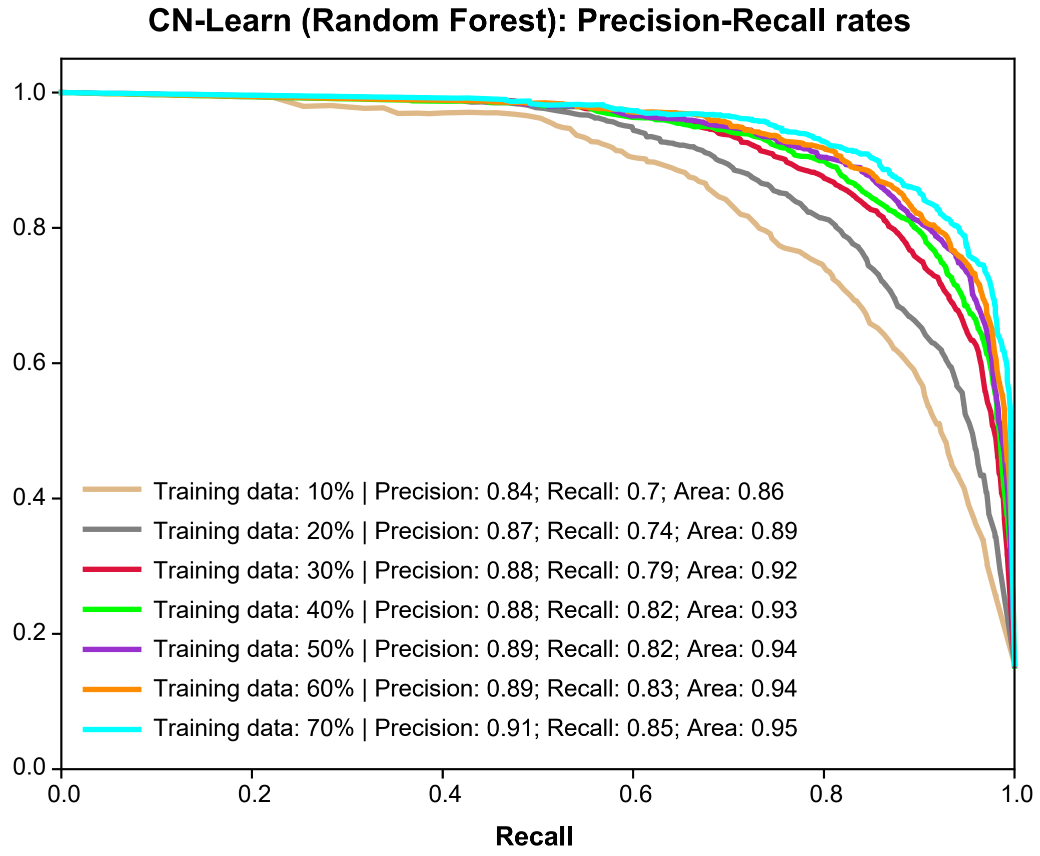

**Supplemental Figure S8: Performance of CN-Learn when breakpoints were resolved by selecting the smallest and largest coordinates (using Method 2) of overlapping CNV predictions.** Each curve represents the performance achieved while using different proportions of samples to train CN-Learn, from 10% to 70% in increments of 10%. The precision and recall values shown here are averages from 10-fold cross-validations.

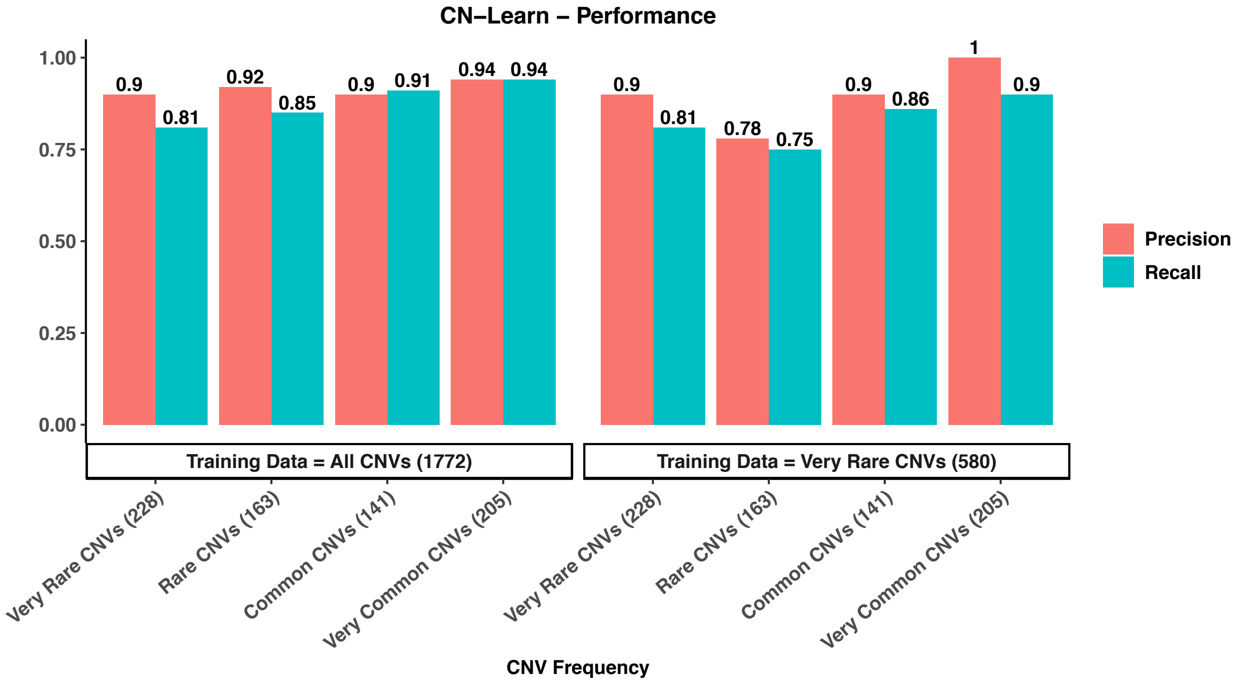

**Supplemental Figure S9: Performance of CN-Learn when trained and tested using CNVs at different frequencies.** CNVs observed once or twice within the cohort are considered very rare; rare if their frequency is between 2 and 10 occurrences; common if their frequency is between 10 and 30; and very common if they occur more than 30 times. The left panel illustrates the performance obtained when CNVs of all frequencies were used for training, while the right panel illustrates the performance obtained when only very rare CNVs were used for training. The four different groups within each panel represent the precision/recall rates obtained when very rare, rare, common and very common CNVs in the test samples were independently classified with CN-Learn, with the number of CNVs in each category listed in parenthesis. The results indicate that the performance of CN-Learn is robust across the entire spectrum of frequencies.

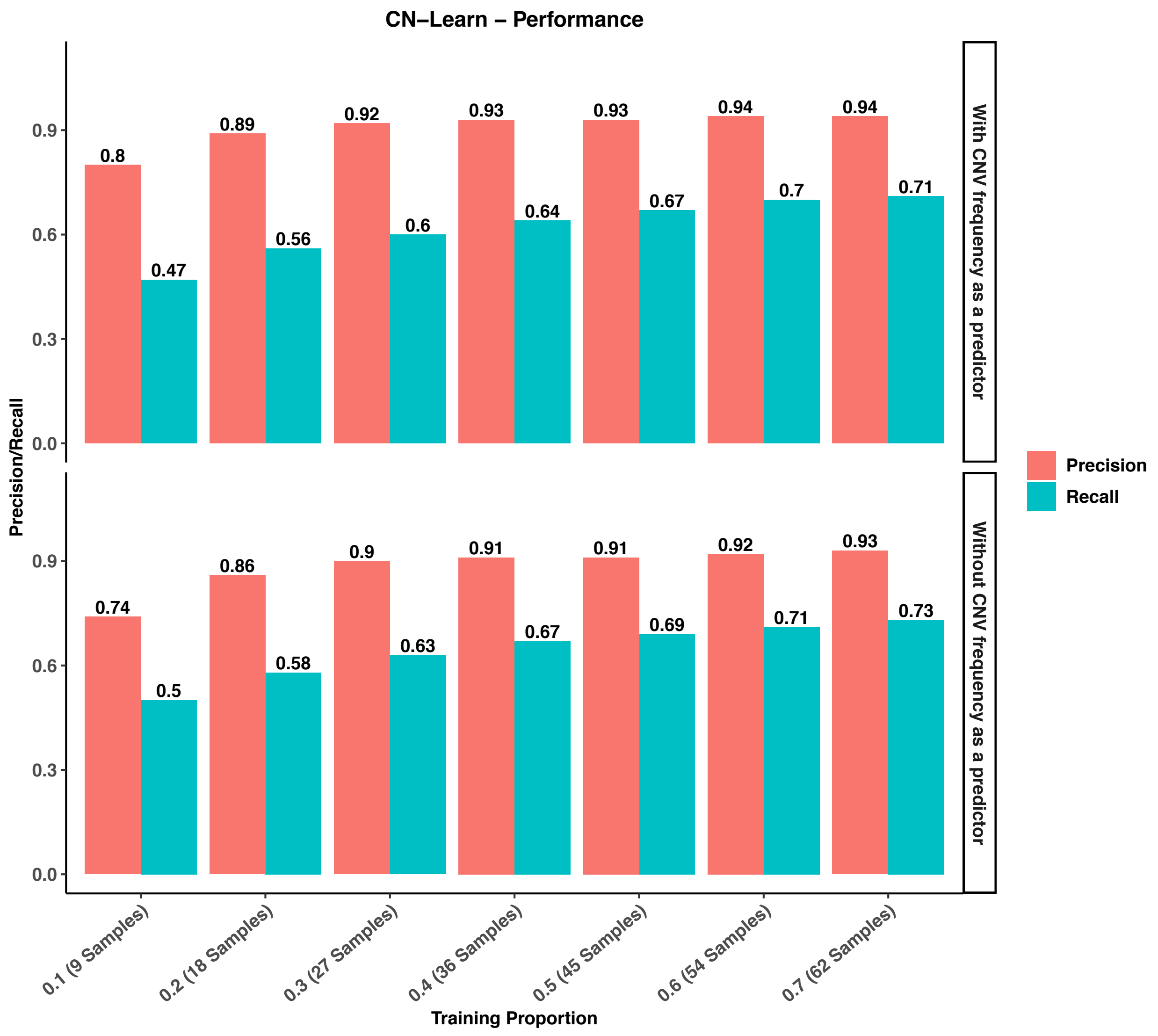

**Supplemental Figure S10: Performance of CN-Learn on 90 samples from the 1000 Genomes project.** Each group on the x-axis represents the fraction of all samples used for the training set, ranging from 10% to 70%. The top panel shows precision and recall when CN-Learn is trained using CNV frequency as one of its predictors, while the bottom panel shows precision and recall when CN-Learn is trained without CNV frequency as a predictor. The precision and recall rates obtained were comparable to the rates obtained without CNV frequency as a predictor to CN-Learn.

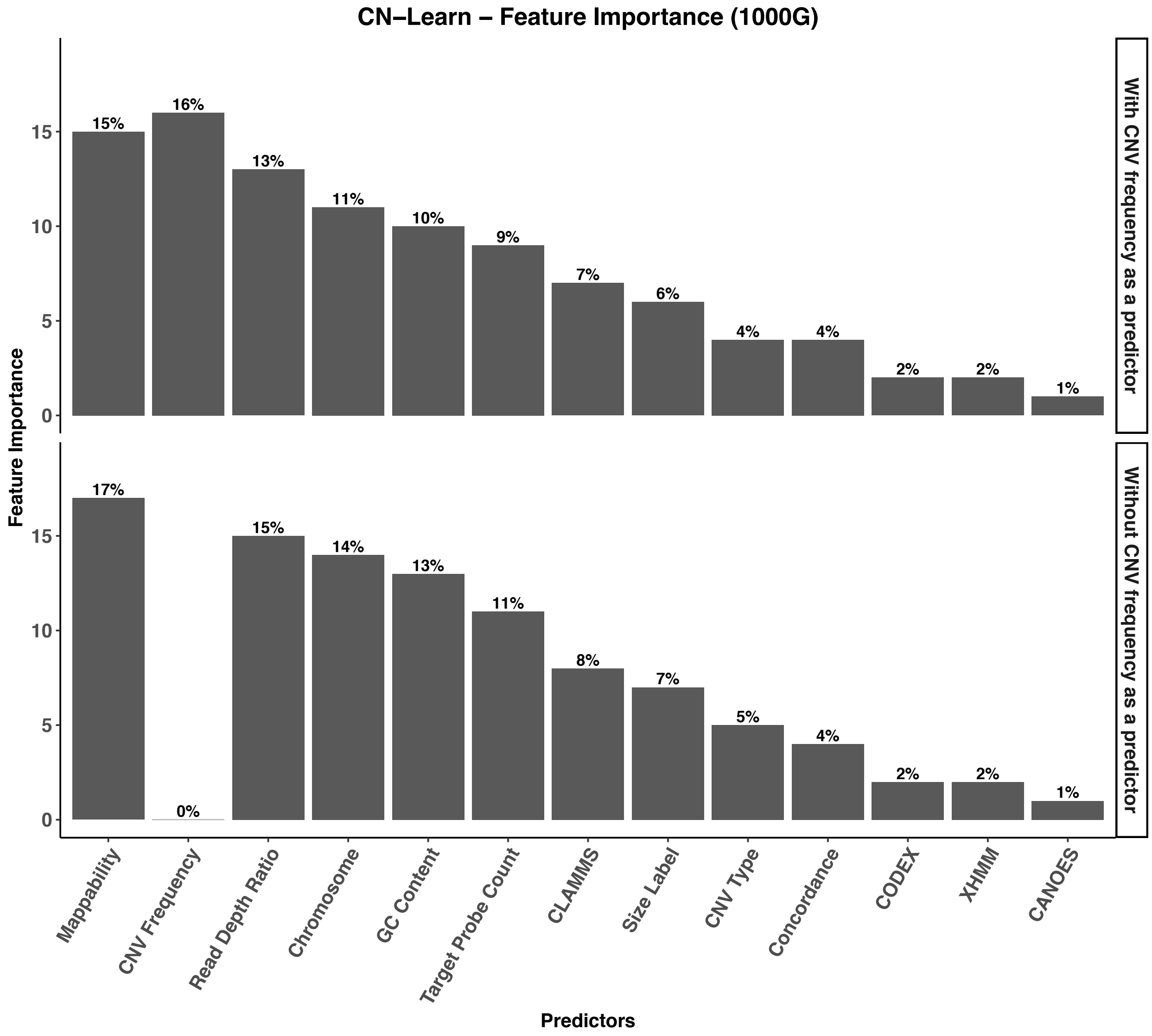

**Supplemental Figure S11: Relative importance of genomic and caller-specific features supplemented to CN-Learn, including CNV frequency.** The frequency of CNVs observed in the sequencing cohort appeared as the top predictor of true CNVs due to the differential performance and sensitivities of CNV callers in detecting true versus common CNVs.

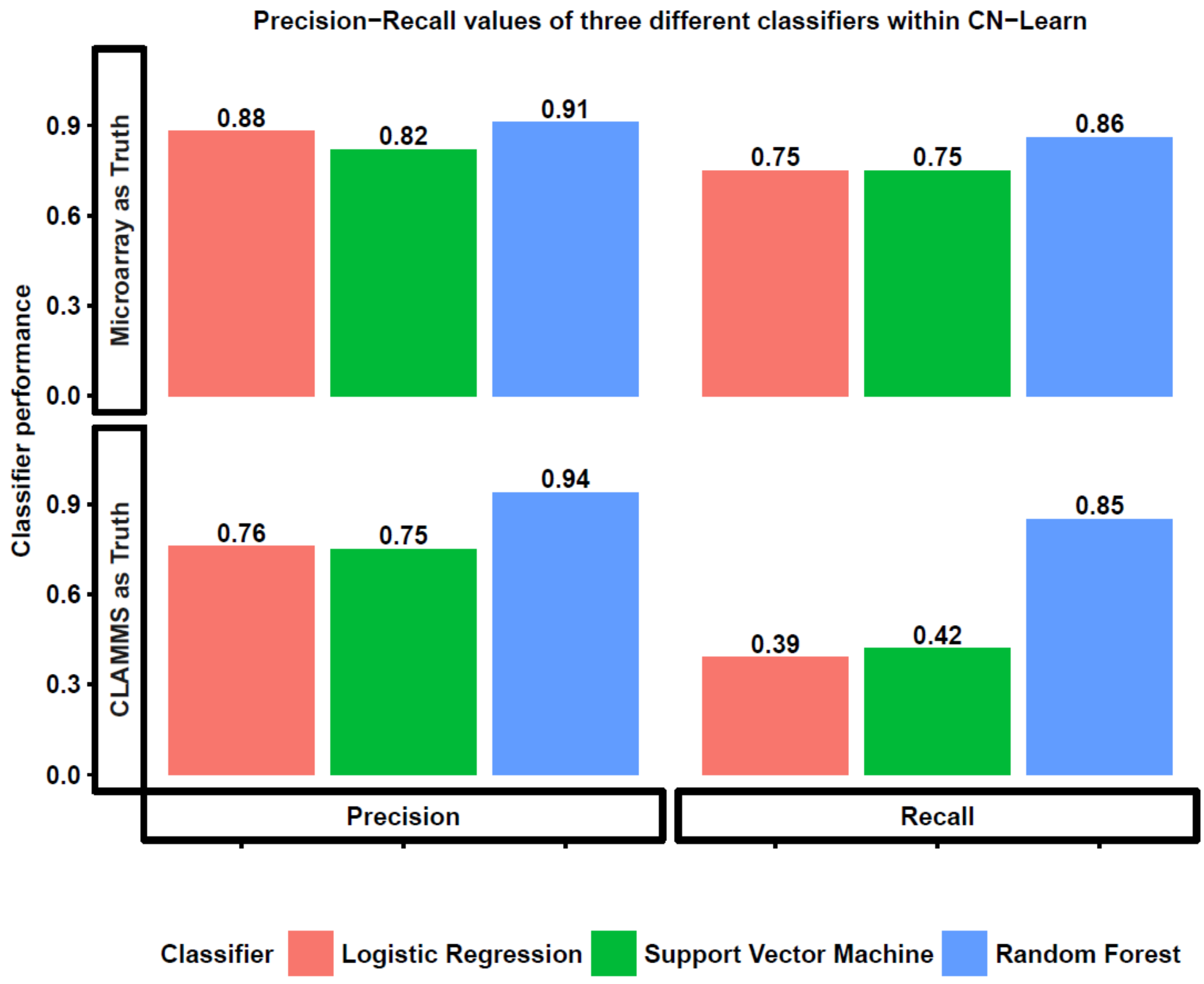

**Supplemental Figure S12: Performance comparison of three binary classifiers supported by CN-Learn.** The aggregate precision and recall values obtained by CN-Learn during 10-fold cross-validation using 70% training data are illustrated in two separate panels along the x-axis. Performance obtained when using microarray (n=291 samples) and CLAMMS (n=503 samples) validations is illustrated along the Y-axis. Each color is associated with one of the three binary classification algorithms supported by CN-Learn. These results demonstrate the superior performance of the Random Forest-based classifier.

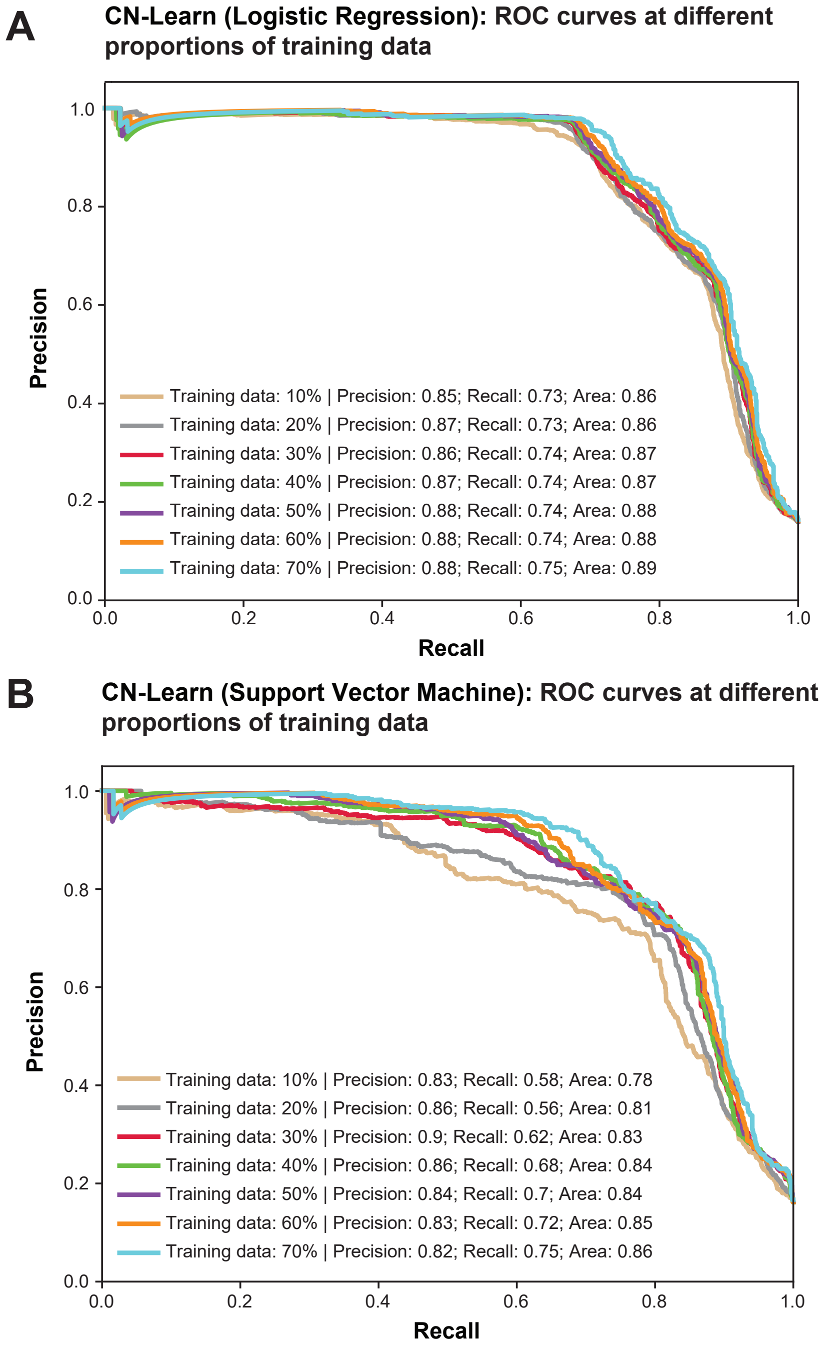

**Supplemental Figure S13: Performance of CN-Learn built as Support Vector Machine and Logistic Regression classifiers.** Each curve represents the performance (precision and recall) achieved while using different proportions of samples to train CN-Learn, from 10% to 70% in increments of 10%. Validations were based on microarray calls. The precision and recall values shown here are averages from 10-fold cross-validation.

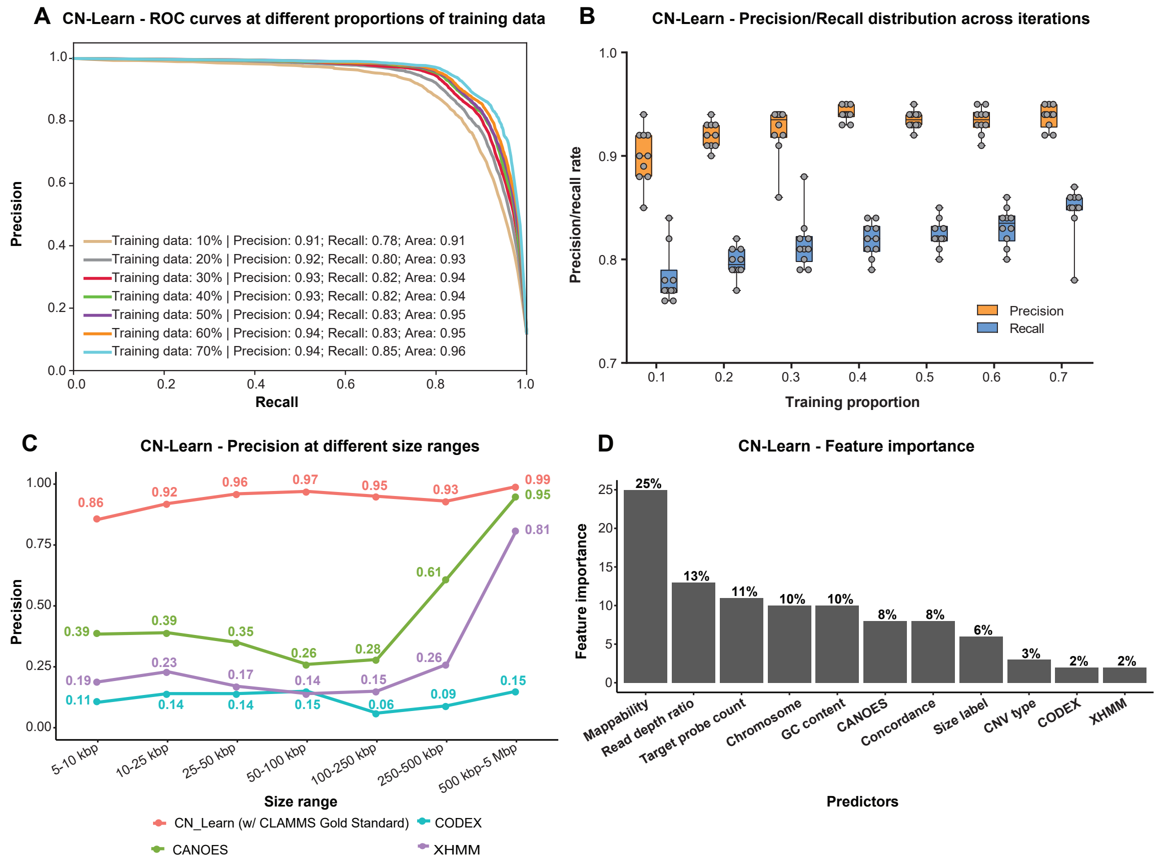

**Supplemental Figure S14: Performance of CN-Learn while using CLAMMS based validation. (A)** Receiver operating characteristic (ROC) curves indicating the trade-off between precision and recall rates when CN-Learn was trained as a Random Forest classifier are shown. Each curve represents the performance achieved when using different proportions of samples to train CN-Learn, between 10% and 70% in increments of 10%. The results shown were from experiments aggregated across 10-fold cross-validation. **(B)** Plot showing variability observed in precision and recall measures during 10-fold cross validation at various proportions of training data. Both measures varied within ±5% of their corresponding averages. **(C)** Precision rates for CNVs when CN-Learn was trained at four different size ranges compared to precision rates for CNVs from individual callers are shown. The precision rates for CN-Learn were estimated as the classification accuracy (True positives/ [True positives + False positives]), while the precision rates for the individual callers were calculated as the proportion of CNVs at each size range that were validated by CLAMMS calls. **(D)** The relative importance of each genomic and caller-specific feature supplemented to CN-Learn is shown. Data shown here are the aggregate of the values obtained across 10-fold cross-validation after using 70% of the samples for training.

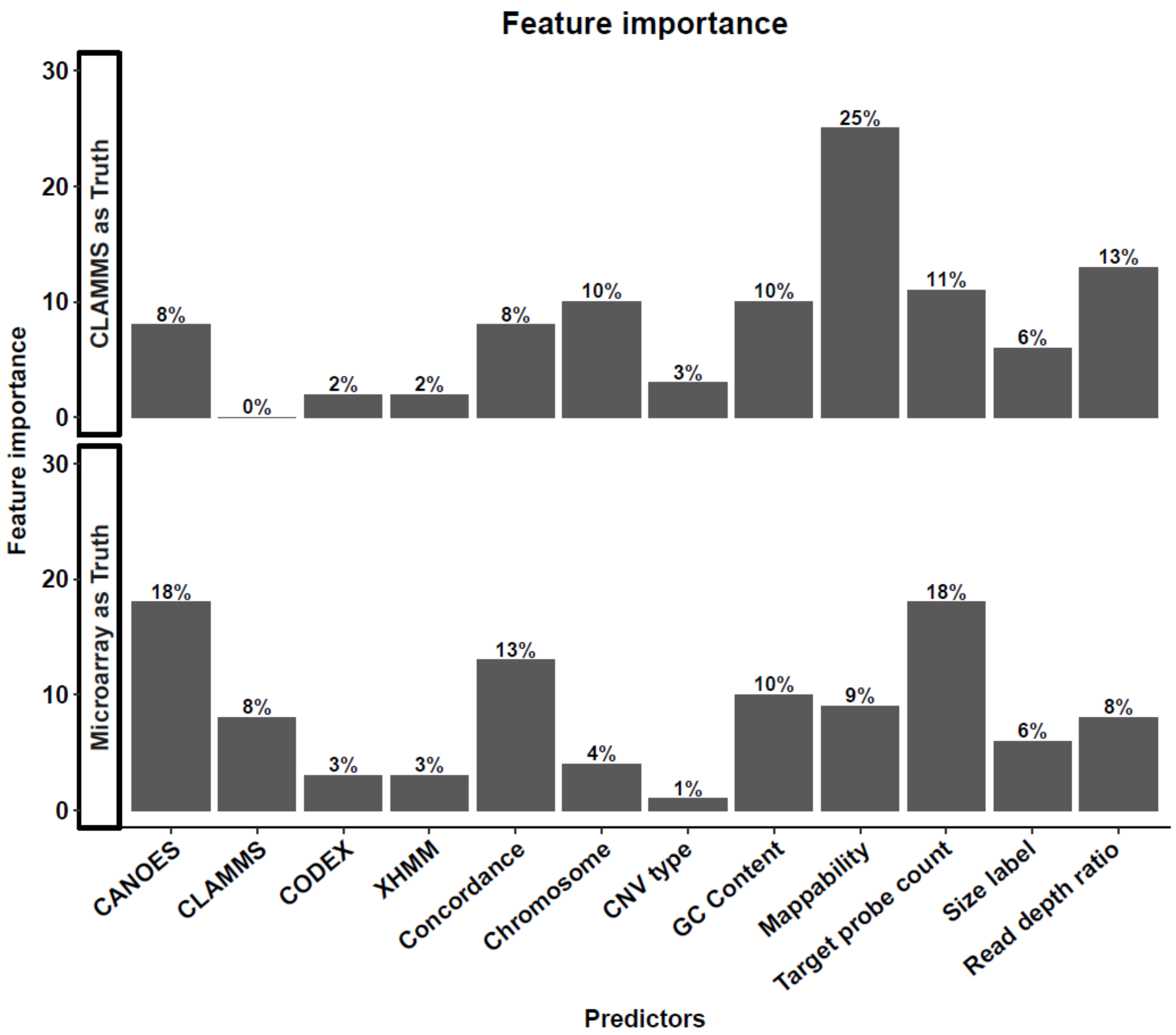

**Supplemental Figure S15: Relative importance of features used by CN-Learn in making predictions.** The feature importance values shown are averages measured based on Gini importance obtained during 10-fold cross-validation with 70% training data. **(Top panel)** Relative importance of each feature when CN-Learn was trained using predictions made by CLAMMS as the truth set (n=503 samples). **(Bottom panel)** Relative importance of each feature when CN-Learn was trained using microarray calls as the truth set (n = 291).

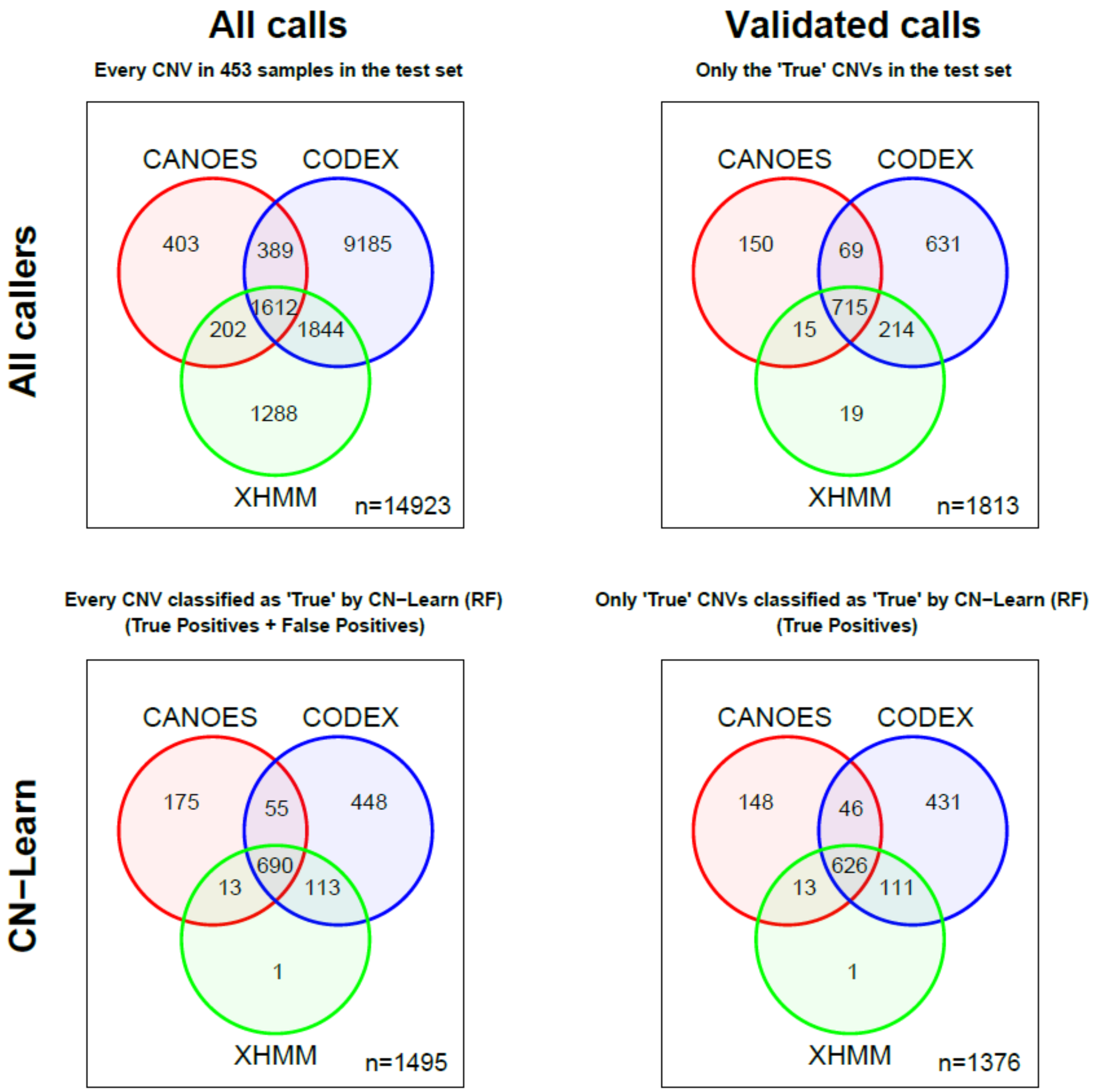

**Supplemental Figure S16: Concordance profile of CNVs before and after classification by CN-Learn using CLAMMS-based validation.** CNVs in 453 (test set) of the 503 samples before and after classification by CN-Learn are illustrated as Venn diagrams based on the CNV callers that provided support for each CNV. (**Top left**) Distribution of 14,923 CNVs in the test set prior to classification. (**Bottom left**) Distribution of 1,489 CNVs classified by CN-Learn as true (True + False positives). (**Top right**) Distribution of 1,813 CNVs in the test set that intersected with the validations. (**Bottom right**) Distribution of 1,376 CNVs classified by CN-Learn as true that intersected with the validations (True positives only).

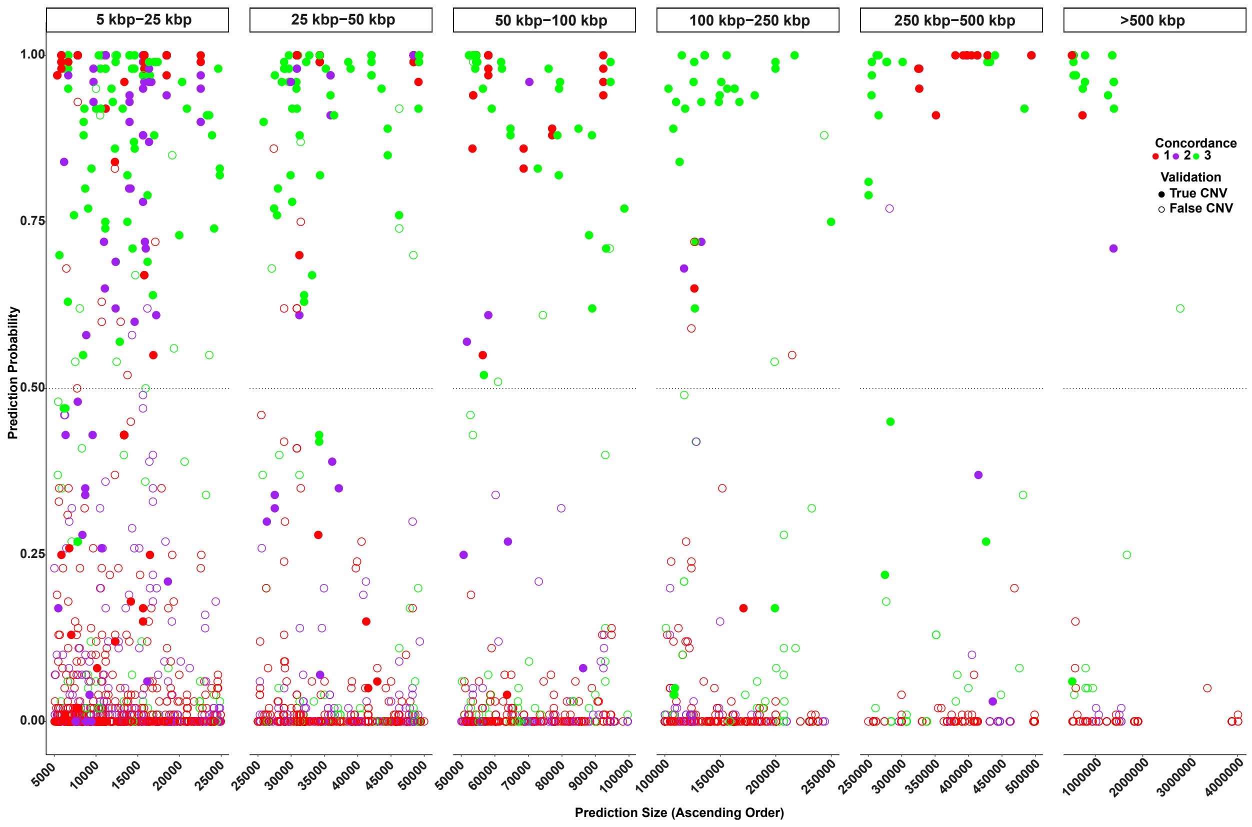

**Supplemental Figure S17: Distribution of CNVs in 453 test samples classified by CN-Learn as true, based on probability scores predicted by CN-Learn across six size ranges.** The level of concordance with one, two or three callers for each CNV is represented by different colors. CNVs that overlapped with predictions made by CLAMMS are indicated by the filled circles, while CNVs that did not overlap with CLAMMS predictions are represented by hollow circles.

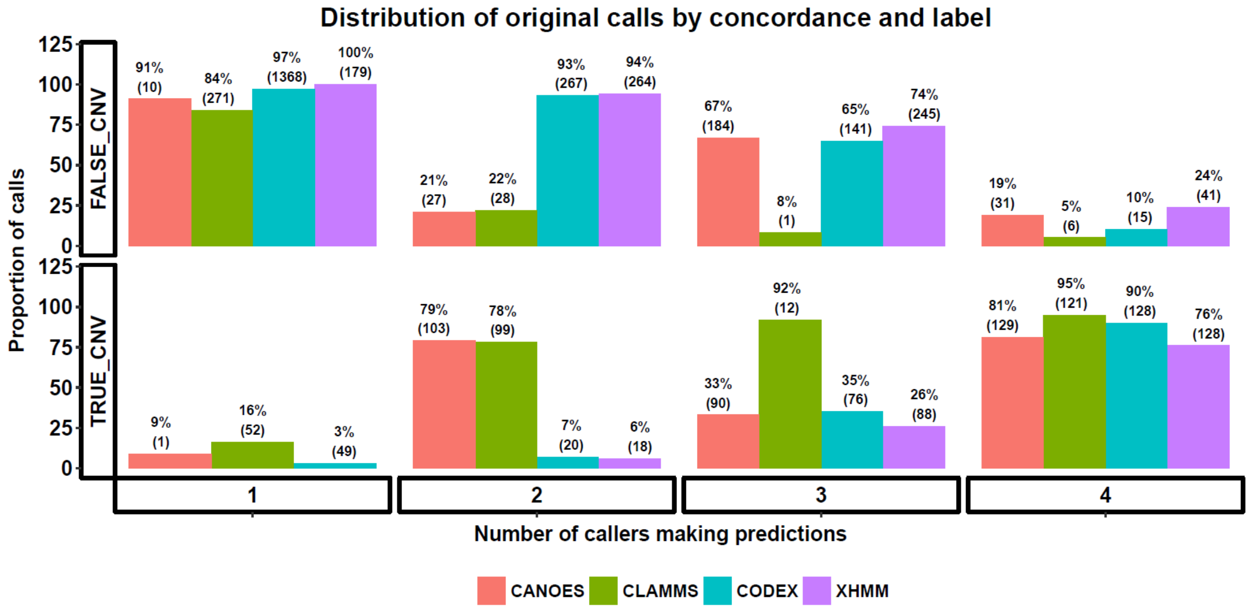

**Supplemental Figure S18: Characterization of CNVs predicted by four exome CNV callers in 291 samples based on concordance and validation labels.** Prior to pre-processing, the overlaps among the initial CNV predictions made by the four exome CNV callers and with validated microarray predictions were measured. Calls were labeled as "True" if their overlap with microarray calls was at least 10%, while the remaining predictions were labeled as "False". Each CNV prediction is represented in one of 32 bins based on the algorithm that made the prediction, its concordance with the other callers, and its overlap with microarray calls. The percentages illustrated in the bar graph indicate the fraction of CNVs that are split between "True" or "False" labels for each concordance bin.

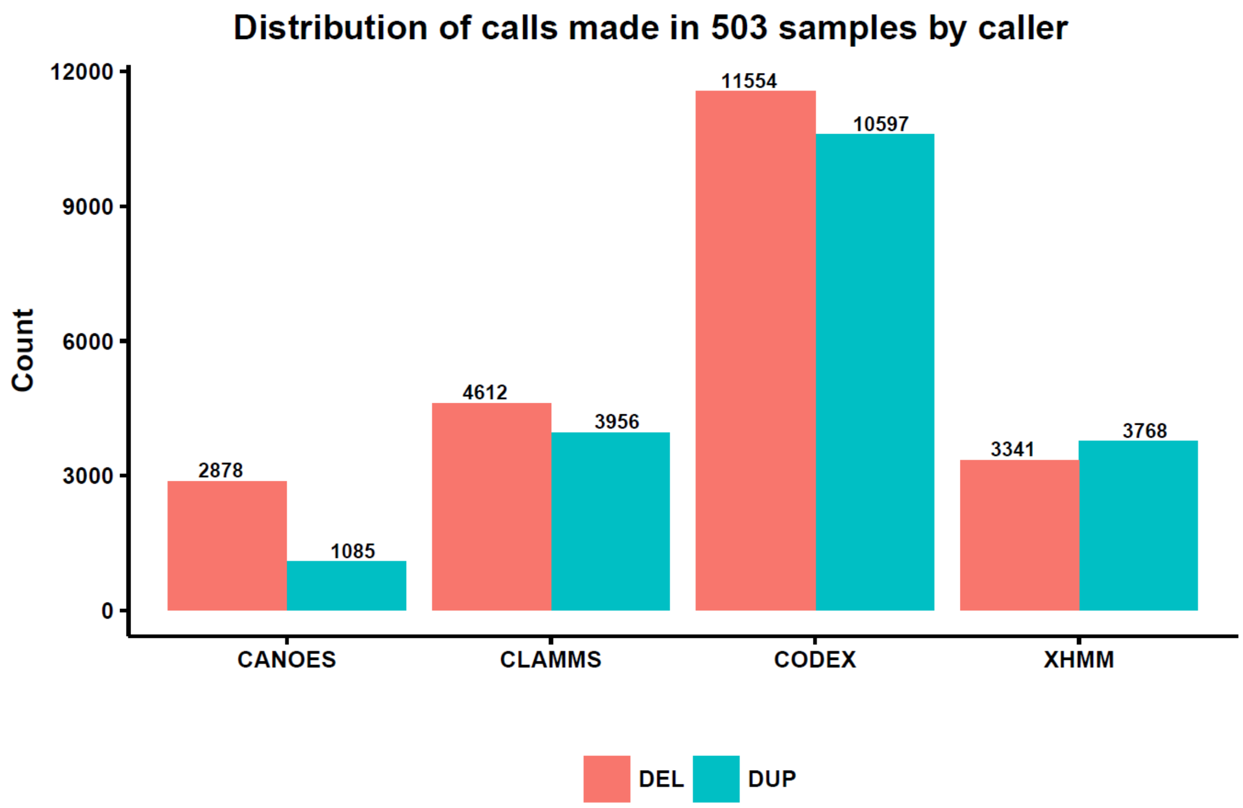

**Supplemental Figure S19: Number of duplications and deletions identified by four exome CNV callers in 503 samples.** Exome CNV callers are listed along the x-axis and the number of variants are listed along the Y-axis. The red bars illustrate the number of deletions, while the blue bars illustrate duplications. Although the distribution of CNVs varied widely among the callers, the general trend of variation within each caller was similar between deletions and duplications.

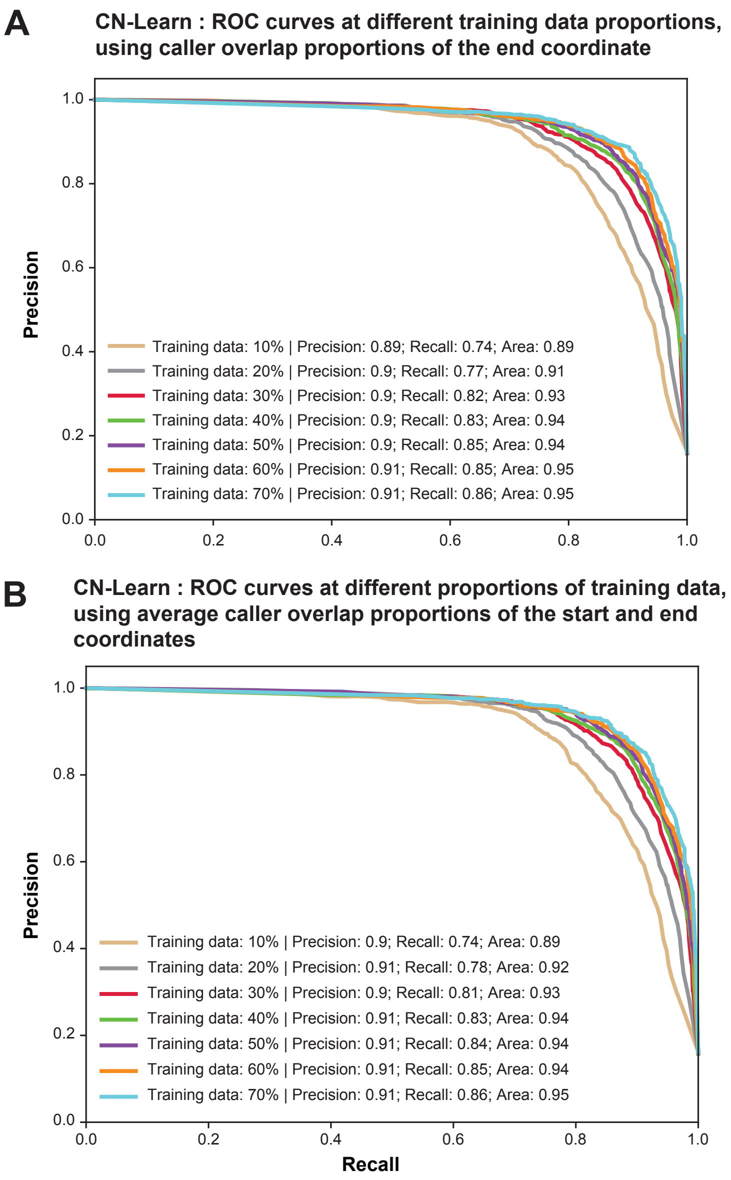

**Supplemental Figure S20: Performance of CN-Learn while using different methods for selecting overlap proportions in breakpoint resolution.** These plots show the performance of CN-Learn while using overlap proportions extracted for the caller whose end coordinate was chosen as the coordinate of the CNV event (A) and while using the average of overlap proportions extracted for the two callers whose start and end coordinates were chosen as the coordinates of the CNV event (B). Each ROC curve indicates the performance (precision and recall) obtained while using microarray calls as the truth set and different proportions of samples to train CN-Learn, from 10% to 70% in increments of 10%. The results shown are averages across 10-fold cross validation.

### SUPPLEMENTAL TABLES

**Supplemental Table 1: Spearman rank correlation among ten quantitative predictors supplied to CN-Learn for classification.**

| | CANOES | CODEX | CLAMMS | XHMM | Concordance | Pred. size | Target count | GC content | Mappability | RD ratio |
|---|---|---|---|---|---|---|---|---|---|---|
| **CANOES** | 1 | 0.05 | 0.06 | 0.39 | 0.65 | 0.1 | 0.26 | 0.12 | -0.1 | -0.15 |
| **CODEX** | 0.05 | 1 | -0.48 | -0.04 | 0.28 | 0.08 | -0.06 | 0.16 | -0.23 | 0.06 |
| **CLAMMS** | 0.06 | -0.48 | 1 | -0.16 | 0.19 | -0.18 | -0.12 | -0.18 | 0.48 | -0.08 |
| **XHMM** | 0.39 | -0.04 | -0.16 | 1 | 0.62 | 0.05 | 0.28 | 0.15 | -0.18 | 0.07 |
| **Concordance** | 0.65 | 0.28 | 0.19 | 0.62 | 1 | 0.01 | 0.17 | 0.15 | -0.04 | -0.04 |
| **Pred size** | 0.1 | 0.08 | -0.18 | 0.05 | 0.01 | 1 | 0.55 | -0.23 | -0.48 | 0.01 |
| **Target count** | 0.26 | -0.06 | -0.12 | 0.28 | 0.17 | 0.55 | 1 | 0.12 | -0.5 | -0.06 |
| **GC content** | 0.12 | 0.16 | -0.18 | 0.15 | 0.15 | -0.23 | 0.12 | 1 | -0.05 | 0.04 |
| **Mappability** | -0.1 | -0.23 | 0.48 | -0.18 | -0.04 | -0.48 | -0.5 | -0.05 | 1 | -0.01 |
| **RD ratio** | -0.15 | 0.06 | -0.08 | 0.07 | -0.04 | 0.01 | -0.06 | 0.04 | -0.01 | 1 |

**Supplemental Table 2: Proportion of predictions labeled as "True" at various overlap proportion cutoffs with microarray validated CNVs.**

| Overlap proportion | True Label CNV count | False Label CNV count | True Label proportion |
|---|---|---|---|
| 2% | 914 | 7171 | 11.3 |
| 3% | 913 | 7172 | 11.3 |
| 5% | 911 | 7174 | 11.3 |
| **10**% | **899** | **7186** | **11.1** |
| 20% | 877 | 7208 | 10.8 |
| 30% | 843 | 7242 | 10.4 |
| 40% | 813 | 7272 | 10.1 |
| 50% | 786 | 7299 | 9.7 |
| 60% | 726 | 7359 | 9 |
| 70% | 690 | 7395 | 8.5 |
| 80% | 637 | 7448 | 7.9 |
| 90% | 569 | 7516 | 7 |

**Supplemental Table 3: Performance indicators for individual CNV callers, combinations of callers, and CN-Learn.**

| Callers | Predicted CNVs | True predictions (compared to validated set) | Precision (%) | Recall (%) |
|---|---|---|---|---|
| CANOES | 399 | 121 | 30.33 | 5.33 |
| CODEX | 2415 | 441 | 18.26 | 19.43 |
| CLAMMS | 366 | 151 | 41.26 | 6.65 |
| XHMM | 593 | 120 | 20.24 | 5.29 |
| **Pairs of callers** | | | | |
| CANOES + CODEX | 286 | 114 | 39.86 | 5.02 |
| CANOES + CLAMMS | 79 | 54 | 68.35 | 2.38 |
| CANOES + XHMM | 178 | 64 | 35.96 | 2.82 |
| CODEX + CLAMMS | 172 | 120 | 69.77 | 5.29 |
| CODEX + XHMM | 381 | 114 | 29.92 | 5.02 |
| CLAMMS + XHMM | 74 | 43 | 58.11 | 1.89 |
| Two callers | 402 | 122 | 30.35 | 5.37 |
| Two or more callers | 611 | 220 | 36.01 | 9.69 |
| **Combinations of three callers** | | | | |
| CANOES + CODEX + CLAMMS | 77 | 54 | 70.13 | 2.38 |
| CANOES + CODEX + XHMM | 160 | 64 | 40 | 2.82 |
| CODEX + CLAMMS + XHMM | 66 | 42 | 63.64 | 1.85 |
| CANOES + CLAMMS + XHMM | 47 | 31 | 65.96 | 1.37 |
| Three callers | 162 | 67 | 41.36 | 2.95 |
| Three or more callers | 162 | 67 | 41.36 | 2.95 |
| **All callers** | 47 | 31 | 65.96 | 1.37 |
| **CN-Learn** | 347 | 334 | 96.25 | 14.71 |

**Supplemental Table 4 (Excel file): Final list of validated CNVs for 1000 Genomes samples.**

**Supplemental Table 5 (Excel file): List of CNVs classified by CN-Learn as true independently using microarray and CLAMMS based validations.**

**Supplemental Table 6 (Excel file): List of CNVs classified by CN-Learn as false independently using microarray and CLAMMS based validations.**
