## Supplemental_Code for "A machine-learning approach for accurate detection of copy-number variants from exome sequencing": calc_valdata_overlap.sh

```
#!/bin/bash
################################################################################
# Script : calc_valdata_overlap.sh                                             #
# Author : Vijay Kumar                                                         #
# Date   : 4/5/2019                                                            #
#                                                                              #
# This script takes each breakpoint-resolved CNV and labels them based on      #
# their overlap with the set of validated CNVs.                                #
#                                                                              #
# Prereqs (Format= <FILE LOCATION> File Description):                          #
# 1) <DATA> File with breakpoint-resolved CNVs along with GC, Mappability and  #
#           exome capture target count information.                            #
# 2) <SOURCE> File with the list of validated CNVs. i.e., gold-standard CNVs   #
# 3) <SOURCE> File with dummy chromosome coordinates. This file is used to     #
#             make sure that CNVs predicted in samples without validated CNVs  #
#             are not ignored. Overlapping them with this file ensures it.     # 
#                                                                              #
# (c) 2019 - Vijay Kumar                                                       #
# Licenced under the GNU General Public License 3.0.                           #
################################################################################
echo "Job started on `hostname` at `date`"

source TBD/config.params

####################################################
# STEP 0: Declare directories, files and variables #
####################################################
# Files:
ALL_PRED_W_GC_MAP_TARG=${DATA_DIR}'final_preds_GC_Map_Targ.txt'


if [ ! -f ${VAL_DATA_FILE} ];
then
    echo "ERROR : The input file with the list of validated CNVs is unavailable.";
    echo "Please place this file in the SOURCE directory and try again."
    exit 1;
fi

if [ -f ${SAMPLE_LIST_TRAIN} ];
then
    echo "ERROR : The input file with the list of training samples is unavailable.";
    echo "Please place this file in the SOURCE directory and try again."
    exit 1;
fi

if [ -f ${SAMPLE_LIST_TEST} ];
then
    echo "ERROR : The input file with the list of testing samples is unavailable.";
    echo "Please place this file in the SOURCE directory and try again."
    exit 1;
fi

########################################################################
# STEP 1: Loop through twice for different 1) Samples and 2) CNV_TYPE  #
########################################################################
if [ -f ${DATA_DIR}overlap_w_valdata.txt ]; 
then
rm ${DATA_DIR}overlap_w_valdata.txt
touch ${DATA_DIR}overlap_w_valdata.txt
fi

for SAMPLE in `cat ${SAMPLE_LIST_TRAIN}`;  
do
for CNV_TYPE in "DUP" "DEL";
do 
cat ${ALL_PRED_W_GC_MAP_TARG} | grep -w ${SAMPLE} | grep ${CNV_TYPE} > ${PRED_DIR}${SAMPLE}_${CNV_TYPE}.txt;
cat ${VAL_DATA_FILE} | grep -w ${SAMPLE} | grep ${CNV_TYPE}  > ${VALD_DIR}${SAMPLE}_${CNV_TYPE}.txt; 

if [ -s ${PRED_DIR}${SAMPLE}_${CNV_TYPE}.txt ] && [ -s ${VALD_DIR}${SAMPLE}_${CNV_TYPE}.txt ];
then 
${DOCKER_COMMAND}${BEDTOOLS_DIR}intersectBed -wao -f ${VALDATA_OV_THRESHOLD} -a ${PRED_DIR}${SAMPLE}_${CNV_TYPE}.txt \
                                                                             -b ${VALD_DIR}${SAMPLE}_${CNV_TYPE}.txt \
                                                                             >> ${DATA_DIR}overlap_w_valdata.txt;
elif [ -s ${PRED_DIR}${SAMPLE}_${CNV_TYPE}.txt ] && [ ! -s ${VALD_DIR}${SAMPLE}_${CNV_TYPE}.txt ];
then
${DOCKER_COMMAND}${BEDTOOLS_DIR}intersectBed -wao -f ${VALDATA_OV_THRESHOLD} -a ${PRED_DIR}${SAMPLE}_${CNV_TYPE}.txt \
                                                                             -b ${SOURCE_DIR}dummy.bed \
                                                                             >> ${DATA_DIR}overlap_w_valdata.txt;
fi

done
done

########################################################################
# STEP 2: Filter samples without validations into a separate test file #
########################################################################
if [ -f ${DATA_DIR}test_data_temp.txt ];
then
rm ${DATA_DIR}test_data_temp.txt
fi

for SAMPLE in `cat ${SAMPLE_LIST_TEST}`;
do
cat ${ALL_PRED_W_GC_MAP_TARG} | grep -w ${SAMPLE} >> ${DATA_DIR}test_data_temp.txt
done

##############################################################################################
# STEP 3: Loop through the files to group predictions and append labels and prediction sizes #
##############################################################################################
${DOCKER_COMMAND}Rscript ${RSCRIPTS_DIR}consolidate_val_ov.r  ${DATA_DIR}  overlap_w_valdata.txt \
        test_data_temp.txt  training_data.txt  test_data.txt  ${CALLER_COUNT}  ${CALLER_LIST}

echo "Job ended on `hostname` at `date`"
```
