## Supplemental_Code for "A machine-learning approach for accurate detection of copy-number variants from exome sequencing": calculate_CNV_overlap.sh

```
#!/bin/bash
################################################################################
# Script : calculate_CNV_overlap.sh                                            #
# Author : Vijay Kumar                                                         #
# Date   : 4/5/2019                                                            #
#                                                                              #
# This script calculates the overlap between CNVs predicted by multiple        #
# callers. Once the overlaps are measured, data is reshaped, and the number    #
# of targets each CNV overlaps with is then calculated. This script is written #                      
# in such a way that it is restartable.                                        # 
#                                                                              #
# Prereqs (Format= <FILE LOCATION> File Description):                          #
# 1) <DATA> File with the consolidated set of CNVs from multiple callers       #
# 2) <SOURCE> File with the list of all sample names                           #
#                                                                              #
# (c) 2019 - Vijay Kumar	                                               #
# Licenced under the GNU General Public License 3.0.                           #
################################################################################
echo "Job started on `hostname` at `date`"

source TBD/config.params

####################################################
# STEP 0: Declare directories, files and variables #
####################################################
# Files:
CONS_PRED_FILE_NAME='consolidated_calls.bed'
CONS_PRED_W_OV_FILE_NAME='cons_calls_w_caller_ov.bed'
CONS_PRED_W_OV_PROP_FILE_NAME='cons_calls_w_caller_ov_prop.bed'

CONS_PRED_FILE=${DATA_DIR}${CONS_PRED_FILE_NAME}
CONS_PRED_W_VAL_FILE=${DATA_DIR}${CONS_PRED_W_VALS_FILE_NAME}
CONS_PRED_W_OV_FILE=${DATA_DIR}${CONS_PRED_W_OV_FILE_NAME}

###################################################################################
# Step 0: Merge the input CNV call files from multiple callers into a single file #
###################################################################################
${DOCKER_COMMAND}Rscript --vanilla ${RSCRIPTS_DIR}merge_init_preds.r ${DATA_DIR} ${CONS_PRED_FILE_NAME}

################################################################################
# Step 1: Make sure the data/format of the input file is consistent & accurate #
################################################################################
cnv_type_count=`cat ${CONS_PRED_FILE} | cut -f4 | sort | uniq | wc -l`

if [ ${cnv_type_count} -eq 2 ];
then
    cnv_type_1=`cat ${CONS_PRED_FILE} | cut -f4 | sort | uniq | sort | head -1`
    cnv_type_2=`cat ${CONS_PRED_FILE} | cut -f4 | sort | uniq | sort | tail -1`

    if [ ${cnv_type_1} != "DEL" ] || [ ${cnv_type_2} != "DUP" ];
    then
        echo "ERROR : Only DUP and DEL are acceptable values for CNV type. 
        Please check the fifth column in the input file for data integrity";
        exit 1;
    fi

else
    echo "ERROR : Only two types of CNVs can be analyzed. The input file has either 
    more or less than two CNV types. Please check the input file for data integrity";
    exit 1;
fi

num_of_callers=`cat ${CONS_PRED_FILE} | cut -f6 | sort | uniq | wc -l`
caller_list_input=`cat ${CONS_PRED_FILE} | cut -f6 | sort | uniq | sort`

if [ ${num_of_callers} -eq ${CALLER_COUNT} ];
then 
    caller_num=1
    for caller in ${CALLER_LIST};
    do
        caller_input=`echo ${caller_list_input} | cut -d' ' -f${caller_num}`
        if [ ${caller} != ${caller_input} ];
        then
            echo "ERROR : Mismatch in the caller names between the input file and ";
            echo "the names supplied in the config.params file ";
            exit 1;
        fi
    let "caller_num+=1"
    done
else
    echo "ERROR : Mismatch in the number of callers between the input file and ";
    echo "the number of callers supplied in the config.params file ";
    exit 1;
fi

if [ ! -f ${SAMPLE_LIST} ];
then
    echo "ERROR : The input file with the list of samples to process is unavailable.";
    echo "Please ensure the presence of this file in the SOURCE directory and try again."
    exit 1;
fi

################################################################################
# STEP 2: Run bedtools to identify predictions between callers that overlap.   #
#         Three nested loops to process  1) Caller 2) Sample 3) CNV Type.      #
################################################################################
if [ -f ${CONS_PRED_W_OV_FILE} ];
then
rm ${CONS_PRED_W_OV_FILE}
fi
touch ${CONS_PRED_W_OV_FILE}


for caller in ${CALLER_LIST};
do

if [ -f ${DATA_DIR}${caller}_caller_ov.txt ];
then
rm ${DATA_DIR}${caller}_caller_ov.txt
fi
touch ${DATA_DIR}${caller}_caller_ov.txt

for sample in `cat ${SAMPLE_LIST}`; 
do
for cnv_type in "DUP" "DEL";
do 
cat ${CONS_PRED_FILE} | grep ${caller} | grep -w ${sample} | grep ${cnv_type} \
                             > ${PRED_DIR}${caller}_${sample}_${cnv_type}.txt; 
cat ${CONS_PRED_FILE} | grep -w ${sample} | grep ${cnv_type} \
                  > ${PRED_DIR}${caller}_complement_${sample}_${cnv_type}.txt; 

if [ -s ${PRED_DIR}${caller}_${sample}_${cnv_type}.txt ] && \
   [ -s ${PRED_DIR}${caller}_complement_${sample}_${cnv_type}.txt ];
then 
${DOCKER_COMMAND}${BEDTOOLS_DIR}intersectBed -wao -a ${PRED_DIR}${caller}_${sample}_${cnv_type}.txt \
                                     -b ${PRED_DIR}${caller}_complement_${sample}_${cnv_type}.txt \
                                     | cut -f1-6,12,13 >> ${DATA_DIR}${caller}_caller_ov.txt;

elif [ -s ${PRED_DIR}${caller}_${sample}_${cnv_type}.txt ] && \
     [ ! -s ${PRED_DIR}${caller}_complement_${sample}_${cnv_type}.txt ];
then
${DOCKER_COMMAND}${BEDTOOLS_DIR}intersectBed -wao -a ${PRED_DIR}${caller}_${sample}_${cnv_type}.txt \
                                     -b ${SOURCE_DIR}dummy.bed | cut -f1-6,12,13 \
                                     >> ${DATA_DIR}${caller}_caller_ov.txt;
fi
done
done
cat ${DATA_DIR}${caller}_caller_ov.txt >> ${CONS_PRED_W_OV_FILE}
done  

################################################################################
# STEP 3: Run the R script to reshape the overlap info from rows to columns.   #
################################################################################
${DOCKER_COMMAND}Rscript --vanilla ${RSCRIPTS_DIR}reshape_caller_overlap_data.r  \
                                   ${DATA_DIR} ${CONS_PRED_W_OV_FILE_NAME} \
                                   ${CONS_PRED_W_OV_PROP_FILE_NAME} \
                                   ${CALLER_COUNT} ${CALLER_LIST}

echo "Job ended on `hostname` at `date`"
```
