## Supplemental_Code for "A machine-learning approach for accurate detection of copy-number variants from exome sequencing": canoes_call_CNVs.sh

```
#!/bin/bash
####################################################################
# Script : canoes_call_CNVs.sh                                     #
# Author : Vijay Kumar                                             #
# Date   : 4/5/2019                                                #
#                                                                  #
# This script uses the read depth information extracted by the     #
# CANOES script canoes_extract.sh and uses the information from    #
# all the samples to predict CNVs.                                 #
#                                                                  #
# (c) 2019 - Vijay Kumar                                           #
# Licenced under the GNU General Public License 3.0.               #    
####################################################################
echo "Job started on `hostname` at `date`"

source TBD/config.params

############################################
# STEP 0: Declare variables and file names #
############################################
CANOES_READS_FILE='canoes_reads'
CONS_READS=${DATA_CANOES_DIR}'cons_canoes_reads' 

NUM_OF_PROBES=`wc -l < ${TARGET_PROBES}`

###################################################################################################
# STEP 1: Make sure the number of rows in each output file matches with the number of target probes 
###################################################################################################
cd ${DATA_CANOES_DIR}
split_file_list=`ls list_of_bam_split* | grep "^list_of_bam_split"`
cat ${TARGET_PROBES} > ${CONS_READS}
for split_file in ${split_file_list};
do
num_of_rows_input=`wc -l < ${DATA_CANOES_DIR}${CANOES_READS_FILE}_${split_file}`
if [ ${NUM_OF_PROBES} = ${num_of_rows_input} ];
then
cat ${DATA_CANOES_DIR}${CANOES_READS_FILE}_${split_file} | cut -f 4,5,6,7 \
                                            > ${DATA_CANOES_DIR}'temp_file'
paste ${CONS_READS} ${DATA_CANOES_DIR}'temp_file' > ${DATA_CANOES_DIR}'temp_file2'
mv ${DATA_CANOES_DIR}'temp_file2' ${CONS_READS}
else
echo "Error in probe counts in file ${DATA_CANOES_DIR}${CANOES_READS_FILE}_${split_file}"
fi
done

##############################################
# STEP 2: Extract GC content for each interval
##############################################
${DOCKER_COMMAND}java -Xmx2000m -Djava.io.tmpdir=${DATA_LOGS_DIR} \
                      -jar ${GATK_SW_DIR}GenomeAnalysisTK.jar \
                      -T GCContentByInterval -L ${TARGET_PROBES} \
                       -R ${REF_GENOME} -o ${DATA_CANOES_DIR}gc.txt

##############################################################################################
# STEP 3: Execute R script to merge data. This is needed because multicov command was executed 
#         for just four samples via separate jobs, to parallalize data extraction manually.
##############################################################################################
${DOCKER_COMMAND}Rscript --vanilla ${RSCRIPTS_DIR}canoes_merge_files.r \
                                   ${RSCRIPTS_DIR} ${DATA_CANOES_DIR} ${CONS_READS} canoes_calls.csv

###############################################################
# STEP 4: Copy the output files to the final output directory #
###############################################################
cp ${DATA_CANOES_DIR}canoes_calls.csv ${DATA_DIR}'canoes_calls.csv'

echo "Job ended on `hostname` at `date`"
```
