## Supplemental_Code for "A machine-learning approach for accurate detection of copy-number variants from exome sequencing": canoes_extract.sh

```
#!/bin/bash
####################################################################
# Script : canoes_extract.sh                                       #
# Author : Vijay Kumar                                             #
# Date   : 4/5/2019                                                #
#                                                                  #
# This script extracts the read depth information for subsequent   #
# analysis by the CANOES pipeline.                                 #
#                                                                  #
# IMPORTANT NOTE: This for loop that processes the samples can be  #
# easily modified to run in parallel on local clusters or on other #
# cloud infrastructures.                                           #
#                                                                  #
# (c) 2019 - Vijay Kumar                                           #
# Licenced under the GNU General Public License 3.0.               #    
####################################################################
echo "Job started on `hostname` at `date`"

source TBD/config.params

############################################
# STEP 0: Declare variables and file names #
############################################
CANOES_READS_FILE='canoes_reads'
CONS_READS=${DATA_CANOES_DIR}'cons_canoes_reads' 

NUM_OF_PROBES=`wc -l < ${TARGET_PROBES}`

######################################################
# STEP 1: Generate the file with the list of samples #
######################################################
echo 'SAMPLE' > ${DATA_CANOES_DIR}sample_seq
num_samples=`cat ${BAM_FILE_LIST_W_PATH} | wc -l`
for samp in $(seq 1 ${num_samples});
do
echo "S${samp}" >> ${DATA_CANOES_DIR}sample_seq
done

echo 'SAMPLE_NAME' > ${DATA_CANOES_DIR}sample_names
cat ${SAMPLE_LIST} >> ${DATA_CANOES_DIR}sample_names
paste -d , ${DATA_CANOES_DIR}sample_seq ${DATA_CANOES_DIR}sample_names \
                             > ${DATA_CANOES_DIR}sample_map_canoes.csv

cp ${DATA_CANOES_DIR}sample_map_canoes.csv ${DATA_DIR}sample_map_canoes.csv

#######################################################################################
# STEP 2: Extract Read Counts for each sample in a single step and submit separate jobs
#######################################################################################
touch ${DATA_CANOES_DIR}${CANOES_READS_FILE}
cd ${DATA_CANOES_DIR}
split -l 4 ${BAM_FILE_LIST_W_PATH} list_of_bam_split

split_file_list=`ls list_of_bam_split* | tail -n+2 | tail -1`

for split_file in ${split_file_list};
do
${DOCKER_COMMAND}${BEDTOOLS_DIR}bedtools multicov -bams `cat ${DATA_CANOES_DIR}${split_file} \
                                     | tr "\n" " "` -bed ${TARGET_PROBES} -q 20 \
                                     > ${DATA_CANOES_DIR}${CANOES_READS_FILE}_${split_file}
done

echo "Job ended on `hostname` at `date`"
```
