## Supplemental_Code for "A machine-learning approach for accurate detection of copy-number variants from exome sequencing": clamms_extract.sh

```
#!/bin/bash
########################################################################
# Script : clamms_extract.sh                                           #
# Author : Vijay Kumar                                                 #
# Date   : 4/5/2019                                                    #
#                                                                      #
# This script uses samtools to extract read depth information for the  #
# windows generated in the prior script clamms_preprocess.sh. This     #
# task is isolated from other steps in the CLAMMS pipeline to make it  #
# easier to parallelize.                                               #
#                                                                      #
# IMPORTANT NOTE: This for loop that processes the samples can be      #
# easily modified to run in parallel on local clusters or on other     #
# cloud infrastructures.                                               #
#                                                                      #
# (c) 2019 - Vijay Kumar                                               #
# Licenced under the GNU General Public License 3.0.                   #
########################################################################
echo "Job started on `hostname` at `date`"

source TBD/config.params

####################################################
# STEP 0: Declare variables, directories and files #
####################################################
export INSERT_SIZE=200

##################################################################
# STEP 1: Calculate actual and normalized coverage for each sample 
##################################################################
for bam_file in `cat ${BAM_FILE_LIST_W_PATH} | head -1`;
do

echo ${bam_file}
sample_name=`echo ${bam_file} | rev | cut -f1 -d/ | rev`

${DOCKER_COMMAND}${SAMTOOLS_DIR}samtools bedcov -Q 30 ${DATA_CLAMMS_DIR}windows.bed ${bam_file} | \
                                  awk '{ printf "%s\t%d\t%d\t%.6g\n", $1, $2, $3, $NF/($3-$2); }' \
                                                 > ${DATA_CLAMMS_DIR}${sample_name}.coverage.bed

${DOCKER_COMMAND}${CLAMMS_DIR}normalize_coverage ${DATA_CLAMMS_DIR}${sample_name}.coverage.bed \
                                                 ${DATA_CLAMMS_DIR}windows.bed \
                                                 > ${DATA_CLAMMS_DIR}${sample_name}.norm.cov.bed
done

echo "Job ended on `hostname` at `date`"
```
