## Supplemental_Code for "A machine-learning approach for accurate detection of copy-number variants from exome sequencing": clamms_postprocess.sh

```
#!/bin/bash
########################################################################
# Script : clamms_postprocess.sh                                       #
# Author : Vijay Kumar                                                 #
# Date   : 4/5/2019                                                    #
#                                                                      #
# This script is part of the CLAMMS pipeline that uses the read depth  #
# information extracted by the script clamms_extract.sh for all        #
# samples and predicts CNVs.                                           #
#                                                                      #
# (c) 2019 - Vijay Kumar                                               #
# Licenced under the GNU General Public License 3.0.                   #
########################################################################
echo "Job started on `hostname` at `date`"

source TBD/config.params

####################################################
# STEP 0: Declare variables, directories and files #
####################################################
export INSERT_SIZE=200

#####################################################################################
# STEP 1: Combine normalized coverage into a single panel prior to fitting the model
#####################################################################################
if [ -f ${DATA_CLAMMS_DIR}clamms_cnvs.bed ];
then
rm ${DATA_CLAMMS_DIR}clamms_cnvs.bed
fi
touch ${DATA_CLAMMS_DIR}clamms_cnvs.bed

sample_name=${SAMPLE_LIST}
ls ${DATA_CLAMMS_DIR}*norm.cov.bed | grep -f ${sample_name} | while read FILE;
do
    echo "${FILE}"
    #grep "^Y" $FILE | awk '{ x += $4; n++; } END { if (x/(n+1) >= 0.1) print "M"; else print "F"; }'
done > ${DATA_CLAMMS_DIR}/ref.panel.files.txt

${DOCKER_COMMAND}${CLAMMS_DIR}fit_models ${DATA_CLAMMS_DIR}ref.panel.files.txt ${DATA_CLAMMS_DIR}windows.bed \
                                                                > ${DATA_CLAMMS_DIR}models.bed

#####################################################################################
# STEP 2: Use the model file and normalized coverage file to make CNV predictions
#####################################################################################
cd ${DATA_CLAMMS_DIR}
sample_name=${SAMPLE_LIST}
for norm_cov_file in `ls *norm.cov.bed | grep -f ${sample_name}`;
do 
${DOCKER_COMMAND}${CLAMMS_DIR}call_cnv ${DATA_CLAMMS_DIR}${norm_cov_file} ${DATA_CLAMMS_DIR}models.bed \
                                                       >> ${DATA_CLAMMS_DIR}clamms_cnvs.bed
done

cp ${DATA_CLAMMS_DIR}clamms_cnvs.bed ${DATA_DIR}clamms_calls.txt

echo "Job ended on `hostname` at `date`"
```
