## Supplemental_Code for "A machine-learning approach for accurate detection of copy-number variants from exome sequencing": clamms_preprocess.sh

```
#!/bin/bash
########################################################################
# Script : clamms_preprocess.sh                                        #
# Author : Vijay Kumar                                                 #
# Date   : 4/5/2019                                                    #
#                                                                      #
# This script is part of the CLAMMS pipeline. It downloads genome      #
# mappability file and uses it with the exome capture target probe     #
# list to produce a custom set of target probe list for subsequent     #
# analyses.                                                            #
#                                                                      #
# (c) 2019 - Vijay Kumar                                               #
# Licenced under the GNU General Public License 3.0.                   #
########################################################################
echo "Job started on `hostname` at `date`"

source TBD/config.params

####################################################
# STEP 0: Declare variables, directories and files #
####################################################
export INSERT_SIZE=200

#################################################################################
# STEP 1: Convert bigwig to wig file to obtain mappability scores               # 
#################################################################################
cd ${SOURCE_DIR}
if [ ! -f wgEncodeCrgMapabilityAlign75mer.bigWig ];
then
wget http://hgdownload.cse.ucsc.edu/goldenPath/hg19/encodeDCC/wgEncodeMapability/wgEncodeCrgMapabilityAlign75mer.bigWig
fi

${DOCKER_COMMAND}${SW_DIR}bigWigToWig ${SOURCE_DIR}wgEncodeCrgMapabilityAlign75mer.bigWig \
                                 ${SOURCE_DIR}wgEncodeCrgMapabilityAlign75mer.wig

grep -v '^#' ${SOURCE_DIR}wgEncodeCrgMapabilityAlign75mer.wig | sed 's/^chr//g' | sort -k1,1 -k2,2n \
                                   > ${DATA_CLAMMS_DIR}mappability.bed

#################################################################################
# STEP 2: Sort the target probe file and the mappability bed file               #
#################################################################################
sort -k1,1 -k2,2n ${TARGET_PROBES} > ${DATA_CLAMMS_DIR}sorted_targets.bed
sort -k1,1 -k2,2n ${DATA_CLAMMS_DIR}mappability.bed > ${DATA_CLAMMS_DIR}mappability_sorted.bed

${DOCKER_COMMAND}sort -k1,1 -k2,2n ${CLAMMS_DIR}data/clamms_special_regions.bed \
                      > ${DATA_CLAMMS_DIR}clamms_special_regions_sorted.bed


#accesses the special_regions.bed in CLAMMS
${DOCKER_COMMAND}${CLAMMS_DIR}annotate_windows.sh ${DATA_CLAMMS_DIR}sorted_targets.bed ${REF_GENOME} \
                                                  ${DATA_CLAMMS_DIR}mappability_sorted.bed $INSERT_SIZE \
                                                  ${DATA_CLAMMS_DIR}clamms_special_regions_sorted.bed \
                                                              > ${DATA_CLAMMS_DIR}windows.bed

echo "Job ended on `hostname` at `date`"
```
