## Supplemental_Code for "A machine-learning approach for accurate detection of copy-number variants from exome sequencing": cn_learn.py

```
################################################################################
# Script : cn_learn.py                                                         # 
# Author : Vijay Kumar                                                         #
# Date   : 4/5/2019                                                            #
#                                                                              #
# This script takes a list of labelled CNVs as input to train CN-Learn. It     #
# takes several input parameters from the bash script to make predictions on   #
# CNVs predicted from new samples.                                             #
#                                                                              #
# (c) 2019 - Vijay Kumar                                                       #
# Licenced under the GNU General Public License 3.0.                           #
################################################################################
from __future__ import division
import sys
import re
import os
import csv   
import pandas as pd
import numpy as np
import sklearn as sk

from sklearn import preprocessing
from sklearn.preprocessing import MinMaxScaler
from sklearn.calibration import calibration_curve
from sklearn.ensemble import RandomForestClassifier
from sklearn import svm
from sklearn.svm import SVC
from sklearn.linear_model import LogisticRegression
from inspect import getmembers
from scipy import interp
from sklearn import tree
from sklearn.externals import six
from itertools import cycle

#############################################################
# Define a class for the binary classifier with predictors, #
# response variables and additional parameters. This class  #
# has several methods that can be used to train, test and   #
# evaluate the performance of the classifier.               #
#############################################################
class BinaryClassifier(object):

    def __init__(self, X, Y, test_data, classifier_name, num_trees):
        self.X = X
        self.Y = Y
        self.test = test_data
        self.classifier_name = classifier_name
        self.num_trees = num_trees    

    ######################################################
    # Method 1: Build the classifier using training data #
    ######################################################
    def run_classifier(self):
        y, _ = pd.factorize(self.X['TYPE_IND'])
        y, _ = pd.factorize(self.X['CHR'])
        y, _ = pd.factorize(self.X['SIZE_LABEL'])

        if self.classifier_name == 'RF':    
            clf = RandomForestClassifier(n_estimators=self.num_trees, n_jobs = -1)
            clf = clf.fit(self.X, self.Y)

        elif self.classifier_name == 'SVM':
            scaling = MinMaxScaler(feature_range=(-1,1)).fit(self.X)
            X_train = scaling.transform(self.X)
            clf = svm.SVC(probability=True,kernel='linear')
            clf = clf.fit(X_train, self.Y)

        elif self.classifier_name == 'LR':
            clf = LogisticRegression()
            clf = clf.fit(self.X, self.Y)

        return clf

    #########################################################
    # Method 2: Use the classifier to predict probabilities #
    #########################################################
    def predict_prob(self, clf):
        self.clf = clf

        if self.classifier_name == 'RF':
            pred_probs = self.clf.predict_proba(self.test)
        elif self.classifier_name == 'SVM':
            scaling = MinMaxScaler(feature_range=(-1,1)).fit(self.test)
            X_test     = scaling.transform(self.test)
            pred_probs = self.clf.predict_proba(X_test)
        elif self.classifier_name == 'LR':
            pred_probs = self.clf.predict_proba(self.test)

        return pred_probs
 
#############################
# Main function starts here #
#############################
def main():
    parameter_count = len(sys.argv) - 1
    print(parameter_count)
    caller_count = int(sys.argv[8])
    if caller_count <=0:
        print('Number of callers used must be greater than 1')
        sys.exit(0)
    else:
        expected_parm_count = 8 + caller_count
    if parameter_count != expected_parm_count:
        error_message =  """Please make sure that all the required parameters are supplied to CN-Learn. 
                            Usage: cn_learn.py <DATA_DIRECTORY> <TRAINING_DATA> <TEST_DATA> <CLASSIFIER_TYPE> 
                                               <LOWER_SIZE_LIMIT> <UPPER_SIZE_LIMIT> <NUMBER_OF_TREES> 
                                               <CALLER_COUNT> <CALLER_LIST>"""
        print(error_message)
        sys.exit(0)
    Source_Path = sys.argv[1]
    training_data = sys.argv[2]
    test_data = sys.argv[3]
    classifier_name = str(sys.argv[4])
    lower_size_limit = int(sys.argv[5])
    upper_size_limit = int(sys.argv[6])
    num_trees = int(sys.argv[7])
    caller_count = int(sys.argv[8])
    caller_list = []
    for caller_number in range(1, caller_count + 1):
        caller_list.append(sys.argv[8 + caller_number])
    print(caller_list)

    ############################################
    # STEP 1: Declare all the global variables #
    ############################################
    size_dict = {'A)<1KB':1, 'B)1KB-5KB':2, 'C)5KB-10KB':3, 'D)10KB-25KB':4, 'E)25KB-50KB':5, 'F)50KB-75KB':6,
             'G)75KB-100KB':7, 'H)100KB-250KB':8, 'I)250KB-500KB':9, 'J)500KB-1MB':10, 'K)1MB-5MB':11, 'L)>5MB':12}

    ############################################################################
    # STEP 2: Read the input files with the list of CNVs, select calls between # 
    #         the preferred size threshold and format the generated dataframe  #
    ############################################################################
    train_df_temp = pd.read_table(Source_Path + training_data)
    test_df_temp = pd.read_table(Source_Path + test_data)
    
    train_df = train_df_temp[(train_df_temp["PRED_SIZE"] >= lower_size_limit) & (train_df_temp["PRED_SIZE"] <= upper_size_limit)]
    train_df.loc[:,'TYPE_IND'] = [1 if x == 'DEL' else 2 for x in train_df['TYPE']]
    train_df.loc[:,'SIZE_LABEL'] = [size_dict.get(x) for x in train_df['SIZE_LABEL']]

    test_df = test_df_temp[(test_df_temp["PRED_SIZE"] >= lower_size_limit) & (test_df_temp["PRED_SIZE"] <= upper_size_limit)]
    test_df.loc[:,'TYPE_IND'] = [1 if x == 'DEL' else 2 for x in test_df['TYPE']]
    test_df.loc[:,'SIZE_LABEL'] = [size_dict.get(x) for x in test_df['SIZE_LABEL']]

    print(train_df.shape)
    print(test_df.shape)

    X_train = pd.DataFrame(train_df, columns = ['CHR', 'TYPE_IND', 'NUM_OVERLAPS', 'RD_PROP', 'GC', 'MAP', 'NUM_TARGETS', 'SIZE_LABEL'] + caller_list)
    X_test = pd.DataFrame(test_df, columns = ['CHR', 'TYPE_IND', 'NUM_OVERLAPS', 'RD_PROP', 'GC', 'MAP', 'NUM_TARGETS', 'SIZE_LABEL'] + caller_list)
    X_test_all_cols = pd.DataFrame(test_df, columns = ['CHR', 'PRED_START', 'PRED_END', 'TYPE', 'SAMPLE', 'NUM_OVERLAPS', 'RD_PROP', \
                          'GC', 'PRED_SIZE', 'MAP', 'NUM_TARGETS', 'SIZE_LABEL'] + caller_list)
    Y = train_df["LABEL_VAL"]

    ####################################################################
    # STEP 3: Create an instance of the Binary Classifier and make predictions #
    ####################################################################
    clf = BinaryClassifier(X_train, Y, X_test, classifier_name, num_trees)
    clf_out  = clf.run_classifier()
    pred_probs = clf.predict_prob(clf_out)
            
    ###########################################################
    # Assign CNVs to a class based on predicted probabilities #
    ###########################################################
    true_lab_prob = [float(item[1]) for item in pred_probs]
    true_lab_pred = [1 if lab > 0.5 else 0 for lab in true_lab_prob]
    X_test_all_cols.loc[:,"PRED_PROBS"] = true_lab_prob
    X_test_all_cols.loc[:,"PRED_LABEL"] = true_lab_pred

    print(X_test_all_cols.head)

    #################################
    # STEP 5: Write the output file #
    #################################
    X_test_all_cols.to_csv(Source_Path + 'CNV_list_with_predictions.csv', index = False)   
 
if __name__ == '__main__':
    main()
```
