## Supplemental_Code for "A machine-learning approach for accurate detection of copy-number variants from exome sequencing": cn_learn.sh

```
#!/bin/bash
################################################################################
# Script : cn_learn.sh                                                         # 
# Author : Vijay Kumar                                                         #
# Date   : 4/5/2019                                                            #
#                                                                              #
# This script executes the python script that takes several parameters to      #
# build CN-Learn. The parameters include classifier type (RF, SVM, LR),        #
# number of trees, number of splits etc                                        #
#                                                                              # 
# (c) 2019 - Vijay Kumar                                                       #
# Licenced under the GNU General Public License 3.0.                           #
################################################################################
echo "Job started on `hostname` at `date`"

source TBD/config.params

#########################################
# STEP 0: Declare variables required    #
#########################################
LOWER_SIZE_LIMIT=50000
UPPER_SIZE_LIMIT=5000000

##########################################
# STEP 2: Run CN-Learn for each strategy #
##########################################
${DOCKER_COMMAND}python3 -u ${SCRIPTS_DIR}cn_learn.py  ${DATA_DIR}  training_data.txt  test_data.txt \
                            ${CLASSIFIER}  ${LOWER_SIZE_LIMIT}  ${UPPER_SIZE_LIMIT}  ${NUM_TREES} \
                            ${CALLER_COUNT}  ${CALLER_LIST}

echo "Job ended on `hostname` at `date`"
```
