## Supplemental_Code for "A machine-learning approach for accurate detection of copy-number variants from exome sequencing": codex_extract.sh

```
#!/bin/bash
####################################################################
# Script : codex_extract.sh                                        #
# Author : Vijay Kumar                                             #
# Date   : 4/5/2019                                                #
#                                                                  #
# This CODEX script extracts and analyzes the read depth from all  #
# the samples and subsequently predicts CNVs. This script analyzes #
# one chromosome at a time, but can be parallelized.               #
#                                                                  #
# IMPORTANT NOTE: The for loop that processes chromosomes can be   #
# easily modified to run in parallel on local clusters or on other #
# cloud infrastructures.                                           #
#                                                                  #
# (c) 2019 - Vijay Kumar	                                   #
# Licenced under the GNU General Public License 3.0.               #
####################################################################
echo "Job started on `hostname` at `date`"

source TBD/config.params

PROJ_NAME='cnlearn'
CODEX_OUTPUT_FILE='codex_calls.txt'

for chr_num in {1..22};do
${DOCKER_COMMAND}Rscript ${RSCRIPTS_DIR}codex.r ${chr_num} ${BAM_FILE_LIST_W_PATH} \
                         ${BAM_FILE_LIST} ${TARGET_PROBES} ${PROJ_NAME} ${DATA_CODEX_DIR}
done

echo "Job ended on `hostname` at `date`"
```
