## Supplemental_Code for "A machine-learning approach for accurate detection of copy-number variants from exome sequencing": codex_postprocess.sh

```
#!/bin/bash
##################################################################
# Script : codex_postprocess.sh                                  # 
# Author : Vijay Kumar                                           #
# Date   : 4/5/2019                                              #
#                                                                #
# This is a helper script that needs to be run right after the   #
# successful execution of codex_extract.sh. This script simply   #
# combines the output files generated by the prior script and    #
# moves the final output with the list of predicted CNVs to      #
# the data directory for subsequent processing.                  # 
#                                                                #
# (c) 2019 - Vijay Kumar	                                 #
# Licenced under the GNU General Public License 3.0.             #
##################################################################
echo "Job started on `hostname` at `date`"

source TBD/config.params

PROJ_NAME='cnlearn'
CODEX_OUTPUT_FILE='codex_calls.txt'

########################################################
# STEP 1: Merge output files while managing headers    #
########################################################
if [ -f ${DATA_CODEX_DIR}${CODEX_OUTPUT_FILE} ];
then
rm ${DATA_CODEX_DIR}${CODEX_OUTPUT_FILE}
fi

first_file="yes"
for codex_file in ${DATA_CODEX_DIR}${PROJ_NAME}*;
do
        if [ ${first_file} = "yes" ]; 
        then
                echo "$codex_file copying"
                cp ${codex_file} ${DATA_CODEX_DIR}${CODEX_OUTPUT_FILE}
                first_file="no"
        else
                echo "${codex_file} omit first line"
                tail -n +2 -q ${codex_file} >> ${DATA_CODEX_DIR}${CODEX_OUTPUT_FILE}
        fi
done

###########################################
# STEP 2: Merge the output files together #
###########################################
cat ${DATA_CODEX_DIR}codex_calls.txt | sed -e 's/\.bam\b//' > ${DATA_DIR}codex_calls.txt

echo "Job ended on `hostname` at `date`"
```
