## Supplemental_Code for "A machine-learning approach for accurate detection of copy-number variants from exome sequencing": extract_bp_coverage.sh

```
#!/bin/bash
########################################################################
# Script : extract_bp_coverage.sh                                      #
# Author : Vijay Kumar                                                 #
# Date   : 4/5/2019                                                    #
#                                                                      #
# This script calculates basepair level coverage information for       #
# each sample. This information is used later to resolve breakpoints   #
# of concordant CNVs.                                                  #
#                                                                      #
# (c) 2019 - Vijay Kumar	                                       #
# Licenced under the GNU General Public License 3.0.                   #
########################################################################
echo "Job started on `hostname` at `date`"

source TBD/config.params

##################################################################
# STEP 1: Calculate actual and normalized coverage for each sample 
##################################################################
for file_loc in `cat ${BAM_FILE_LIST_W_PATH}`;
do

sample_name=`echo ${file_loc} | rev | cut -f1 -d/| rev`

${DOCKER_COMMAND}${BEDTOOLS_DIR}genomeCoverageBed -ibam ${file_loc} -bga \
                             > ${DATA_BPCOV_DIR}${sample_name}.bpcov.bed
done

echo "Job ended on `hostname` at `date`"
```
