## Supplemental_Code for "A machine-learning approach for accurate detection of copy-number variants from exome sequencing": extract_gc_map_vals.sh

```
#!/bin/bash
################################################################################
# Script : extract_gc_map_vals.sh                                              # 
# Author : Vijay Kumar                                                         #
# Date   : 4/5/2019                                                            #
#                                                                              #
# This script extracts GC content and mappability scores for the set of CNV    #
# predictions after the overlapping calls are merged together.                 #
# Prereqs (Format= <DIRECTORY_NAME> File Description):                         #
# 1) <DATA> File with the list of breakpoint-resolved CNVs w/ read depth ratio #
# 2) <SOURCE> Mappability scores file from the ENCODE project. This script     #
#             is designed to download the source file if it is not found.      #
# 3) <SOURCE> File with the list of exome capture probe coordinates            #
#                                                                              #
# (c) 2019 - Vijay Kumar                                                       #
# Licenced under the GNU General Public License 3.0.                           #
################################################################################
echo "Job started on `hostname` at `date`"

source TBD/config.params

#####################################
# STEP 0: Declare variables         #
#####################################
ALL_PRED_FILE=${DATA_DIR}'final_preds.txt'
ALL_PRED_W_GC=${DATA_DIR}'final_preds_GC.txt'
ALL_PRED_W_GC_MAP=${DATA_DIR}'final_preds_GC_Map.txt'
ALL_PRED_W_GC_MAP_TARG=${DATA_DIR}'final_preds_GC_Map_Targ.txt'

########################################################
# STEP 1: Extract mappability scores for each interval #
########################################################
cd ${SOURCE_DIR}
if [ ! -f ${SOURCE_DIR}wgEncodeCrgMapabilityAlign100mer.bigWig ];
then
wget http://hgdownload.cse.ucsc.edu/goldenpath/hg19/encodeDCC/wgEncodeMapability/wgEncodeCrgMapabilityAlign100mer.bigWig
fi

input_file_col_count=`head -1 ${ALL_PRED_FILE} | wc -w`

last_caller_col=$((5 + ${CALLER_COUNT}))
overlap_count_col=$((${last_caller_col} + 1))


#############################################################
# STEP 2: (Read Depth) Extract GC content for each interval #
#############################################################
if [ ${input_file_col_count} -eq $((${overlap_count_col} + 2)) ];
then
rd_ratio_col=$((${last_caller_col} + 3))
gc_content_col=$((${rd_ratio_col} + 2))
cnv_size_col=$((${gc_content_col} + 7))

${DOCKER_COMMAND}${BEDTOOLS_DIR}bedtools nuc -fi ${REF_GENOME} -bed ${ALL_PRED_FILE} | tail -n +2 | \
         cut -f1-${overlap_count_col},${rd_ratio_col},${gc_content_col},${cnv_size_col} > ${ALL_PRED_W_GC}

################################################################################
# Format the bed files to a format required to extract mappability scores      #
################################################################################
cat -n ${ALL_PRED_W_GC} | awk '{printf "%s\t%s\t%s\t%s\n", "chr"$2, $3, $4, $1}' \
                          > ${DATA_DIR}cons_pred_intvls_four_cols.bed

######################################################################################################
# Extract only the mappability score column along with the unique identifier column; sort the output #
######################################################################################################
${DOCKER_COMMAND}${SW_DIR}bigWigAverageOverBed ${SOURCE_DIR}wgEncodeCrgMapabilityAlign100mer.bigWig \
                                  ${DATA_DIR}cons_pred_intvls_four_cols.bed ${DATA_DIR}map_output.tab

cut -f1,6 ${DATA_DIR}map_output.tab | sort -k1 -n > ${DATA_DIR}sorted_map_scores

################################################################
# Add the mappability score column to the file with GC content #
################################################################
paste -d'\t' ${ALL_PRED_W_GC} ${DATA_DIR}sorted_map_scores | \
      cut -f1-${rd_ratio_col},$((${rd_ratio_col} + 1)),$((${rd_ratio_col} + 3)) > ${ALL_PRED_W_GC_MAP}

###################################################################
# This step is needed to count the number of targets that overlap #
# with each predicted CNV region. This is needed to ignore CNVs   #
# that do not overlap with atleast one target probe.              #
###################################################################
${DOCKER_COMMAND}${BEDTOOLS_DIR}intersectBed -c -a ${ALL_PRED_W_GC_MAP} -b ${TARGET_PROBES} \
                                                   > ${ALL_PRED_W_GC_MAP_TARG}


elif [ ${input_file_col_count} -eq ${overlap_count_col} ];
then
gc_content_col=$((${overlap_count_col} + 2))
cnv_size_col=$((${gc_content_col} + 7))

###########################################################
# STEP 2 (End join): Extract GC content for each interval #
###########################################################
${DOCKER_COMMAND}${BEDTOOLS_DIR}bedtools nuc -fi ${REF_GENOME} -bed ${ALL_PRED_FILE} | tail -n +2 | \
         cut -f1-${overlap_count_col},${gc_content_col},${cnv_size_col} > ${ALL_PRED_W_GC}

################################################################################
# Format the bed files to a format required to extract mappability scores      #
################################################################################
cat -n ${ALL_PRED_W_GC} | awk '{printf "%s\t%s\t%s\t%s\n", "chr"$2, $3, $4, $1}' \
                          > ${DATA_DIR}cons_pred_intvls_four_cols.bed

######################################################################################################
# Extract only the mappability score column along with the unique identifier column; sort the output #
######################################################################################################
${DOCKER_COMMAND}${SW_DIR}bigWigAverageOverBed ${SOURCE_DIR}wgEncodeCrgMapabilityAlign100mer.bigWig \
                                  ${DATA_DIR}cons_pred_intvls_four_cols.bed ${DATA_DIR}map_output.tab

cut -f1,6 ${DATA_DIR}map_output.tab | sort -k1 -n > ${DATA_DIR}sorted_map_scores

################################################################
# Add the mappability score column to the file with GC content #
################################################################
paste -d'\t' ${ALL_PRED_W_GC} ${DATA_DIR}sorted_map_scores | \
      cut -f1-$((${overlap_count_col} +1)),$((${overlap_count_col} + 2)),$((${overlap_count_col} + 4)) > ${ALL_PRED_W_GC_MAP}

###################################################################
# This step is needed to count the number of targets that overlap #
# with each predicted CNV region. This is needed to ignore CNVs   #
# that do not overlap with atleast one target probe.              #
###################################################################
${DOCKER_COMMAND}${BEDTOOLS_DIR}intersectBed -c -a ${ALL_PRED_W_GC_MAP} -b ${TARGET_PROBES} \
                                                   > ${ALL_PRED_W_GC_MAP_TARG}

fi

echo "Job ended on `hostname` at `date`" 
```
