## Supplemental_Code for "A machine-learning approach for accurate detection of copy-number variants from exome sequencing": merge_overlapping_CNVs_endjoin.sh

```
#!/bin/bash
################################################################################
# Script : merge_overlapping_CNVs_endjoin.sh                                   # 
# Author : Vijay Kumar                                                         #
# Date   : 4/5/2019                                                            #
#                                                                              #
# This is the master program that merges the calls with consensus among        #
# multiple callers and extracts overlap of each call with the consensus calls. #
# Prereqs (Format= <FILE LOCATION> File Description):                          #
# 1) <DATA> File with the list of CNVs along with overlap among callers        #
# 2) <SOURCE> File with the list of coordinates of the exome capture probes    #
# 3) <DATA> Files with the basepair level coverage info to measure read depth  #
#           ratio for each potential coordinate of CNVs with concordance       #
#                                                                              #
# (c) 2019 - Vijay Kumar                                                       #
# Licenced under the GNU General Public License 3.0.                           #
################################################################################
echo "Job started on `hostname` at `date`"

source /data/test_installation/CN_Learn/config.params

####################################################
# STEP 0: Declare directories, files and variables #
####################################################
CONS_PRED_W_OV_PROP_FILE_NAME='cons_calls_w_caller_ov_prop.bed'
TARGET_PROBES_W_LEN_ID='targets_auto_no_chr_w_uniq_id.bed'
ALL_PREDS_W_ALL_INTVL='preds_all_intvl_combos.txt'
ALL_PREDS_W_ALL_INTVL_TARG='all_samples_all_intvls_w_targ.txt'

##################################################################
# STEP 1: Make sure the input files are available for processing #
##################################################################
if [ ! -f ${TARGET_PROBES} ];
then
    echo "ERROR : The input file with the list of exome capture probes is unavailable.";
    echo "Please place this file in the SOURCE directory and try again."
    exit 1;
fi

################################################################
# Add a column with unique number for each exome capture probe #
################################################################
cat ${TARGET_PROBES} | awk 'BEGIN{OFS="\t"} {print $1, $2, $3, $3-$2, NR}' \
                     > ${PRED_DIR}${TARGET_PROBES_W_LEN_ID}

################################################################################
# STEP 2: Merge the overlapping calls made by multiple callers, in each sample #
################################################################################
for sample in `cat ${SAMPLE_LIST}`;
do

########################################################
# Loop through the CNV Type : Deletions & Duplications #
########################################################
for cnv_type in "DEL" "DUP";
do
cat ${DATA_DIR}${CONS_PRED_W_OV_PROP_FILE_NAME} | grep -w ${sample} \
    | grep ${cnv_type} > ${PRED_DIR}${sample}_${cnv_type}_preds.txt

################################################################################################
# Extract the predictions for each sample and CNV type and group them based on their intervals #
################################################################################################
${DOCKER_COMMAND}Rscript ${RSCRIPTS_DIR}generate_interval_combo.r  ${PRED_DIR} \
                         ${sample}_${cnv_type}_preds.txt \
                         ${sample}_${cnv_type}_preds_w_grps.txt  ${sample}_${cnv_type}_grouped_preds.txt \
                         ${sample}_${cnv_type}_preds_all_intvl_combos.txt  ${CALLER_COUNT}  ${CALLER_LIST}

done
done

###############################################################################
# STEP 3: Consolidate the output files from each sample into single output file
###############################################################################
# Remove the final files and create new ones prior to the for loop that loops #
# through sample and CNV type                                                 #
###############################################################################
if [ -f ${DATA_DIR}preds_w_grps_info.txt ];
then
rm ${DATA_DIR}preds_w_grps_info.txt
fi
touch ${DATA_DIR}preds_w_grps_info.txt

if [ -f ${DATA_DIR}extn_grouped_preds.txt ];
then
rm ${DATA_DIR}extn_grouped_preds.txt
fi
touch ${DATA_DIR}extn_grouped_preds.txt

#############################################################
# STEP 4: Loop through each sample and CNV type to generate #
#         two separate/consolidated output files.           #
#############################################################
for sample in `cat ${SAMPLE_LIST}`;
do
for cnv_type in "DEL" "DUP";
do

cat ${PRED_DIR}${sample}_${cnv_type}_preds_w_grps.txt >> ${DATA_DIR}preds_w_grps_info.txt
cat ${PRED_DIR}${sample}_${cnv_type}_grouped_preds.txt >> ${DATA_DIR}extn_grouped_preds.txt

done
done

last_caller_column=$((6 + ${CALLER_COUNT}))
concordance_column=$((${last_caller_column} + 1))
col_after_conc_column=$((${concordance_column} + 1))


awk -v OFS='\t' -v conc_col=${concordance_column} '{if ($conc_col != 1) print $1,$2,$3,$4,$5,"CONSENSUS";}' \
                                 ${DATA_DIR}extn_grouped_preds.txt  > ${DATA_DIR}extn_grouped_conc_preds.txt 
awk -v conc_col=${concordance_column} '{if ($conc_col == 1) print $0;}' ${DATA_DIR}extn_grouped_preds.txt  \
                                 > ${DATA_DIR}extn_grouped_nonconc_preds.txt

if [ -f ${DATA_DIR}CONSENSUS_caller_ov.txt ];
then
rm ${DATA_DIR}CONSENSUS_caller_ov.txt
fi
touch ${DATA_DIR}CONSENSUS_caller_ov.txt

#########################################################################
# STEP 5: Loop through each sample to measure the overlap for each call #
#         with the consensus CNV obtained for regions with overlaps.    #
#########################################################################
for sample in `cat ${SAMPLE_LIST}`;
do
for cnv_type in "DUP" "DEL";
do
cat ${DATA_DIR}extn_grouped_conc_preds.txt | grep -w ${sample} | grep ${cnv_type} \
                                  > ${PRED_DIR}CONSENSUS_${sample}_${cnv_type}.txt;

for caller in ${CALLER_LIST};
do
cat ${DATA_DIR}${CONS_PRED_W_OV_PROP_FILE_NAME} | grep -w ${caller} | grep -w ${sample} \
    | grep ${cnv_type} > ${PRED_DIR}${caller}_${sample}_${cnv_type}.txt

if [ -s ${PRED_DIR}CONSENSUS_${sample}_${cnv_type}.txt ] && [ -s ${PRED_DIR}${caller}_${sample}_${cnv_type}.txt ];
then

${DOCKER_COMMAND}${BEDTOOLS_DIR}intersectBed -wao -a ${PRED_DIR}CONSENSUS_${sample}_${cnv_type}.txt \
                                                  -b ${PRED_DIR}${caller}_${sample}_${cnv_type}.txt \
                                     | cut -f1-6,${col_after_conc_column},$((${col_after_conc_column} + 1)) \
                                     | awk -v OFS='\t' '{if ($8 > 0) print $0;}' \
                                     >> ${DATA_DIR}CONSENSUS_caller_ov.txt;

elif [ -s ${PRED_DIR}CONSENSUS_${sample}_${cnv_type}.txt ] && [ ! -s ${PRED_DIR}${caller}_${sample}_${cnv_type}.txt ];
then

${DOCKER_COMMAND}${BEDTOOLS_DIR}intersectBed -wao -a ${PRED_DIR}CONSENSUS_${sample}_${cnv_type}.txt \
                                     -b ${SOURCE_DIR}dummy.bed \
                                     | cut -f1-6,${col_after_conc_column},$((${col_after_conc_column} + 1)) \
                                     | awk -v OFS='\t' '{if ($8 > 0) print $0;}' \
                                     >> ${DATA_DIR}CONSENSUS_caller_ov.txt;
fi

done
done
done

${DOCKER_COMMAND}Rscript --vanilla ${RSCRIPTS_DIR}reshape_caller_overlap_data.r  ${DATA_DIR} \
                      ${DATA_DIR}CONSENSUS_caller_ov.txt  \
                      ${DATA_DIR}CONSENSUS_caller_ov_prop.txt \
                      ${CALLER_COUNT} ${CALLER_LIST}

cut -f6,7   --complement ${DATA_DIR}CONSENSUS_caller_ov_prop.txt > ${DATA_DIR}final_preds.txt
cut -f6,${col_after_conc_column}  --complement ${DATA_DIR}extn_grouped_nonconc_preds.txt >> ${DATA_DIR}final_preds.txt


echo "Job ended on `hostname` at `date`"
```
