## Supplemental_Code for "A machine-learning approach for accurate detection of copy-number variants from exome sequencing": merge_overlapping_CNVs_readdepth.sh

```
#!/bin/bash
################################################################################
# Script : merge_overlapping_CNVs_readdepth.sh                                 # 
# Author : Vijay Kumar                                                         #
# Date   : 4/5/2019                                                            #
#                                                                              #
# This is the master program that merges the calls with consensus among        #
# multiple callers and extracts additional read depth info.                    #
# Prereqs (Format= <FILE LOCATION> File Description):                          #
# 1) <DATA> File with the list of CNVs along with overlap among callers        #
# 2) <SOURCE> File with the list of coordinates of the exome capture probes    #
# 3) <DATA> Files with the basepair level coverage info to measure read depth  #
#           ratio for each potential coordinate of CNVs with concordance       #
#                                                                              #
# (c) 2019 - Vijay Kumar                                                       #
# Licenced under the GNU General Public License 3.0.                           #
################################################################################
echo "Job started on `hostname` at `date`"

source /data/test_installation/CN_Learn/config.params

####################################################
# STEP 0: Declare directories, files and variables #
####################################################
CONS_PRED_W_OV_PROP_FILE_NAME='cons_calls_w_caller_ov_prop.bed'
TARGET_PROBES_W_LEN_ID='targets_auto_no_chr_w_uniq_id.bed'
ALL_PREDS_W_ALL_INTVL='preds_all_intvl_combos.txt'
ALL_PREDS_W_ALL_INTVL_TARG='all_samples_all_intvls_w_targ.txt'

##################################################################
# STEP 1: Make sure the input files are available for processing #
##################################################################
if [ ! -f ${TARGET_PROBES} ];
then
    echo "ERROR : The input file with the list of exome capture probes is unavailable.";
    echo "Please place this file in the SOURCE directory and try again."
    exit 1;
fi

for sample in `cat ${SAMPLE_LIST}`;
do
    if [ ! -f ${DATA_BPCOV_DIR}${sample}.bpcov.bed ];
    then
        echo "ERROR : The file with basepair level coverage is missing for one ";
        echo "or more samples. Make sure the coverage file for each sample is  ";
        echo "available and rerun the script.";
        exit 1;
    fi
done

################################################################
# Add a column with unique number for each exome capture probe #
################################################################
cat ${TARGET_PROBES} | awk 'BEGIN{OFS="\t"} {print $1, $2, $3, $3-$2, NR}' \
                     > ${PRED_DIR}${TARGET_PROBES_W_LEN_ID}

################################################################################
# STEP 2: Merge the overlapping calls made by multiple callers, in each sample #
################################################################################
for sample in `cat ${SAMPLE_LIST}`;
do

########################################################
# Loop through the CNV Type : Deletions & Duplications #
########################################################
for cnv_type in "DEL" "DUP";
do
cat ${DATA_DIR}${CONS_PRED_W_OV_PROP_FILE_NAME} | grep -w ${sample} \
    | grep ${cnv_type} > ${PRED_DIR}${sample}_${cnv_type}_preds.txt

#######################################################################
# Extract the predictions for each sample and CNV type and group them #
# based on their intervals.                                           #   
#######################################################################
${DOCKER_COMMAND}Rscript ${RSCRIPTS_DIR}generate_interval_combo.r ${PRED_DIR}  \
                         ${sample}_${cnv_type}_preds.txt \
                         ${sample}_${cnv_type}_preds_w_grps.txt  \
                         ${sample}_${cnv_type}_grouped_preds.txt \
                         ${sample}_${cnv_type}_preds_all_intvl_combos.txt  \
                         ${CALLER_COUNT}  ${CALLER_LIST}

${DOCKER_COMMAND}${BEDTOOLS_DIR}intersectBed -wao \
                     -a ${PRED_DIR}${sample}_${cnv_type}_preds_all_intvl_combos.txt \
                     -b ${PRED_DIR}${TARGET_PROBES_W_LEN_ID} \
                     > ${PRED_DIR}${sample}_${cnv_type}_preds_all_intvl_targs.txt

${DOCKER_COMMAND}Rscript ${RSCRIPTS_DIR}identify_targets_of_interest.r  ${PRED_DIR} \
                         ${sample}_${cnv_type}_preds_all_intvl_targs.txt  ${TARGET_PROBES_W_LEN_ID} \
                         ${sample}_${cnv_type}_all_intvl_info.txt  \
                         ${sample}_${cnv_type}_targets_of_interest.txt \
                         ${CALLER_COUNT} ${CALLER_LIST}

################################################################################################
# Generate separate files with probe information for predicted and left/right flanking regions #
################################################################################################
last_caller_column=$((5 + ${CALLER_COUNT}))
addl_info_col_start=$((${last_caller_column} + 1))
addl_info_col_end=$((${last_caller_column} + 4))
left_flank_start_col=$((${last_caller_column} + 5))
left_flank_end_col=$((${last_caller_column} + 6))
pred_region_start_col=$((${last_caller_column} + 7))
pred_region_end_col=$((${last_caller_column} + 8))
right_flank_start_col=$((${last_caller_column} + 9))
right_flank_end_col=$((${last_caller_column} + 10))
comb_column=$((${last_caller_column} + 11))

cat ${PRED_DIR}${sample}_${cnv_type}_all_intvl_info.txt | \
    cut -f1-${addl_info_col_end},${comb_column},${left_flank_start_col},${left_flank_end_col} \
    > ${PRED_DIR}${sample}_${cnv_type}_all_intvl_info_left_flank.txt
cat ${PRED_DIR}${sample}_${cnv_type}_all_intvl_info.txt | \
    cut -f1-${addl_info_col_end},${comb_column},${pred_region_start_col},${pred_region_end_col} \
    > ${PRED_DIR}${sample}_${cnv_type}_all_intvl_info_pred_region.txt
cat ${PRED_DIR}${sample}_${cnv_type}_all_intvl_info.txt | \
    cut -f1-${addl_info_col_end},${comb_column},${right_flank_start_col},${right_flank_end_col} \
    > ${PRED_DIR}${sample}_${cnv_type}_all_intvl_info_right_flank.txt

################################################
# Extract coverage for each target of interest #
################################################
${DOCKER_COMMAND}${BEDTOOLS_DIR}intersectBed -wao \
                    -a ${PRED_DIR}${sample}_${cnv_type}_targets_of_interest.txt \
                    -b ${DATA_BPCOV_DIR}${sample}.bpcov.bed \
                    > ${PRED_DIR}${sample}_${cnv_type}_targets_of_interest_w_cov.txt

${DOCKER_COMMAND}Rscript ${RSCRIPTS_DIR}measure_rd_stats.r  ${PRED_DIR} \
                         ${sample}_${cnv_type}_all_intvl_info_left_flank.txt  \
                         ${sample}_${cnv_type}_all_intvl_info_pred_region.txt \
                         ${sample}_${cnv_type}_all_intvl_info_right_flank.txt \
                         ${sample}_${cnv_type}_targets_of_interest_w_cov.txt  \
                         ${sample}_${cnv_type}_w_rd_stats.txt \
                         ${CALLER_COUNT} ${CALLER_LIST}

done
done

###############################################################################
# STEP 3: Consolidate the output files from each sample into single output file
###############################################################################
# Remove the final files and create new ones prior to the for loop that loops #
# through sample and CNV type                                                 #
###############################################################################
if [ -f ${DATA_DIR}preds_w_grps_info.txt ];
then
rm ${DATA_DIR}preds_w_grps_info.txt
fi
touch ${DATA_DIR}preds_w_grps_info.txt

if [ -f ${DATA_DIR}extn_grouped_preds.txt ];
then
rm ${DATA_DIR}extn_grouped_preds.txt
fi
touch ${DATA_DIR}extn_grouped_preds.txt

if [ -f ${DATA_DIR}all_intvl_info.txt ];
then
rm ${DATA_DIR}all_intvl_info.txt
fi
touch ${DATA_DIR}all_intvl_info.txt

if [ -f ${DATA_DIR}final_preds.txt ];
then
rm ${DATA_DIR}final_preds.txt
fi
touch ${DATA_DIR}final_preds.txt

#############################################################
# STEP 4: Loop through each sample and CNV type to generate #
#         four separate/consolidated output files.          #
#############################################################
for sample in `cat ${SAMPLE_LIST}`;
do
for cnv_type in "DEL" "DUP";
do

cat ${PRED_DIR}${sample}_${cnv_type}_preds_w_grps.txt >> ${DATA_DIR}preds_w_grps_info.txt
cat ${PRED_DIR}${sample}_${cnv_type}_grouped_preds.txt >> ${DATA_DIR}extn_grouped_preds.txt
cat ${PRED_DIR}${sample}_${cnv_type}_all_intvl_info.txt >> ${DATA_DIR}all_intvl_info.txt
cat ${PRED_DIR}${sample}_${cnv_type}_w_rd_stats.txt >> ${DATA_DIR}final_preds.txt

done
done

echo "Job ended on `hostname` at `date`"
```
