## Supplemental_Code for "A machine-learning approach for accurate detection of copy-number variants from exome sequencing": plots_for_paper.r

```
#######################################################################################
# Author: Vijay Kumar                                                                 #
# Date  : 4/5/2019                                                                    #
# This is the main script that generates every summary plot required for the project. #
# The first section of the script imports a set of input files while the following    #
# sections use ggplot2 to generate the plots and save them as tiff files.             #
#                                                                                     #
# For readability and ease of use, the input files are listed below                   #
#    1) snp_overlap_data_w_callerinfo.txt - Original set of calls with MA SNP count   #
#    2) cons_overlap_w_caller_0.1.txt - Calls that overlap with atleast 1 SNP M.array #
#    3) cons_overlap_strat1_0.1.txt - Calls consolidated by the first strategy        #
#    4) cons_overlap_strat2_0.1.txt - Calls consolidated by the second strategy       #
#    5) cons_overlap_strat3_0.1.txt - Calls with a particular caller as validation    #
#    6) val_summ_by_sample.txt - List of samples with number of CNVs called by MA     #
#    7) RF_Metrics_SVIP.csv - Classifier metrics for all classifiers                  #
#    8) True_Pos_Calls_SVIP.csv - List of TP calls from first strategy                #
#    9) False_Pos_Calls_SVIP.csv - List of FP calls from third strategy               #
#   10) Samplelist_MA_SVIP.csv - List of samples in the test set (MA)                 #
#   11) Samplelist_CLAMMS_SVIP.csv - List of samples in the test set (CLAMMS)         #
#   12) Overlap_Stats_strat2 - Overlap stats with validated data at different props   #
#   13) Overlap_Stats_strat3 - Overlap stats with validated data at different props   #
#   14) 16p_dups_dels.txt - List of samples with 16p duplication or deletion          #
#   15) 16p_overlap.txt - Subset of calls on Chromosome 16                            #
#   16) 16p_overlap_only.txt - Subset of calls that overlapped with 16p events        #
#   17) 16p_overlap_cnlearn.txt - Subset of 16p overlap calls w/ first strategy       #
#######################################################################################
library(sqldf)
library(ggplot2)
library(reshape2)
library(gplots)
library(limma)
library(Rtsne)
library(ggfortify)
library(rgl)
library(corrplot)
library(Hmisc)

####################################################################
# Read the files that include CALLER information to generate metrics
####################################################################
original_calls_df <- read.table("snp_overlap_data_w_callerinfo.txt")
colnames(original_calls_df) <- c("CHR", "PRED_START", "PRED_END", "TYPE", "SAMPLE", "CALLER", "PRED_QUAL", "PRED_SIZE", "UNIQ_PRED_ID", 
                                 "CANOES_PROP", "CODEX_PROP", "CLAMMS_PROP", "XHMM_PROP", 
                                 "CANOES_OV", "CODEX_OV", "CLAMMS_OV", "XHMM_OV", "NUM_OVERLAPS", "NUM_TARGETS", "NUM_SNPS")
dim(original_calls_df)
retained_calls_df <- read.table("cons_overlap_w_caller_0.1.txt",  header = TRUE)
retained_calls_df$NUM_OVERLAPS <- retained_calls_df$NUM_OVERLAPS + 1

retained_calls_gt5kb_df <- sqldf("select * from retained_calls_df where PRED_SIZE > 5000")
retained_calls_gt50kb_df <- sqldf("select * from retained_calls_df where PRED_SIZE > 50000")

ma_val_count_df <- read.table("val_summ_by_sample.txt")
colnames(ma_val_count_df) <- c("SAMPLE", "COUNT")

######################################################################
# Read the files prior to and after merging the calls with concordance
######################################################################
all_calls_before_bpres_df <- read.table("cons_calls_w_caller_ov_prop.bed")
colnames(all_calls_before_bpres_df) <- c("CHR", "PRED_START", "PRED_END", "TYPE", "SAMPLE", "CALLER", "PRED_QUAL", "PRED_SIZE", "UNIQ_PRED_ID", 
                                         "CANOES_PROP", "CODEX_PROP", "CLAMMS_PROP", "XHMM_PROP", 
                                         "CANOES_OV", "CODEX_OV", "CLAMMS_OV", "XHMM_OV", "NUM_OVERLAPS")

all_calls_after_bpres_df <- read.table("final_preds_w_rd_stats.txt")
colnames(all_calls_after_bpres_df) <- c("CHR", "PRED_START", "PRED_END", "TYPE", "SAMPLE", "CANOES_PROP", "CODEX_PROP", "CLAMMS_PROP", "XHMM_PROP", 
                                        "CANOES_OV", "CODEX_OV", "CLAMMS_OV", "XHMM_OV", "NUM_OVERLAPS", "GROUP", "RD_PROP")

strat3_calls_after_bpres_df <- read.table("wo_clamms_final_preds_w_rd_stats.txt")
colnames(strat3_calls_after_bpres_df) <- c("CHR", "PRED_START", "PRED_END", "TYPE", "SAMPLE", "CANOES_PROP", "CODEX_PROP", "CLAMMS_PROP", "XHMM_PROP", 
                                           "CANOES_OV", "CODEX_OV", "CLAMMS_OV", "XHMM_OV", "NUM_OVERLAPS", "GROUP", "RD_PROP")
head(strat3_calls_after_bpres_df)
head(all_calls_before_bpres_df)
head(all_calls_after_bpres_df)

val_overlap_stats_strat2_df <- read.table("Overlap_Stats_strat2")
val_overlap_stats_strat3_df <- read.table("Overlap_Stats_strat3")

###############################################################
# Read the file with the list of exome target probe coordinates
###############################################################
target_list <- read.table("targets_auto_no_chr.bed")

#############################################################
# Read the files to generate metrics from consolidated calls
#############################################################
strat1_df <- read.table("cons_overlap_strat1_0.1.txt",  header = TRUE)
strat2_df <- read.table("cons_overlap_strat2_0.1.txt",  header = TRUE)
strat3_df <- read.table("cons_overlap_strat3_0.1.txt",  header = TRUE)

strat2_df$NUM_OVERLAPS <- strat2_df$NUM_OVERLAPS + 1

#############################################################
# Read the files to generate classifier performance measures
#############################################################
rf_metrics_df <- read.csv("RF_Metrics_SVIP.csv")
rf_metrics_size_df <- read.csv("RF_Metrics_Size.csv")


#############################################################
# Read the files to generate true positive distribution
#############################################################
tp_df <- read.csv("True_Pos_Calls_SVIP.csv")
fp_df <- read.csv("False_Pos_Calls_SVIP.csv")
fn_df <- read.csv("False_Neg_Calls_SVIP.csv")
tn_df <- read.csv("True_Neg_Calls_SVIP.csv")

tp_df$CLASS <- 'True Positive'
fp_df$CLASS <- 'False Positive'
fn_df$CLASS <- 'False Negative'
tn_df$CLASS <- 'True Negative'

tp_df$VAL_LABEL <- 'True CNV'
fn_df$VAL_LABEL <- 'True CNV'
tn_df$VAL_LABEL <- 'False CNV'
fp_df$VAL_LABEL <- 'False CNV'


tp_df$NUM_OVERLAPS <- tp_df$NUM_OVERLAPS + 1
fp_df$NUM_OVERLAPS <- fp_df$NUM_OVERLAPS + 1
tn_df$NUM_OVERLAPS <- tn_df$NUM_OVERLAPS + 1
fn_df$NUM_OVERLAPS <- fn_df$NUM_OVERLAPS + 1

tp_fp_df <- rbind(tp_df, fp_df)
tp_fp_fn_df <- rbind(tp_df, fp_df, fn_df)
tp_fp_tn_fn_df <- rbind(tp_df, fp_df, tn_df, fn_df)
tn_fn_df <- rbind(tn_df, fn_df)

tp_fp_tn_fn_df$VAL_LABEL <- ordered(tp_fp_tn_fn_df$VAL_LABEL, levels = c("True CNV", "False CNV"))
tp_fp_tn_fn_df$CLASS <- ordered(tp_fp_tn_fn_df$CLASS, levels = c("True Positive", "False Positive", "True Negative", "False Negative"))

head(tp_df)
####################################################################
# This portion generates the report on the number of 16P deletions #
# and duplications identified by each caller                       #
####################################################################
ma_sample_list_df <- as.data.frame(read.csv("Samplelist_MA_SVIP.csv",header = TRUE))
clamms_sample_list_df <- as.data.frame(read.csv("Samplelist_CLAMMS_SVIP.csv", header = TRUE))

chr16p_samples_df <- as.data.frame(read.table("16p_dups_dels.txt"))
chr16p_ind_callers_df <- as.data.frame(read.table("16p_overlap_only.txt"))
chr16p_cnlearn_df <- as.data.frame(read.table("16p_overlap_cnlearn.txt"))

colnames(chr16p_samples_df) <- c("CHR", "START", "END", "SAMPLE", "CNV_TYPE")
colnames(chr16p_ind_callers_df) <- c("CHR", "PRED_START", "PRED_END", "CNV_TYPE", "SAMPLE", "CALLER", "PRED_QUAL", "PRED_SIZE", "UNIQ_PRED_ID",  "CANOES_PROP", "CODEX_PROP", 
                                     "CLAMMS_PROP", "XHMM_PROP", "CANOES_OV", "CODEX_OV", "CLAMMS_OV", "XHMM_OV", "NUM_OVERLAPS",
                                     "NUM_TARGETS", "NUM_SNPS", "OV_STATUS")

colnames(chr16p_cnlearn_df) <- c("CHR", "PRED_START", "PRED_END", "CNV_TYPE", "SAMPLE", "CANOES_PROP", "CODEX_PROP", "CLAMMS_PROP", "XHMM_PROP", "NUM_OVERLAPS", "RD_PROP",
                                 "GC_CONTENT", "PRED_SIZE", "MAPPABILITY","NUM_TARGETS","NUM_SNPS", "OVERLAP_PROP", "SIZE_LABEL", "PRED_PROBS", "CLASSIFIER_NAME", "NUM_SPLITS", "STRATEGY", "TRAINING_PROP", "OV_STATUS")

##########################################################
# Read the files needed to generate supplementary tables #
##########################################################
sample_all_intvls <- read.table("14734.x11_DEL_preds_all_intvl_combos.txt")
colnames(sample_all_intvls) <- c("CHR", "PRED_START", "PRED_END", "TYPE", "SAMPLE", "CALLER_ST", "CALLER_END", "UNIQ_PRED_ST", "UNIQ_PRED_END", 
                                 "CANOES_PROP", "CODEX_PROP", "CLAMMS_PROP", "XHMM_PROP", 
                                 "CANOES_OV", "CODEX_OV", "CLAMMS_OV", "XHMM_OV", "NUM_OVERLAPS", "GROUP_NAME")

########################################
# Global variables for subsequent uses #
########################################
venn_colors <- c("red", "blue", "green", "orchid")

##########################################################
# Figure 2B: Correlation between predictors (Microarray) #
##########################################################
corr_predictors_df <- strat2_df[,c("CANOES_PROP", "CODEX_PROP", "CLAMMS_PROP", "XHMM_PROP", "NUM_OVERLAPS", "PRED_SIZE", "NUM_TARGETS", "GC", "MAP", "RD_PROP")]
colnames(corr_predictors_df) <- c("CANOES", "CODEX", "CLAMMS", "XHMM", "Concordance", "Pred Size", "Target Count", "GC Content", "Mappability", "Read Depth Ratio")
corr_results <- cor(corr_predictors_df)

flattenCorrMatrix <- function(cormat, pmat) {
  ut <- upper.tri(cormat)
  data.frame(
    row = rownames(cormat)[row(cormat)[ut]],
    column = rownames(cormat)[col(cormat)[ut]],
    cor  =(cormat)[ut],
    p = pmat[ut]
  )
}

options(scipen=999)
result_spearman <-rcorr(as.matrix(corr_predictors_df), type = "spearman")
flattenCorrMatrix(round(result_spearman$r,3), round(result_spearman$P,3))

correlations <- round(as.data.frame(result_spearman$r),2)
write.csv(correlations, file="supp_table_1_pred_correlations_ma.csv")


#Build correlation plot
pdf("fig_2b_predictor_correlations_microarray.pdf", width = 8, height = 5)
corrplot(result_spearman$r, type="upper", p.mat = result_spearman$P, sig.level = 0.01, insig = "blank", tl.col = "black", tl.srt = 45)
dev.off()


#############################################################
# Figure 2C: Prediction probability analysis - (Microarray) #
#############################################################
tp_fp_tn_fn_ma_df <- tp_fp_tn_fn_df
tp_fp_tn_fn_ma_df$CANOES <- ifelse(tp_fp_tn_fn_ma_df$CANOES_PROP > 0, 1, 0)
tp_fp_tn_fn_ma_df$CLAMMS <- ifelse(tp_fp_tn_fn_ma_df$CLAMMS_PROP > 0, 1, 0)
tp_fp_tn_fn_ma_df$CODEX<- ifelse(tp_fp_tn_fn_ma_df$CODEX_PROP > 0, 1, 0)
tp_fp_tn_fn_ma_df$XHMM <- ifelse(tp_fp_tn_fn_ma_df$XHMM_PROP > 0, 1, 0)
tp_fp_tn_fn_ma_df$OV_CODE <- paste(tp_fp_tn_fn_ma_df$CANOES, tp_fp_tn_fn_ma_df$CLAMMS, tp_fp_tn_fn_ma_df$CODEX, tp_fp_tn_fn_ma_df$XHMM, sep = '') 


tp_fp_tn_fn_probs_size_ma_df <- sqldf("select TYPE, PRED_SIZE, NUM_OVERLAPS, PRED_PROBS, CLASS, VAL_LABEL
                                      from tp_fp_tn_fn_ma_df 
                                      where CLASSIFIER = 'RF' and STRATEGY = '2' and NUM_SPLITS = 1 and TRAINING_PROP = 0.7
                                      and PRED_PROBS > 0")

#Generate plot by Validation Label (Distribution Plot)
pdf("fig_2c_predicted_probability_distribution_microarray_0.7.pdf", width = 8, height = 5)
ggplot(data=tp_fp_tn_fn_probs_size_ma_df, aes(x=PRED_PROBS, fill = VAL_LABEL)) + theme_classic() + 
  geom_histogram(position = "dodge", aes(y=..count../sum(..count..)), binwidth = 0.05) +
  ggtitle ("Distribution of predicted probability scores (Excludes zero)") +
  scale_fill_manual(values = c("True CNV" = "blue","False CNV" = "red")) +
  theme(plot.title = element_text(lineheight=.8, size=9, face="bold", hjust = 0.5), 
        strip.text = element_text(size = 9, face = "bold"),
        axis.title = element_text(size = 9, face="bold"),
        axis.text = element_text(size = 9, face="bold"), 
        legend.title = element_text(size=9, face="bold"),
        legend.text = element_text(size=9, face="bold"),
        legend.position = "bottom") +
  geom_vline(xintercept = 0.5, linetype="dotted") + 
  labs(x = "Predicted Probabilities", y = "Frequency", fill = "") 
dev.off()


#############################################################
# Figure 3B: Prediction probability analysis - (Microarray) #
#############################################################
tp_summary_df <- sqldf("select STRATEGY,  SPLIT, TRAINING_PROP, CLASS, count(*) as COUNT
                       from tp_df
                       where NUM_SPLITS = 10 and STRATEGY = 2 and CLASSIFIER = 'RF'
                       group by STRATEGY,  SPLIT, TRAINING_PROP, CLASS")
fp_summary_df <- sqldf("select STRATEGY, SPLIT, TRAINING_PROP, CLASS, count(*) as COUNT
                       from fp_df
                       where NUM_SPLITS = 10 and STRATEGY = 2 and CLASSIFIER = 'RF'
                       group by STRATEGY, SPLIT, TRAINING_PROP, CLASS")
fn_summary_df <- sqldf("select STRATEGY, SPLIT, TRAINING_PROP, CLASS, count(*) as COUNT
                       from fn_df
                       where NUM_SPLITS = 10 and STRATEGY = 2 and CLASSIFIER = 'RF'
                       group by STRATEGY, SPLIT, TRAINING_PROP, CLASS")
tn_summary_df <- sqldf("select STRATEGY,  SPLIT, TRAINING_PROP, CLASS, count(*) as COUNT
                       from tn_df
                       where NUM_SPLITS = 10 and STRATEGY = 2 and CLASSIFIER = 'RF'
                       group by STRATEGY, SPLIT, TRAINING_PROP, CLASS")

all_prec_recall_df <- sqldf("select tp.TRAINING_PROP, tp.SPLIT, round(tp.COUNT/((tp.COUNT + fp.COUNT) * 1.0),2) as Precision,
                            round(tp.COUNT/((tp.COUNT + fn.COUNT) * 1.0),2) as Recall
                            from tp_summary_df tp, fp_summary_df fp, fn_summary_df fn
                            where tp.SPLIT = fp.SPLIT and tp.TRAINING_PROP = fp.TRAINING_PROP and
                            tp.SPLIT = fn.SPLIT and tp.TRAINING_PROP = fn.TRAINING_PROP
                            order by tp.TRAINING_PROP")

write.csv(all_prec_recall_df, file="supp_table_2_precision_dist_ma.csv", row.names = FALSE)

melted_all_prec_recall_df <- melt(all_prec_recall_df, id=c("TRAINING_PROP", "SPLIT"))
means <- aggregate(value ~  TRAINING_PROP + variable, melted_all_prec_recall_df, mean)
means$value <- round(means$value,2)

pdf(file="fig_3b_precision_distribution_microarray.pdf", width = 5, height = 5)
ggplot(data=melted_all_prec_recall_df, aes(x=as.character(TRAINING_PROP), y=value, fill = variable)) + theme_classic() +
  geom_boxplot(position=position_dodge(1)) + ylim(0.5,1.0) + 
  geom_point(position=position_jitterdodge(dodge.width=0.9)) +
  geom_text(data = means, aes(label = value, y = value + 0.08)) +
  #  geom_text(aes(label=label_val), position = position_dodge(width = 0.9), vjust = -.25, size = 2, fontface = "bold") +
  ggtitle ("Distribution of performance obtained during 10-fold cross-validation") + 
  theme(plot.title = element_text(lineheight=.8, size=9, face="bold", hjust = 0.5), 
        strip.text = element_text(size = 8, face = "bold"),
        axis.title = element_text(size = 8, face="bold"),
        axis.text = element_text(size = 8, face="bold"), 
        #        axis.text.x = element_blank(),
        legend.title = element_text(size=8, face="bold"),
        legend.text = element_text(size=8, face="bold"),
        legend.position = "bottom") +
  labs(x = "Training Proportion", y = "Precision-Recall", fill = "") 
dev.off()

###########################################################################
# Figure 3C: Feature Importances for the main merge strategy (Microarray) #
###########################################################################
feat_imp_strat2 <- sqldf ("select Training_Proportion, Chromosome, CNV_Type, Canoes_Proportion, Codex_Proportion, Clamms_Proportion, XHMM_Proportion, Overlap_Count, 
                          RD_Proportion, GC_Content, Mappability, Number_Of_Targets, Size_Label, 'Genomic' as Predictor_Group 
                          from rf_metrics_df 
                          where Classifier = 'RF' and Strategy = 2 and Num_Splits = 10 and Training_Proportion = 0.7")
melted_feat_imp_strat2 <- melt(feat_imp_strat2, id=c("Training_Proportion", "Predictor_Group"))
melted_feat_imp_strat2$prop <- melted_feat_imp_strat2$value *100
melted_feat_imp_strat2$Predictor_Group[3:7] <- "Caller"
melted_feat_imp_strat2$pred_var_label <- c("Chromosome", "CNV Type", "CANOES", "CODEX", "CLAMMS", "XHMM", "Concordance", "Read Depth Ratio", "GC Content", 
                                           "Mappability", "Target Probe Count", "Size Label")
melted_feat_imp_strat2 <- sqldf("select * from melted_feat_imp_strat2 order by prop desc")

melted_feat_imp_strat2$pred_var_label <- 
  factor(melted_feat_imp_strat2$pred_var_label, levels = melted_feat_imp_strat2$pred_var_label[order(as.numeric(melted_feat_imp_strat2$prop))])

pdf("fig_3c_feature_importance_randomforest_microarray_0.7.pdf", width = 8, height = 5)
ggplot(data=melted_feat_imp_strat2, aes(x=reorder(pred_var_label,-prop), y=prop)) + theme_classic() +
  geom_bar(position = "dodge", stat="identity") + 
  #  facet_grid(~Predictor_Group, scale = "free", switch = "x") +
  geom_text(aes(label=paste0(prop,"%")), position = position_dodge(width = 0.9), vjust = -.25, size = 3.5, fontface = "bold") +
  ggtitle ("CN-Learn - Feature Importance") +
  theme(plot.title = element_text(lineheight=.8, size=15, face="bold", hjust = 0.5), 
        strip.text = element_text(size = 12, face = "bold"),
        axis.title = element_text(size = 12, face="bold"),
        axis.text = element_text(size = 12, face="bold"), 
        #        axis.text.x = element_blank(),
        axis.text.x = element_text(angle = 60, hjust = 1),
        legend.title = element_text(size=12, face="bold"),
        legend.text = element_text(size=10, face="bold")) +
  labs(x = "Predictors", y = "Feature Importance", fill = "       Predictors") 
dev.off()

#########################################################################################################
# Figure 3D: Characterization of callers based on precision rates at different size ranges (Microarray) #
#########################################################################################################
retained_calls_df <- read.table("cons_overlap_w_caller_0.1.txt",  header = TRUE)
rf_metrics_size_df <- read.csv("RF_Metrics_Size.csv")

rf_metrics_size_df$Size <- ''
rf_metrics_size_df$Size[c(1,2,3,4,17,18,19,20)] <- '50-100kb'
rf_metrics_size_df$Size[c(5,6,7,8,21,22,23,24)] <- '100-250kb'
rf_metrics_size_df$Size[c(9,10,11,12,25,26,27,28)] <- '250-500kb'
rf_metrics_size_df$Size[c(13,14,15,16,29,30,31,32)] <- '500kb-5mb'
rf_metrics_size_df$Size[c(33,34,35,36)] <- '5-10kb'
rf_metrics_size_df$Size[c(37,38,39,40)] <- '10-25kb'
rf_metrics_size_df$Size[c(41,42,43,44)] <- '25-50kb'
rf_metrics_size_df$Size <- ordered(rf_metrics_size_df$Size, levels = c("5-10kb", "10-25kb", "25-50kb", "50-100kb", "100-250kb",
                                                                       "250-500kb", "500kb-5mb"))

rf_metrics_size_modified_df <- sqldf("select (case when Strategy = 2 then 'CN_Learn' 
                                     else 'CN_Learn (w/ CLAMMS Gold Standard)' end) as Caller, Precision, Recall, AUC, Size 
                                     from rf_metrics_size_df where Training_Proportion = 0.7")

rf_metrics_size_modified_df <- melt(rf_metrics_size_modified_df[1:4,],id = c("Caller", "Size"))

retained_calls_modified_df <- retained_calls_df
retained_calls_modified_df$Size <- ''
retained_calls_modified_df$Size <- ifelse(retained_calls_modified_df$SIZE_LABEL=='C)5KB-10KB', '5-10kb', retained_calls_modified_df$Size)
retained_calls_modified_df$Size <- ifelse (retained_calls_modified_df$SIZE_LABEL=='D)10KB-25KB', '10-25kb', retained_calls_modified_df$Size)
retained_calls_modified_df$Size <- ifelse (retained_calls_modified_df$SIZE_LABEL=='E)25KB-50KB', '25-50kb', retained_calls_modified_df$Size)
retained_calls_modified_df$Size <- ifelse (retained_calls_modified_df$SIZE_LABEL=='F)50KB-75KB', '50-100kb', retained_calls_modified_df$Size)
retained_calls_modified_df$Size <- ifelse (retained_calls_modified_df$SIZE_LABEL=='G)75KB-100KB', '50-100kb', retained_calls_modified_df$Size)
retained_calls_modified_df$Size <- ifelse (retained_calls_modified_df$SIZE_LABEL=='H)100KB-250KB', '100-250kb', retained_calls_modified_df$Size)
retained_calls_modified_df$Size <- ifelse (retained_calls_modified_df$SIZE_LABEL=='I)250KB-500KB', '250-500kb', retained_calls_modified_df$Size)
retained_calls_modified_df$Size <- ifelse (retained_calls_modified_df$SIZE_LABEL=='J)500KB-1MB', '500kb-5mb', retained_calls_modified_df$Size)
retained_calls_modified_df$Size <- ifelse (retained_calls_modified_df$SIZE_LABEL=='K)1MB-5MB', '500kb-5mb', retained_calls_modified_df$Size)

retained_calls_modified_df$Size <- ordered(retained_calls_modified_df$Size, levels = c("5-10kb", "10-25kb", "25-50kb", "50-100kb", "100-250kb",
                                                                                       "250-500kb", "500kb-5mb"))

head(retained_calls_modified_df)

caller_precisions_by_size <- sqldf("select A.CALLER as Caller, round(A.COUNT/(B.COUNT * 1.0), 2) as Precision, 
                                   A.Size from 
                                   (select CALLER, Size, LABEL, count(*) as COUNT from retained_calls_modified_df 
                                   group by CALLER, Size, LABEL) A,
                                   (select CALLER, Size, count(*) as COUNT from retained_calls_modified_df 
                                   group by CALLER, Size) B
                                   where A.CALLER = B.CALLER and A.Size = B.Size
                                   and label = 'TRUE_CNV'")

caller_precisions_by_size <- melt(caller_precisions_by_size, id = c("Caller", "Size"))

all_precisions_by_size <- rbind(caller_precisions_by_size[,], rf_metrics_size_modified_df[,])
all_precisions_by_size <- all_precisions_by_size[which(all_precisions_by_size$Size != '5-10kb' &
                                                         all_precisions_by_size$Size != '10-25kb' &
                                                         all_precisions_by_size$Size != '25-50kb'), ]
all_precisions_by_size$Caller <- paste0(all_precisions_by_size$Caller, " (", all_precisions_by_size$variable, ")")
all_precisions_by_size$Caller <- ordered(all_precisions_by_size$Caller, levels = c("CN_Learn (Precision)", "CN_Learn (Recall)", "CN_Learn (AUC)", 
                                                                                   "CANOES (Precision)", "CLAMMS (Precision)", 
                                                                                   "CODEX (Precision)", "XHMM (Precision)")) 

pdf("fig_3d.pdf", width = 7, height = 5)
ggplot(all_precisions_by_size, aes(x = Size, y = value, group = Caller, color = Caller)) +
  ylim(0.0, 1.00) + geom_point(size=2.5, shape=1, stroke = 1.5) + geom_line(size=1) + theme_classic() +
  geom_text(aes(label=value), vjust = -0.5, size = 2.5, hjust = 1.25, fontface = "bold") +
  theme(plot.title = element_text(lineheight=.8, size=9, face="bold", hjust = 0.5), 
        strip.text = element_text(size = 9, face = "bold"),
        axis.title = element_text(size = 9, face="bold"),
        axis.text = element_text(size = 9, face="bold"), 
        legend.title = element_text(size=9, face="bold"),
        legend.text = element_text(size=9, face="bold"), 
        legend.position = "bottom") +
  labs(x = "Size Range", y = "Precision", color = "") 
dev.off()


################################################################################################################################
# Figure 4A: Distribution of all calls in test samples vs. classified as 'True' by the classifier (Microarray) (Venn Diagrams) #
################################################################################################################################
# Profile of calls made in all the samples in the test set
all_test_calls_strat2_df <- sqldf("select a.* from  strat2_df a, ma_sample_list_df b
                                  where a.PRED_SIZE > 50000 and a.PRED_SIZE < 5000000
                                  and b.TRAINING_PROP = 0.1 and b.CLASSIFIER_NAME = 'RF' and b.STRATEGY = 2
                                  and a.SAMPLE = b.SAMPLE")
all_test_calls_venn_df <- data.frame(matrix(,ncol=0,nrow=dim(all_test_calls_strat2_df)[1]))
all_test_calls_venn_df$CANOES <- ifelse(all_test_calls_strat2_df$CANOES_PROP > 0, 1, 0)
all_test_calls_venn_df$CODEX <- ifelse(all_test_calls_strat2_df$CODEX_PROP > 0, 1, 0)
all_test_calls_venn_df$CLAMMS <- ifelse(all_test_calls_strat2_df$CLAMMS_PROP > 0, 1, 0)
all_test_calls_venn_df$XHMM <- ifelse(all_test_calls_strat2_df$XHMM_PROP > 0, 1, 0)
all_test_calls_venn_counts_df <- vennCounts(all_test_calls_venn_df)
all_test_calls_venn_counts_df[1,5] <- paste0("N=", sum(all_test_calls_venn_counts_df[,5]))

all_true_test_calls_strat2_df <- sqldf("select a.* from strat2_df a, ma_sample_list_df b
                                       where a.PRED_SIZE > 50000 and a.PRED_SIZE < 5000000 and a.LABEL = 'TRUE_CNV'
                                       and b.TRAINING_PROP = 0.1 and b.CLASSIFIER_NAME = 'RF' and b.STRATEGY = 2
                                       and a.SAMPLE = b.SAMPLE and LABEL = 'TRUE_CNV'")
all_true_test_calls_venn_df <- data.frame(matrix(,ncol=0,nrow=dim(all_true_test_calls_strat2_df)[1]))
all_true_test_calls_venn_df$CANOES <- ifelse(all_true_test_calls_strat2_df$CANOES_PROP > 0, 1, 0)
all_true_test_calls_venn_df$CODEX <- ifelse(all_true_test_calls_strat2_df$CODEX_PROP > 0, 1, 0)
all_true_test_calls_venn_df$CLAMMS <- ifelse(all_true_test_calls_strat2_df$CLAMMS_PROP > 0, 1, 0)
all_true_test_calls_venn_df$XHMM <- ifelse(all_true_test_calls_strat2_df$XHMM_PROP > 0, 1, 0)
all_true_test_calls_venn_counts_df <- vennCounts(all_true_test_calls_venn_df)
all_true_test_calls_venn_counts_df[1,5] <- paste0("N=", sum(all_true_test_calls_venn_counts_df[,5]))

# Profile of all the samples in the test set classified as 'TRUE'
tp_strat2_rf_df <- sqldf("select * from tp_df where CLASSIFIER = 'RF' and NUM_SPLITS = 1 and STRATEGY = 2 and TRAINING_PROP = '0.1'")
tp_fp_strat2_rf_df <- sqldf("select * from tp_fp_df where CLASSIFIER = 'RF' and NUM_SPLITS = 1 and STRATEGY = 2 and TRAINING_PROP = '0.1'")

tp_fp_rf_venn_df <- data.frame(matrix(,ncol=0,nrow=dim(tp_fp_strat2_rf_df)[1]))
tp_fp_rf_venn_df$CANOES <- ifelse(tp_fp_strat2_rf_df$CANOES_PROP > 0, 1, 0)
tp_fp_rf_venn_df$CODEX <- ifelse(tp_fp_strat2_rf_df$CODEX_PROP > 0, 1, 0)
tp_fp_rf_venn_df$CLAMMS <- ifelse(tp_fp_strat2_rf_df$CLAMMS_PROP > 0, 1, 0)
tp_fp_rf_venn_df$XHMM <- ifelse(tp_fp_strat2_rf_df$XHMM_PROP > 0, 1, 0)
tp_fp_rf_venn_counts_df <- vennCounts(tp_fp_rf_venn_df)
tp_fp_rf_venn_counts_df[1,5] <- paste0("N=", sum(tp_fp_rf_venn_counts_df[,5]))

tp_rf_venn_df <- data.frame(matrix(,ncol=0,nrow=dim(tp_strat2_rf_df)[1]))
tp_rf_venn_df$CANOES <- ifelse(tp_strat2_rf_df$CANOES_PROP > 0, 1, 0)
tp_rf_venn_df$CODEX <- ifelse(tp_strat2_rf_df$CODEX_PROP > 0, 1, 0)
tp_rf_venn_df$CLAMMS <- ifelse(tp_strat2_rf_df$CLAMMS_PROP > 0, 1, 0)
tp_rf_venn_df$XHMM <- ifelse(tp_strat2_rf_df$XHMM_PROP > 0, 1, 0)
tp_rf_venn_counts_df <- vennCounts(tp_rf_venn_df)
tp_rf_venn_counts_df[1,5] <- paste0("N=", sum(tp_rf_venn_counts_df[,5]))

num_samples_in_testset <- sqldf("select count(*) from ma_sample_list_df where TRAINING_PROP = 0.1 and CLASSIFIER_NAME = 'RF' and STRATEGY = 2")
dyn_title = paste0("Every CNV in ", num_samples_in_testset, " samples in the test set")

pdf("fig_4a_truepositives_venn_randomforest_microarray_0.1.pdf", width = 12, height = 12)
par(mfrow=c(2,2),oma=c(0,2,2,0))
vennDiagram(all_test_calls_venn_counts_df, cex = c(2,2.5,2), lwd = 3, circle.col=venn_colors, include=c("both")) + 
  title(main = dyn_title, cex.main=1.5)
vennDiagram(all_true_test_calls_venn_counts_df, cex = c(2,2.5,2), lwd = 3, circle.col=venn_colors, include=c("both")) + 
  title(main = "Validated CNVs in the test set", cex.main=1.5)
vennDiagram(tp_fp_rf_venn_counts_df, cex = c(2,2.5,2), lwd = 3, circle.col=venn_colors, include=c("both")) + 
  title(main = "Every CNV classified as 'True' by CN-Learn \n (True Positives + False Positives) \n", cex.main=1.5)
vennDiagram(tp_rf_venn_counts_df, cex = c(2,2.5,2), lwd = 3, circle.col=venn_colors, include=c("both")) + 
  title("Only 'True' CNVs classified as 'True' by CN-Learn \n (True Positives) \n", cex.main=1.5)
mtext("      All Calls                                        Validated Calls", outer=TRUE, cex = 2.25, font = 2) 
mtext("  CN-Learn                                     All Callers", side = 2, outer=TRUE, cex = 2.25, font = 2) 
dev.off()


################################################################################################
# Figure 4B: Distribution of prediction scored by CNV size and concordance values (Microarray) #
################################################################################################
tp_fp_tn_fn_probs_size_ma_df <- sqldf("select TYPE, PRED_SIZE, NUM_OVERLAPS, PRED_PROBS, CLASS, VAL_LABEL
                                      from tp_fp_tn_fn_ma_df 
                                      where CLASSIFIER = 'RF' and STRATEGY = '2' and NUM_SPLITS = 1 and TRAINING_PROP = 0.7")

tp_fp_tn_fn_probs_size_ma_df$SIZE_LABEL[tp_fp_tn_fn_probs_size_ma_df$PRED_SIZE > 50000 & tp_fp_tn_fn_probs_size_ma_df$PRED_SIZE <= 100000] <- '50KB-100KB'
tp_fp_tn_fn_probs_size_ma_df$SIZE_LABEL[tp_fp_tn_fn_probs_size_ma_df$PRED_SIZE > 100000 & tp_fp_tn_fn_probs_size_ma_df$PRED_SIZE <= 250000] <- '100KB-250KB'
tp_fp_tn_fn_probs_size_ma_df$SIZE_LABEL[tp_fp_tn_fn_probs_size_ma_df$PRED_SIZE > 250000 & tp_fp_tn_fn_probs_size_ma_df$PRED_SIZE <= 500000] <- '250KB-500KB'
tp_fp_tn_fn_probs_size_ma_df$SIZE_LABEL[tp_fp_tn_fn_probs_size_ma_df$PRED_SIZE > 500000] <- '>500KB'

tp_fp_tn_fn_probs_size_ma_df$SIZE_LABEL <- ordered(tp_fp_tn_fn_probs_size_ma_df$SIZE_LABEL, levels = c("50KB-100KB", "100KB-250KB", "250KB-500KB", ">500KB"))

# Generate plot by size and concordance (MA) [Circles]
pdf("fig_4b_dist_of_tpfpCNVs_by_size_and_prob_concordance_microarray_0.7.pdf", width = 24, height = 16)
ggplot(data=tp_fp_tn_fn_probs_size_ma_df, aes(x=as.numeric(PRED_SIZE), y=as.numeric(PRED_PROBS), 
                                              color = as.factor(NUM_OVERLAPS), shape = VAL_LABEL)) + theme_classic() + 
  geom_point(size=4.5, stroke = 0.75) +
  facet_grid(~SIZE_LABEL, scale = "free") +
  scale_shape_manual(values=c(19, 1)) + 
  scale_color_manual(values=c("red", "purple", "green", "blue")) +
  theme(plot.title = element_text(lineheight=.8, size=9, face="bold", hjust = 0.5), 
        strip.text = element_text(size = 12, face = "bold"),
        axis.title = element_text(size = 12, face="bold"),
        axis.text = element_text(size = 12, face="bold"), 
        axis.text.x = element_text(angle = 45, hjust = 1),
        legend.title = element_text(size=12, face="bold"),
        legend.text = element_text(size=12, face="bold"),
        legend.position = "bottom",
        panel.spacing = unit(2, "lines")) +
  geom_hline(yintercept = 0.5, linetype="dotted") + 
  labs(color = "Concordance", x = "Prediction Size (Ascending Order)", y = "Prediction Probability", shape = "Validation") 
dev.off()


########################
# Supplementary Figures:
########################

#################################################################
# Supp Figure 4: Distribution of True Positives in each strategy
#################################################################
tp_fp_dist_all_strat_df <- sqldf("select A.Strategy, A.NUM_OVERLAPS, count(*) as Frequency, 
                                 round((count(*) * 100)/(B.Tot_Freq * 1.0), 1) as Proportion, round(B.Tot_Freq/(C.Tot_Freq * 1.0),3) * 100 as Prop,
                                 CASE WHEN A.STRATEGY = 2 THEN 'Microarray as Truth'
                                 WHEN A.STRATEGY = 3 THEN 'CLAMMS as Truth'
                                 ELSE 'CLAMMS Labels + Based on Read-Depth' END as Strategy_Name, A.VAL_LABEL, A.CLASS
                                 from tp_fp_df A, 
                                 (select Strategy, NUM_OVERLAPS, count(*) as Tot_Freq from tp_fp_df 
                                 where Classifier = 'RF' and Num_Splits = 10 and Training_Prop = 0.7
                                 group by Strategy, NUM_OVERLAPS) B,
                                 (select Strategy, count(*) as Tot_Freq from tp_fp_df 
                                 where Classifier = 'RF' and Num_Splits = 10 and Training_Prop = 0.7
                                 group by Strategy) C
                                 where A.Strategy in (2,3) and A.Classifier = 'RF' and A.Num_Splits = 10 and A.Training_Prop = 0.7
                                 and A.Strategy = B.Strategy and A.NUM_OVERLAPS = B.NUM_OVERLAPS
                                 and A.Strategy = C.Strategy 
                                 group by A.Strategy, A.NUM_OVERLAPS, A.VAL_LABEL, A.CLASS
                                 order by A.Strategy, A.NUM_OVERLAPS, A.VAL_LABEL desc")


tp_fp_dist_all_strat_df$Label <- paste0(tp_fp_dist_all_strat_df$Proportion, '%\n(', tp_fp_dist_all_strat_df$Frequency, ')')
tp_fp_dist_all_strat_df$VAL_LABEL <- ordered(tp_fp_dist_all_strat_df$VAL_LABEL , levels = c("True CNV", "False CNV"))
tp_fp_dist_all_strat_df$Strategy_Name <- ordered(tp_fp_dist_all_strat_df$Strategy_Name, levels=c("Microarray as Truth","CLAMMS as Truth"))
tp_fp_dist_all_strat_df$new_prop <- round((tp_fp_dist_all_strat_df$Proportion * tp_fp_dist_all_strat_df$Prop)/100, 1)
tp_fp_dist_all_strat_df$new_prop[c(1,3,5,7,9,11,13)] <-tp_fp_dist_all_strat_df$Prop[c(1,3,5,7,9,11,13)]


pdf("supp_fig_4_concordance_profile_of_truepositives_microarray_clamms_0.7.pdf", width = 6, height = 6)
ggplot(data=tp_fp_dist_all_strat_df, aes(x=factor(NUM_OVERLAPS), y=as.numeric(new_prop), fill = CLASS)) + theme_classic() +
  geom_bar(position = "dodge", stat="identity") + 
  #  geom_bar(aes(fill = factor(VAL_LABEL) ), stat="identity", position = position_stack(reverse = TRUE)) + 
  facet_grid(Strategy_Name~NUM_OVERLAPS, scale = "free", switch = "both") + ylim(0,50) +
  geom_text(aes(label=Label), position = position_dodge(width = 0.9), vjust = -.25, size = 3, fontface = "bold") +
  ggtitle ("Distribution of predictions classified as 'True' by CN-Learn") +
  scale_fill_manual(values = c("True Positive" = "blue","False Positive" = "red")) +
  theme(plot.title = element_text(lineheight=.8, size=9, face="bold", hjust = 0.5), 
        strip.text = element_text(size = 9, face = "bold"),
        axis.title = element_text(size = 9, face="bold"),
        axis.text = element_text(size = 9, face="bold"), 
        axis.text.x = element_blank(),
        legend.title = element_text(size=9, face="bold"),
        legend.text = element_text(size=9, face="bold"),
        legend.position = "bottom") +
  labs(x = "Number of Callers", y = "", fill = "") 
dev.off()


##############################################################
# Supp Figure 6B: Feature importance of CN-Learn with 1000G
##############################################################
rf_metrics_nocnvfreq_1000g_df <- read.csv("RF_Metrics_No_CNVFreq_1000G.csv", header = TRUE)
feat_imp_nocnvfreq_1000g_df <- sqldf ("select Training_Proportion, Chromosome, CNV_Type, Canoes_Proportion, Codex_Proportion, Clamms_Proportion, XHMM_Proportion, Overlap_Count, 
                                      RD_Proportion, GC_Content, Mappability, Number_Of_Targets, Size_Label
                                      from rf_metrics_nocnvfreq_1000g_df 
                                      where Classifier = 'RF' and Strategy = 2 and Num_Splits = 10 and Training_Proportion = 0.7")

melted_feat_imp_nocnvfreq_1000g_df <- melt(feat_imp_nocnvfreq_1000g_df, id=c("Training_Proportion"))
melted_feat_imp_nocnvfreq_1000g_df$prop <- melted_feat_imp_nocnvfreq_1000g_df$value *100
melted_feat_imp_nocnvfreq_1000g_df$pred_var_label <- c("Chromosome", "CNV Type", "CANOES", "CODEX", "CLAMMS", "XHMM", "Concordance", "Read Depth Ratio", "GC Content", 
                                                       "Mappability", "Target Probe Count", "Size Label")
melted_feat_imp_nocnvfreq_1000g_df <- sqldf("select * from melted_feat_imp_nocnvfreq_1000g_df order by prop desc")
melted_feat_imp_nocnvfreq_1000g_df$pred_var_label <- factor(melted_feat_imp_nocnvfreq_1000g_df$pred_var_label, 
                                                            levels = melted_feat_imp_nocnvfreq_1000g_df$pred_var_label[order(as.numeric(melted_feat_imp_nocnvfreq_1000g_df$prop))])


pdf("supp_fig_6b", width = 8, height = 5)
ggplot(data=melted_feat_imp_nocnvfreq_1000g_df, aes(x=reorder(pred_var_label,-prop), y=prop)) + theme_classic() +
  geom_bar(position = "dodge", stat="identity") + 
  geom_text(aes(label=paste0(prop,"%")), position = position_dodge(width = 0.9), vjust = -.25, size = 3.5, fontface = "bold") +
  ggtitle ("CN-Learn - Feature Importance (1000G)") +
  theme(plot.title = element_text(lineheight=.8, size=15, face="bold", hjust = 0.5), 
        strip.text = element_text(size = 12, face = "bold"),
        axis.title = element_text(size = 12, face="bold"),
        axis.text = element_text(size = 12, face="bold"), 
        #        axis.text.x = element_blank(),
        axis.text.x = element_text(angle = 60, hjust = 1),
        legend.title = element_text(size=12, face="bold"),
        legend.text = element_text(size=10, face="bold")) +
  labs(x = "Predictors", y = "Feature Importance", fill = "       Predictors") 
dev.off()

#####################################################################################
# Supp Figure 6C: Concordance profile of CNVs before and after classification (1000G)
#####################################################################################
venn_colors <- c("red", "blue", "green", "orchid")
strat2_nocnvfreq_1000g_df <- read.table("cons_overlap_strat2_0.1_1000g.txt", header = TRUE)
head(strat2_nocnvfreq_1000g_df)
dim(strat2_nocnvfreq_1000g_df)

sample_list_nocnvfreq_df <- as.data.frame(read.csv("Samplelist_No_CNVFreq_1000G.csv",header = FALSE))
colnames(sample_list_nocnvfreq_df) <- c("SAMPLE", "TRAINING_PROP", "CLASSIFIER_NAME", "STRATEGY")

all_test_calls_strat2_nocnvfreq_df <- sqldf("select a.* from  strat2_nocnvfreq_1000g_df a, sample_list_nocnvfreq_df b
                                            where a.PRED_SIZE >= 5000 and a.PRED_SIZE <= 5000000
                                            and b.TRAINING_PROP = 0.7 and b.CLASSIFIER_NAME = 'RF' and b.STRATEGY = 2
                                            and a.SAMPLE = b.SAMPLE")

tp_nocnvfreq_1000g_df <- read.csv("True_Pos_Calls_No_CNVFreq_1000G.csv")
fp_nocnvfreq_1000g_df <- read.csv("False_Pos_Calls_No_CNVFreq_1000G.csv")
tp_fp_nocnvfreq_1000g_df <- rbind(tp_nocnvfreq_1000g_df, fp_nocnvfreq_1000g_df)

all_test_calls_nocnvfreq_venn_df <- data.frame(matrix(,ncol=0,nrow=dim(all_test_calls_strat2_nocnvfreq_df)[1]))
all_test_calls_nocnvfreq_venn_df$CANOES <- ifelse(all_test_calls_strat2_nocnvfreq_df$CANOES_PROP > 0, 1, 0)
all_test_calls_nocnvfreq_venn_df$CODEX <- ifelse(all_test_calls_strat2_nocnvfreq_df$CODEX_PROP > 0, 1, 0)
all_test_calls_nocnvfreq_venn_df$CLAMMS <- ifelse(all_test_calls_strat2_nocnvfreq_df$CLAMMS_PROP > 0, 1, 0)
all_test_calls_nocnvfreq_venn_df$XHMM <- ifelse(all_test_calls_strat2_nocnvfreq_df$XHMM_PROP > 0, 1, 0)
all_test_calls_nocnvfreq_venn_counts_df <- vennCounts(all_test_calls_nocnvfreq_venn_df)
all_test_calls_nocnvfreq_venn_counts_df[1,5] <- paste0("N=", sum(all_test_calls_nocnvfreq_venn_counts_df[,5]))

all_true_test_calls_strat2_nocnvfreq_df <- sqldf("select a.* from strat2_nocnvfreq_1000g_df a, sample_list_nocnvfreq_df b
                                                 where a.PRED_SIZE > 5000 and a.PRED_SIZE < 5000000 and a.LABEL = 'TRUE_CNV'
                                                 and b.TRAINING_PROP = 0.7 and b.CLASSIFIER_NAME = 'RF' and b.STRATEGY = 2
                                                 and a.SAMPLE = b.SAMPLE and LABEL = 'TRUE_CNV'")

all_true_test_calls_nocnvfreq_venn_df <- data.frame(matrix(,ncol=0,nrow=dim(all_true_test_calls_strat2_nocnvfreq_df)[1]))
all_true_test_calls_nocnvfreq_venn_df$CANOES <- ifelse(all_true_test_calls_strat2_nocnvfreq_df$CANOES_PROP > 0, 1, 0)
all_true_test_calls_nocnvfreq_venn_df$CODEX <- ifelse(all_true_test_calls_strat2_nocnvfreq_df$CODEX_PROP > 0, 1, 0)
all_true_test_calls_nocnvfreq_venn_df$CLAMMS <- ifelse(all_true_test_calls_strat2_nocnvfreq_df$CLAMMS_PROP > 0, 1, 0)
all_true_test_calls_nocnvfreq_venn_df$XHMM <- ifelse(all_true_test_calls_strat2_nocnvfreq_df$XHMM_PROP > 0, 1, 0)
all_true_test_calls_nocnvfreq_venn_counts_df <- vennCounts(all_true_test_calls_nocnvfreq_venn_df)
all_true_test_calls_nocnvfreq_venn_counts_df[1,5] <- paste0("N=", sum(all_true_test_calls_nocnvfreq_venn_counts_df[,5]))

# Profile of all the samples in the test set classified as 'TRUE'
tp_strat2_nocnvfreq_rf_df <- sqldf("select * from tp_nocnvfreq_1000g_df where CLASSIFIER = 'RF' and NUM_SPLITS = 1 and STRATEGY = 2 and TRAINING_PROP = '0.7'")
tp_fp_strat2_nocnvfreq_rf_df <- sqldf("select * from tp_fp_nocnvfreq_1000g_df where CLASSIFIER = 'RF' and NUM_SPLITS = 1 and STRATEGY = 2 and TRAINING_PROP = '0.7'")

head(tp_1000g_df)

tp_fp_rf_nocnvfreq_venn_df <- data.frame(matrix(,ncol=0,nrow=dim(tp_fp_strat2_nocnvfreq_rf_df)[1]))
tp_fp_rf_nocnvfreq_venn_df$CANOES <- ifelse(tp_fp_strat2_nocnvfreq_rf_df$CANOES_PROP > 0, 1, 0)
tp_fp_rf_nocnvfreq_venn_df$CODEX <- ifelse(tp_fp_strat2_nocnvfreq_rf_df$CODEX_PROP > 0, 1, 0)
tp_fp_rf_nocnvfreq_venn_df$CLAMMS <- ifelse(tp_fp_strat2_nocnvfreq_rf_df$CLAMMS_PROP > 0, 1, 0)
tp_fp_rf_nocnvfreq_venn_df$XHMM <- ifelse(tp_fp_strat2_nocnvfreq_rf_df$XHMM_PROP > 0, 1, 0)
tp_fp_rf_nocnvfreq_venn_counts_df <- vennCounts(tp_fp_rf_nocnvfreq_venn_df)
tp_fp_rf_nocnvfreq_venn_counts_df[1,5] <- paste0("N=", sum(tp_fp_rf_nocnvfreq_venn_counts_df[,5]))

tp_rf_nocnvfreq_venn_df <- data.frame(matrix(,ncol=0,nrow=dim(tp_strat2_nocnvfreq_rf_df)[1]))
tp_rf_nocnvfreq_venn_df$CANOES <- ifelse(tp_strat2_nocnvfreq_rf_df$CANOES_PROP > 0, 1, 0)
tp_rf_nocnvfreq_venn_df$CODEX <- ifelse(tp_strat2_nocnvfreq_rf_df$CODEX_PROP > 0, 1, 0)
tp_rf_nocnvfreq_venn_df$CLAMMS <- ifelse(tp_strat2_nocnvfreq_rf_df$CLAMMS_PROP > 0, 1, 0)
tp_rf_nocnvfreq_venn_df$XHMM <- ifelse(tp_strat2_nocnvfreq_rf_df$XHMM_PROP > 0, 1, 0)
tp_rf_nocnvfreq_venn_counts_df <- vennCounts(tp_rf_nocnvfreq_venn_df)
tp_rf_nocnvfreq_venn_counts_df[1,5] <- paste0("N=", sum(tp_rf_nocnvfreq_venn_counts_df[,5]))

num_samples_in_testset_nocnvfreq <- sqldf("select count(*) from sample_list_nocnvfreq_df where TRAINING_PROP = 0.7 and CLASSIFIER_NAME = 'RF' and STRATEGY = 2")
dyn_title = paste0("Every CNV in ", num_samples_in_testset_nocnvfreq, " samples in the test set")

pdf("supp_fig_6c.pdf", width = 12, height = 12)
par(mfrow=c(2,2),oma=c(0,2,2,0))
vennDiagram(all_test_calls_nocnvfreq_venn_counts_df, cex = c(2,2.5,2), lwd = 3, circle.col=venn_colors, include=c("both")) + 
  title(main = dyn_title, cex.main=1.5)
vennDiagram(all_true_test_calls_nocnvfreq_venn_counts_df, cex = c(2,2.5,2), lwd = 3, circle.col=venn_colors, include=c("both")) + 
  title(main = "Validated CNVs in the test set", cex.main=1.5)
vennDiagram(tp_fp_rf_nocnvfreq_venn_counts_df, cex = c(2,2.5,2), lwd = 3, circle.col=venn_colors, include=c("both")) + 
  title(main = "Every CNV classified as 'True' by CN-Learn \n (True Positives + False Positives) \n", cex.main=1.5)
vennDiagram(tp_rf_nocnvfreq_venn_counts_df, cex = c(2,2.5,2), lwd = 3, circle.col=venn_colors, include=c("both")) + 
  title("Only 'True' CNVs classified as 'True' by CN-Learn \n (True Positives) \n", cex.main=1.5)
mtext("      All Calls                                        Validated Calls", outer=TRUE, cex = 2.25, font = 2) 
mtext("  CN-Learn                                     All Callers", side = 2, outer=TRUE, cex = 2.25, font = 2) 
dev.off()


##############################################################
# Supp Figure 9: Performance for CNVs in different freq bins
##############################################################
rf_metrics_cnvfreq_df <- read.csv("RF_Metrics_CNVFreq_SVIP.csv")

cnvfreq_perf_df <- sqldf("select Training_Data, CNV_Freq, Precision, Recall, CNV_Count_Train, CNV_Count_Test 
                         from rf_metrics_cnvfreq_df where Num_Splits = 10 and Training_Prop = 0.7
                         order by Training_Data")
cnvfreq_perf_df$Training_Data <- c(rep("Training Data = All CNVs",4),rep("Training Data = Very Rare CNVs",4))
cnvfreq_perf_df$CNV_Freq <- rep(c("Very Rare CNVs", "Rare CNVs", "Common CNVs", "Very Common CNVs"),2)
cnvfreq_perf_df$Training_Data <- paste0(cnvfreq_perf_df$Training_Data, " (", cnvfreq_perf_df$CNV_Count_Train, ")")
cnvfreq_perf_df$CNV_Freq <- paste0(cnvfreq_perf_df$CNV_Freq, " (", cnvfreq_perf_df$CNV_Count_Test, ")")
cnvfreq_perf_df <- cnvfreq_perf_df[,-c(5,6)]

melted_cnvfreq_perf_df <- melt(cnvfreq_perf_df, id = c("Training_Data", "CNV_Freq"))
melted_cnvfreq_perf_df$CNV_Freq <- ordered(melted_cnvfreq_perf_df$CNV_Freq, levels = c("Very Rare CNVs (228)", "Rare CNVs (163)", 
                                                                                       "Common CNVs (141)", "Very Common CNVs (205)"))

pdf("supp_fig_9.pdf", width = 9, height = 5)
ggplot(data=melted_cnvfreq_perf_df, aes(x=CNV_Freq, y=value, fill = variable)) + theme_classic() +
  geom_bar(position = "dodge", stat="identity")  +
  facet_grid(~Training_Data, scale = "free", switch = "x") +
  geom_text(aes(label=value), position = position_dodge(width = 0.9), vjust = -.25, size = 3.5, fontface = "bold") +
  ggtitle ("CN-Learn - Performance") +
  theme(plot.title = element_text(lineheight=.8, size=12, face="bold", hjust = 0.5), 
        strip.text = element_text(size = 10, face = "bold"),
        axis.title = element_text(size = 10, face="bold"),
        axis.text = element_text(size = 10, face="bold"), 
        #axis.text.x = element_blank(),
        axis.text.x = element_text(angle = 40, hjust = 1),
        legend.title = element_text(size=10, face="bold"),
        legend.text = element_text(size=10, face="bold")) +
  labs(x = "CNV Frequency", y = "", fill = "") 
dev.off()

############################################################
# Supp Figure 10: Precision-Recall with and without CNV freq
############################################################
rf_metrics_nocnvfreq_1000g_df <- read.csv("RF_Metrics_No_CNVFreq_1000G.csv")
rf_metrics_nocnvfreq_1000g_df <- sqldf("select Training_Proportion, Precision, Recall from rf_metrics_nocnvfreq_1000g_df where Num_Splits = 10 and Classifier = 'RF'")
rf_metrics_nocnvfreq_1000g_df$Training_Proportion <- paste0(rf_metrics_nocnvfreq_1000g_df$Training_Proportion, " (", c("9","18","27","36","45","54","62"), " Samples)")
melted_rf_metrics_nocnvfreq_1000g_df <- melt(rf_metrics_nocnvfreq_1000g_df, id = "Training_Proportion")
melted_rf_metrics_nocnvfreq_1000g_df$TYPE <- 'Without CNV frequency as a predictor'

rf_metrics_cnvfreq_1000g_df <- read.csv("RF_Metrics_1000G.csv", header = TRUE)
rf_metrics_cnvfreq_1000g_df <- sqldf("select Training_Proportion, Precision, Recall from rf_metrics_cnvfreq_1000g_df where Num_Splits = 10 and Classifier = 'RF'")
rf_metrics_cnvfreq_1000g_df$Training_Proportion <- paste0(rf_metrics_cnvfreq_1000g_df$Training_Proportion, " (", c("9","18","27","36","45","54","62"), " Samples)")
melted_rf_metrics_cnvfreq_1000g_df <- melt(rf_metrics_cnvfreq_1000g_df, id = "Training_Proportion")
melted_rf_metrics_cnvfreq_1000g_df$TYPE <- 'With CNV frequency as a predictor'

rf_metrics_1000g_cnvfreq_df <- rbind(melted_rf_metrics_nocnvfreq_1000g_df, melted_rf_metrics_cnvfreq_1000g_df)

pdf("supp_fig_10.pdf", width = 9, height = 8)
ggplot(data=rf_metrics_1000g_cnvfreq_df, aes(x=Training_Proportion, y=value, fill = variable)) + theme_classic() +
  geom_bar(position = "dodge", stat="identity")  + ylim(0,1.1) + 
  facet_grid(TYPE~., scale = "free", switch = "x") +
  geom_text(aes(label=value), position = position_dodge(width = 0.9), vjust = -.25, size = 3.5, fontface = "bold") +
  ggtitle ("CN-Learn - Performance") +
  theme(plot.title = element_text(lineheight=.8, size=12, face="bold", hjust = 0.5), 
        strip.text = element_text(size = 10, face = "bold"),
        axis.title = element_text(size = 10, face="bold"),
        axis.text = element_text(size = 10, face="bold"), 
        #axis.text.x = element_blank(),
        axis.text.x = element_text(angle = 40, hjust = 1),
        legend.title = element_text(size=10, face="bold"),
        legend.text = element_text(size=10, face="bold")) +
  labs(x = "Training Proportion", y = "Precision/Recall", fill = "") 
dev.off()

##############################################################
# Supp Figure 11: Feature importance with and without CNV freq
##############################################################
rf_metrics_nocnvfreq_1000g_df <- read.csv("RF_Metrics_No_CNVFreq_1000G.csv", header = TRUE)
feat_imp_nocnvfreq_1000g_df <- sqldf ("select Training_Proportion, Chromosome, CNV_Type, Canoes_Proportion, Codex_Proportion, Clamms_Proportion, XHMM_Proportion, Overlap_Count, 
                                      RD_Proportion, GC_Content, Mappability, Number_Of_Targets, Size_Label, 0 as CNV_Frequency
                                      from rf_metrics_nocnvfreq_1000g_df 
                                      where Classifier = 'RF' and Strategy = 2 and Num_Splits = 10 and Training_Proportion = 0.7")

melted_feat_imp_nocnvfreq_1000g_df <- melt(feat_imp_nocnvfreq_1000g_df, id=c("Training_Proportion"))
melted_feat_imp_nocnvfreq_1000g_df$prop <- melted_feat_imp_nocnvfreq_1000g_df$value *100
melted_feat_imp_nocnvfreq_1000g_df$pred_var_label <- c("Chromosome", "CNV Type", "CANOES", "CODEX", "CLAMMS", "XHMM", "Concordance", "Read Depth Ratio", "GC Content", 
                                                       "Mappability", "Target Probe Count", "Size Label", "CNV Frequency")
melted_feat_imp_nocnvfreq_1000g_df <- sqldf("select * from melted_feat_imp_nocnvfreq_1000g_df order by prop desc")
melted_feat_imp_nocnvfreq_1000g_df$TYPE <- 'Without CNV frequency as a predictor'

rf_metrics_cnvfreq_1000g_df <- read.csv("RF_Metrics_1000G.csv", header = TRUE)
feat_imp_cnvfreq_1000g_df <- sqldf ("select Training_Proportion, Chromosome, CNV_Type, Canoes_Proportion, Codex_Proportion, Clamms_Proportion, XHMM_Proportion, Overlap_Count, 
                                    RD_Proportion, GC_Content, Mappability, Number_Of_Targets, Size_Label, CNV_Frequency
                                    from rf_metrics_cnvfreq_1000g_df 
                                    where Classifier = 'RF' and Strategy = 2 and Num_Splits = 10 and Training_Proportion = 0.7")

melted_feat_imp_cnvfreq_1000g_df <- melt(feat_imp_cnvfreq_1000g_df, id=c("Training_Proportion"))
melted_feat_imp_cnvfreq_1000g_df$prop <- melted_feat_imp_cnvfreq_1000g_df$value *100
melted_feat_imp_cnvfreq_1000g_df$pred_var_label <- c("Chromosome", "CNV Type", "CANOES", "CODEX", "CLAMMS", "XHMM", "Concordance", "Read Depth Ratio", "GC Content", 
                                                     "Mappability", "Target Probe Count", "Size Label", "CNV Frequency")
melted_feat_imp_cnvfreq_1000g_df <- sqldf("select * from melted_feat_imp_cnvfreq_1000g_df order by prop desc")

melted_feat_imp_cnvfreq_1000g_df$TYPE <- 'With CNV frequency as a predictor'
melted_1000g_feat_imp_df <- rbind(melted_feat_imp_nocnvfreq_1000g_df, melted_feat_imp_cnvfreq_1000g_df)
melted_1000g_feat_imp_df$pred_var_label <- ordered(melted_1000g_feat_imp_df$pred_var_label, levels = c("Mappability", "CNV Frequency", "Read Depth Ratio",
                                                                                                       "Chromosome", "GC Content", "Target Probe Count",
                                                                                                       "CLAMMS", "Size Label", "CNV Type", "Concordance",
                                                                                                       "CODEX", "XHMM", "CANOES"))


pdf("supp_fig_11.pdf", width = 10, height = 9)
ggplot(data=melted_1000g_feat_imp_df, aes(x=pred_var_label, y=prop)) + theme_classic() +
  geom_bar(position = "dodge", stat="identity") + 
  geom_text(aes(label=paste0(prop,"%")), position = position_dodge(width = 0.9), vjust = -.25, size = 3.5, fontface = "bold") +
  facet_grid(TYPE~., scale = "free", switch = "x") + ylim(0,19) + 
  ggtitle ("CN-Learn - Feature Importance (1000G)") +
  theme(plot.title = element_text(lineheight=.8, size=15, face="bold", hjust = 0.5), 
        strip.text = element_text(size = 12, face = "bold"),
        axis.title = element_text(size = 12, face="bold"),
        axis.text = element_text(size = 12, face="bold"), 
        #        axis.text.x = element_blank(),
        axis.text.x = element_text(angle = 60, hjust = 1),
        legend.title = element_text(size=12, face="bold"),
        legend.text = element_text(size=10, face="bold")) +
  labs(x = "Predictors", y = "Feature Importance", fill = "       Predictors") 
dev.off()

############################################################
# Supp Figure 12: Precision-Recall values of each classifier
############################################################
prec_rec_all_classifiers <- sqldf("select Precision, Recall, 
                                  CASE WHEN STRATEGY = 2 THEN 'Microarray as Truth'
                                  WHEN STRATEGY = 3 THEN 'CLAMMS as Truth' END as Strategy_Name,
                                  CASE WHEN Classifier = 'RF' THEN 'Random Forest' WHEN Classifier = 'LR' THEN 'Logistic Regression'
                                  WHEN Classifier = 'SVM' THEN 'Support Vector Machine' END as Classifier_Name
                                  from rf_metrics_df where Training_Proportion = 0.7 and Num_Splits = 10 and STRATEGY in (2,3)")
melted_prec_rec_all_classifiers <- melt(prec_rec_all_classifiers, id = c("Classifier_Name", "Strategy_Name"))
melted_prec_rec_all_classifiers$Strategy_Name <- ordered(melted_prec_rec_all_classifiers$Strategy_Name, levels=c("Microarray as Truth","CLAMMS as Truth"))
melted_prec_rec_all_classifiers$Classifier_Name <- ordered(melted_prec_rec_all_classifiers$Classifier_Name, levels = c("Logistic Regression",
                                                                                                                       "Support Vector Machine", "Random Forest"))

pdf("supp_fig_12_prec_rec_RF_SVM_LR_microarray_clamms_0.7.pdf", width = 6, height = 5)
ggplot(data=melted_prec_rec_all_classifiers, aes(x=Classifier_Name, y=value, fill=Classifier_Name)) + theme_classic() +
  geom_bar(position = "dodge", stat="identity") + 
  #  geom_errorbar(aes(ymin=value-0.1, ymax=value+0.1),width=.2,position=position_dodge(.9)) +
  facet_grid(Strategy_Name~variable, scale = "free", switch = "both") + ylim(0,1.1) +
  geom_text(aes(label=value), position = position_dodge(width = 0.9), vjust = -.25, size = 3, fontface = "bold") +
  ggtitle ("Precision-Recall values of three different classifiers within CN-Learn") +
  theme(plot.title = element_text(lineheight=.8, size=9, face="bold", hjust = 0.5), 
        strip.text = element_text(size = 9, face = "bold"),
        axis.title = element_text(size = 9, face="bold"),
        axis.text = element_text(size = 9, face="bold"), 
        axis.text.x = element_blank(),
        legend.title = element_text(size=9, face="bold"),
        legend.text = element_text(size=9, face="bold"),
        legend.position = "bottom") +
  labs(x = "", y = "Classifier Performance", fill = "Classifier") 
dev.off()

#############################################################
# Supp Figure 15: Feature Importances for all the strategies
#############################################################
feat_imp_all_strat <- sqldf ("select Training_Proportion, Chromosome, CNV_Type, Canoes_Proportion, Codex_Proportion, Clamms_Proportion, XHMM_Proportion, 
                             Overlap_Count, RD_Proportion, GC_Content, Mappability, Number_Of_Targets, Size_Label, 'Genomic' as Predictor_Group,
                             CASE WHEN STRATEGY = 2 THEN 'Microarray as Truth'
                             WHEN STRATEGY = 3 THEN 'CLAMMS as Truth'
                             ELSE 'CLAMMS Labels + Based on Read-Depth' END as Strategy_Name
                             from rf_metrics_df 
                             where strategy in (2,3) and
                             classifier = 'RF' and Num_Splits = 10 and Training_Proportion = 0.7")

melted_feat_imp_all_strat <- melt(feat_imp_all_strat, id=c("Training_Proportion", "Strategy_Name", "Predictor_Group"))
melted_feat_imp_all_strat$prop <- melted_feat_imp_all_strat$value *100
melted_feat_imp_all_strat$Predictor_Group[5:14] <- "Caller"

melted_feat_imp_all_strat$pred_var_label <- rep(c("Chromosome", "CNV Type", "CANOES", "CODEX", "CLAMMS", "XHMM", "Concordance", "Read Depth Ratio", "GC Content", 
                                                  "Mappability", "Target Probe Count", "Size Label"), each = 2)
melted_feat_imp_all_strat <- sqldf("select * from melted_feat_imp_all_strat order by Strategy_Name desc, prop desc")
melted_feat_imp_all_strat$pred_var_label <- ordered(melted_feat_imp_all_strat$pred_var_label, levels = c("CANOES", "CLAMMS", "CODEX", "XHMM", "Concordance", 
                                                                                                         "Chromosome", "CNV Type", "GC Content", "Mappability", "Target Probe Count", "Size Label", "Read Depth Ratio"))

pdf("supp_fig_15_feature_importance_microarray_clamms_0.7.pdf", width = 8, height = 7)
ggplot(data=melted_feat_imp_all_strat, aes(x=pred_var_label, y=prop)) + theme_classic() +
  geom_bar(position = "dodge", stat="identity") + ylim(0,30) +
  facet_grid(Strategy_Name~., scale = "free", switch = "both") +
  geom_text(aes(label=paste0(prop,"%")), position = position_dodge(width = 0.9), vjust = -.25, size = 3.5, fontface = "bold") +
  ggtitle ("Feature Importance") +
  theme(plot.title = element_text(lineheight=.8, size=15, face="bold", hjust = 0.5), 
        strip.text = element_text(size = 12, face = "bold"),
        axis.title = element_text(size = 12, face="bold"),
        axis.text = element_text(size = 12, face="bold"), 
        #axis.text.x = element_blank(),
        axis.text.x = element_text(angle = 40, hjust = 1),
        legend.title = element_text(size=12, face="bold"),
        legend.text = element_text(size=10, face="bold")) +
  labs(x = "Predictors", y = "Feature Importance", fill = "       Predictors") 
dev.off()

###################################################################################################################################
# Supp Fig 16: Distribution of all calls in test samples vs. classified as 'True' with CLAMMS validation (Venn Diagrams) (CLAMMS) #
###################################################################################################################################
# Profile of all the samples in the test set classified as 'TRUE'
# Profile of calls made in all the samples in the test set
all_test_calls_strat3_df <- sqldf("select a.* from  strat3_df a, clamms_sample_list_df b
                                  where a.PRED_SIZE > 5000 and a.PRED_SIZE < 5000000
                                  and b.TRAINING_PROP = 0.1 and b.CLASSIFIER_NAME = 'RF' and b.STRATEGY = 3
                                  and a.SAMPLE = b.SAMPLE")

all_test_calls_clamms_venn_df <- data.frame(matrix(,ncol=0,nrow=dim(all_test_calls_strat3_df)[1]))
all_test_calls_clamms_venn_df$CANOES <- ifelse(all_test_calls_strat3_df$CANOES_PROP > 0, 1, 0)
all_test_calls_clamms_venn_df$CODEX <- ifelse(all_test_calls_strat3_df$CODEX_PROP > 0, 1, 0)
all_test_calls_clamms_venn_df$XHMM <- ifelse(all_test_calls_strat3_df$XHMM_PROP > 0, 1, 0)
all_test_calls_clamms_venn_counts_df <- vennCounts(all_test_calls_clamms_venn_df)
all_test_calls_clamms_venn_counts_df[1,4] <- paste0("N=", sum(all_test_calls_clamms_venn_counts_df[,4]))

all_true_test_calls_strat3_df <- sqldf("select a.* from strat3_df a, clamms_sample_list_df b
                                       where a.PRED_SIZE > 5000 and a.PRED_SIZE < 5000000 and a.LABEL = 'TRUE_CNV'
                                       and b.TRAINING_PROP = 0.1 and b.CLASSIFIER_NAME = 'RF' and b.STRATEGY = 3
                                       and a.SAMPLE = b.SAMPLE and LABEL = 'TRUE_CNV'")
all_true_test_calls_clamms_venn_df <- data.frame(matrix(,ncol=0,nrow=dim(all_true_test_calls_strat3_df)[1]))
all_true_test_calls_clamms_venn_df$CANOES <- ifelse(all_true_test_calls_strat3_df$CANOES_PROP > 0, 1, 0)
all_true_test_calls_clamms_venn_df$CODEX <- ifelse(all_true_test_calls_strat3_df$CODEX_PROP > 0, 1, 0)
all_true_test_calls_clamms_venn_df$XHMM <- ifelse(all_true_test_calls_strat3_df$XHMM_PROP > 0, 1, 0)
all_true_test_calls_clamms_venn_counts_df <- vennCounts(all_true_test_calls_clamms_venn_df)
all_true_test_calls_clamms_venn_counts_df[1,4] <- paste0("N=", sum(all_true_test_calls_clamms_venn_counts_df[,4]))

tp_strat3_rf_df <- sqldf("select * from tp_df where CLASSIFIER = 'RF' and NUM_SPLITS = 1 and STRATEGY = 3 and TRAINING_PROP = '0.1'")
tp_fp_strat3_rf_df <- sqldf("select * from tp_fp_df where CLASSIFIER = 'RF' and NUM_SPLITS = 1 and STRATEGY = 3 and TRAINING_PROP = '0.1'")

tp_strat3_svm_df <- sqldf("select * from tp_df where CLASSIFIER = 'SVM' and NUM_SPLITS = 1 and STRATEGY = 3 and TRAINING_PROP = '0.1'")
tp_fp_strat3_svm_df <- sqldf("select * from tp_fp_df where CLASSIFIER = 'SVM' and NUM_SPLITS = 1 and STRATEGY = 3 and TRAINING_PROP = '0.1'")

tp_strat3_lr_df <- sqldf("select * from tp_df where CLASSIFIER = 'LR' and NUM_SPLITS = 1 and STRATEGY = 3 and TRAINING_PROP = '0.1'")
tp_fp_strat3_lr_df <- sqldf("select * from tp_fp_df where CLASSIFIER = 'LR' and NUM_SPLITS = 1 and STRATEGY = 3 and TRAINING_PROP = '0.1'")

tp_fp_rf_venn_clamms_df <- data.frame(matrix(,ncol=0,nrow=dim(tp_fp_strat3_rf_df)[1]))
tp_fp_rf_venn_clamms_df$CANOES <- ifelse(tp_fp_strat3_rf_df$CANOES_PROP > 0, 1, 0)
tp_fp_rf_venn_clamms_df$CODEX <- ifelse(tp_fp_strat3_rf_df$CODEX_PROP > 0, 1, 0)
tp_fp_rf_venn_clamms_df$XHMM <- ifelse(tp_fp_strat3_rf_df$XHMM_PROP > 0, 1, 0)
tp_fp_rf_venn_clamms_counts_df <- vennCounts(tp_fp_rf_venn_clamms_df)
tp_fp_rf_venn_clamms_counts_df[1,4] <- paste0("N=", sum(tp_fp_rf_venn_clamms_counts_df[,4]))

tp_rf_venn_clamms_df <- data.frame(matrix(,ncol=0,nrow=dim(tp_strat3_rf_df)[1]))
tp_rf_venn_clamms_df$CANOES <- ifelse(tp_strat3_rf_df$CANOES_PROP > 0, 1, 0)
tp_rf_venn_clamms_df$CODEX <- ifelse(tp_strat3_rf_df$CODEX_PROP > 0, 1, 0)
tp_rf_venn_clamms_df$XHMM <- ifelse(tp_strat3_rf_df$XHMM_PROP > 0, 1, 0)
tp_rf_venn_clamms_counts_df <- vennCounts(tp_rf_venn_clamms_df)
tp_rf_venn_clamms_counts_df[1,4] <- paste0("N=", sum(tp_rf_venn_clamms_counts_df[,4]))

tp_fp_svm_venn_clamms_df <- data.frame(matrix(,ncol=0,nrow=dim(tp_fp_strat3_svm_df)[1]))
tp_fp_svm_venn_clamms_df$CANOES <- ifelse(tp_fp_strat3_svm_df$CANOES_PROP > 0, 1, 0)
tp_fp_svm_venn_clamms_df$CODEX <- ifelse(tp_fp_strat3_svm_df$CODEX_PROP > 0, 1, 0)
tp_fp_svm_venn_clamms_df$XHMM <- ifelse(tp_fp_strat3_svm_df$XHMM_PROP > 0, 1, 0)
tp_fp_svm_venn_clamms_counts_df <- vennCounts(tp_fp_svm_venn_clamms_df)
tp_fp_svm_venn_clamms_counts_df[1,4] <- paste0("N=", sum(tp_fp_svm_venn_clamms_counts_df[,4]))

tp_svm_venn_clamms_df <- data.frame(matrix(,ncol=0,nrow=dim(tp_strat3_svm_df)[1]))
tp_svm_venn_clamms_df$CANOES <- ifelse(tp_strat3_svm_df$CANOES_PROP > 0, 1, 0)
tp_svm_venn_clamms_df$CODEX <- ifelse(tp_strat3_svm_df$CODEX_PROP > 0, 1, 0)
tp_svm_venn_clamms_df$XHMM <- ifelse(tp_strat3_svm_df$XHMM_PROP > 0, 1, 0)
tp_svm_venn_clamms_counts_df <- vennCounts(tp_svm_venn_clamms_df)
tp_svm_venn_clamms_counts_df[1,4] <- paste0("N=", sum(tp_svm_venn_clamms_counts_df[,4]))

tp_fp_lr_venn_clamms_df <- data.frame(matrix(,ncol=0,nrow=dim(tp_fp_strat3_lr_df)[1]))
tp_fp_lr_venn_clamms_df$CANOES <- ifelse(tp_fp_strat3_lr_df$CANOES_PROP > 0, 1, 0)
tp_fp_lr_venn_clamms_df$CODEX <- ifelse(tp_fp_strat3_lr_df$CODEX_PROP > 0, 1, 0)
tp_fp_lr_venn_clamms_df$XHMM <- ifelse(tp_fp_strat3_lr_df$XHMM_PROP > 0, 1, 0)
tp_fp_lr_venn_clamms_counts_df <- vennCounts(tp_fp_lr_venn_clamms_df)
tp_fp_lr_venn_clamms_counts_df[1,4] <- paste0("N=", sum(tp_fp_lr_venn_clamms_counts_df[,4]))

tp_lr_venn_clamms_df <- data.frame(matrix(,ncol=0,nrow=dim(tp_strat3_lr_df)[1]))
tp_lr_venn_clamms_df$CANOES <- ifelse(tp_strat3_lr_df$CANOES_PROP > 0, 1, 0)
tp_lr_venn_clamms_df$CODEX <- ifelse(tp_strat3_lr_df$CODEX_PROP > 0, 1, 0)
tp_lr_venn_clamms_df$XHMM <- ifelse(tp_strat3_lr_df$XHMM_PROP > 0, 1, 0)
tp_lr_venn_clamms_counts_df <- vennCounts(tp_lr_venn_clamms_df)
tp_lr_venn_clamms_counts_df[1,4] <- paste0("N=", sum(tp_lr_venn_clamms_counts_df[,4]))

num_samples_in_testset <- sqldf("select count(*) from clamms_sample_list_df where TRAINING_PROP = 0.1 and CLASSIFIER_NAME = 'RF' and STRATEGY = 3")
dyn_title = paste0("Every CNV in ", num_samples_in_testset, " samples in the test set")

pdf("supp_fig_16_tp_venn_randomforest_clamms_0.1.pdf", width = 12, height = 12)
par(mfrow=c(2,2),oma=c(0,2,2,0))
vennDiagram(all_test_calls_clamms_venn_counts_df, cex = c(2,2.5,2), lwd = 3, circle.col=venn_colors) + 
  title(main = dyn_title , cex.main=1.25)
vennDiagram(all_true_test_calls_clamms_venn_counts_df, cex = c(2,2.5,2), lwd = 3, circle.col=venn_colors) + 
  title(main = "Only the 'True' CNVs in the test set", cex.main=1.25)
vennDiagram(tp_fp_rf_venn_clamms_counts_df, cex = c(2,2.5,2), lwd = 3, circle.col=venn_colors) + 
  title(main = "Every CNV classified as 'True' by CN-Learn (RF) \n (True Positives + False Positives)\n", cex.main=1.25)
vennDiagram(tp_rf_venn_clamms_counts_df, cex = c(2,2.5,2), lwd = 3, circle.col=venn_colors) + 
  title("Only 'True' CNVs classified as 'True' by CN-Learn (RF) \n (True Positives)\n", cex.main=1.25)
mtext("      All Calls                                        Validated Calls", outer=TRUE, cex = 2.25, font = 2) 
mtext("  CN-Learn                                     All Callers", side = 2, outer=TRUE, cex = 2.25, font = 2) 
dev.off()

#################################################################################################
# Supp Figure 17: Distribution of prediction scored by CNV size and concordance values (CLAMMS) #
#################################################################################################
tp_fp_tn_fn_probs_size_clamms_df <- sqldf("select TYPE, PRED_SIZE, NUM_OVERLAPS, PRED_PROBS, CLASS, VAL_LABEL
                                          from tp_fp_tn_fn_clamms_df 
                                          where CLASSIFIER = 'RF' and STRATEGY = '3' and NUM_SPLITS = 1 and TRAINING_PROP = 0.7")

tp_fp_tn_fn_probs_size_clamms_df$SIZE_LABEL[tp_fp_tn_fn_probs_size_clamms_df$PRED_SIZE > 5000 & tp_fp_tn_fn_probs_size_clamms_df$PRED_SIZE <= 25000] <- '5KB-25KB'
tp_fp_tn_fn_probs_size_clamms_df$SIZE_LABEL[tp_fp_tn_fn_probs_size_clamms_df$PRED_SIZE > 25000 & tp_fp_tn_fn_probs_size_clamms_df$PRED_SIZE <= 50000] <- '25KB-50KB'
tp_fp_tn_fn_probs_size_clamms_df$SIZE_LABEL[tp_fp_tn_fn_probs_size_clamms_df$PRED_SIZE > 50000 & tp_fp_tn_fn_probs_size_clamms_df$PRED_SIZE <= 100000] <- '50KB-100KB'
tp_fp_tn_fn_probs_size_clamms_df$SIZE_LABEL[tp_fp_tn_fn_probs_size_clamms_df$PRED_SIZE > 100000 & tp_fp_tn_fn_probs_size_clamms_df$PRED_SIZE <= 250000] <- '100KB-250KB'
tp_fp_tn_fn_probs_size_clamms_df$SIZE_LABEL[tp_fp_tn_fn_probs_size_clamms_df$PRED_SIZE > 250000 & tp_fp_tn_fn_probs_size_clamms_df$PRED_SIZE <= 500000] <- '250KB-500KB'
tp_fp_tn_fn_probs_size_clamms_df$SIZE_LABEL[tp_fp_tn_fn_probs_size_clamms_df$PRED_SIZE > 500000] <- '>500KB'

sqldf("select count(*) from tp_fp_tn_fn_probs_size_clamms_df where PRED_SIZE < 25000")

tp_fp_tn_fn_probs_size_clamms_df$SIZE_LABEL <- ordered(tp_fp_tn_fn_probs_size_clamms_df$SIZE_LABEL, levels = c("5KB-25KB","25KB-50KB", "50KB-100KB", "100KB-250KB", "250KB-500KB", ">500KB"))

pdf("supp_fig_17_dist_of_tpfpCNVs_by_size_and_prob_concordance_clamms_0.7.pdf", width = 24, height = 16)
ggplot(data=tp_fp_tn_fn_probs_size_clamms_df, aes(x=as.numeric(PRED_SIZE), y=as.numeric(PRED_PROBS), 
                                                  color = as.factor(NUM_OVERLAPS), shape = VAL_LABEL)) + theme_classic() + 
  geom_point(size=4.5, stroke = 0.75) +
  facet_grid(~SIZE_LABEL, scale = "free") +
  scale_shape_manual(values=c(19, 1)) + 
  scale_color_manual(values=c("red", "purple", "green", "blue")) +
  theme(plot.title = element_text(lineheight=.8, size=9, face="bold", hjust = 0.5), 
        strip.text = element_text(size = 12, face = "bold"),
        axis.title = element_text(size = 12, face="bold"),
        axis.text = element_text(size = 12, face="bold"), 
        axis.text.x = element_text(angle = 45, hjust = 1),
        legend.title = element_text(size=12, face="bold"),
        legend.text = element_text(size=12, face="bold"),
        legend.position = "bottom",
        panel.spacing = unit(2, "lines")) +
  geom_hline(yintercept = 0.5, linetype="dotted") + 
  labs(color = "Concordance", x = "Prediction Size (Ascending Order)", y = "Prediction Probability", shape = "Validation") 
dev.off()

###############################################################
# Supp Figure 18: Calls by Overlap Count & Label (Original Data)
###############################################################
calls_by_ovcount_vallab <- sqldf("select A.CALLER, A.NUM_OVERLAPS, A.LABEL, A.COUNT, round((A.COUNT/(B.COUNT * 1.0) * 100)) as PROP from 
                                 (select CALLER, NUM_OVERLAPS, LABEL, count(*) as COUNT from retained_calls_gt50kb_df group by CALLER, NUM_OVERLAPS, LABEL) A,
                                 (select CALLER, NUM_OVERLAPS, count(*) as COUNT from retained_calls_gt50kb_df group by CALLER, NUM_OVERLAPS) B
                                 where A.CALLER = B.CALLER and A.NUM_OVERLAPS = B.NUM_OVERLAPS")
calls_by_ovcount_vallab$label_val <- paste0(calls_by_ovcount_vallab$PROP, "%\n(", calls_by_ovcount_vallab$COUNT, ')')

pdf("supp_fig_18_concordance_by_validationlabel_w_microarrays.pdf", width = 10, height = 5)
ggplot(data=calls_by_ovcount_vallab, aes(x=as.character(NUM_OVERLAPS), y=as.numeric(PROP), fill=CALLER)) + theme_classic() +
  geom_bar(position = "dodge", stat="identity") + 
  facet_grid(LABEL~NUM_OVERLAPS, scale = "free", switch = "both") + ylim(0,120) + 
  geom_text(aes(label=label_val), position = position_dodge(width = 1), vjust = -.25, size = 2.75, hjust = 0.5, fontface = "bold") + 
  ggtitle ("Distribution of the original calls by concordance and label") + 
  theme(plot.title = element_text(lineheight=.8, size=15, face="bold", hjust = 0.5), 
        strip.text = element_text(size = 12, face = "bold"),
        axis.title = element_text(size = 12, face="bold"),
        axis.text = element_text(size = 12, face="bold"), 
        axis.text.x = element_blank(),
        legend.title = element_text(size=12, face="bold"),
        legend.text = element_text(size=10, face="bold"),
        legend.position = "bottom") +
  labs(fill = "", x = "Number of callers making the predictions", y = "Proportion of calls") 
dev.off()

########################################################
# Supp Fig 19: Call count on all 503 samples by caller 
########################################################
call_count_summary_503_samples <-  sqldf("select CALLER, TYPE, count(*) as COUNT from all_calls_before_bpres_df
                                         group by CALLER, TYPE")

pdf("supp_fig_19_503samples_original_call_count_summary.pdf", width = 6, height = 4)
ggplot(data=call_count_summary_503_samples, aes(x=CALLER, y=COUNT, fill=TYPE)) + theme_classic() +
  geom_bar(position = "dodge", stat="identity") + 
  #  facet_grid(~TYPE, scale = "free", switch = "x") + 
  geom_text(aes(label=COUNT), position = position_dodge(width = 1), vjust = -.25, size = 2.5, fontface = "bold") +
  ggtitle ("Distribution of the calls made in 503 samples samples, by caller") +
  theme(plot.title = element_text(lineheight=.8, size=12, face="bold", hjust = 0.5), 
        strip.text = element_text(size = 10, face = "bold"),
        axis.title = element_text(size = 10, face="bold"),
        axis.text = element_text(size = 8, face="bold"), 
        #axis.text.x = element_blank(),
        legend.title = element_text(size=8, face="bold"),
        legend.text = element_text(size=8, face="bold"),
        legend.position = "bottom") +
  labs(x = "", y = "Count", fill = "") 
dev.off()


########################
# SUPPLEMENTARY TABLES #
########################

#########################
# Supplementary Table 2 #
#########################
val_overlap_stats_strat2 <- sqldf("select V2 as True_Label,V4 as False_Label, round(V2/((V2 + V4) * 1.0) * 100,1) as Label_Prop 
                                  from val_overlap_stats_strat2_df")
val_overlap_stats_strat2$Proportion <- c(2,3,5,10,20,30,40,50,60,70,80,90) 
val_overlap_stats_strat2 <- val_overlap_stats_strat2[c(4,1,2,3)]
write.csv(val_overlap_stats_strat2, file="supp_table_1_label_proportion.csv", row.names = FALSE)

##########################
#  Supplementary Table 3 #
##########################
# Overall Precision (without sample CNV frequency and CNV size between 5000 and 5000000)
sample_list_df_1000g <- sqldf("select sample from sample_list_df where TRAINING_PROP = 0.7")

all_1000g_calls <- rbind(tp_fp_1000g_df, tn_fn_1000g_df)
all_1000g_calls <- sqldf("select CANOES_PROP, CODEX_PROP, CLAMMS_PROP, XHMM_PROP, CLASS_LABEL, VAL_LABEL 
                         from all_1000g_calls where SAMPLE in (select SAMPLE from sample_list_df_1000g) 
                         and NUM_SPLITS = 1 and TRAINING_PROP = 0.7")
dim(all_1000g_calls)
all_1000g_calls$UNIQ_ID <- seq(1,dim(all_1000g_calls)[1])

all_1000g_calls$CANOES_IND <- ifelse(all_1000g_calls$CANOES_PROP > 0, 1, 0)
all_1000g_calls$CODEX_IND <- ifelse(all_1000g_calls$CODEX_PROP > 0, 1, 0)
all_1000g_calls$CLAMMS_IND <- ifelse(all_1000g_calls$CLAMMS_PROP > 0, 1, 0)
all_1000g_calls$XHMM_IND <- ifelse(all_1000g_calls$XHMM_PROP > 0, 1, 0)
all_1000g_calls <- all_1000g_calls[,-c(1,2,3,4)]
head(all_1000g_calls)
dim(all_1000g_calls)

all_1000g_calls$CANOES_CODEX <- ifelse((all_1000g_calls$CANOES_IND + all_1000g_calls$CODEX_IND) >= 2, 1, 0)
all_1000g_calls$CANOES_CLAMMS <- ifelse((all_1000g_calls$CANOES_IND + all_1000g_calls$CLAMMS_IND) >= 2, 1, 0)
all_1000g_calls$CANOES_XHMM <- ifelse((all_1000g_calls$CANOES_IND + all_1000g_calls$XHMM_IND) >= 2, 1, 0)
all_1000g_calls$CODEX_CLAMMS <- ifelse((all_1000g_calls$CODEX_IND + all_1000g_calls$CLAMMS_IND) >= 2, 1, 0)
all_1000g_calls$CODEX_XHMM <- ifelse((all_1000g_calls$CODEX_IND + all_1000g_calls$XHMM_IND) >= 2, 1, 0)
all_1000g_calls$CLAMMS_XHMM <- ifelse((all_1000g_calls$CLAMMS_IND + all_1000g_calls$XHMM_IND) >= 2, 1, 0)
all_1000g_calls$TWO_CALLERS <- ifelse((all_1000g_calls$CANOES_IND + all_1000g_calls$CODEX_IND + all_1000g_calls$CLAMMS_IND + all_1000g_calls$XHMM_IND) == 2, 1, 0)
all_1000g_calls$TWO_OR_MORE_CALLERS <- ifelse((all_1000g_calls$CANOES_IND + all_1000g_calls$CODEX_IND + all_1000g_calls$CLAMMS_IND + all_1000g_calls$XHMM_IND) >= 2, 1, 0)
all_1000g_calls$CANOES_CODEX_CLAMMS <- ifelse((all_1000g_calls$CANOES_IND + all_1000g_calls$CODEX_IND + all_1000g_calls$CLAMMS_IND) >= 3, 1, 0)
all_1000g_calls$CANOES_CODEX_XHMM <- ifelse((all_1000g_calls$CANOES_IND + all_1000g_calls$CODEX_IND + all_1000g_calls$XHMM_IND) >= 3, 1, 0)
all_1000g_calls$CODEX_CLAMMS_XHMM <- ifelse((all_1000g_calls$CODEX_IND + all_1000g_calls$CLAMMS_IND + all_1000g_calls$XHMM_IND) >= 3, 1, 0)
all_1000g_calls$CANOES_CLAMMS_XHMM <- ifelse((all_1000g_calls$CANOES_IND + all_1000g_calls$CLAMMS_IND + all_1000g_calls$XHMM_IND) >= 3, 1, 0)
all_1000g_calls$THREE_CALLERS <- ifelse((all_1000g_calls$CANOES_IND + all_1000g_calls$CODEX_IND + all_1000g_calls$CLAMMS_IND + all_1000g_calls$XHMM_IND) == 3, 1, 0)
all_1000g_calls$THREE_OR_MORE_CALLERS <- ifelse((all_1000g_calls$CANOES_IND + all_1000g_calls$CODEX_IND + all_1000g_calls$CLAMMS_IND + all_1000g_calls$XHMM_IND) >= 3, 1, 0)
all_1000g_calls$CANOES_CODEX_CLAMMS_XHMM <- ifelse((all_1000g_calls$CANOES_IND + all_1000g_calls$CODEX_IND + all_1000g_calls$CLAMMS_IND + all_1000g_calls$XHMM_IND) >= 4, 1, 0)

head(all_1000g_calls)

canoes_df <- sqldf("select VAL_LABEL, CANOES_IND, count(*) as COUNT, 'CANOES' as CALLER from all_1000g_calls group by VAL_LABEL, CANOES_IND")
codex_df <- sqldf("select VAL_LABEL, CODEX_IND, count(*) as COUNT, 'CODEX' as CALLER from all_1000g_calls group by VAL_LABEL, CODEX_IND")
clamms_df <- sqldf("select VAL_LABEL, CLAMMS_IND, count(*) as COUNT, 'CLAMMS' as CALLER from all_1000g_calls group by VAL_LABEL, CLAMMS_IND")
xhmm_df <- sqldf("select VAL_LABEL, XHMM_IND, count(*) as COUNT, 'XHMM' as CALLER from all_1000g_calls group by VAL_LABEL, XHMM_IND")
canoes_codex_df <- sqldf("select VAL_LABEL, CANOES_CODEX, count(*) as COUNT, 'CANOES + CODEX' as CALLER from all_1000g_calls group by VAL_LABEL, CANOES_CODEX")
canoes_clamms_df <- sqldf("select VAL_LABEL, CANOES_CLAMMS, count(*) as COUNT, 'CANOES + CLAMMS' as CALLER from all_1000g_calls group by VAL_LABEL, CANOES_CLAMMS")
canoes_xhmm_df <- sqldf("select VAL_LABEL, CANOES_XHMM, count(*) as COUNT, 'CANOES + XHMM' as CALLER from all_1000g_calls group by VAL_LABEL, CANOES_XHMM")
codex_clamms_df <- sqldf("select VAL_LABEL, CODEX_CLAMMS, count(*) as COUNT, 'CODEX + CLAMMS' as CALLER from all_1000g_calls group by VAL_LABEL, CODEX_CLAMMS")
codex_xhmm_df <- sqldf("select VAL_LABEL, CODEX_XHMM, count(*) as COUNT, 'CODEX + XHMM' as CALLER from all_1000g_calls group by VAL_LABEL, CODEX_XHMM")
clamms_xhmm_df <- sqldf("select VAL_LABEL, CLAMMS_XHMM, count(*) as COUNT, 'CLAMMS + XHMM' as CALLER from all_1000g_calls group by VAL_LABEL, CLAMMS_XHMM")
two_callers_df <- sqldf("select VAL_LABEL, TWO_CALLERS, count(*) as COUNT, 'Two Callers' as CALLER from all_1000g_calls group by VAL_LABEL, TWO_CALLERS")
two_or_more_callers_df <- sqldf("select VAL_LABEL, TWO_OR_MORE_CALLERS, count(*) as COUNT, 'Two or more callers' as CALLER from all_1000g_calls group by VAL_LABEL, TWO_OR_MORE_CALLERS")

canoes_codex_clamms_df <- sqldf("select VAL_LABEL, CANOES_CODEX_CLAMMS, count(*) as COUNT, 'CANOES + CODEX + CLAMMS' as CALLER from all_1000g_calls group by VAL_LABEL, CANOES_CODEX_CLAMMS")
canoes_codex_xhmm_df <- sqldf("select VAL_LABEL, CANOES_CODEX_XHMM, count(*) as COUNT, 'CANOES + CODEX + XHMM' as CALLER from all_1000g_calls group by VAL_LABEL, CANOES_CODEX_XHMM")
codex_clamms_xhmm_df <- sqldf("select VAL_LABEL, CODEX_CLAMMS_XHMM, count(*) as COUNT, 'CODEX + CLAMMS + XHMM' as CALLER from all_1000g_calls group by VAL_LABEL, CODEX_CLAMMS_XHMM")
canoes_clamms_xhmm_df <- sqldf("select VAL_LABEL, CANOES_CLAMMS_XHMM, count(*) as COUNT, 'CANOES + CLAMMS + XHMM' as CALLER from all_1000g_calls group by VAL_LABEL, CANOES_CLAMMS_XHMM")

three_callers_df <- sqldf("select VAL_LABEL, THREE_CALLERS, count(*) as COUNT, 'Three Callers' as CALLER from all_1000g_calls group by VAL_LABEL, THREE_CALLERS")
three_or_more_callers_df <- sqldf("select VAL_LABEL, THREE_CALLERS, count(*) as COUNT, 'Three or more callers' as CALLER from all_1000g_calls group by VAL_LABEL, THREE_CALLERS")


all_df <- sqldf("select VAL_LABEL, CANOES_CODEX_CLAMMS_XHMM, count(*) as COUNT, 'ALL' as CALLER from all_1000g_calls group by VAL_LABEL, CANOES_CODEX_CLAMMS_XHMM")

colnames(canoes_df) <- c("VAL_LABEL", "PRED_IND", "COUNT", "CALLER")
colnames(codex_df) <- c("VAL_LABEL", "PRED_IND", "COUNT", "CALLER")
colnames(clamms_df) <- c("VAL_LABEL", "PRED_IND", "COUNT", "CALLER")
colnames(xhmm_df) <- c("VAL_LABEL", "PRED_IND", "COUNT", "CALLER")
colnames(canoes_codex_df) <- c("VAL_LABEL", "PRED_IND", "COUNT", "CALLER")
colnames(canoes_clamms_df) <- c("VAL_LABEL", "PRED_IND", "COUNT", "CALLER")
colnames(canoes_xhmm_df) <- c("VAL_LABEL", "PRED_IND", "COUNT", "CALLER")
colnames(codex_clamms_df) <- c("VAL_LABEL", "PRED_IND", "COUNT", "CALLER")
colnames(codex_xhmm_df) <- c("VAL_LABEL", "PRED_IND", "COUNT", "CALLER")
colnames(clamms_xhmm_df) <- c("VAL_LABEL", "PRED_IND", "COUNT", "CALLER")
colnames(two_callers_df) <- c("VAL_LABEL", "PRED_IND", "COUNT", "CALLER")
colnames(two_or_more_callers_df) <- c("VAL_LABEL", "PRED_IND", "COUNT", "CALLER")
colnames(canoes_codex_clamms_df) <- c("VAL_LABEL", "PRED_IND", "COUNT", "CALLER")
colnames(canoes_codex_xhmm_df) <- c("VAL_LABEL", "PRED_IND", "COUNT", "CALLER")
colnames(codex_clamms_xhmm_df) <- c("VAL_LABEL", "PRED_IND", "COUNT", "CALLER")
colnames(canoes_clamms_xhmm_df) <- c("VAL_LABEL", "PRED_IND", "COUNT", "CALLER")
colnames(three_callers_df) <- c("VAL_LABEL", "PRED_IND", "COUNT", "CALLER")
colnames(three_or_more_callers_df) <- c("VAL_LABEL", "PRED_IND", "COUNT", "CALLER")
colnames(all_df) <- c("VAL_LABEL", "PRED_IND", "COUNT", "CALLER")

summary_df <- rbind(canoes_df,codex_df,clamms_df,xhmm_df,canoes_codex_df,canoes_clamms_df,
                    canoes_xhmm_df,codex_clamms_df,codex_xhmm_df,clamms_xhmm_df,two_callers_df, two_or_more_callers_df,
                    canoes_codex_clamms_df,canoes_codex_xhmm_df, codex_clamms_xhmm_df,
                    canoes_clamms_xhmm_df,three_callers_df,three_or_more_callers_df,all_df)
summary_df <- sqldf("select * from summary_df where PRED_IND = 1")

head(summary_df)

caller_summary_df <- sqldf("select CALLER, sum(COUNT) as COUNT from summary_df group by CALLER")

prec_recall_1000g_callers <- sqldf("select a.CALLER, b.count as CONCORDANT_CNVs, a.COUNT as TRUE_PREDICTIONS, 
                                   round(a.COUNT/(b.COUNT * 1.0) * 100,2) as PRECISION, round(a.count/(2270 * 1.0) * 100,2) as RECALL
                                   from summary_df a, caller_summary_df b 
                                   where a.CALLER = b.CALLER and a.VAL_LABEL = 'TRUE_CNV'")

# CN-Learn
prec_recall_1000g_cnlearn <- sqldf("select 'CN-Learn' as CALLER, b.count as CONCORDANT_CNVs, count(*) as TRUE_PREDICTIONS, 
                                   round(count(*)/(b.count * 1.0) * 100,2) as PRECISION, round(count(*)/(2270 * 1.0) * 100,2) as RECALL
                                   from tp_fp_1000g_df a, (select count(*) as COUNT from tp_fp_1000g_df where TRAINING_PROP = 0.7 and STRATEGY = 2 AND NUM_SPLITS = 1) b
                                   where a.TRAINING_PROP = 0.7 and a.STRATEGY = 2 AND a.NUM_SPLITS = 1 and a.VAL_LABEL = 'TRUE_CNV'
                                   group by VAL_LABEL")

prec_recall_1000g <- rbind(prec_recall_1000g_callers, prec_recall_1000g_cnlearn)
getwd()
write.csv(prec_recall_1000g, "prec_recall_caller_combo.csv")

#############################
# Supplementary Table 5 & 6 #
#############################
all_tp_fp_calls_ma <- tp_fp_df[which(tp_fp_df$STRATEGY == 2 & tp_fp_df$CLASSIFIER == 'RF' & tp_fp_df$NUM_SPLITS == 1 & 
                                       tp_fp_df$TRAINING_PROP == 0.1),c(1,2,3,4,5,6,7,8,9,10,19,26)]

all_tp_fp_calls_clamms <- tp_fp_df[which(tp_fp_df$STRATEGY == 3 & tp_fp_df$CLASSIFIER == 'RF' & tp_fp_df$NUM_SPLITS == 1 & 
                                           tp_fp_df$TRAINING_PROP == 0.1),c(1,2,3,4,5,6,7,9,10,19,26)]


all_tp_fp_calls_ma$CANOES <- ifelse(all_tp_fp_calls_ma$CANOES_PROP > 0, 'YES', 'NO')
all_tp_fp_calls_ma$CODEX <- ifelse(all_tp_fp_calls_ma$CODEX_PROP > 0, 'YES', 'NO')
all_tp_fp_calls_ma$CLAMMS <- ifelse(all_tp_fp_calls_ma$CLAMMS_PROP > 0, 'YES', 'NO')
all_tp_fp_calls_ma$XHMM <- ifelse(all_tp_fp_calls_ma$XHMM_PROP > 0, 'YES', 'NO')

all_tp_fp_calls_ma$CLASSIFICATION <- ifelse(all_tp_fp_calls_ma$PRED_PROBS >= 0.5, 'True', 'False')
all_tp_fp_calls_ma <- all_tp_fp_calls_ma[,-c(6,7,8,9)]

all_tp_fp_calls_clamms$CANOES <- ifelse(all_tp_fp_calls_clamms$CANOES_PROP > 0, 'YES', 'NO')
all_tp_fp_calls_clamms$CODEX <- ifelse(all_tp_fp_calls_clamms$CODEX_PROP > 0, 'YES', 'NO')
all_tp_fp_calls_clamms$XHMM <- ifelse(all_tp_fp_calls_clamms$XHMM_PROP > 0, 'YES', 'NO')

all_tp_fp_calls_clamms$CLASSIFICATION <- ifelse(all_tp_fp_calls_clamms$PRED_PROBS > 0.5, 'True', 'False')
all_tp_fp_calls_clamms <- all_tp_fp_calls_clamms[,-c(6,7,8)]

all_tp_fp_calls_ma <- all_tp_fp_calls_ma[order(-all_tp_fp_calls_ma$PRED_PROBS, all_tp_fp_calls_ma$NUM_OVERLAPS),]
all_tp_fp_calls_clamms <- all_tp_fp_calls_clamms[order(-all_tp_fp_calls_clamms$PRED_PROBS, all_tp_fp_calls_clamms$NUM_OVERLAPS),]

dim(all_tp_fp_calls_ma)
dim(all_tp_fp_calls_clamms)

write.csv(all_tp_fp_calls_ma,file="tp_fp_calls_w_caller_info_ma.csv", row.names = FALSE)
write.csv(all_tp_fp_calls_clamms, file="tp_fp_calls_w_caller_info_clamms.csv", row.names = FALSE)

#############################
# Supplementary Table 5 & 6 #
#############################
all_tn_fn_calls_ma <- tn_fn_df[which(tn_fn_df$STRATEGY == 2 & tn_fn_df$CLASSIFIER == 'RF' & tn_fn_df$NUM_SPLITS == 1 & 
                                       tn_fn_df$TRAINING_PROP == 0.1),c(1,2,3,4,5,6,7,8,9,10,19,26)]
all_tn_fn_calls_clamms <- tn_fn_df[which(tn_fn_df$STRATEGY == 3 & tn_fn_df$CLASSIFIER == 'RF' & tn_fn_df$NUM_SPLITS == 1 & 
                                           tn_fn_df$TRAINING_PROP == 0.1),c(1,2,3,4,5,6,7,8,9,10,19,26)]

all_tn_fn_calls_ma$CANOES <- ifelse(all_tn_fn_calls_ma$CANOES_PROP > 0, 'YES', 'NO')
all_tn_fn_calls_ma$CODEX <- ifelse(all_tn_fn_calls_ma$CODEX_PROP > 0, 'YES', 'NO')
all_tn_fn_calls_ma$CLAMMS <- ifelse(all_tn_fn_calls_ma$CLAMMS_PROP > 0, 'YES', 'NO')
all_tn_fn_calls_ma$XHMM <- ifelse(all_tn_fn_calls_ma$XHMM_PROP > 0, 'YES', 'NO')

all_tn_fn_calls_ma$CLASSIFICATION <- ifelse(all_tn_fn_calls_ma$PRED_PROBS > 0.5, 'True', 'False')
all_tn_fn_calls_ma <- all_tn_fn_calls_ma[,-c(6,7,8,9)]

all_tn_fn_calls_clamms$CANOES <- ifelse(all_tn_fn_calls_clamms$CANOES_PROP > 0, 'YES', 'NO')
all_tn_fn_calls_clamms$CODEX <- ifelse(all_tn_fn_calls_clamms$CODEX_PROP > 0, 'YES', 'NO')
all_tn_fn_calls_clamms$XHMM <- ifelse(all_tn_fn_calls_clamms$XHMM_PROP > 0, 'YES', 'NO')

all_tn_fn_calls_clamms$CLASSIFICATION <- ifelse(all_tn_fn_calls_clamms$PRED_PROBS > 0.5, 'True', 'False')
all_tn_fn_calls_clamms <- all_tn_fn_calls_clamms[,-c(6,7,8)]

all_tn_fn_calls_ma <- all_tn_fn_calls_ma[order(-all_tn_fn_calls_ma$PRED_PROBS, all_tn_fn_calls_ma$NUM_OVERLAPS),]
all_tn_fn_calls_clamms <- all_tn_fn_calls_clamms[order(-all_tn_fn_calls_clamms$PRED_PROBS, all_tn_fn_calls_clamms$NUM_OVERLAPS),]

getwd()
write.csv(all_tn_fn_calls_ma,file="tn_fn_calls_w_caller_info_ma.csv", row.names = FALSE)
write.csv(all_tn_fn_calls_clamms, file="tn_fn_calls_w_caller_info_clamms.csv", row.names = FALSE)

dim(all_tn_fn_calls_ma)
print(all_tn_fn_calls_ma)
dim(all_tp_fp_calls_clamms)


#######################################
# Numbers for the 'Results' section   #
#######################################
#Venn Diagram - Microarray:
all_test_calls_strat2_df
all_true_test_calls_strat2_df
tp_fp_rf_venn_df
tp_strat2_rf_df
head(all_test_calls_strat2_df)
# Number of CNV calls in the test set
sqldf("select count(*) from all_test_calls_strat2_df")

#Proportion of true CNVs in the test set
sqldf("select count(*)/((select count(*) from all_test_calls_strat2_df) * 1.0) as TRUE_PROP from all_test_calls_strat2_df where LABEL = 'TRUE_CNV'")

sqldf("select count(*) from tp_fp_rf_venn_df")
sqldf("select count(*) from tp_strat2_rf_df")

# Microarray : Number of calls by concordance, prior to breakpoint resolution
sqldf("select NUM_OVERLAPS, count(*) from all_calls_before_bpres_df group by NUM_OVERLAPS")

# Microarray : Number of calls by concordance, after breakpoint resolution
sqldf("select count(*) from all_calls_after_bpres_df")
sqldf("select NUM_OVERLAPS, count(*) from all_calls_after_bpres_df group by NUM_OVERLAPS")

# Microarray : Number of calls available to the CN-Learn classifier
sqldf("select count(*) from all_calls_after_bpres_df where (PRED_END - PRED_START) > 50000 and (PRED_END - PRED_START)  < 5000000")
sqldf("select * from all_calls_after_bpres_df where (PRED_END - PRED_START) > 50000 and (PRED_END - PRED_START)  < 5000000")


# CLAMMS : Number of calls available to the CN-Learn classifier
sqldf("select count(*) from strat3_calls_after_bpres_df where (PRED_END - PRED_START) > 5000 and (PRED_END - PRED_START)  < 5000000")
# CLAMMS : Number of calls in the test set
sqldf("select count(*) from strat3_calls_after_bpres_df where SAMPLE in (select * from clamms_sample_list_df)
      and (PRED_END - PRED_START) > 5000 and (PRED_END - PRED_START)  < 5000000")


##########################
# MISCELLANEOUS ANALYSIS #
##########################
###############################################
# Calculate the median distance between three #
# exome capture probes                        #
###############################################
number_of_probes <- dim(target_list)[1]

probe_distance <- integer(number_of_probes)
for (i in 1:number_of_probes){
  if (i+2 <= number_of_probes & target_list[(i + 2),1] == target_list[i,1] ){
    probe_distance[i] <- target_list[(i + 2),3] - target_list[i,2]
  }
}
probe_distance <- probe_distance[probe_distance!=0]
summary(probe_distance)

####################################################
# Calculate the median distance between SNP probes #
####################################################
omni1_snps <- read.table("omni1_snps.bed")
omni1_snps <- sqldf("select * from omni1_snps order by 1,2")
num_of_snp_probes <- dim(omni1_snps)[1]

snp_probe_distance <- integer(num_of_snp_probes)
for (i in 1:num_of_snp_probes){
  if (i+1 <= num_of_snp_probes & omni1_snps[(i + 1),1] == omni1_snps[i,1] ){
    snp_probe_distance[i] <- omni1_snps[(i + 1),3] - omni1_snps[i,3]
  }
}
summary(snp_probe_distance)

############################################################
# Preparatory analysis for identifying samples to illustrate
############################################################
calls_w_3_overlaps <- sqldf("select distinct a.CALLER, a.SAMPLE, a.CHR, a.PRED_START, a.PRED_END 
                            from retained_calls_df a,  strat2_df b
                            where a.NUM_OVERLAPS = 3 and a.LABEL = 'TRUE_CNV'
                            and a.SAMPLE = b.SAMPLE 
                            order by a.SAMPLE, a.CHR, a.PRED_START, a.PRED_END, a.CALLER")
write.csv(calls_w_3_overlaps, file = "calls_w_3_overlaps.csv", row.names = FALSE)


# Query to find calls that are relatively smaller in size using the second merge strategy 
sqldf("select a.CHR, a.SAMPLE, a.PRED_START, a.PRED_END, b.PRED_START, b.PRED_END, round((b.PRED_END - b.PRED_START)/((a.PRED_END - a.PRED_START) * 1.0),2) as COORD_PROP
      from strat1_df a, strat2_df b
      where a.SAMPLE = b.SAMPLE and a.TYPE = b.TYPE and a.CHR = b.CHR
      and a.NUM_OVERLAPS = 3 and a.LABEL = 'TRUE_CNV' and b.LABEL = 'TRUE_CNV'
      and b.PRED_START > a.PRED_START and b.PRED_START < a.PRED_END
      and COORD_PROP < 0.6
      order by a.SAMPLE, a.CHR, a.PRED_START")

sqldf("select * from strat2_df where sample = '14825.x1' and chr = 2")
sqldf("select * from retained_calls_df where sample = '14825.x1' and chr = 2")

sqldf("select * from strat2_df where sample = '14725.x51' and chr = 7")
sqldf("select * from retained_calls_df where sample = '14725.x51' and chr = 7")


###################################################################
# Supp Fig 2B Equivalent :Correlation between predictors (CLAMMS) #
###################################################################
corr_predictors_3_df <- strat3_df[,c("CANOES_PROP", "CODEX_PROP", "XHMM_PROP", "NUM_OVERLAPS", "PRED_SIZE", "NUM_TARGETS", "GC", "MAP", "RD_PROP")]
colnames(corr_predictors_3_df) <- c("CANOES", "CODEX", "XHMM", "Concordance", "Pred Size", "Target Count", "GC Content", "Mappability", "Read Depth Ratio")
corr_results <- cor(corr_predictors_3_df)

options(scipen=999)
result_3_spearman <-rcorr(as.matrix(corr_predictors_3_df), type = "spearman")
flattenCorrMatrix(round(result_3_spearman$r,3), round(result_3_spearman$P,3))

correlations_3 <- round(as.data.frame(result_3_spearman$r),2)
write.csv(correlations_3, file="supp_table_3_pred_correlations_clamms.csv", row.names = FALSE)

#Build correlation plot
pdf("supp_fig_2b_eq_predictor_correlations_clamms.pdf", width = 8, height = 5)
corrplot(result_3_spearman$r, type="upper", p.mat = result_3_spearman$P, sig.level = 0.01, insig = "blank", tl.col = "black", tl.srt = 45)
dev.off()


#######################################################################
# Supp Figure 2C Equivalent :Prediction probability analysis - CLAMMS #
#######################################################################
tp_fp_tn_fn_clamms_df <- tp_fp_tn_fn_df
tp_fp_tn_fn_clamms_df$CANOES <- ifelse(tp_fp_tn_fn_clamms_df$CANOES_PROP > 0, 1, 0)
tp_fp_tn_fn_clamms_df$CODEX<- ifelse(tp_fp_tn_fn_clamms_df$CODEX_PROP > 0, 1, 0)
tp_fp_tn_fn_clamms_df$XHMM <- ifelse(tp_fp_tn_fn_clamms_df$XHMM_PROP > 0, 1, 0)
tp_fp_tn_fn_clamms_df$OV_CODE <- paste(tp_fp_tn_fn_clamms_df$CANOES, tp_fp_tn_fn_clamms_df$CODEX, tp_fp_tn_fn_clamms_df$XHMM, sep = '') 

tp_fp_tn_fn_probs_size_clamms_df <- sqldf("select TYPE, PRED_SIZE, NUM_OVERLAPS, PRED_PROBS, CLASS, VAL_LABEL
                                          from tp_fp_tn_fn_clamms_df 
                                          where CLASSIFIER = 'RF' and STRATEGY = '3' and NUM_SPLITS = 1 and TRAINING_PROP = 0.7
                                          and PRED_PROBS > 0")

#Generate plot by Validation Label (Distribution Plot)
pdf("supp_fig_2c_eq_predicted_probability_distribution_clamms_0.7.pdf", width = 8, height = 5)
ggplot(data=tp_fp_tn_fn_probs_size_clamms_df, aes(x=PRED_PROBS, fill = VAL_LABEL)) + theme_classic() + 
  geom_histogram(position = "dodge", aes(y=..count../sum(..count..)), binwidth = 0.05) +
  #  facet_grid(~TYPE, scale = "free") + 
  ggtitle ("Distribution of predicted probability scores (Excludes zero)") +
  scale_fill_manual(values = c("True CNV" = "blue","False CNV" = "red")) +
  theme(plot.title = element_text(lineheight=.8, size=9, face="bold", hjust = 0.5), 
        strip.text = element_text(size = 9, face = "bold"),
        axis.title = element_text(size = 9, face="bold"),
        axis.text = element_text(size = 9, face="bold"), 
        legend.title = element_text(size=9, face="bold"),
        legend.text = element_text(size=9, face="bold"),
        legend.position = "bottom") +
  geom_vline(xintercept = 0.5, linetype="dotted") + 
  labs(x = "Predicted Probabilities", y = "Frequency", fill = "") 
dev.off()


#########################################################################
# Supp Figure 3B Equivalent: Prediction probability analysis - (CLAMMS) #
#########################################################################
tp_summary_clamms_df <- sqldf("select STRATEGY,  SPLIT, TRAINING_PROP, CLASS, count(*) as COUNT
                              from tp_df
                              where NUM_SPLITS = 10 and STRATEGY = 3 and CLASSIFIER = 'RF'
                              group by STRATEGY,  SPLIT, TRAINING_PROP, CLASS")
fp_summary_clamms_df <- sqldf("select STRATEGY, SPLIT, TRAINING_PROP, CLASS, count(*) as COUNT
                              from fp_df
                              where NUM_SPLITS = 10 and STRATEGY = 3 and CLASSIFIER = 'RF'
                              group by STRATEGY, SPLIT, TRAINING_PROP, CLASS")
fn_summary_clamms_df <- sqldf("select STRATEGY, SPLIT, TRAINING_PROP, CLASS, count(*) as COUNT
                              from fn_df
                              where NUM_SPLITS = 10 and STRATEGY = 3 and CLASSIFIER = 'RF'
                              group by STRATEGY, SPLIT, TRAINING_PROP, CLASS")
tn_summary_clamms_df <- sqldf("select STRATEGY,  SPLIT, TRAINING_PROP, CLASS, count(*) as COUNT
                              from tn_df
                              where NUM_SPLITS = 10 and STRATEGY = 3 and CLASSIFIER = 'RF'
                              group by STRATEGY, SPLIT, TRAINING_PROP, CLASS")

all_prec_recall_clamms_df <- sqldf("select tp.TRAINING_PROP, tp.SPLIT, round(tp.COUNT/((tp.COUNT + fp.COUNT) * 1.0),2) as Precision,
                                   round(tp.COUNT/((tp.COUNT + fn.COUNT) * 1.0),2) as Recall
                                   from tp_summary_clamms_df tp, fp_summary_clamms_df fp, fn_summary_clamms_df fn
                                   where tp.SPLIT = fp.SPLIT and tp.TRAINING_PROP = fp.TRAINING_PROP and
                                   tp.SPLIT = fn.SPLIT and tp.TRAINING_PROP = fn.TRAINING_PROP
                                   order by tp.TRAINING_PROP")

write.csv(all_prec_recall_clamms_df, file="supp_table_4_precision_dist_clamms.csv", row.names = FALSE)
melted_all_prec_recall_clamms_df <- melt(all_prec_recall_clamms_df, id=c("TRAINING_PROP", "SPLIT"))

means_clamms <- aggregate(value ~  TRAINING_PROP + variable, melted_all_prec_recall_clamms_df, mean)
means_clamms$value <- round(means_clamms$value,2)

pdf(file="supp_fig_3b_eq_precision_distribution_clamms.pdf", width = 5, height = 5)
ggplot(data=melted_all_prec_recall_clamms_df, aes(x=as.character(TRAINING_PROP), y=value, fill = variable)) + theme_classic() +
  geom_boxplot(position=position_dodge(1)) + ylim(0.5,1.0) + 
  geom_point(position=position_jitterdodge(dodge.width=0.9)) +
  geom_text(data = means_clamms, aes(label = value, y = value + 0.03)) +
  #  geom_text(aes(label=label_val), position = position_dodge(width = 0.9), vjust = -.25, size = 2, fontface = "bold") +
  ggtitle ("Distribution of performance obtained during 10-fold cross-validation") + 
  theme(plot.title = element_text(lineheight=.8, size=9, face="bold", hjust = 0.5), 
        strip.text = element_text(size = 8, face = "bold"),
        axis.title = element_text(size = 8, face="bold"),
        axis.text = element_text(size = 8, face="bold"), 
        #        axis.text.x = element_blank(),
        legend.title = element_text(size=8, face="bold"),
        legend.text = element_text(size=8, face="bold"),
        legend.position = "bottom") +
  labs(x = "", y = "Precision-Recall", fill = "") 
dev.off()


######################################################################################################################
# Supp Figure 3C Equivalent : Characterization of callers based on precision rates at different size ranges (CLAMMS) #
######################################################################################################################
clamms_precision_df <- all_calls_before_bpres_df
clamms_precision_df$CLAMMS_VAL_LABEL <- ifelse(clamms_precision_df$CLAMMS_PROP > 0.1, 1, 0)
clamms_precision_df$Size <- ''
clamms_precision_df$Size <- ifelse(clamms_precision_df$PRED_SIZE >= 5000 & clamms_precision_df$PRED_SIZE < 10000, '5-10kb', clamms_precision_df$Size)
clamms_precision_df$Size <- ifelse(clamms_precision_df$PRED_SIZE >= 10000 & clamms_precision_df$PRED_SIZE < 25000, '10-25kb', clamms_precision_df$Size)
clamms_precision_df$Size <- ifelse(clamms_precision_df$PRED_SIZE >= 25000 & clamms_precision_df$PRED_SIZE < 50000, '25-50kb', clamms_precision_df$Size)
clamms_precision_df$Size <- ifelse(clamms_precision_df$PRED_SIZE >= 50000 & clamms_precision_df$PRED_SIZE < 100000, '50-100kb', clamms_precision_df$Size)
clamms_precision_df$Size <- ifelse(clamms_precision_df$PRED_SIZE >= 100000 & clamms_precision_df$PRED_SIZE < 250000, '100-250kb', clamms_precision_df$Size)
clamms_precision_df$Size <- ifelse(clamms_precision_df$PRED_SIZE >= 250000 & clamms_precision_df$PRED_SIZE < 500000, '250-500kb', clamms_precision_df$Size)
clamms_precision_df$Size <- ifelse(clamms_precision_df$PRED_SIZE >= 500000 & clamms_precision_df$PRED_SIZE < 5000000, '500kb-5mb', clamms_precision_df$Size)


caller_precision_clamms <- sqldf("select A.CALLER as Caller, round(A.COUNT/(B.COUNT * 1.0),2) as Precision, A.SIZE as Size from
                                 (select CALLER, SIZE, CLAMMS_VAL_LABEL, count(*) as COUNT from clamms_precision_df where CALLER <> 'CLAMMS'
                                 and CLAMMS_VAL_LABEL = 1
                                 GROUP BY CALLER, SIZE, CLAMMS_VAL_LABEL) as A, 
                                 (select CALLER, SIZE, count(*) as COUNT from clamms_precision_df where CALLER <> 'CLAMMS'
                                 GROUP BY CALLER, SIZE) as B
                                 where A.CALLER = B.CALLER and A.SIZE = B.SIZE and A.Size <> ''")

all_precisions_by_size_clamms <- rbind(caller_precision_clamms, rf_metrics_size_modified_df[5:11,])
all_precisions_by_size_clamms$Caller <- ordered(all_precisions_by_size_clamms$Caller, 
                                                levels = c("CN_Learn (w/ CLAMMS Gold Standard)", "CANOES", "CODEX", "XHMM")) 
all_precisions_by_size_clamms$Size <- ordered(all_precisions_by_size_clamms$Size, levels = c("5-10kb", "10-25kb", "25-50kb", "50-100kb", "100-250kb",
                                                                                             "250-500kb", "500kb-5mb"))

pdf("supp_fig_3c_eq_precision_values_by_sizerange_clamms_0.7.pdf", width = 8, height = 5)
ggplot(all_precisions_by_size_clamms, aes(x = Size, y = Precision, group = Caller, color = Caller)) +
  ylim(0.0, 1.00) + geom_point(size=2.5, shape=1, stroke = 1.5) + geom_line(size=1) + theme_classic() +
  geom_text(aes(label=Precision), vjust = -0.5, size = 2.5, hjust = 1.25, fontface = "bold") +
  theme(plot.title = element_text(lineheight=.8, size=9, face="bold", hjust = 0.5), 
        strip.text = element_text(size = 9, face = "bold"),
        axis.title = element_text(size = 9, face="bold"),
        axis.text = element_text(size = 9, face="bold"), 
        legend.title = element_text(size=9, face="bold"),
        legend.text = element_text(size=9, face="bold"),
        legend.position = "bottom") +
  labs(x = "Size Range", y = "Precision", color = "") 
dev.off()


#######################################################################################
# Supp Figure 3D Equivalent: Feature Importances for the main merge strategy (CLAMMS) #
#######################################################################################
feat_imp_strat3 <- sqldf ("select Training_Proportion, Chromosome, CNV_Type, Canoes_Proportion, Codex_Proportion, Clamms_Proportion, XHMM_Proportion, Overlap_Count, 
                          RD_Proportion, GC_Content, Mappability, Number_Of_Targets, Size_Label, 'Genomic' as Predictor_Group 
                          from rf_metrics_df 
                          where Classifier = 'RF' and Strategy = 3 and Num_Splits = 10 and Training_Proportion = 0.7")
melted_feat_imp_strat3 <- melt(feat_imp_strat3, id=c("Training_Proportion", "Predictor_Group"))
melted_feat_imp_strat3$prop <- melted_feat_imp_strat3$value *100
melted_feat_imp_strat3$Predictor_Group[3:7] <- "Caller"
melted_feat_imp_strat3$pred_var_label <- c("Chromosome", "CNV Type", "CANOES", "CODEX", "CLAMMS", "XHMM", "Concordance", "Read Depth Ratio", "GC Content", 
                                           "Mappability", "Target Probe Count", "Size Label")
melted_feat_imp_strat3 <- sqldf("select * from melted_feat_imp_strat3 order by prop desc")

melted_feat_imp_strat3$pred_var_label <- 
  factor(melted_feat_imp_strat3$pred_var_label, levels = melted_feat_imp_strat3$pred_var_label[order(as.numeric(melted_feat_imp_strat3$prop))])
melted_feat_imp_strat3 <- melted_feat_imp_strat3[which(melted_feat_imp_strat3$pred_var_label != "CLAMMS"),]

pdf("supp_fig_3d_eq_feature_importance_randomforest_clamms_0.7.pdf", width = 8, height = 5)
ggplot(data=melted_feat_imp_strat3, aes(x=reorder(pred_var_label,-prop), y=prop)) + theme_classic() +
  geom_bar(position = "dodge", stat="identity") + 
  #  facet_grid(~Predictor_Group, scale = "free", switch = "x") +
  geom_text(aes(label=paste0(prop,"%")), position = position_dodge(width = 0.9), vjust = -.25, size = 3.5, fontface = "bold") +
  ggtitle ("CN-Learn - Feature Importance") +
  theme(plot.title = element_text(lineheight=.8, size=15, face="bold", hjust = 0.5), 
        strip.text = element_text(size = 12, face = "bold"),
        axis.title = element_text(size = 12, face="bold"),
        axis.text = element_text(size = 12, face="bold"), 
        #        axis.text.x = element_blank(),
        axis.text.x = element_text(angle = 60, hjust = 1),
        legend.title = element_text(size=12, face="bold"),
        legend.text = element_text(size=10, face="bold")) +
  labs(x = "Predictors", y = "Feature Importance", fill = "       Predictors") 
dev.off()
```
