## Supplemental_Code for "A machine-learning approach for accurate detection of copy-number variants from exome sequencing": CANOES.R

```
# DEPENDENCIES
#    nnls, Hmisc, mgcv, plyr

Test <- function(){
  # read in the data
  gc <- read.table("gc.txt")$V2
  canoes.reads <- read.table("canoes.reads.txt")
  # rename the columns of canoes.reads
  sample.names <- paste("S", seq(1:26), sep="")
  names(canoes.reads) <- c("chromosome", "start", "end", sample.names)
  # create a vector of consecutive target ids
  target <- seq(1, nrow(canoes.reads))
  # combine the data into one data frame
  canoes.reads <- cbind(target, gc, canoes.reads)
  # call CNVs in each sample
  # create a vector to hold the results for each sample
  xcnv.list <- vector('list', length(sample.names))
  for (i in 1:length(sample.names)){
    xcnv.list[[i]] <- CallCNVs(sample.names[i], canoes.reads) 
  }
  # combine the results into one data frame
  xcnvs <- do.call('rbind', xcnv.list)
  # inspect the first two CNV calls
  print(head(xcnvs, 2))
  # plot all the CNV calls to a pdf
  pdf("CNVplots.pdf")
  for (i in 1:nrow(xcnvs)){
     PlotCNV(canoes.reads, xcnvs[i, "SAMPLE"], xcnvs[i, "TARGETS"])
  }
  dev.off()
  # genotype all the CNVs calls made above in sample S2
  genotyping.S2 <- GenotypeCNVs(xcnvs, "S2", canoes.reads)
  # inspect the genotype scores for the first two CNV calls
  print(head(genotyping.S2, 2))
}

# Constants
NUM.ABNORMAL.STATES=2
NUM.STATES=3
DELETION=1
NORMAL=2
DUPLICATION=3

# PlotCNV
#     Plots count data for targets of interest
#     highlights sample of interest in red, 
#     highlights area of interest with a black line
#     highlights probe locations with black dots
# Arguments:
#   counts: 
#     count matrix, with column "target" with target numbers 
#     and sample data in columns 6:end
#   sample.name:
#     sample of interest (will be highlighted in red in figure)
#     (should correspond to a column in counts)
#   targets:
#     targets of interest in the form start.target..end.target
#   offset:
#     number of targets to add on either end (default=1)
# Returns: 
#   returns nothing
PlotCNV <- function(counts, sample.name, targets, offset=1){
  sample.name <- as.character(sample.name)
  if (!sample.name %in% names(counts)){stop("No column for sample ", sample.name, " in counts matrix")}
  if (length(setdiff("target", names(counts)[1:5]) > 0)){
    stop("counts matrix must have column named target")
  }
  t <- as.character(targets)
  start.target <- as.numeric(unlist(strsplit(t, "..", fixed=T))[1])
  end.target <- as.numeric(unlist(strsplit(t, "..", fixed=T))[2])
  if (!start.target %in% counts$target){
    stop("no data for start.target in counts matrix")
  }
  if (!end.target %in% counts$target){
    stop("no data for end.target in counts matrix")
  }
  if ((start.target - offset) %in% counts$target){
    start.target <- start.target - offset
  }
  if ((end.target + offset) %in% counts$target){
    end.target <- end.target + offset
  }
  ref.sample.names <- setdiff(as.character(names(counts)[-seq(1,5)]), 
                              sample.name)
  data <- subset(counts, target >= start.target & target <= end.target)
  sample.data <- data[, sample.name]
  means <- apply(data[, ref.sample.names], 1, mean)
  sd <- sqrt(apply(data[, ref.sample.names], 1, var))
  refs.z.scores <- matrix(NA, nrow(data), length(ref.sample.names))
  sample.z.score <- numeric(length = nrow(data))
  for (i in seq(1, dim(data)[1])){
    refs.z.scores[i, ] <- as.numeric((data[i, ref.sample.names] - means[i]) / 
                                       max(0.000001, sd[i]))
    sample.z.score[i] <- (sample.data[i] - means[i]) / max(0.000001, sd[i])
  }
  ylim <- max(abs(refs.z.scores), abs(sample.z.score))
  plot(seq(-6, 6), seq(-6, 6), 
       xlim=c(data[1, "start"], data[dim(data)[1], "start"]), 
       ylim=c(-ylim - 0.1, ylim + 0.1), type="n", xlab="", ylab="Z-score")
  for (i in seq(1, length(ref.sample.names))){
    lines(data[, "start"], refs.z.scores[, i], col="#2f4f4f85")
  }
  lines(data[, "start"], sample.z.score, col="red", lwd=3)
  points(data[, "start"], rep(-ylim - 0.05, length(data[, "start"])), pch=20)
  lines( c(data[1 + offset, "start"], data[nrow(data) - offset, "end"]) , 
         c(ylim+0.2, ylim+0.2), lwd=2)
  title(main=paste("Sample ", sample.name, ", ", 
                   counts$chromosome[start.target], ":", 
                   data$start[1], "-", data$end[nrow(data)], sep=""))
}

# CallCNVs
#     Calls CNVs in sample of interest
# Arguments:
#   sample.name:
#     sample to call CNVs in (should correspond to a column in counts)
#   counts: 
#     count matrix, first five columns should be 
#       target: consecutive numbers for targets (integer)
#       chromosome: chromosome number (integer-valued) 
#         (support for sex chromosomes to come)
#       start: start position of probe (integer)
#       end: end position of probe (integer)
#       gc: gc content (real between 0 and 1)
#       subsequent columns should include counts for each probe for samples
#   p:
#     average rate of occurrence of CNVs (real) default is 1e-08
#   D:
#     expected distance between targets in a CNV (integer) default is 70,000
#   Tnum:
#     expected number of targets in a CNV (integer) default is 6
#   numrefs
#     maximum number of reference samples to use (integer) default is 30
#     the weighted variance calculations will take a long time if too 
#     many reference samples are used
# Returns: 
#   data frame with the following columns:
#      SAMPLE: name of sample
#      CNV: DEL of DUP
#      INTERVAL: CNV coordinates in the form chr:start-stop
#      KB: length of CNV in kilobases
#      CHR: chromosome
#      MID_BP: middle base pair of CNV
#      TARGETS: target numbers of CNV in the form start..stop
#      NUM_TARG: how many targets are in the CNV
#      Q_SOME: a Phred-scaled quality score for the CNV
CallCNVs <- function(sample.name, counts, p=1e-08, Tnum=6, D=70000, numrefs=30, get.dfs=F, homdel.mean=0.2){
  if (!sample.name %in% names(counts)){stop("No column for sample ", sample.name, " in counts matrix")}
  if (length(setdiff(names(counts)[1:5], c("target", "chromosome", "start", "end", "gc"))) > 0){
    stop("First five columns of counts matrix must be target, chromosome, start, end, gc")
  }
  if (length(setdiff(unique(counts$chromosome), seq(1:22))) > 0) {
    # remove sex chromosomes
    cat("Trying to remove sex chromosomes and 'chr' prefixes\n")
    counts <- subset(counts, !chromosome %in% c("chrX", "chrY", "X", "Y"))
    if (sum(grepl("chr", counts$chromosome))==length(counts$chromosome)){
      counts$chromosome <- gsub("chr", "", counts$chromosome)
    }
    counts$chromosome <- as.numeric(counts$chromosome)
    if (length(setdiff(unique(counts$chromosome), seq(1:22))) > 0) 
      stop("chromosome must take value in range 1-22 (support for sex chromosomes to come)")
  }
  library(plyr)
  counts <- arrange(counts, chromosome, start)
  if (p <= 0){
    stop("parameter p must be positive")
  }
  if (Tnum <= 0){
    stop("parameter Tnum must be positive")
  }
  if (D <= 0){
    stop("parameter D must be positive")
  }
  if (numrefs <= 0){
    stop("parameter numrefs must be positive")
  }
  sample.names <- colnames(counts)[-seq(1,5)]
  # find mean coverage of probes
  mean.counts <- mean(apply(counts[, sample.names], 2, mean))
  # normalize counts; round so we can use negative binomial
  counts[, sample.names] <- apply(counts[, sample.names], 2, 
        function(x, mean.counts) 
                 round(x * mean.counts / mean(x)), mean.counts)
  # calculate covariance of read count across samples
  cov <- cor(counts[, sample.names], counts[, sample.names])
  reference.samples <- setdiff(sample.names, sample.name)
  covariances <- cov[sample.name, reference.samples]
  reference.samples <- names(sort(covariances, 
          decreasing=T)[1:min(numrefs, length(covariances))])
  sample.mean.counts <- mean(counts[, sample.name])
  sample.sumcounts <- apply(counts[, reference.samples], 2, sum)
  # normalize reference samples to sample of interest
  counts[, reference.samples] <- apply(counts[, reference.samples], 2, 
        function(x, sample.mean.counts) 
                round(x * sample.mean.counts / 
                mean(x)), sample.mean.counts)  
  # select reference samples and weightings using non-negative least squares
  b <- counts[, sample.name]
  A <- as.matrix(counts[, reference.samples])
  library(nnls)
  all <- nnls(A, b)$x
  est <- matrix(0, nrow=50, ncol=length(reference.samples))
  set.seed(1)
  for (i in 1:50){
    d <- sample(nrow(A), min(500, nrow(A)))
    est[i, ] <- nnls(A[d, ], b[d])$x
  }
  weights <- colMeans(est)
  sample.weights <- weights / sum(weights)
  library(Hmisc)
  # calculate weighted mean of read count
  # this is used to calculate emission probabilities
  counts$mean <- apply(counts[, reference.samples], 
                       1, wtd.mean, sample.weights)
  targets <- counts$target
  # exclude probes with all zero counts
  nonzero.rows <- counts$mean > 0
  nonzero.rows.df <- data.frame(target=counts$target, 
                                nonzero.rows=nonzero.rows)

  counts <- counts[nonzero.rows, ]
  # get the distances between consecutive probes
  distances <- GetDistances(counts)
  # estimate the read count variance at each probe
  var.estimate <- EstimateVariance(counts, reference.samples, 
                                               sample.weights)
  emission.probs <- EmissionProbs(counts[, sample.name], 
                        counts$mean, var.estimate$var.estimate, 
                        counts[, "target"])
  if (get.dfs){
    return(list(emission.probs=emission.probs, distances=distances))
  }
  # call CNVs with the Viterbi algorithm
  viterbi.state <- Viterbi(emission.probs, distances, p, Tnum, D)  
  # format the CNVs
  cnvs <- PrintCNVs(sample.name, viterbi.state, 
                         counts)
  # if there aren't too many CNVs, calculate the Q_SOME
  if (nrow(cnvs) > 0 & nrow(cnvs) <= 50){
    qualities <- GenotypeCNVs(cnvs, sample.name, counts, p, Tnum, D, numrefs, 
                          emission.probs=emission.probs, 
                          distances=distances)
    for (i in 1:nrow(cnvs)){
      cnvs$Q_SOME[i] <- ifelse(cnvs$CNV[i]=="DEL", qualities[i, "SQDel"], 
                               qualities[i, "SQDup"])
    }
  }
  data <- as.data.frame(cbind(counts$target, counts$mean, var.estimate$var.estimate, counts[, sample.name]))
  names(data) <- c("target", "countsmean", "varestimate", "sample")
  if (nrow(cnvs) > 0){
    cnvs <- CalcCopyNumber(data, cnvs, homdel.mean)
  }
  return(cnvs)
}

# GenotypeCNVs
#     Genotype CNVs in sample of interest
# Arguments:
#   xcnv
#     data frame with the following columns, and one row for each
#     CNV to genotype
#      INTERVAL: CNV coordinates in the form chr:start-stop
#      TARGETS: target numbers of CNV in the form start..stop
#               these should correspond to the target numbers in counts
#   sample.name:
#     sample to genotype CNVs in (should correspond to a column in counts)
#   counts: 
#     count matrix, first five columns should be 
#       target: consecutive numbers for targets (integer)
#       chromosome: chromosome number (integer-valued) 
#         (support for sex chromosomes to come)
#       start: start position of probe (integer)
#       end: end position of probe (integer)
#       gc: gc content (real between 0 and 1)
#       subsequent columns should include counts for each probe for samples
#   p:
#     average rate of occurrence of CNVs (real) default is 1e-08
#   D:
#     expected distance between targets in a CNV (integer) default is 70,000
#   Tnum:
#     expected number of targets in a CNV (integer) default is 6
#   numrefs
#     maximum number of reference samples to use (integer) default is 30
#     the weighted variance calculations will take a long time if too 
#     many reference samples are used
#   emission.probs and distances are for internal use only
# Returns: 
#   data frame with the following columns and one row for each genotyped CNV:
#      INTERVAL: CNV coordinates in the form chr:start-stop
#      NQDEL: a Phred-scaled quality score that sample.name has no deletion 
#             in the interval
#      SQDEL: a Phred-scaled quality score that sample.name has a deletion 
#             in the interval
#      NQDUP and SQDUP: same, but for a duplication
GenotypeCNVs <- function(xcnvs, sample.name, counts, p=1e-08, Tnum=6, 
                    D=70000, numrefs=30,
                    emission.probs=NULL, 
                    distances=NULL){
  if (!sample.name %in% names(counts)){stop("No column for sample ", sample.name, " in counts matrix")}
  if (length(setdiff(names(counts)[1:5], c("target", "chromosome", "start", "end", "gc"))) > 0){
    stop("First five columns of counts matrix must be target, chromosome, start, end, gc")
  }
  if (length(setdiff(unique(counts$chromosome), seq(1:22))) > 0) {
    # remove sex chromosomes
    cat("Trying to remove sex chromosomes and 'chr' prefixes\n")
    counts <- subset(counts, !chromosome %in% c("chrX", "chrY", "X", "Y"))
    if (sum(grepl("chr", counts$chromosome))==length(counts$chromosome)){
      counts$chromosome <- gsub("chr", "", counts$chromosome)
    }
    counts$chromosome <- as.numeric(counts$chromosome)
    if (length(setdiff(unique(counts$chromosome), seq(1:22))) > 0) 
      stop("chromosome must take value in range 1-22 (support for sex chromosomes to come)")
  }
  library(plyr)
  counts <- arrange(counts, chromosome, start)
  if (p <= 0){
    stop("parameter p must be positive")
  }
  if (Tnum <= 0){
    stop("parameter Tnum must be positive")
  }
  if (D <= 0){
    stop("parameter D must be positive")
  }
  if (numrefs <= 0){
    stop("parameter numrefs must be positive")
  }
  num.cnvs <- nrow(xcnvs)
  cnv.intervals <- as.character(xcnvs$INTERVAL)
  # if no emission probs matrix is passed in, generate a new one
  if (is.null(emission.probs)){
    l <- CallCNVs(sample.name, counts, p, Tnum=6, D=70000, numrefs=30, get.dfs=T)
    emission.probs <- l[['emission.probs']]
    distances <- l[['distances']]
  }
  forward.m <- GetForwardMatrix(emission.probs, distances, p, Tnum, D)
  backward.m <- GetBackwardMatrix(emission.probs, distances, p, Tnum, D)
  qualities <- matrix(0, nrow=num.cnvs, ncol=5, 
                      dimnames=list(cnv.intervals, 
                                    c("INTERVAL", "NQDel", "SQDel", "NQDup", "SQDup")))
  for (i in 1:num.cnvs){
    interval <- as.character(xcnvs[i, "INTERVAL"])
    targets <- as.numeric(strsplit(as.character(xcnvs[i, "TARGETS"]), ".", fixed=T)[[1]][c(1,3)])
    left.target <- targets[1]
    right.target <- targets[2]
    likelihoods <- GetModifiedLikelihood(forward.m, backward.m, 
                                         emission.probs, distances, 
                                         left.target, right.target, 
                                         c(DUPLICATION, DELETION), p, Tnum, D)
    modified.likelihood <- likelihoods[1]; 
    unmodified.likelihood <- likelihoods[2]
    Prob.All.Normal <- exp(modified.likelihood - unmodified.likelihood)
    likelihoods <- GetModifiedLikelihood(forward.m, backward.m, 
                                         emission.probs, distances, 
                                         left.target, right.target, DELETION, p, Tnum, D)
    modified.likelihood <- likelihoods[1]; 
    unmodified.likelihood <- likelihoods[2]
    Prob.No.Deletion <- exp(modified.likelihood - unmodified.likelihood)
    likelihoods <- GetModifiedLikelihood(forward.m, backward.m, 
                                         emission.probs, distances, 
                                         left.target, right.target, DUPLICATION, p, Tnum, D)
    modified.likelihood <- likelihoods[1]; 
    unmodified.likelihood <- likelihoods[2]
    Prob.No.Duplication <- exp(modified.likelihood - unmodified.likelihood)
    # Check if probabilities greater than 1 are numerical error or bug
    Phred <- function(prob){
      return(round(min(99, -10 * log10(1 - prob))))
    }
    qualities[i, "NQDel"] <- Phred(Prob.No.Deletion)       
    qualities[i, "SQDel"] <- Phred(Prob.No.Duplication - Prob.All.Normal)
    qualities[i, "NQDup"] <- Phred(Prob.No.Duplication)       
    qualities[i, "SQDup"] <- Phred(Prob.No.Deletion - Prob.All.Normal)
    qualities[i, "INTERVAL"] <- interval
  }
  qualities <- as.data.frame(qualities, stringsAsFactors=F)
  qualities$NQDel <- as.integer(qualities$NQDel)
  qualities$NQDup <- as.integer(qualities$NQDup)
  qualities$SQDel <- as.integer(qualities$SQDel)
  qualities$SQDup <- as.integer(qualities$SQDup)
  return(qualities)
}

# returns data frame with distance to each target from the previous target 
# (0 in the case of the first target on chromosome 1, a very big number
# for the first target on each other chromosome--this resets the HMM
# for each chromosome)
GetDistances <- function(counts){
  chromosome <- counts[, "chromosome"]
  startbase <- counts[, "start"]
  num.nonzero.exons <- length(startbase)
  distances <- c(0, startbase[2:num.nonzero.exons] - 
                   startbase[1:(num.nonzero.exons - 1)] + 
                   1000000000000 * (chromosome[2:num.nonzero.exons] - 
                                      chromosome[1:(num.nonzero.exons - 1)]))
  return(data.frame(target=counts[, "target"], distance=distances))
}

EstimateVariance <- function(counts, ref.sample.names, sample.weights){
  library(Hmisc)
  counts$var <- apply(counts[, ref.sample.names], 1, wtd.var, sample.weights, normwt=T)
  set.seed(1)
  counts.subset <- counts[sample(nrow(counts), min(36000, nrow(counts))), ]
  library(mgcv)
  # can't do gamma regression with negative 
  counts.subset$var[counts.subset$var==0] <- 0.1 
  fit <- gam(var ~ s(mean) + s(gc), family=Gamma(link=log), data=counts.subset)
  # we don't want variance less than Poisson
  # we take maximum of genome-wide estimate, method of moments estimate
  # and Poisson variance
  v.estimate <- pmax(predict(fit, counts, type="response"), counts$var, 
                     counts$mean * 1.01)
  return(data.frame(target=counts$target, var.estimate=v.estimate))
}

EmissionProbs <- function(test.counts, target.means, 
                                      var.estimate, targets){
  num.targets <- length(test.counts)
  # calculate the means for the deletion, normal and duplication states
  state.target.means <- t(apply(data.frame(x=target.means), 1, function(x) c(x*1/2, x, x*3/2)))
  # calculate the expected size (given the predicted variance)
  size <- target.means ^ 2 / (var.estimate - target.means)
  emission.probs <- matrix(NA, num.targets, 4)
  colnames(emission.probs) <- c("target", "delprob", "normalprob", "dupprob")
  # calculate the emission probabilities given the read count
  size.del <- size
  size.dup <- size
  size.del <- size / 2
  size.dup <- size * 3 / 2
  emission.probs[, "delprob"] <- dnbinom(
    test.counts,
    mu=state.target.means[, 1],
    size=size.del, log=T)
  emission.probs[, "normalprob"] <- dnbinom(
    test.counts,
    mu=state.target.means[, 2],
    size=size, log=T)
  emission.probs[, "dupprob"] <- dnbinom(
    test.counts,
    mu=state.target.means[, 3],
    size=size.dup, log=T)
  emission.probs[, "target"] <- targets
  # some values may be infinite as a result of extreme read count
  row.all.inf <- which(apply(emission.probs, 1, function(x){all(is.infinite(x))}))
  if (length(row.all.inf) > 0){
    for (i in row.all.inf){
      if (test.counts[i] >= state.target.means[i, 3]){
        emission.probs[i, 2:4] <- c(-Inf, -Inf, -0.01)
      }
      else if (test.counts[i] <= state.target.means[i, 1]){
        emission.probs[i, 2:4] <- c(-0.01, -Inf, -Inf)
      }
      else emission.probs[i, 2:4] <- c(-Inf, -0.01, -Inf)
    }
  }
  return(emission.probs)
}

# Viterbi algorithm
Viterbi <- function(emission.probs.matrix, distances, p, Tnum, D){
  targets <- emission.probs.matrix[, 1]
  emission.probs.matrix <- as.matrix(emission.probs.matrix[, 2:4])
  num.exons <- dim(emission.probs.matrix)[1]
  viterbi.matrix <- matrix(NA, nrow=num.exons, ncol=NUM.STATES)
  viterbi.pointers <- matrix(NA, nrow=num.exons, ncol=NUM.STATES)
  initial.state <- log(c(0.0075 / NUM.ABNORMAL.STATES, 1 - 0.0075, 0.0075 / NUM.ABNORMAL.STATES))
  viterbi.matrix[1, ] <- initial.state + emission.probs.matrix[1,]
  for (i in 2:num.exons) {
    temp.matrix <- viterbi.matrix[i - 1, ] + GetTransitionMatrix(distances$distance[i], p, Tnum, D)
    viterbi.matrix[i, ] <- apply(temp.matrix, 2, max)
    emission.probs <- c(emission.probs.matrix[i,])
    dim(emission.probs) <- c(NUM.STATES, 1)
    viterbi.matrix[i, ] <- viterbi.matrix[i, ] + emission.probs
    viterbi.pointers[i, ] <- apply(temp.matrix, 2, which.max)
  }
  viterbi.states = vector(length = num.exons)
  viterbi.states[num.exons] = which.max(viterbi.matrix[num.exons, ])
  for (i in (num.exons - 1):1) {
    viterbi.states[i] <- viterbi.pointers[i + 1, viterbi.states[i + 1]]
  }
  return(data.frame(target=targets, viterbi.state=viterbi.states))
}

# returns a transition matrix
#                              to state
#                    deletion   normal    duplication
#           deletion   
#from state   normal
#        duplication
GetTransitionMatrix <- function(distance, p, Tnum, D){
  q <- 1 / Tnum
  f = exp(-distance/D)
  prob.abnormal.abnormal <- f * (1 - q) + (1 - f) * p
  prob.abnormal.normal <- f * q + (1 - f) * (1 - 2 * p)
  prob.abnormal.diff.abnormal <- (1 - f) * p
  prob.normal.normal <- 1 - 2 * p
  prob.normal.abnormal <- p
  transition.probs <- 
    c(prob.abnormal.abnormal, prob.abnormal.normal, prob.abnormal.diff.abnormal, 
      prob.normal.abnormal, prob.normal.normal, prob.normal.abnormal,
      prob.abnormal.diff.abnormal, prob.abnormal.normal, prob.abnormal.abnormal)
  transition.m = log(matrix(transition.probs, NUM.STATES, NUM.STATES, byrow=TRUE))
  return(transition.m)
}

# adds two log-space probabilities using the identity
# log (p1 + p2) = log p1 + log(1 + exp(log p2 - log p1))
AddTwoProbabilities <- function(x, y){
  if (is.infinite(x)) return (y)
  if (is.infinite(y)) return (x)
  sum.probs <- max(x, y) + log1p(exp(-abs(x - y)))
}

# adds multiple log-space probabilities
SumProbabilities <- function(x){
  sum.probs <- x[1]
  for (i in 2:length(x)){
    sum.probs <- AddTwoProbabilities(sum.probs, x[i])
  }
  return(sum.probs)
}

# finds the data likelihood by summing the product of the corresponding 
# forward and backward probabilities at any token (should give the same value
# regardless of the token)
GetLikelihood <- function(forward.matrix, backward.matrix, x){
  SumProbabilities(forward.matrix[x, ] + backward.matrix[x, ])
}

# get the forward probabilities
GetForwardMatrix <- function(emission.probs.matrix, distances, p, Tnum, D){
  emission.probs.matrix <- as.matrix(emission.probs.matrix[, 2:4])
  num.exons <- dim(emission.probs.matrix)[1]
  forward.matrix <- matrix(NA, nrow=num.exons, ncol=NUM.STATES)   # matrix to hold forward probabilities
  initial.state <- log(c(0.0075 / NUM.ABNORMAL.STATES, 1 - 0.0075, 0.0075 / NUM.ABNORMAL.STATES))
  forward.matrix[1, ] <- initial.state + emission.probs.matrix[1, ]
  for (i in 2:num.exons){
    # compute matrix with probability we were in state j and are now in state i
    # in temp.matrix[j, i] (ignoring emission of current token)
    temp.matrix <- forward.matrix[i - 1, ] + GetTransitionMatrix(distances$distance[i], p, Tnum, D)
    # find the probability that we are in each of the three states
    sum.probs <- apply(temp.matrix, 2, SumProbabilities)
    forward.matrix[i, ] <- sum.probs + emission.probs.matrix[i, ]
  }  
  return(forward.matrix)  
}

# get the backward probabilities
GetBackwardMatrix <- function(emission.probs.matrix, distances, 
                                  p, Tnum, D){
  emission.probs.matrix <- as.matrix(emission.probs.matrix[, 2:4])
  num.exons <- dim(emission.probs.matrix)[1]
  backward.matrix <- matrix(NA, nrow=num.exons, ncol=NUM.STATES)   # matrix to hold backward probabilities
  initial.state <- log(c(0.0075 / NUM.ABNORMAL.STATES, 1 - 0.0075, 0.0075 / NUM.ABNORMAL.STATES))
  backward.matrix[num.exons, ] <- rep(0, NUM.STATES)
  for (i in (num.exons - 1):1){
    temp.matrix <- GetTransitionMatrix(distances$distance[i+1], p, Tnum, D) + 
      matrix(backward.matrix[i + 1, ], 3, 3, byrow=T) +
      matrix(emission.probs.matrix[i+1, ], 3, 3, byrow=T)
    backward.matrix[i, ] <- apply(temp.matrix, 1, SumProbabilities)
  }  
  final.prob <- backward.matrix[1, ] + emission.probs.matrix[1, ] + initial.state
  return(backward.matrix)  
}

# find the likelihood of the data given that certain states are disallowed
# between start target and end target
GetModifiedLikelihood <- function(forward.matrix, backward.matrix, emission.probs.matrix, distances, 
                                      start.target, end.target, disallowed.states, p, Tnum, D){
  targets <- emission.probs.matrix[, 1]
  emission.probs.matrix <- as.matrix(emission.probs.matrix[, 2:4])
  # there may be missing targets in this sample, we genotype the largest stretch of 
  # targets that lie in the CNV
  left.target <- min(which(targets >= start.target))
  right.target <- max(which(targets <= end.target))
  num.exons <- dim(emission.probs.matrix)[1]
  unmodified.likelihood <- GetLikelihood(forward.matrix, 
                                             backward.matrix, min(right.target + 1, num.exons))
  #right.target or left.target may be empty
  
  #if (right.target >= left.target) return(c(NA, unmodified.likelihood))
  stopifnot(right.target >= left.target)
  modified.emission.probs.matrix <- emission.probs.matrix
  modified.emission.probs.matrix[left.target:right.target, 
                                 disallowed.states] <- -Inf
  
  # if the start target is the first target we need to recalculate the 
  # forward probabilities
  # for that target, using the modified emission probabilities
  if (left.target == 1){
    initial.state <- log(c(0.0075 / NUM.ABNORMAL.STATES, 1 - 0.0075, 0.0075 / NUM.ABNORMAL.STATES))
    forward.matrix[1, ] <- initial.state + modified.emission.probs.matrix[1, ]
    left.target <- left.target + 1
  } 
  for (i in seq(left.target, min(right.target + 1, num.exons))){
    # compute matrix with probability we were in state j and are now in state i
    # in temp.matrix[j, i] (ignoring emission of current token)
    temp.matrix <- forward.matrix[i - 1, ] + GetTransitionMatrix(distances$distance[i], p, Tnum, D)
    # find the probability that we are in each of the three states
    sum.probs <- apply(temp.matrix, 2, SumProbabilities) 
    if (!i == (right.target + 1)){
      forward.matrix[i, ] <- sum.probs + modified.emission.probs.matrix[i, ]
    } else{
      forward.matrix[i, ] <- sum.probs + emission.probs.matrix[i, ]
    }
  }  
  # find the modified likelihood of the sequence
  modified.likelihood <- GetLikelihood(forward.matrix, backward.matrix, min(right.target + 1, num.exons))
  return(c(modified.likelihood, unmodified.likelihood))
}

SummarizeCNVs <- function(cnv.targets, counts, sample.name, state){
  sample.name <- sample.name
  cnv.type <- ifelse(state==3, "DUP", "DEL")
  cnv.start <- min(cnv.targets$target)
  cnv.end <- max(cnv.targets$target)
  cnv.chromosome <- counts[cnv.start, "chromosome"]
  cnv.start.base <- counts[cnv.start, "start"]
  cnv.start.target <- counts[cnv.start, "target"]
  cnv.end.base <- counts[cnv.end, "end"]
  cnv.end.target <- counts[cnv.end, "target"]
  cnv.kbs <- (cnv.end.base - cnv.start.base) / 1000
  cnv.midbp <- round((cnv.end.base - cnv.start.base) / 2) + cnv.start.base
  cnv.targets <- paste(cnv.start.target, "..", cnv.end.target, sep="")
  cnv.interval <- paste(cnv.chromosome, ":", cnv.start.base, "-", cnv.end.base, sep="")
  num.targets <- cnv.end.target - cnv.start.target + 1
  return(data.frame(sample.name=sample.name, cnv.type=cnv.type, cnv.interval=cnv.interval, 
                    cnv.kbs=cnv.kbs, cnv.chromosome=cnv.chromosome, 
                    cnv.midbp=cnv.midbp, cnv.targets=cnv.targets, num.targets=num.targets))
}

PrintCNVs <- function(test.sample.name, viterbi.state, 
                      nonzero.counts){  
  consecutiveGroups <- function(sequence){
    num <- length(sequence)
    group <- 1
    groups <- rep(0, num)
    groups[1] <- group
    if (num > 1){
      for (i in 2:num){
        if (!sequence[i] == (sequence[i - 1] + 1)) group <- group + 1
        groups[i] <- group
      }
    }
    return(groups)
  }
  num.duplications <- 0
  num.deletions <- 0
  for (state in c(1, 3)){
    cnv.targets <- which(viterbi.state$viterbi.state == state)
    if (!length(cnv.targets) == 0){
      groups <- consecutiveGroups(cnv.targets)
      library(plyr)
      cnvs.temp.df <- ddply(data.frame(target=cnv.targets, group=groups), 
                            "group", SummarizeCNVs, nonzero.counts, test.sample.name, 
                            state)
      if (state == 1){
        deletions.df <- cnvs.temp.df
        if (!is.null(dim(deletions.df))){
          num.deletions <- dim(deletions.df)[1]
        }
      } else {
        duplications.df <- cnvs.temp.df
        if (!is.null(dim(duplications.df))){
          num.duplications <- dim(duplications.df)[1]
        }
      }
    }
  }
  num.calls <- num.deletions + num.duplications
  cat(num.calls, "CNVs called in sample", test.sample.name, "\n")
  if (num.deletions == 0 & num.duplications == 0){
    df <- data.frame(SAMPLE=character(0), CNV=character(0), INTERVAL=character(0), 
                     KB=numeric(0), CHR=character(0), 
                     MID_BP=numeric(), TARGETS=character(0), NUM_TARG=numeric(0), Q_SOME=numeric(0), MLCN=numeric(0))
    return(df)
  }
  if (num.deletions > 0 & num.duplications > 0){
    cnvs.df <- rbind(deletions.df, duplications.df)
  } else {
    ifelse(num.deletions > 0, 
           cnvs.df <- deletions.df, cnvs.df <- duplications.df)
  }
  xcnv <- cbind(cnvs.df[, c("sample.name", "cnv.type", "cnv.interval", 
                      "cnv.kbs", "cnv.chromosome", "cnv.midbp", 
                      "cnv.targets", "num.targets")], 0)
  colnames(xcnv) <- c("SAMPLE", "CNV", "INTERVAL", "KB", "CHR", "MID_BP", "TARGETS",
                      "NUM_TARG", "MLCN")
  xcnv$Q_SOME <- NA
  return(xcnv)
}

CalcCopyNumber <- function(data, cnvs, homdel.mean){
  for (i in 1:nrow(cnvs)){
    cnv <- cnvs[i, ]
    targets <- as.numeric(unlist(strsplit(as.character(cnv$TARGETS), "..", fixed=T)))
    cnv.data <- subset(data, target >= targets[1] & target <= targets[2])
    state.target.means <- t(apply(data.frame(x=cnv.data$countsmean), 1, 
                                  function(x) c(C1=x*1/2, C2=x, C3=x*3/2, 
                                                C4=x * 2, C5=x * 5/2, C6=x*6/2)))
    # calculate the expected size (given the predicted variance)
    size <- cnv.data$countsmean ^ 2 / (cnv.data$varestimate - cnv.data$countsmean)
    emission.probs <- matrix(NA, nrow(cnv.data), 7)
    colnames(emission.probs) <- c("C0", "C1", "C2", "C3", "C4", "C5", "C6")
    #colnames(emission.probs) <- c("target", "delprob", "normalprob", "dupprob")
    # calculate the emission probabilities given the read count
    emission.probs[, 1] <- dpois(cnv.data$sample, homdel.mean, log=T)
    for (s in 1:6){
      size.state <- size * s/2
      emission.probs[, s+1] <- dnbinom(cnv.data$sample, mu=state.target.means[, s], 
                                       size=size.state, log=T)
    }
    cs <- colSums(emission.probs)
    ml.state <- which.max(cs) - 1
    if (ml.state==2){
      ml.state <- ifelse(cnv$CNV=="DEL", 1, 3)
    }
    cnvs$MLCN[i] <- ml.state
  }  
  return(cnvs)
}
```
