## Supplemental_Code for "A machine-learning approach for accurate detection of copy-number variants from exome sequencing": canoes_merge_files.r

```
#############################################################
# Author: Vijay Kumar                                       #
# Date  : 4/5/2019                                          #
# This script is supplied as part of the CANOES pipeline.   #
# It has been modified to take in parameters from the shell # 
# script. It has been made more readable with additional    # 
# comments. It also sources the R script CANOES.R from the  #
# rscripts directory.                                       #
#############################################################
############################################################################
# STEP 1: Fetch the directory locations and file names from the shell script
############################################################################
args <- commandArgs(trailingOnly=TRUE)
scripts_dir <- args[1]
data_dir <- args[2]
input_file_name <- args[3]
output_file_name <- args[4]

######################################################
# STEP 2: Read the read count file and GC content file
######################################################
setwd(data_dir)
gc <- read.table("gc.txt")$V2
canoes.reads <- read.table(input_file_name)

########################################################
# STEP 3: Assign column names and merge columns together
########################################################
num_cols <- dim(canoes.reads)[2]
num_samples <- num_cols - 3
sample.names <- paste("S", seq(1: num_samples), sep="")
names(canoes.reads) <- c("chromosome", "start", "end", sample.names)
target <- seq(1, nrow(canoes.reads))
canoes.reads <- cbind(target, gc, canoes.reads)

######################################################
# STEP 4: Load R script and process each column/sample
######################################################
setwd(scripts_dir)
source("CANOES.R")
xcnv.list <- vector('list', length(sample.names))
for (i in 1:length(sample.names)){
  xcnv.list[[i]] <- CallCNVs(sample.names[i], canoes.reads)
}
################################################################################
# STEP 5: Append output as rows onto a single variable and write the output file
################################################################################
xcnvs <- do.call('rbind', xcnv.list)

setwd(data_dir)
write.table(xcnvs, file=output_file_name, sep=",")
```
