## Supplemental_Code for "A machine-learning approach for accurate detection of copy-number variants from exome sequencing": codex.r

```
library(CODEX)

args <- commandArgs(1)

#Creates codex objects from input info
chr <- args[1]
dirPath <- as.matrix(read.table(args[2]))
sampname <- as.matrix(read.table(args[3]))
bedFile <- args[4]
projName <- args[5]
output_dir <- args[6]

###################
#Merges input 
###################

#bambedObj <- getbambed(bamdir = bamdir, bedFile = bedFile, sampname = sampname, projectname = projName, chr)
bambedObj <- getbambed(bamdir = dirPath, bedFile = bedFile, sampname = sampname, projectname = projName, chr)
bamdir <- bambedObj$bamdir; sampname <- bambedObj$sampname
ref <- bambedObj$ref; projectname <- bambedObj$projectname; chr <- bambedObj$chr

#gets coverage, read lengths
coverageObj <- getcoverage(bambedObj, mapqthres = 20)
Y <- coverageObj$Y; readlength <- coverageObj$readlength

#gets gc content, mappability
gc <- getgc(chr, ref)
mapp <- getmapp(chr, ref)

#quality control
qcObj <- qc(Y, sampname, chr, ref, mapp, gc, cov_thresh = c(20, 4000), length_thresh = c(20, 2000), mapp_thresh = 0.9, gc_thresh = c(20, 80))
Y_qc <- qcObj$Y_qc; sampname_qc <- qcObj$sampname_qc; gc_qc <- qcObj$gc_qc
mapp_qc <- qcObj$mapp_qc; ref_qc <- qcObj$ref_qc; qcmat <- qcObj$qcmat
#write.table(qcmat, file = paste(projectname, _, chr, _qcmat, .txt, sep=), sep=\t, quote=FALSE, row.names=FALSE)

#normalization
normObj <- normalize(Y_qc, gc_qc, K = 1:9)
Yhat <- normObj$Yhat; AIC <- normObj$AIC; BIC <- normObj$BIC
RSS <- normObj$RSS; K <- normObj$K

#looks at different metrics to create deviance reduction plots, puts them in a pdf
#choiceofK(AIC, BIC, RSS, K, filename = paste(projectname, "_", chr, "_choiceofK", ".pdf", sep = ""))

#looks for CNVs, outputs to table
optK = K[which.max(BIC)]
finalcall <- segment(Y_qc, Yhat, optK = optK, K = K, sampname_qc, ref_qc, chr, lmax = 200, mode = "integer")
write.table(finalcall, file = paste(output_dir,'/',projectname, '_', chr,'_', optK, '_CODEX_frac.txt', sep=''), sep='\t', quote=FALSE, row.names=FALSE)
```
