## Supplemental_Code for "A machine-learning approach for accurate detection of copy-number variants from exome sequencing": consolidate_val_ov.r

```
################################################################################
# Script : consolidate_val_ov.r                                                #
# Author : Vijay Kumar                                                         #
# Date   : 4/5/2019                                                            #
#                                                                              #
# This script groups the final set of CNVs after they are labelled using       #
# validated CNVs. This is done to handle situations where the same CNV overlap # 
# with multiple validated CNVs. It also creates new variables that capture CNV #
# size and validation label.                                                   #
#                                                                              #
# (c) 2019 - Vijay Kumar                                                       #
# Licenced under the GNU General Public License 3.0.                           #
################################################################################
options(gsubfn.engine = "R")
library(sqldf)

args = commandArgs(trailingOnly=TRUE)
input_dir = args[1]
train_file = args[2]
test_file = args[3]
train_out_file = args[4]
test_out_file = args[5]
caller_count = args[6]

caller_list <- character(caller_count)
for (caller_number in 1:caller_count){
    caller_list[caller_number] <- args[6 + caller_number]
} 

#Set working directory
setwd(input_dir)

train_data <- as.data.frame(read.table(train_file))
test_data <- as.data.frame(read.table(test_file))

#####################################################################
# STEP 1: Label the CNVs based on their overlap with validated CNVs #
#####################################################################
dyn_col_names <- character(0)
for (caller_name in caller_list) {
    dyn_col_names <- append(dyn_col_names, caller_name)
}

colnames(train_data) = c('CHR','PRED_START', 'PRED_END','TYPE', 'SAMPLE', dyn_col_names, 
                         'NUM_OVERLAPS', 'RD_PROP', 'GC', 'PRED_SIZE', 'MAP', 'NUM_TARGETS', 
                         'OV_IND', 'OV_START','OV_END', 'MA_TYPE', 'MA_SIZE', 'MA_SAMPLE', 'OV_SIZE')
colnames(test_data) = c('CHR','PRED_START', 'PRED_END','TYPE', 'SAMPLE', 
                       dyn_col_names, 'NUM_OVERLAPS', 'RD_PROP', 'GC', 
                       'PRED_SIZE', 'MAP', 'NUM_TARGETS') 

dyn_query_string <- character(0)
for (caller_name in caller_list) {
    dyn_query_string <- paste(dyn_query_string, caller_name, sep =", ") 
}

sql_query_1 <- paste0("select CHR, PRED_START, PRED_END, TYPE, SAMPLE ", dyn_query_string, 
                      ", NUM_OVERLAPS, RD_PROP, round(GC,2) as GC, PRED_SIZE, ",
                      "round(MAP) as MAP, NUM_TARGETS, max(OV_IND) as OV_IND ",
                      "from train_data ",
                      "group by CHR, PRED_START, PRED_END, TYPE, SAMPLE ", dyn_query_string, 
                      ", NUM_OVERLAPS, RD_PROP, round(GC,2), PRED_SIZE, round(MAP,2), NUM_TARGETS")

train_df <- sqldf(sql_query_1)

sql_query_2 <- paste0("select CHR, PRED_START, PRED_END, TYPE, SAMPLE ", dyn_query_string, ", 
                       NUM_OVERLAPS, RD_PROP, round(GC,2) as GC, PRED_SIZE, round(MAP) as MAP, 
                       NUM_TARGETS ",
                       "from test_data ",
                       "group by CHR, PRED_START, PRED_END, TYPE, SAMPLE ", 
                       dyn_query_string, ", NUM_OVERLAPS, RD_PROP, round(GC,2), 
                       PRED_SIZE, round(MAP,2), NUM_TARGETS")

test_df <- sqldf(sql_query_2)

########################################################################
# STEP 2: Generate additional variables needed for subsequent analysis #
########################################################################
train_df$SIZE_LABEL[train_df$PRED_SIZE <= 1000] <- 'A)<1KB'
train_df$SIZE_LABEL[train_df$PRED_SIZE > 1000 & train_df$PRED_SIZE <= 5000] <- 'B)1KB-5KB'
train_df$SIZE_LABEL[train_df$PRED_SIZE > 5000 & train_df$PRED_SIZE <= 10000] <- 'C)5KB-10KB'
train_df$SIZE_LABEL[train_df$PRED_SIZE > 10000 & train_df$PRED_SIZE <= 25000] <- 'D)10KB-25KB'
train_df$SIZE_LABEL[train_df$PRED_SIZE > 25000 & train_df$PRED_SIZE <= 50000] <- 'E)25KB-50KB'
train_df$SIZE_LABEL[train_df$PRED_SIZE > 50000 & train_df$PRED_SIZE <= 75000] <- 'F)50KB-75KB'
train_df$SIZE_LABEL[train_df$PRED_SIZE > 75000 & train_df$PRED_SIZE <= 100000] <- 'G)75KB-100KB'
train_df$SIZE_LABEL[train_df$PRED_SIZE > 100000 & train_df$PRED_SIZE <= 250000] <- 'H)100KB-250KB'
train_df$SIZE_LABEL[train_df$PRED_SIZE > 250000 & train_df$PRED_SIZE <= 500000] <- 'I)250KB-500KB'
train_df$SIZE_LABEL[train_df$PRED_SIZE > 500000 & train_df$PRED_SIZE <= 1000000] <- 'J)500KB-1MB'
train_df$SIZE_LABEL[train_df$PRED_SIZE > 1000000 & train_df$PRED_SIZE <= 5000000] <- 'K)1MB-5MB'
train_df$SIZE_LABEL[train_df$PRED_SIZE > 5000000 ] <- 'L)>5MB'

train_df$OV_IND <- gsub("\\.", 0, train_df$OV_IND)
train_df$LABEL_VAL <- ifelse((as.numeric(train_df$OV_IND) > 0), 1,0)
train_df$OV_IND <- NULL

test_df$SIZE_LABEL[test_df$PRED_SIZE <= 1000] <- 'A)<1KB'
test_df$SIZE_LABEL[test_df$PRED_SIZE > 1000 & test_df$PRED_SIZE <= 5000] <- 'B)1KB-5KB'
test_df$SIZE_LABEL[test_df$PRED_SIZE > 5000 & test_df$PRED_SIZE <= 10000] <- 'C)5KB-10KB'
test_df$SIZE_LABEL[test_df$PRED_SIZE > 10000 & test_df$PRED_SIZE <= 25000] <- 'D)10KB-25KB'
test_df$SIZE_LABEL[test_df$PRED_SIZE > 25000 & test_df$PRED_SIZE <= 50000] <- 'E)25KB-50KB'
test_df$SIZE_LABEL[test_df$PRED_SIZE > 50000 & test_df$PRED_SIZE <= 75000] <- 'F)50KB-75KB'
test_df$SIZE_LABEL[test_df$PRED_SIZE > 75000 & test_df$PRED_SIZE <= 100000] <- 'G)75KB-100KB'
test_df$SIZE_LABEL[test_df$PRED_SIZE > 100000 & test_df$PRED_SIZE <= 250000] <- 'H)100KB-250KB'
test_df$SIZE_LABEL[test_df$PRED_SIZE > 250000 & test_df$PRED_SIZE <= 500000] <- 'I)250KB-500KB'
test_df$SIZE_LABEL[test_df$PRED_SIZE > 500000 & test_df$PRED_SIZE <= 1000000] <- 'J)500KB-1MB'
test_df$SIZE_LABEL[test_df$PRED_SIZE > 1000000 & test_df$PRED_SIZE <= 5000000] <- 'K)1MB-5MB'
test_df$SIZE_LABEL[test_df$PRED_SIZE > 5000000 ] <- 'L)>5MB'

#################################
# STEP 3: Write the output file #
#################################
setwd(input_dir)
write.table(train_df, file=train_out_file, quote=FALSE, row.names=FALSE, sep='\t')
write.table(test_df, file=test_out_file, quote=FALSE, row.names=FALSE, sep='\t')
```
