## Supplemental_Code for "A machine-learning approach for accurate detection of copy-number variants from exome sequencing": generate_interval_combo.r

```
###############################################################################
# Script : generate_interval_combo.r                                          #
# Author : Vijay Kumar                                                        # 
# Date   : 4/5/2019                                                           #
#                                                                             #
# This script implements the algorithm that identifies CNVs that overlap and  #
# groups them together. Once the calls are grouped, it generates all possible #
# start and end coordinates based on the CNV coordinates within each group.   #
#                                                                             #
# (c) 2019 - Vijay Kumar                                                      #
# Licenced under the GNU General Public License 3.0.                          #
###############################################################################
library(sqldf)

##########################################################
# STEP 1: Parse the arguements passed from the bash script
##########################################################
args = commandArgs(trailingOnly=TRUE)
data_dir = args[1]
input_file_name = args[2]
output_file_w_grps = args[3]
output_file_grouped_preds = args[4]
output_file_name_2 = args[5]
caller_count = args[6]

caller_list <- character(caller_count)
for (caller_number in 1:caller_count){
    caller_list[caller_number] <- args[6 + caller_number]
} 

setwd(data_dir)

sample_all_preds_df <- as.data.frame(read.table(input_file_name))

num_obs <- nrow(sample_all_preds_df)
num_cols <- ncol(sample_all_preds_df)

sample_all_preds_df[,(num_cols + 1)] <- NA

dyn_col_names <- character(0)
for (caller_name in caller_list) {
    dyn_col_names <- append(dyn_col_names, caller_name)
}

colnames(sample_all_preds_df) <- c('CHR', 'START', 'END', 'CNV_TYPE', 'SAMPLE',
             'CALLER', 'PRED_SIZE',  dyn_col_names, 'OVERLAP_COUNT', 'GRP_NAME')

################################################################################
# STEP 2: Sort the CNVs based on their chromosome, start and end position in   #
#         order to facilitate the subsequent alggorithm to read CNVs one after #
#         another to accurately identify overlapping calls                     #
################################################################################
dyn_query_string_1 <- character(0)
dyn_query_string_2 <- character(0)
dyn_query_string_3 <- character(0)

for (caller_name in caller_list) {
    dyn_query_string_1 <- paste(dyn_query_string_1, caller_name, sep =", ") 
    dyn_query_string_2 <- paste(dyn_query_string_2, paste0(" sum(", caller_name, ") 
                                                     as ", caller_name), sep =", ")
    dyn_query_string_3 <- paste(dyn_query_string_3, paste0(" a.", caller_name), sep =", ")
}

sql_query_1 <- paste0("select CHR, START, END, CNV_TYPE, SAMPLE, CALLER, PRED_SIZE ", 
                      dyn_query_string_1, ", OVERLAP_COUNT, GRP_NAME",
                    " from sample_all_preds_df",
                    " order by CHR, length(START), cast(substr(START,1,3) as int), 
                      cast(substr(END,1,5) as int), CAST(OVERLAP_COUNT as int) desc")

sample_all_preds_df <- sqldf(sql_query_1)

################################################################################
# STEP 3: Read the CNVs within the sample and identify the ones that overlap   #
#   by comparing the coordinates of THE current CNV with the next in the list  # 
################################################################################
row_num = 1
group_counter = 1
group_list <- c()
group_list <- append(group_list,row_num)
row_num = row_num + 1
while (row_num <= num_obs) {
    prev_row_num = row_num - 1
    curr_chr = sample_all_preds_df[row_num,"CHR"]
    prev_chr = sample_all_preds_df[prev_row_num,"CHR"]
    curr_start = sample_all_preds_df[row_num,"START"]
    prev_end = sample_all_preds_df[prev_row_num,"END"]
    if (curr_chr == prev_chr) {
      if (curr_start < prev_end) {
        row_num = row_num + 1
      }
      else if (curr_start > prev_end){
        group_list <- append(group_list,row_num)
        row_num = row_num + 1
      }
    }
    else if (curr_chr != prev_chr){
      group_list <- append(group_list,row_num)
      row_num = row_num + 1
    }
}

###############################################################
# STEP 4: Loop through each CNV and assign them group numbers #
###############################################################
row_num = 1
num_groups <- length(group_list) + 1
group_counter = 0
while (row_num <= num_obs) {
  if (row_num %in% group_list){
    group_counter = group_counter + 1
    sample_all_preds_df[row_num,"GRP_NAME"] <- paste0('G',group_counter)
  }
  else{
    sample_all_preds_df[row_num,"GRP_NAME"] <- paste0('G',group_counter)
  }
  row_num = row_num + 1
}

valid_sample_all_preds_df <- sample_all_preds_df[sample_all_preds_df$CHR %in% seq(1:22), ]

sql_query_2 <- paste0("select CHR, min(START) as START, max(END) as END, CNV_TYPE, SAMPLE, 
                              max(END) - min(START) as PRED_SIZE ", dyn_query_string_2, 
                              ", max(OVERLAP_COUNT) as OVERLAP_COUNT, GRP_NAME",
                    " from valid_sample_all_preds_df",
                    " group by CHR, CNV_TYPE, SAMPLE, GRP_NAME")

grouped_all_preds_df <- sqldf(sql_query_2)

write.table(valid_sample_all_preds_df, output_file_w_grps, quote=FALSE, 
            row.names=F, col.names=F, sep='\t')
write.table(grouped_all_preds_df, output_file_grouped_preds, quote=FALSE, 
            row.names=F, col.names=F, sep='\t')

#################################################################################
# STEP 5: Generate every possible start and end coordinate for overlapping CNVs.#
# This is accomplished using a self join on the list of CNVs with group names.  #
#################################################################################
sql_query_3 <- paste0("select a.CHR, a.START, b.END, a.CNV_TYPE, a.SAMPLE ", 
                      dyn_query_string_3, " ,a.OVERLAP_COUNT, a.GRP_NAME",
                      " from sample_all_preds_df a, sample_all_preds_df b",
                      " where a.CHR = b.CHR and a.GRP_NAME = b.GRP_NAME and a.START < b.END")

all_interval_combo <- sqldf(sql_query_3)

############################################################################
# STEP 6: Write the output file with the list of all interval combinations #
############################################################################
write.table(all_interval_combo, output_file_name_2, quote=FALSE, 
                             row.names=F, col.names=F, sep='\t')
```
