## Supplemental_Code for "A machine-learning approach for accurate detection of copy-number variants from exome sequencing": identify_targets_of_interest.r

```
################################################################################
# Script : identify_targets_of_interest.r                                      #
# Author : Vijay Kumar                                                         #
# Date   : 4/5/2019                                                            #
#                                                                              #
# This script identifies the probes in the left and right flanking regions of  #
# each predicted CNV to facilitate the calculation of read depth ratio between #
# the predicted and their corresponding flanking regions.                      #
#                                                                              #
# (c) 2019 - Vijay Kumar                                                       #
# Licenced under the GNU General Public License 3.0.                           #
################################################################################
library(sqldf)

##########################################################
# STEP 1: Parse the parameters passed by the bash script #
##########################################################
args = commandArgs(trailingOnly=TRUE)
data_dir = args[1]
input_file_name_1 = args[2]
input_file_name_2 = args[3]
output_file_name = args[4]
targ_of_interest_file = args[5]
caller_count = args[6]

caller_list <- character(caller_count)
for (caller_number in 1:caller_count){
    caller_list[caller_number] <- args[6 + caller_number]
} 

##################################################################################
# STEP 2: Identify the targets of interest and extract them from the target file #
##################################################################################
setwd(data_dir)
preds_w_targ_df <- as.data.frame(read.table(input_file_name_1))
targets_file_df <- as.data.frame(read.table(input_file_name_2))

dyn_col_names <- character(0)
for (caller_name in caller_list) {
    dyn_col_names <- append(dyn_col_names, caller_name)
}

colnames(preds_w_targ_df) <- c('CHR','START','END','CNV_TYPE','SAMPLE', dyn_col_names,  'OVERLAP_COUNT', 'GRP_NAME',
                               'CHR_TRG','START_TARG','END_TARG','TARG_SIZE','UNIQ_ID_TARGETS','TARG_OV_SIZE')
colnames(targets_file_df) <- c("CHR", "START", "END", "TARG_SIZE", "UNIQ_TARG_ID")

dyn_query_string <- character(0)
for (caller_name in caller_list) {
    dyn_query_string <- paste0(dyn_query_string, caller_name, sep =", ") 
}

sql_query <- paste0("select CHR, START, END, CNV_TYPE, SAMPLE, ", dyn_query_string, 
                     "OVERLAP_COUNT, GRP_NAME, count(*) as NUM_TARGETS, count(*)/2 as HALF_NUM_TARGETS, ",
                     "CASE WHEN count(*)/2 > 0 THEN min(UNIQ_ID_TARGETS) - count(*)/2 
                           ELSE min(UNIQ_ID_TARGETS) - 1 END as LEFT_FLANK_START, ",
                     "min(UNIQ_ID_TARGETS) - 1 as LEFT_FLANK_END, ", 
                     "min(UNIQ_ID_TARGETS) as START_TARGET, max(UNIQ_ID_TARGETS) as END_TARGET, ",
                     "max(UNIQ_ID_TARGETS) + 1 as RIGHT_FLANK_START, ",
                     "CASE WHEN count(*)/2 > 0 THEN max(UNIQ_ID_TARGETS) + count(*)/2 
                           ELSE max(UNIQ_ID_TARGETS) + 1 END as RIGHT_FLANK_END ",
                     "from preds_w_targ_df ",
                     "GROUP BY CHR, START, END, CNV_TYPE, SAMPLE, ", dyn_query_string, "OVERLAP_COUNT, GRP_NAME ",
                     "ORDER BY cast(trim(substr(GRP_NAME,2,3)) as int), CHR, START, END")

cons_w_all_info <- sqldf(sql_query)

cons_w_all_info$UNIQ_PRED_COMBO_ID <- paste0('C', seq.int(nrow(cons_w_all_info)))

num_obs <- nrow(cons_w_all_info)
num_cols <- ncol(cons_w_all_info)

####################################################################
# STEP 3: Add the list of targets of interest into a separate file #
####################################################################
targets_of_interest <- c()
for (row in 1:num_obs){
  for (target in cons_w_all_info[row,'LEFT_FLANK_START']:cons_w_all_info[row,'RIGHT_FLANK_END']){
    targets_of_interest <- append(targets_of_interest,target)
  }
}
targets_of_interest <- as.data.frame(targets_of_interest)

chosen_targets <- sqldf("select distinct all_targets.* from targets_file_df all_targets, targets_of_interest interest 
                         where all_targets.UNIQ_TARG_ID = interest.targets_of_interest")

##################################
# STEP 4: Write the output files #
##################################
write.table(cons_w_all_info, output_file_name, quote=FALSE, row.names=F, col.names=F, sep='\t')
write.table(chosen_targets, targ_of_interest_file, quote=FALSE, row.names=F, col.names=F, sep='\t')
```
