## Supplemental_Code for "A machine-learning approach for accurate detection of copy-number variants from exome sequencing": measure_rd_stats.r

```
################################################################################
# Script : measure_rd_stats.r                                                  #
# Author : Vijay Kumar                                                         #
# Date   : 4/5/2019                                                            #
#                                                                              #
# This script calculates the average read depth within the predicted regions   #
# and their corresponding flaking regions for each possible CNV start and end  #
# coords. It then selects the endpoints with the highest (Dup) or lowest (Del) #
# average read depth ratio as the most likely breakpoints.                     # 
#                                                                              #
# (c) 2019 - Vijay Kumar                                                       #
# Licenced under the GNU General Public License 3.0.                           #
################################################################################
library(sqldf)

###########################################################
# STEP 1: Parse the parameters passed by the shell script #
###########################################################
args = commandArgs(trailingOnly=TRUE)
calls_dir = args[1]
left_flank_file = args[2]
pred_region_file = args[3]
right_flank_file = args[4]
targets_rd_file = args[5]
output_file = args[6]
caller_count = args[7]

caller_list <- character(caller_count)
for (caller_number in 1:caller_count){
    caller_list[caller_number] <- args[7 + caller_number]
} 

################################################################
# STEP 2: Create dataframes for the three separate regions and #
#         for read depth in exome capture targets              #
################################################################
setwd(calls_dir)
left_flank_df <- as.data.frame(read.table(left_flank_file))
pred_region_df <- as.data.frame(read.table(pred_region_file))
right_flank_df <- as.data.frame(read.table(right_flank_file))
target_rd_df <- as.data.frame(read.table(targets_rd_file))

dyn_col_names <- character(0)
for (caller_name in caller_list) {
    dyn_col_names <- append(dyn_col_names, caller_name)
}

colnames(left_flank_df) <- c("CHR", "START", "END", "CNV_TYPE", "SAMPLE", dyn_col_names, "OVERLAP_COUNT", "GRP_NAME",
                             "NUM_TARGETS", "HALF_NUM_TARGETS", "FIRST_TARGET", "LAST_TARGET","UNIQ_PRED_COMBO_ID" )
colnames(pred_region_df) <- c("CHR", "START", "END", "CNV_TYPE", "SAMPLE", dyn_col_names, "OVERLAP_COUNT", "GRP_NAME",
                              "NUM_TARGETS", "HALF_NUM_TARGETS", "FIRST_TARGET", "LAST_TARGET", "UNIQ_PRED_COMBO_ID")
colnames(right_flank_df) <- c("CHR", "START", "END", "CNV_TYPE", "SAMPLE", dyn_col_names, "OVERLAP_COUNT", "GRP_NAME",
                              "NUM_TARGETS", "HALF_NUM_TARGETS", "FIRST_TARGET", "LAST_TARGET", "UNIQ_PRED_COMBO_ID")
colnames(target_rd_df) <- c("CHR", "START", "END", "TARG_SIZE", "UNIQ_TARG_ID", "OV_CHR", "OV_START", "OV_END", 
                            "READ_DEPTH", "NUM_BPS")

flank_df <- rbind(left_flank_df, right_flank_df)

target_rd_df$READ_DEPTH <- sub(".", "0", target_rd_df$READ_DEPTH, fixed=TRUE)
target_rd_df <- transform(target_rd_df, TARG_SIZE=as.numeric(TARG_SIZE), UNIQ_TARG_ID=as.numeric(UNIQ_TARG_ID),
                          READ_DEPTH=as.numeric(READ_DEPTH),NUM_BPS=as.numeric(NUM_BPS))
pred_region_df <- transform(pred_region_df, FIRST_TARGET=as.numeric(FIRST_TARGET), LAST_TARGET=as.numeric(LAST_TARGET))
flank_df <- transform(flank_df, FIRST_TARGET=as.numeric(FIRST_TARGET), LAST_TARGET=as.numeric(LAST_TARGET))

#################################################################
# STEP 3: Extract read depth information for both predicted and #
#         flanking regions
#################################################################
pred_region_rd_df <- sqldf("select cnv.*, targ.READ_DEPTH, targ.NUM_BPS, targ.READ_DEPTH * targ.NUM_BPS as RD_VAL
                           from pred_region_df cnv left outer join target_rd_df targ
                           where targ.UNIQ_TARG_ID >= cnv.FIRST_TARGET and targ.UNIQ_TARG_ID <= cnv.LAST_TARGET")

flank_rd_df <- sqldf("select cnv.*, targ.READ_DEPTH, targ.NUM_BPS, targ.READ_DEPTH * targ.NUM_BPS as RD_VAL
                     from flank_df cnv left outer join target_rd_df targ
                     where targ.UNIQ_TARG_ID >= cnv.FIRST_TARGET and targ.UNIQ_TARG_ID <= cnv.LAST_TARGET")

###########################################################################
# STEP 4: Calculate the total read depth within each interval of interest #
###########################################################################
dyn_query_string_1 <- character(0)
dyn_query_string_2 <- character(0)
for (caller_name in caller_list) {
    dyn_query_string_1 <- paste(dyn_query_string_1, caller_name, sep =", ") 
    dyn_query_string_2 <- paste(dyn_query_string_2, paste0(" max(", caller_name, ") as ", caller_name), sep =", ")
}

sql_query_1 <- paste0("select CHR, START, END, CNV_TYPE, SAMPLE ", dyn_query_string_1, 
                      ", OVERLAP_COUNT, NUM_TARGETS, HALF_NUM_TARGETS, GRP_NAME, UNIQ_PRED_COMBO_ID, ", 
                      "sum(NUM_BPS) as PR_NUM_BPS, sum(READ_DEPTH * NUM_BPS) as PR_TOT_RD, ",
                      "round(sum(READ_DEPTH * NUM_BPS)/sum(NUM_BPS),2) as PR_AVG_RD ",
                      "from pred_region_rd_df ", 
                      "group by CHR, START, END, CNV_TYPE, SAMPLE ", dyn_query_string_1, 
                      ", OVERLAP_COUNT, NUM_TARGETS, HALF_NUM_TARGETS, ",
                      "GRP_NAME, UNIQ_PRED_COMBO_ID")

rd_freq_pred <- sqldf(sql_query_1)

sql_query_2 <- paste0("select CHR, START, END, CNV_TYPE, SAMPLE ", dyn_query_string_1, 
                      ", OVERLAP_COUNT, NUM_TARGETS, HALF_NUM_TARGETS, GRP_NAME, UNIQ_PRED_COMBO_ID, ",
                      "sum(NUM_BPS) as FL_NUM_BPS, sum(READ_DEPTH * NUM_BPS) as FL_TOT_RD, ",
                      "round(sum(READ_DEPTH * NUM_BPS)/sum(NUM_BPS),2) as FL_AVG_RD ",
                      "from flank_rd_df ",
                      "group by CHR, START, END, CNV_TYPE, SAMPLE ", dyn_query_string_1, 
                      ", OVERLAP_COUNT, NUM_TARGETS, HALF_NUM_TARGETS, ",
                      "GRP_NAME, UNIQ_PRED_COMBO_ID")

rd_freq_flank <- sqldf(sql_query_2)

############################################################
# STEP 5: Aggregate the read depth information within each #
#         predicted and flanking region.                   #
############################################################
cons_rd_info <- sqldf("select pr.*,  fl.FL_NUM_BPS, fl.FL_AVG_RD, pr.PR_NUM_BPS/fl.FL_NUM_BPS as BP_Ratio, 
                         round(pr.PR_AVG_RD/fl.FL_AVG_RD,2) as RD_Ratio
                       from rd_freq_pred pr left outer join rd_freq_flank fl
                       where pr.GRP_NAME = fl.GRP_NAME and 
                         pr.UNIQ_PRED_COMBO_ID = fl.UNIQ_PRED_COMBO_ID and
                         pr.CHR = fl.CHR")

del_rd_info <- sqldf("select GRP_NAME, min(RD_Ratio) as MIN_RD_RATIO from cons_rd_info
                      where CNV_TYPE = 'DEL'
                      group by GRP_NAME
                      order by cast(trim(substr(GRP_NAME,2,3)) as int)")

sql_query_3 <- paste0("select CHR, MIN(START) as START, MAX(END) as END, CNV_TYPE, SAMPLE ", dyn_query_string_2, 
                      ", MAX(OVERLAP_COUNT) AS OVERLAP_COUNT, cons_rd_info.GRP_NAME, RD_Ratio ",
                      "from cons_rd_info left outer join del_rd_info ",
                      "where cons_rd_info.GRP_NAME = del_rd_info.GRP_NAME and cons_rd_info.RD_Ratio = del_rd_info.MIN_RD_RATIO ",
                      "group by CHR, CNV_TYPE, SAMPLE, cons_rd_info.GRP_NAME ",
                      "order by cast(trim(substr(cons_rd_info.GRP_NAME,2,3)) as int)")

deletions <- sqldf(sql_query_3)

dup_rd_info <- sqldf("select GRP_NAME, max(RD_Ratio) as MIN_RD_RATIO from cons_rd_info
                      where CNV_TYPE = 'DUP'
                      group by GRP_NAME
                      order by cast(trim(substr(GRP_NAME,2,3)) as int)")

sql_query_4 <- paste0("select CHR, MIN(START) as START, MAX(END) as END, CNV_TYPE, SAMPLE ", dyn_query_string_2, 
                      ", MAX(OVERLAP_COUNT) AS OVERLAP_COUNT, cons_rd_info.GRP_NAME, RD_Ratio ",
                      "from cons_rd_info left outer join dup_rd_info ",
                      "where cons_rd_info.GRP_NAME = dup_rd_info.GRP_NAME and cons_rd_info.RD_Ratio = dup_rd_info.MIN_RD_RATIO ",
                      "group by CHR, CNV_TYPE, SAMPLE, cons_rd_info.GRP_NAME ",
                      "order by cast(trim(substr(cons_rd_info.GRP_NAME,2,3)) as int)")

duplications <- sqldf(sql_query_4)

#############################################################
# STEP 6: Write the final set of duplications and deletions #
#############################################################
final_list_of_calls <- rbind(deletions,duplications)

write.table(final_list_of_calls, file=output_file, row.names=F, col.names=F, quote=F, sep='\t')
```
