## Supplemental_Code for "A machine-learning approach for accurate detection of copy-number variants from exome sequencing": merge_init_preds.r

```
#################################################################################
# Author : Vijay Kumar                                                          #
# Date   : 4/5/2019                                                             #
#                                                                               #
# This script reads the outputs of the CNV calling algorithms and formats them  #
# to ensure data consistency.                                                   #
#                                                                               #
# (c) 2019 - Vijay Kumar                                                        #
# Licenced under the GNU General Public License 3.0.                            #
#################################################################################
options(gsubfn.engine = "R")
library(reshape)

args = commandArgs(trailingOnly=TRUE)
input_dir = args[1]
output_file = args[2]

################################
# Copy the files to dataframes #
################################
#Set working directory
setwd(input_dir)
canoes_calls <- as.data.frame(read.csv("canoes_calls.csv", header=TRUE))
xhmm_calls <- as.data.frame(read.table("xhmm_calls.txt", header=TRUE))
codex_calls <- as.data.frame(read.table("codex_calls.txt", header=TRUE))
clamms_calls <- as.data.frame(read.table("clamms_calls.txt"))

samples <- as.data.frame(read.csv("sample_map_canoes.csv"))

########################
# Process Canoes calls #
########################
canoes_calls_temp <- transform(canoes_calls, INTERVAL = colsplit(INTERVAL, split = "\\:", names = c('chr', 'intvl')))
canoes_calls_tr <- transform(canoes_calls_temp, COORD = colsplit(canoes_calls_temp$INTERVAL$intvl, split = "\\-", names = c('start', 'end')))
canoes_calls_all <- merge(canoes_calls_tr, samples, by = "SAMPLE")

canoes_calls_all <- canoes_calls_all[,-3]
canoes_calls_all$CALLER <- "CANOES"
canoes_calls_final <- canoes_calls_all[c(4,10,11,2,12,13)]

######################
# Process XHMM calls #
######################
xhmm_calls_temp <- transform(xhmm_calls, INTERVAL = colsplit(INTERVAL, split = "\\:", names = c('chr', 'intvl')))
xhmm_calls_tr <- transform(xhmm_calls_temp, COORD = colsplit(xhmm_calls_temp$INTERVAL$intvl, split = "\\-", names = c('start', 'end')))
xhmm_calls_tr$CALLER <- "XHMM"
xhmm_calls_final <- xhmm_calls_tr[c(5,16,17,2,1,18)]

#######################
# Process CODEX calls #
#######################
codex_calls$CALLER <- "CODEX"
codex_calls[,3] <- toupper(codex_calls[,3])
codex_calls_final <- codex_calls[c(2,4,5,3,1,14)]

########################
# Process CLAMMS calls #
########################
clamms_calls$CALLER <- "CLAMMS"
clamms_calls_final <- clamms_calls[c(1,2,3,6,5,19)]

colnames(canoes_calls_final) <- c("CHR", "START", "END", "TYPE", "SAMPLE", "CALLER")
colnames(xhmm_calls_final) <- c("CHR", "START", "END", "TYPE", "SAMPLE", "CALLER")
colnames(codex_calls_final) <- c("CHR", "START", "END", "TYPE", "SAMPLE", "CALLER")
colnames(clamms_calls_final) <- c("CHR", "START", "END", "TYPE", "SAMPLE", "CALLER")

# Add the predictions to the newly created dataframe
cons_calls <- as.data.frame(rbind(canoes_calls_final, codex_calls_final, clamms_calls_final, xhmm_calls_final))

######################################
# Write the consolidated output file #
######################################
sorted_cons_calls <- cons_calls[order(cons_calls$CALLER, cons_calls$SAMPLE, cons_calls$CHR, cons_calls$START, cons_calls$END),]
write.table(sorted_cons_calls,file=output_file, row.names=F, col.names=F, quote=F, sep='\t')
```
