## Supplemental_Code for "A machine-learning approach for accurate detection of copy-number variants from exome sequencing": reshape_caller_overlap_data.r

```
##############################################################################
# Script: reshape_caller_overlap_data.r                                      #
# Author: Vijay Kumar                                                        #
# Date  : 4/5/2019                                                           #
#                                                                            #
# This script reshapes the overlap data obtained in the prior step using     #
# bedtools. Specifically, it transforms row level info to column level.      #
#                                                                            #
# (c) 2019 - Vijay Kumar                                                     #
# Licenced under the GNU General Public License 3.0.                         #
##############################################################################
library(reshape2)
library(sqldf)

############################################
# STEP 1: Parse the arguements to the script 
############################################
args = commandArgs(trailingOnly=TRUE)
input_dir <- args[1]
input_file <- args[2]
output_file <- args[3]
caller_count <- as.integer(args[4])

caller_list <- character(caller_count)
for (caller_number in 1:caller_count){
    caller_list[caller_number] <- args[4 + caller_number]
} 

caller_cols = seq(8,(8 + caller_count - 1))

#################################################################
# STEP 2: Assign column names and create other necessary columns.
#################################################################
setwd(input_dir)
calls_w_ov_df <- read.table(input_file)
colnames(calls_w_ov_df) <- c("CHR", "START", "END", "CNV_TYPE", "SAMPLE", "CALLER", "OV_CALLER", "OV_SIZE")
calls_w_ov_df$PRED_SIZE <- calls_w_ov_df$END - calls_w_ov_df$START
calls_w_ov_df$OV_PROP <- round(calls_w_ov_df$OV_SIZE/calls_w_ov_df$PRED_SIZE,2)
calls_w_ov_df$OV_CALLER <- gsub("\\.", "NO_OVERLAPS", calls_w_ov_df$OV_CALLER)

# Add a dummy unique variable for the dcast step to work since it requires every row to be unique.
# This column will not be processed further.
calls_w_ov_df$DUMMY_UNIQ_ID <- seq.int(nrow(calls_w_ov_df))

calls_w_ov_reshaped_df <- dcast(calls_w_ov_df, CHR + START + END + CNV_TYPE + SAMPLE + CALLER + 
                                PRED_SIZE + DUMMY_UNIQ_ID ~ OV_CALLER, value.var='OV_PROP')
calls_w_ov_reshaped_df[is.na(calls_w_ov_reshaped_df)] <- 0

#########################################################################
# Dynamically construct a string with the list of callers used to predict 
# CNVs prior to aggregating the overlap information.
#########################################################################
dyn_query_string <- character(0)
for (caller_name in caller_list) {
    dyn_query_string <- paste(dyn_query_string, paste0(" sum(", caller_name, ") as ", caller_name), sep =",")
}

sql_query <- paste0("select CHR, START, END, CNV_TYPE, SAMPLE, CALLER, PRED_SIZE", dyn_query_string,
                    " from calls_w_ov_reshaped_df ",
                    "group by CHR, START, END, CNV_TYPE, SAMPLE, CALLER, PRED_SIZE ",
                    "order by CHR, START, END, CNV_TYPE, SAMPLE, CALLER")

calls_w_ov_reshaped_grouped_df <- sqldf(sql_query)
calls_w_ov_reshaped_grouped_df$NUM_OVERLAPS <- rowSums(calls_w_ov_reshaped_grouped_df[,(caller_cols)]!=0)

###############################
# STEP 3: Write the output file
###############################
write.table(calls_w_ov_reshaped_grouped_df, file=output_file, row.names=F, col.names=F, quote=F, sep='\t')
```
