## Supplemental_Code for "A machine-learning approach for accurate detection of copy-number variants from exome sequencing": xhmm_call_CNVs.sh

```
#!/bin/bash
############################################################################
# Script : xhmm_call_CNVs.sh                                               #
# Author : Vijay Kumar                                                     #
# Date   : 4/5/2019                                                        #
#                                                                          #
# This script is part of the XHMM pipeline. It uses the read depth info    #
# extracted in the script xhmm_extract.sh to predict CNVs.                 #
#                                                                          #
# (c) 2019 - Vijay Kumar                                                   #
# Licenced under the GNU General Public License 3.0.                       #
############################################################################
echo "Job started on `hostname` at `date`"

source TBD/config.params

################################################################################
# STEP 1: Declare variables, directory locations and other required parameters #
################################################################################
if [ -f ${DATA_XHMM_DIR}xcnv ]; 
then
rm ${DATA_XHMM_DIR}xcnv
fi
touch ${DATA_XHMM_DIR}xcnv

##################################################################
# STEP 2: Combine all GATK depth-of-coverage outputs into one file
##################################################################
# --GATKdepthsList is a list of all *.list.DATA.sample_interval_summary files to be combined
ls ${DATA_XHMM_DIR} | grep GATK_OUT.sample_interval_summary  | awk -v path="${DATA_XHMM_DIR}" '{print path$0}' \
                     > ${DATA_XHMM_DIR}read_depth_file_list

${DOCKER_COMMAND}${XHMM_DIR}xhmm --mergeGATKdepths -o ${DATA_XHMM_DIR}combined_RD.txt \
                                 --GATKdepthsList ${DATA_XHMM_DIR}read_depth_file_list

##############################
# STEP 3: Calculate GC Content
##############################
${DOCKER_COMMAND}java -Xmx3072m -jar ${GATK_SW_DIR}GenomeAnalysisTK.jar -T GCContentByInterval \
                      -L ${TARGET_PROBES} -R ${REF_GENOME} -o ${DATA_XHMM_DIR}GC.txt
cat ${DATA_XHMM_DIR}GC.txt | awk '{if ($2 < 0.1 || $2 > 0.9) print $1}' > ${DATA_XHMM_DIR}extreme_gc_targets.txt

#########################################################################
# STEP 4: Run PLINK/Seq to filter GATK targets with repeat-maked bases
#########################################################################
echo -e "#CHR\tBP1\tBP2\tID" > ${DATA_XHMM_DIR}targets_no_chr.reg
awk '{print $0"\t"NR}' ${TARGET_PROBES} >> ${DATA_XHMM_DIR}targets_no_chr.reg

${DOCKER_COMMAND}${PLINK_DIR}pseq . loc-load --locdb ${DATA_XHMM_DIR}targets.LOCDB \
                                             --file ${DATA_XHMM_DIR}targets_no_chr.reg --group targets \
                                             --out ${DATA_XHMM_DIR}targets.LOCDB.loc-load

#####################################
# STEP 5: Identify complexity of loci
#####################################
${DOCKER_COMMAND}${PLINK_DIR}pseq . loc-stats --locdb ${DATA_XHMM_DIR}targets.LOCDB --group targets \
                                              --seqdb ${PLINK_DIR}seqdb.hg19 \
                                        | awk '{if (NR > 1) print $_}' \
                                        | sort -k1 -g | awk '{print $10}' \
                                        | paste ${DATA_XHMM_DIR}targets_no_chr.reg - \
                                        > ${DATA_XHMM_DIR}locus_complexity.txt

cat ${DATA_XHMM_DIR}locus_complexity.txt | awk '{if ($4 > 0.25) print $1 ":" $2 "-" $3}' \
                             > ${DATA_XHMM_DIR}low_complexity_targets.txt

#################################################################
# STEP 6: Filter samples by GC content/low complexity, and then # 
          find mean-center of targets. Use default parameters.  #
#################################################################
${DOCKER_COMMAND}${XHMM_DIR}xhmm --matrix -r ${DATA_XHMM_DIR}combined_RD.txt --centerData --centerType \
                                   target -o ${DATA_XHMM_DIR}centered_RD.txt --outputExcludedTargets \
                                             ${DATA_XHMM_DIR}filt_targets.txt --outputExcludedSamples \
                                             ${DATA_XHMM_DIR}filt_samples.txt --excludeTargets \
                                             ${DATA_XHMM_DIR}extreme_gc_targets.txt --excludeTargets \
                                             ${DATA_XHMM_DIR}low_complexity_targets.txt --minTargetSize 10 \
                                             --maxTargetSize 10000 --minMeanTargetRD 10 \
                                             --maxMeanTargetRD 500 --minMeanSampleRD 25 \
                                             --maxMeanSampleRD 200 --maxSdSampleRD 150

###########################################################################
# STEP 7: Run PCA on the mean-centered data & normalize based on PCA values
###########################################################################
${DOCKER_COMMAND}${XHMM_DIR}xhmm --PCA -r ${DATA_XHMM_DIR}centered_RD.txt \
                                 --PCAfiles ${DATA_XHMM_DIR}PCA_output.txt

#Normalize data based on PCA values
${DOCKER_COMMAND}${XHMM_DIR}xhmm --normalize -r ${DATA_XHMM_DIR}centered_RD.txt \
                                 --PCAfiles ${DATA_XHMM_DIR}PCA_output.txt \
                                 --normalizeOutput ${DATA_XHMM_DIR}PCA_normalized.txt \
                                 --PCnormalizeMethod PVE_mean --PVE_mean_factor 0.7

############################################################################
# STEP 8: Filter and calculate z-scores for targets from PCA-normalized data
############################################################################
${DOCKER_COMMAND}${XHMM_DIR}xhmm --matrix -r ${DATA_XHMM_DIR}PCA_normalized.txt \
                                 --centerData --centerType sample --zScoreData \
                                 -o ${DATA_XHMM_DIR}PCA_zscores.txt \
                                 --outputExcludedTargets ${DATA_XHMM_DIR}PCA_z_filt_targets.txt \
                                 --outputExcludedSamples ${DATA_XHMM_DIR}PCA_z_filt_samp.txt \
                                 --maxSdTargetRD 30

#Filter original read-depth data against normalized data
${DOCKER_COMMAND}${XHMM_DIR}xhmm --matrix -r ${DATA_XHMM_DIR}combined_RD.txt \
                                 --excludeTargets ${DATA_XHMM_DIR}filt_targets.txt \
                                 --excludeTargets ${DATA_XHMM_DIR}PCA_z_filt_targets.txt \
                                 --excludeSamples ${DATA_XHMM_DIR}PCA_z_filt_samp.txt \
                                  -o ${DATA_XHMM_DIR}same_filtered_RD.txt

######################################
# STEP 9: Call CNVs on normalized data
######################################
${DOCKER_COMMAND}${XHMM_DIR}xhmm --discover -p ${XHMM_DIR}params.txt -r ${DATA_XHMM_DIR}PCA_zscores.txt \
                                  -R ${DATA_XHMM_DIR}same_filtered_RD.txt -c ${DATA_XHMM_DIR}xcnv \
                                  -a ${DATA_XHMM_DIR}aux_xcnv -s ${DATA_XHMM_DIR}

cp ${DATA_XHMM_DIR}'xcnv' ${DATA_DIR}'xhmm_calls.txt'

echo "Job ended on `hostname` at `date`"
```
