## Supplemental_Code for "A machine-learning approach for accurate detection of copy-number variants from exome sequencing": xhmm_extract.sh

```
#!/bin/bash
########################################################################
# Script : xhmm_extract.sh                                             #
# Author : Vijay Kumar                                                 #
# Date   : 4/5/2019                                                    #
#                                                                      # 
# This script is part of the XHMM pipeline that extracts read depth    #
# information for each sample. This read depth information is further  #
# used in the xhmm_call_CNVs.sh script to predict CNVs.                #
#                                                                      #
# IMPORTANT NOTE: The for loop that processes the samples can be       #
# easily modified to run in parallel on local clusters or on other     #
# cloud infrastructures.                                               #
#                                                                      #
# (c) 2019 - Vijay Kumar                                               #
# Licenced under the GNU General Public License 3.0.                   #
########################################################################
echo "Job started on `hostname` at `date`"

source TBD/config.params

################################################################################
# STEP 0: Declare variables, directory locations and other required parameters #
################################################################################
for bam_file in `cat ${BAM_FILE_LIST_W_PATH}`;
do

sample_name=`echo ${bam_file} | rev | cut -f1 -d/ | rev`

#############################################################################
# STEP 1: Generate separate scripts to calculate read counts for each sample
#############################################################################
${DOCKER_COMMAND}java -Xmx3072m -Djava.io.tmpdir=${DATA_LOGS_DIR}xhmm -jar ${GATK_SW_DIR}GenomeAnalysisTK.jar \
                      -T DepthOfCoverage -I ${bam_file} -L ${TARGET_PROBES} -R ${REF_GENOME} -dt BY_SAMPLE \
                      -dcov 5000 -l INFO --omitDepthOutputAtEachBase --omitLocusTable --minBaseQuality 0 \
                      --minMappingQuality 20 --start 1 --stop 5000 --nBins 200 --includeRefNSites \
                      --countType COUNT_FRAGMENTS -o ${DATA_XHMM_DIR}${sample_name}_GATK_OUT

done

echo "Job ended on `hostname` at `date`"
```
